# Transcriptomic analysis of chronic chikungunya in the Reunionese CHIKGene cohort uncovers a shift in gene expression more than 10 years after infection

**DOI:** 10.1101/2025.01.20.633311

**Authors:** Patrick Gérardin, Raissa Medina-Santos, Sigrid Le Clerc, Léa Bruneau, Adrien Maillot, Taoufik Labib, Myriam Rahmouni, Jean-Louis Spadoni, Jean-Philippe Meyniel, Clémence Cornet, Cécile Lefebvre, Nora El Jahrani, Jakub Savara, Mano Joseph Mathew, Christine Fontaine, Christine Payet, Nathalie Ah-You, Cécile Chabert, Corinne Mussard, Sylvaine Porcherat, Samir Medjane, Josselin Noirel, Catherine Marimoutou, Hakim Hocini, Jean-François Zagury

## Abstract

**Aim:** In 2005-2006, a chikungunya epidemic of unprecedented magnitude hit Reunion Island, which raised a public health concern through the substantial proportions of long-lasting manifestations. To understand the pathophysiology underlying chronic chikungunya (CC), we designed the CHIKGene cohort study and collected blood samples from 133 subjects diagnosed with CC and from 86 control individuals that had recovered within 3 months, 12-to-15 years after exposure.

**Methods:** We conducted bulk RNAseq analysis on peripheral blood mononuclear cells to find differentially expressed genes (DEGs), gene set enrichment analysis (GSEA) and gene ontologies to uncover top-level enriched terms associated with DEGs, and weighted gene correlation network analysis (WGCNA) to elucidate underlying cellular processes.

**Results:** Among 1549 DEGs, gene expression analysis identified 10 top genes including *NR4A2* and *TRIM58* (upregulated in CC), *IGHG3* and *IGHV3-49* (downregulated in CC) linked to immune regulation, *OSBP2* (upregulated in CC) and *SEMA6B* (downregulated in CC) linked to neuronal homeostasis and axon guidance, respectively. GSEA and WGCNA unveiled cellular processes such as "Metabolism of RNA" and "Cell Cycle”.

**Conclusions:** This study uncovers a shift in gene expression of CC subjects. *IGHG3* and *IGHV3-49* gene shut-offs spotlight the importance of neutralizing antibodies against chikungunya virus in the progression to chronic disease. Human diseases associations highlight connections to rheumatoid arthritis, nervous and cardiac systems. GSEA and WGCNA bounce the hypotheses of a persistent viral reservoir or an increased susceptibility to RNA viral pathogens with new onset infections. Together, our findings might offer potential targets for therapeutic options aimed at alleviating chronic chikungunya.

## 1. Introduction

Since its re-emergence in 2004 and global spread over the last two decades, Chikungunya virus (CHIKV), an Aedes mosquito-borne alphavirus of *Togaviridae* family, has become a public health concern through its propension to cause a huge disease burden with chronic disabilities, substantial economic losses [1], and the potential for increased mortality [2].

Most CHIKV infections present as an acute self-limiting illness mimicking dengue with fever, joint pain and rash as cardinal symptoms [1]. Systematic reviews and meta-analyses found prevalence of CHIKV-related persistent disability to be within 35% to 58% with some genotype-linked differences [3–6], while the prevalence of fully asymptomatic infections estimate to be within a very loose range of 3% to 82%, conditioned to the force of infection with inverse correlation [7]. Chikungunya chronic manifestations display a wide of range of symptoms and disabilities including arthralgia (joint pain), arthritis (joint pain with swelling and/or stiffness), muscle ache (*i.e.*, this ensemble sometimes referred to as rheumatic musculoskeletal manifestations), fatigue, neurological, neurocognitive, sensorineural, mood, sleep, digestive and skin disorders [4, 6]. They have been defined by expert consensuses in symptomatic individuals when lasting over three months after the onset of the acute illness [8–11], and all have the potential to impair the quality of life [4, 7]. Pathophysiologically, they are regarded as the long-lasting *sequelae* of the disease [6], or the results of a prolonged inflammation due to the persistence of CHIKV immune debris rather than the persistence of replicative live viruses [1], albeit CHIKV RNA has been detected in myeloid cell sanctuaries from diverse animal models and a single human observation [12, 14].

There is no licensed antiviral treatment and in the acute and subacute stages of the disease, the therapeutic options consist in alleviating pain with analgesics. The therapeutic management of chronic chikungunya (>3 months) is reminiscent of rheumatoid arthritis (RA) [8–11]. One single live-attenuated vaccine candidate (VLA1553) demonstrated efficacy in phase 3 randomised clinical trials and is in the development pipeline for industrial production and immunization in endemic countries and travellers [15, 16].

Despite the progress in chikungunya research, the molecular aetiology of chikungunya chronic manifestations remains poorly understood. In 2005-2006, a high-magnitude epidemic occurred on Reunion Island, where approximately 266,000 individuals (∼33% of the island’s population) contracted the disease, and almost 40% (∼300,000 individuals) were infected, with *Ae albopictus* mosquito identified as the likely vector [17]. This allowed to observe rare forms of the disease and characterise the full spectrum of its manifestations.

In the framework of the CHIKGene cohort study, we took benefit of several pre-existing cohorts that were used for following-up the outbreak impact and sought to decipher with a 12-to-15-year hindsight, the gene expression profile and immune drivers associated with the evolution pattern of the CHIKV infection. In this bulk RNA-seq analysis of the peripheral blood mononuclear cells (PBMCs), we aimed to characterise the gene expression profiles in the group of subjects with different chikungunya chronic syndromes as compared to subjects who have recovered, prior to delve deeper into the potentially different endotypes.

## 2. Methods

### 2.1 Study design, participants, and setting

The CHIKGene cohort consisted of individuals taken from subsets of five historical cohorts: 1-the regionwide population-based cohort issued from the post-epidemic serosurvey (SEROCHIK/TELECHIK); 2-The population-based REDIA (REunion-DIAbète) cohort primarily aimed to assess the prevalence of type-2 diabetes; 3-the RHUMATOCHIK cohort of CHIKV-infected outpatients monitored by rheumatologists; 4-two catch-up hospital-based inception cohorts for which exposure was well established and reliable follow-up data were available.

### 2.2 Ethical statement

Each participant provided written consent to be enrolled in the CHIKGene cohort. Further details on ethics and regulations are given in Supplementary file 1 and ethical approval statement.

### 2.3 Exposure and outcome

Exposure to CHIKV had been defined virologically or serologically with positive RT-PCR and/or positive CHIKV-specific IgM and/or positive CHIKV-specific IgG antibodies as specified at cohort inception or follow-up.

"Chronic chikungunya" (CC) has been defined as a cluster of three persistent syndromes (>3 months): chikungunya rheumatism (CHIK-R), chronic fatigue (CFS)-like syndrome and idiopathic chronic fatigue (ICF). These three clinically relevant subgroups of patients were deemed indicative of CC as they reached aetiologic fractions over 60% in the last-in-date evaluation of phenotypes.

Participants who recovered within 3 months of acute chikungunya illness or who complained of non-specific manifestations were classified as "nonchronic chikungunya" (NCC). Fully asymptomatic subjects were ruled out from this analysis, as done classically for the assessment of chronic chikungunya.

The clinical data and questionnaires were collected and processed in an electronic case report form (Clinsight v6, Ennov). The samples were collected by nurses and sent fresh to biological resource operators in a timely manner for preanalytical conditioning.

### 2.4 Blood sampling, RNA extraction and sequencing

Heparinized whole blood was collected from the participants. PBMCs were isolated using centrifugation with Ficoll (≤ April 15, 2019) or vacutainer Cell Preparation Tube (CPT; > April 15, 2019), and stored at -80℃ until RNA extraction. Total RNA was extracted using the R1018 RNeasy Plus Mini Kit (Qiagen). The quality of the extracted mRNA was assessed using the Bioanalyzer 2100 and the 5067-1513 RNA 6000 Pico Kit (Agilent Technologies). Subsequently, the samples underwent RNA purification using the R1018 ZymoResearch "RNA clean and concentrator-25" as per the manufacturer’s instructions.

Libraries were prepared using the 20020595 TruSeq Stranded mRNA Library Prep Kit (Illumina) according to the manufacturer’s instructions. Sequencing was performed on Illumina HiSeq 2500 sequencer according to the manufacturer’s instructions.

### 2.5 Bioinformatic analysis

#### 2.5.1 Quality assessment and Alignment

The quality of the sequencing reads was assessed using FastQC (v0.11.5) and the results compiled by MultiQC tool (v1.20). Alignment of the reads was performed against the Genome Reference Consortium Human Genome Build 38 (GRCh38) using the Spliced Transcripts Alignment to a Reference Aligner (STAR) (v2.5.2b).

#### 2.5.2 Differential gene expression analysis

For comprehensive differential expression analysis and quality control, we defined "Primary DEGs" (*P*_adj_ <0.05) and "High Confidence DEGs" (*P*_adj_ <0.01) using DESeq2 package (v1.38.3). In addition, in order to select the Top 10 of the strongly DEGs, the genes with an absolute log2 fold change (log2FC > 0.7) and an average of normalized count data across the samples (baseMean) higher than 50 were selected for further characterization in subgroups: CC (chronic chikungunya), NCC (nonchronic chikungunya), females with CC (fCC), females with NCC (fNCC), males with CC (mCC) and males with NCC (mNCC). Boxplots were generated using the *Boxplot* R function to represent the normalized expression of each of these genes.

#### 2.5.3 Pathway enrichment

We conducted a gene set enrichment analysis (GSEA) and searched for associations with human diseases (DisGeNET) using the Metascape portal (https://metascape.org) to uncover the top-level enriched terms associated with both Primary DEGs and High Confidence DEGs. To identify the significant terms, a threshold of q-value <0.05 was applied (log10(q-value) <-1.3).

In another approach, with the full list of genes (*P*_adj_ <0.01) sorted by the absolute values of log2FC, we used the Search Tool for the Retrieval of Interacting Genes (STRING v12) to provide a protein-protein interaction (PPI) enrichment analysis with KEGG, tailored to *Homo Sapiens*.

#### 2.5.4 Weighted gene co-expression network analysis

In the aim to elucidate the underlying cellular processes based on the coordinated co-expression of genes encoding the interacting proteins correlated across all individuals (regardless of disease outcome), we performed a weighted gene co-expression network analysis using identified DEGs (WGCNA, v1.72.5).

We used the normalized and transformed matrix of DEGs counts (*P*_adj_ <0.05) generated with DESeq2. To elucidate the biological relevance of the identified co-expression clusters, we performed an annotation in GSEA using Metascape and DisGeNET tools (q-value <0.05).

The full methods of the bulk RNAseq analysis are detailed in Supplementary file 1 and summarized in Fig. 1.

**Figure 1.**
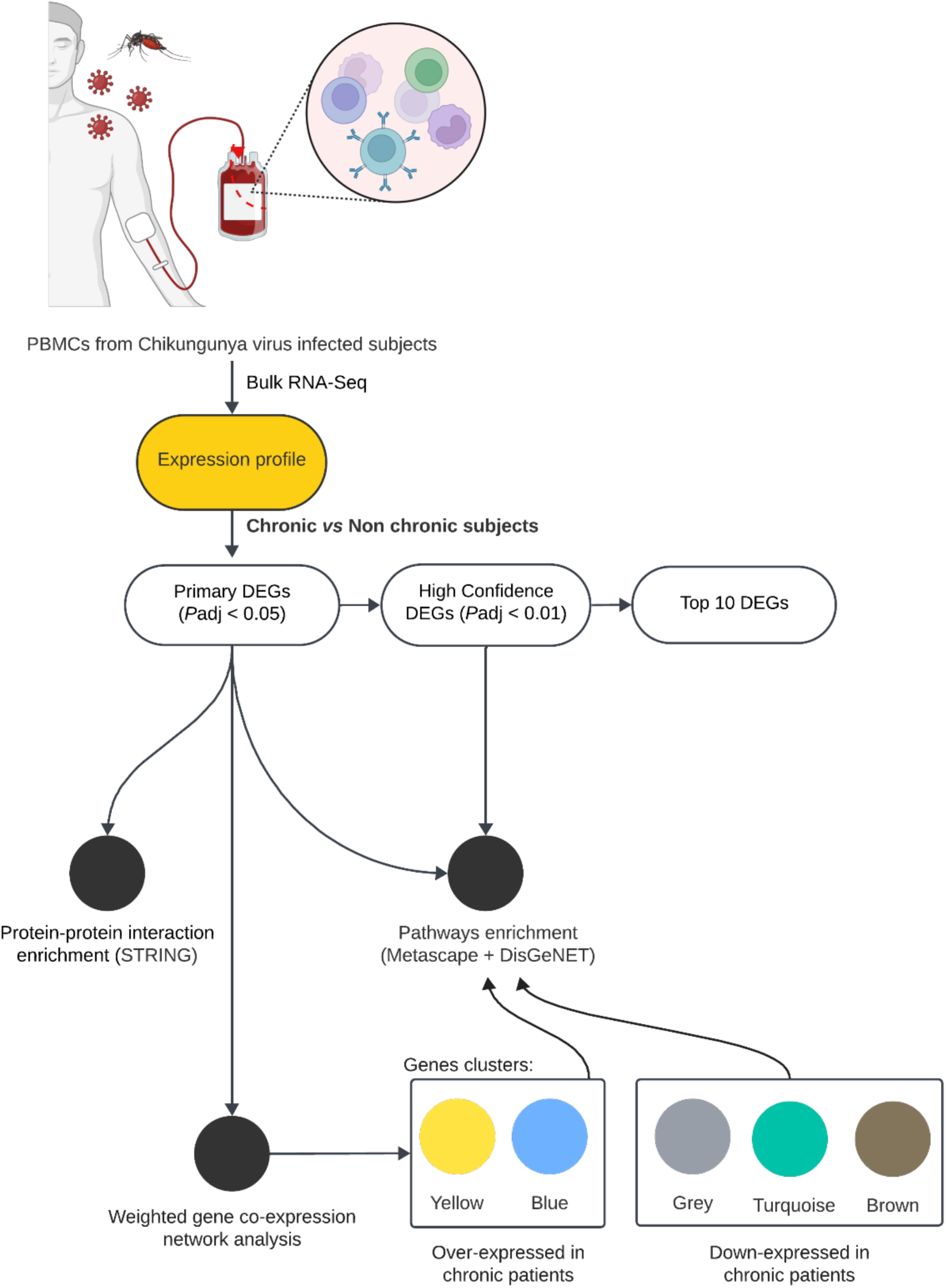
Methods overview. This detailed overview illustrates the different methods used in this bulk RNAseq analysis, from the extraction of PBMCs in CHIKV-exposed subjects. RNA sequencing was performed to identify expression profiles in chronic and nonchronic participants. The resulting list of differentially expressed genes (DEGs), termed Primary DEGs (Padj <0.05), underwent protein-protein interaction enrichment analysis using STRING. Moreover, the Primary DEGs list was employed for pathway enrichment analysis using Metascape. We also subjected the gene clusters resulting from a weighted gene co-expression network analysis to a pathway enrichment in Metascape. High Confidence DEGs (Padj <0.01) were similarly analysed for pathway enrichment using Metascape. Finally, a Top 10 DEGs list (log2FC 0.7 and baseMean >50) was selected for individual gene evaluation and discussion of potential pathomechanisms.

#### 2.5.5 Role of funding source and reporting

The funder of the study had no role in the study design, data collection, analysis, interpretation, or writing of the report. The study used the guidance of STREGA checklist (Supplementary file 2), an extension of STROBE guideline for the reporting of genetic association studies.

## 3. Results

### 3.1 Participants

Of the 609 subjects enrolled in the CHIKGene study cohort, 456 had been infected with the CHIKV on Reunion Island during the 2005-2006 outbreak. Among these, 244 were invited to blood sampling and were offered to participate in the bulk RNAseq analysis from their PBMCs. In this subset, 20 subjects were ruled out due to a fully asymptomatic course since onset of infection, which precluded the study of chronic manifestations and endotypes, and five subjects due to genetic relatedness, resulting in a final count of 219 participants. Among these, 133 (60.7%) had presented CC syndromes and 86 (39.3%) had recovered and were classified as NCC. The study population is presented in Fig. 2. Table 1 provides detailed information about the participants. A detailed interpretation of the cohort is presented in Supplementary file 3.

**Figure 2.**
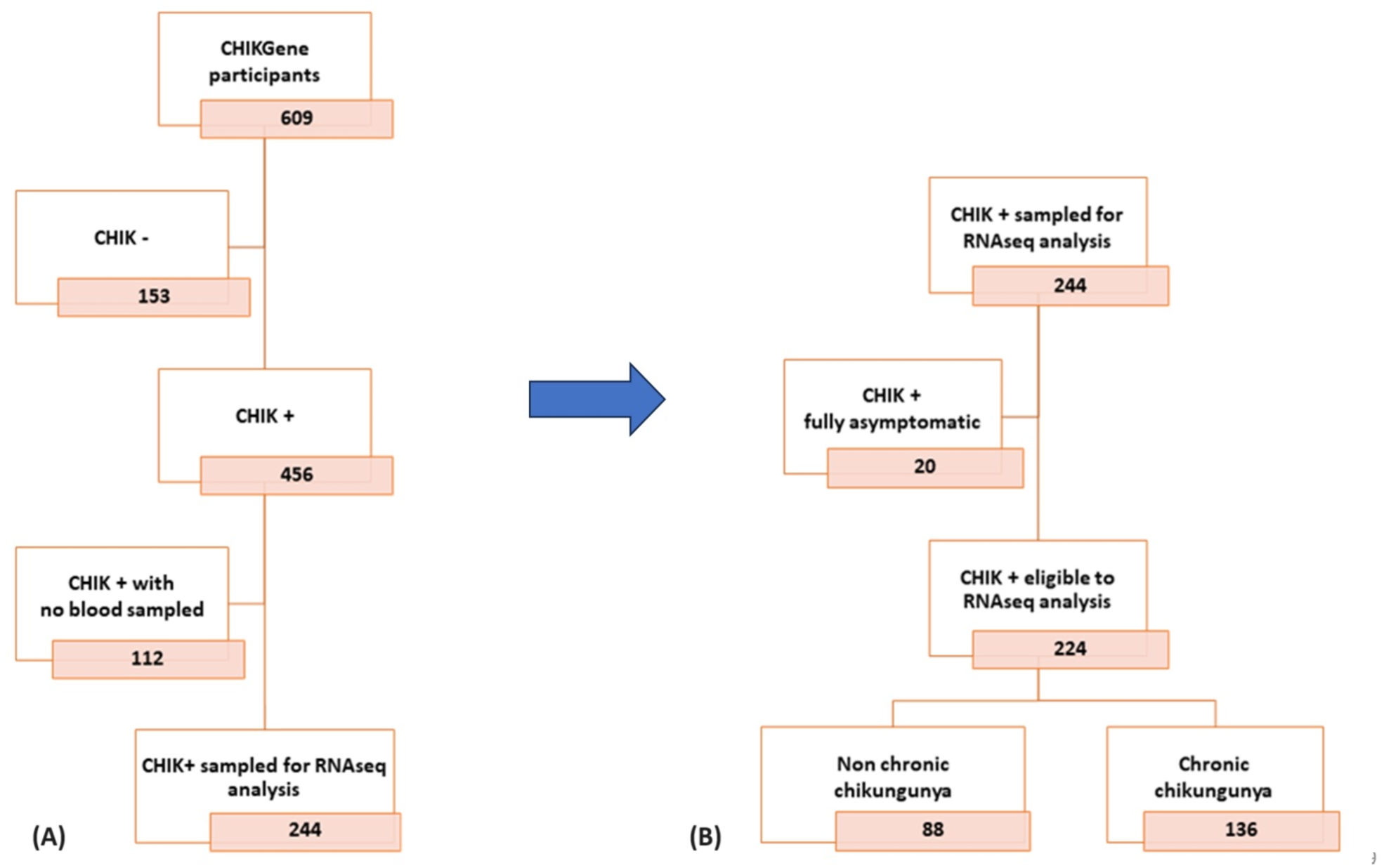
Flow chart of the study population. Selection of study participants from the CHIKGene study population to the subset sampled for PBMCs, showing populations excluded and studied for bulk RNAseq of disease outcomes.

**Table 1.** *C*HIKV-infected participants to bulk RNAseq analysis, CHIKGene cohort study, Reunion, 2018-2020.

| Variables | Total (n=219) |  | Non chronic (n=86) |  | Chronic (n=133) |  | p-value |
| --- | --- | --- | --- | --- | --- | --- | --- |
|  | N | (%) | N | (%) | N | (%) |  |
| **Cohort (group-level variable)** |  |  |  |  |  |  | < 0.001 |
| SEROCHIK/TELECHIK | 70 | (32.0) | 35 | (50.0) | 35 | (50.0) |  |
| REDIA | 39 | (17.8) | 22 | (56.4) | 17 | (43.6) |  |
| RHUMATOCHIK | 29 | (13.2) | 0 | (0.0) | 29 | (100) |  |
| HOSPITAL | 81 | (37.0) | 29 | (35.8) | 52 | (64.2) |  |
| **Individual-level variables** |  |  |  |  |  |  |  |
| **Female sex** | 162 | (74.0) | 56 | (34.6) | 106 | (65.4) | 0.016 |
| Age, years ($\mu$, SD) | 52.5 | (12.6) | 50.5 | (12.8) | 53.8 | (12.4) | 0.063 |
| < 45 | 68 | (31.0) | 32 | (47.1) | 36 | (52.9) | 0.235 |
| 45-64 | 110 | (50.2) | 41 | (37.3) | 69 | (62.7) |  |
| $\geq 65$ | 41 | (18.7) | 13 | (31.7) | 28 | (68.3) | |
| **Smoker \*** | 56 | (25.6) | 26 | (46.4) | 30 | (53.6) | 0.204 |
| **Pregnant at infection onset** | 50 | (22.8) | 24 | (48.0) | 26 | (52.0) | 0.150 |
| **Body mass index, kg/m<sup>2</sup> ($\mu$, SD) (n=170) #** | 26.9 | (5.8) | 26.4 | (5.3) | 27.2 | (6.2) | 0.474 |
| 25-29.9 | 45 | (26.5) | 24 | (53.3) | 21 | (46.7) | 0.285 |
| $\geq 30$ | 50 | (29.4) | 19 | (38.0) | 31 | (62.0) | |
| **Diabetes mellitus** |  |  |  |  |  |  | 0.779 |
| Pre-chikungunya <sup>†</sup> | 24 | (11.0) | 8 | (33.3) | 16 | (66.7) |  |
| Post-chikungunya <sup>‡</sup> | 32 | (14.6) | 12 | (37.5) | 20 | (62.5) |  |
| **Cardiovascular disease** |  |  |  |  |  |  | < 0.001 |
| Pre-chikungunya | 24 | (11.0) | 15 | (62.5) | 9 | (37.5) |  |
| Post-chikungunya | 54 | (24.7) | 10 | (18.5) | 44 | (81.5) |  |
| **Comorbidities** |  |  |  |  |  |  | < 0.001 |
| 1 | 83 | (37.9) | 34 | (41.0) | 49 | (59.0) |  |
| 2 | 73 | (33.3) | 23 | (31.5) | 50 | (68.5) |  |
| $\geq 3$ | 35 | (15.1) | 5 | (15.1) | 28 | (84.9) | |
| **Chronic chikungunya phenotype** |  |  |  |  |  |  |  |
| Chikungunya rheumatism |  |  |  |  | 113 | (85.0) |  |
| Chronic fatigue syndrome like |  |  |  |  | 58 | (43.6) |  |
| Idiopathic chronic fatigue |  |  |  |  | 31 | (23.3) |  |

Data are column (prevalence of the condition) and row percentages (proportion of disease outcome among the exposed to the condition), means ( $\mu$ ) and standard deviations (SD). \* Past or current smoking habit. # 50 subjects with the data missing. <sup>†</sup> The condition was diagnosed prior chikungunya onset. <sup>‡</sup> The condition was diagnosed after chikungunya onset. <sup>¶</sup> Osteoarthritis, diabetes mellitus, cardiovascular disease, asthma, chronic bronchitis, chronic kidney disease, cancer.

### 3.2 Reads quality, principal component analysis and covariate identification

The quality assessment of RNA sequences is displayed as a multiQC quality report in Supplementary file 4. To identify some putative batch effects and check the reliability of sampling thereof, a PCA was conducted (S1 fig.). PCA analysis specifications details are presented in Supplementary file 3.

### 3.3 Bioinformatic analysis

DEGs first-step comparisons between CC and NCC groups allowed us to identify 1549 genes (Primary DEGs -*P*_adj_ <0.05) and 169 high confidence DEGs (*P*_adj_ <0.01) (Fig. 3; Supplementary file 5).

**Figure 3.**
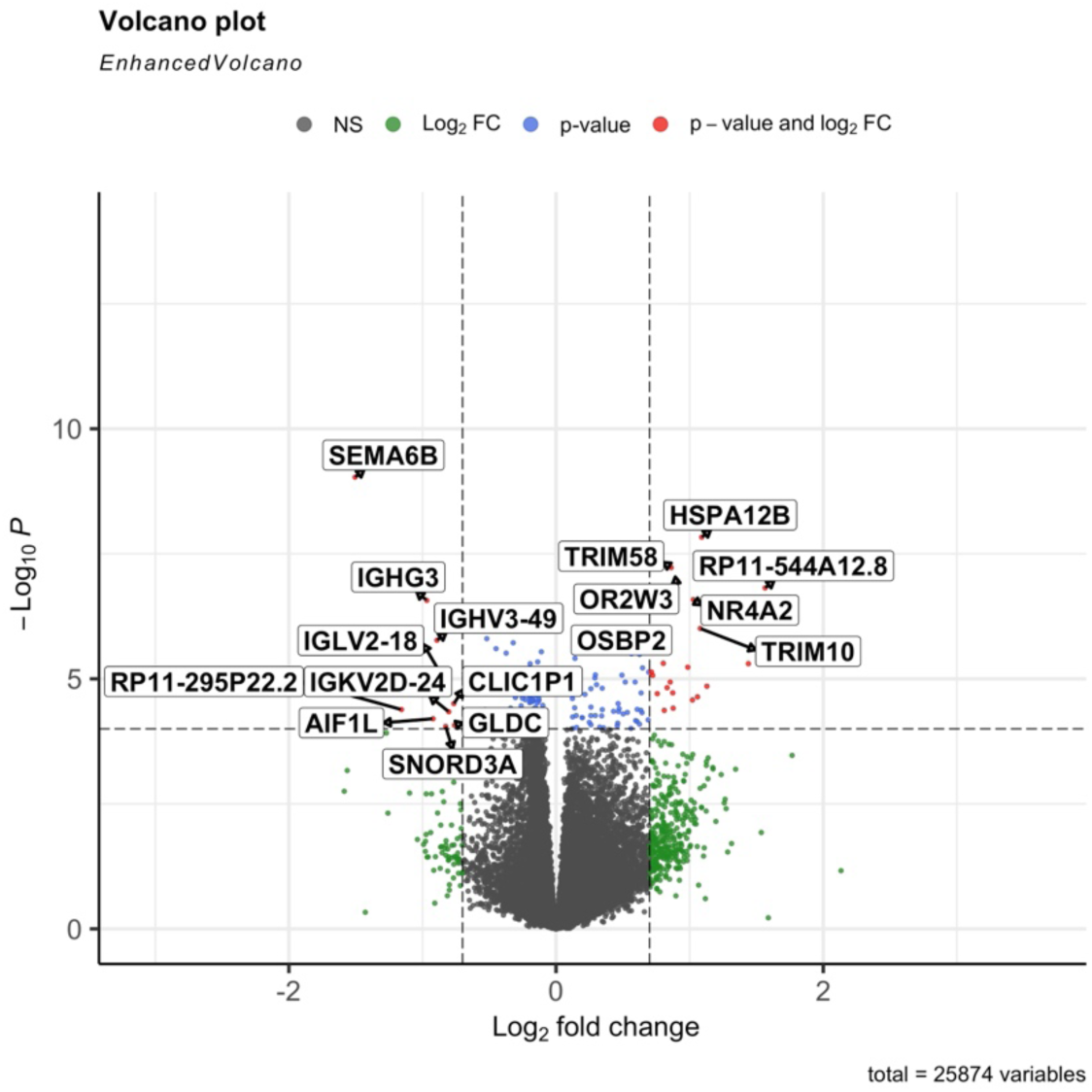
Volcano plot from all 1549 DEGs (*P*_adj_ <0.05) differentiating chronic *vs* nonchronic individuals at first step of differential expression analysis, plotted by *EnhancedVolcano* function. Up and down-regulated DEGs differentiating chronic and nonchronic individuals are plotted vertically in the volcano plots by *EnhancedVolcano* function on the right and on the left on either side of 0 x axis, respectively, and the intensity of changes is plotted horizontally on the y axis.

#### 3.3.1 Top genes identification

Among those DEGs, we selected the Top 10 ranked, based on *P*_adj_ with a log2FC >0.7 and a baseMean >50. Seven of those genes were more expressed in the CC group compared to the NCC group. The expression of the top 10 ranked genes was analysed across different subgroups: CC, NCC, fCC, fNCC, mCC and mNCC. The first set of genes boxplots presents distinct expression patterns in the CC group: *NR4A2*, *TMOD1*, *OSBP2*, *SPTB, TRIM58*, *OR2W3* and *EGR3* (Fig. 4). Five of these genes were upregulated in the CC group regardless of the sex of the individuals. By contrast, the gene *NR4A2* exhibited a sex-biased opposite direction, with a lower expression among mCC than in mNCC subgroup, while it was more strongly expressed within the fCC compared to the fNCC subgroup. The gene *OSBP2*, presented a higher expression in mCC compared to mNCC, but a similar expression in fCC compared to fNCC (Fig. 4).

**Figure 4.**
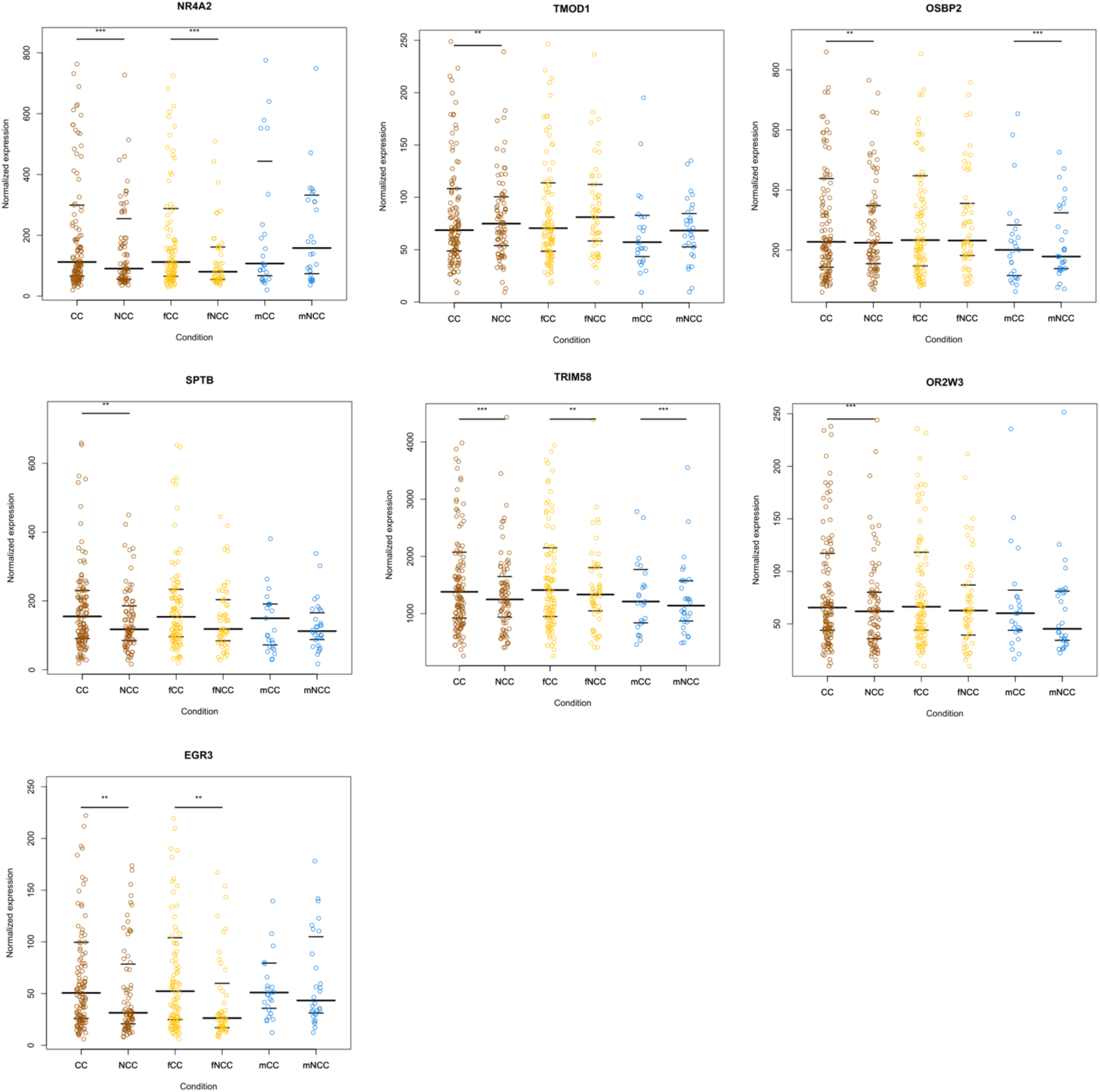
Boxplot of the upregulated genes in chronic vs non chronic individuals. 136 chronic chikungunya (CC) individuals, 88 nonchronic chikungunya (NCC) individuals, 106 CC female individuals, 56 NC female individuals, 27 CC male individuals and 30 NCC male individuals, plotted by R *Boxplot* function. Results are presented as normalized expression of the obtained values, with significance * p<0.05; ** p<0.01 and *** p<0.001.

On the other hand, three genes were more expressed in the NCC group compared to the CC group: *IGHG3*, *IGHV3-49* and *SEMA6B* (Fig 5). Both *IGHG3* and *IGHV3-49* were more expressed among NCC individuals (or downregulated in CC individuals), regardless of sex. In contrast, *SEMA6B* showed a higher expression within mNCC subgroup and a lower expression in fNCC subgroup.

**Figure 5.**
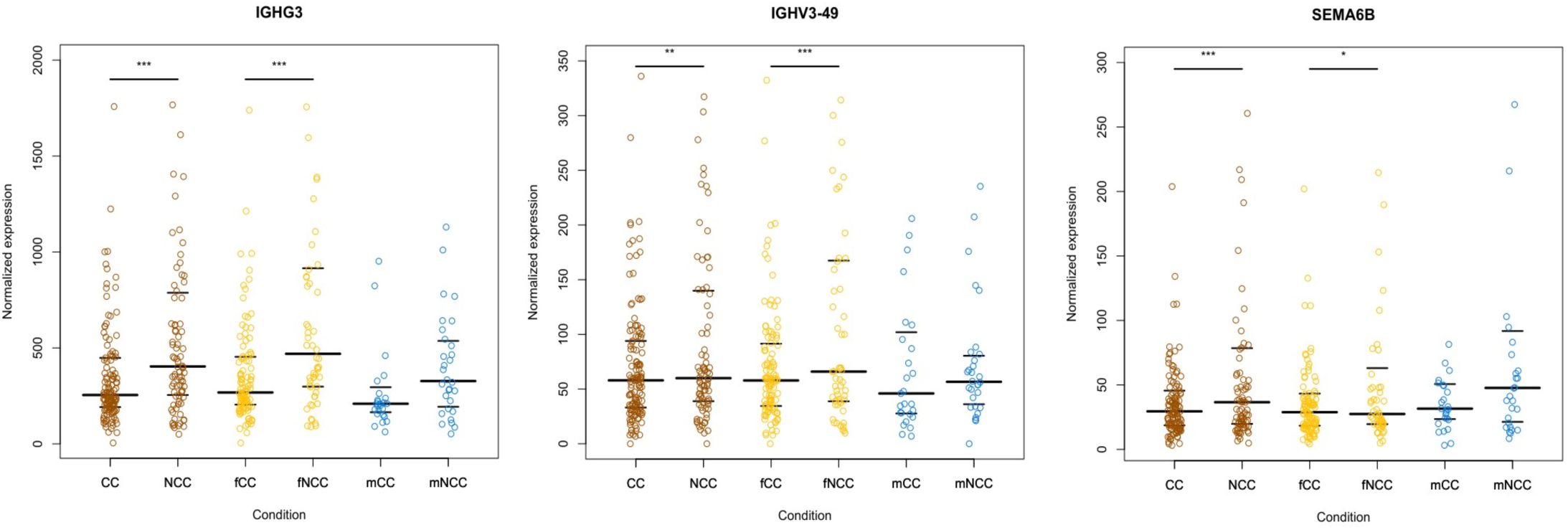
Boxplot of the downregulated genes in chronic vs non chronic individuals. 136 chronic chikungunya (CC) individuals, 88 nonchronic chikungunya (NCC) individuals, 106 CC female individuals, 56 NC female individuals, 27 CC male individuals and 30 NCC male individuals, plotted by R *Boxplot* function. Results are presented as normalized expression of the obtained values, with significance * p<0.05; ** p<0.01 and *** p<0.01.

#### 3.3.2 Pathway enrichment

The top-level 20 from the Metascape GSEA and the top-level 20 enriched and significant terms from the DisGeNET GSEA are displayed in Fig. 6.A and Fig. 6.B, respectively.

**Figure 6.**
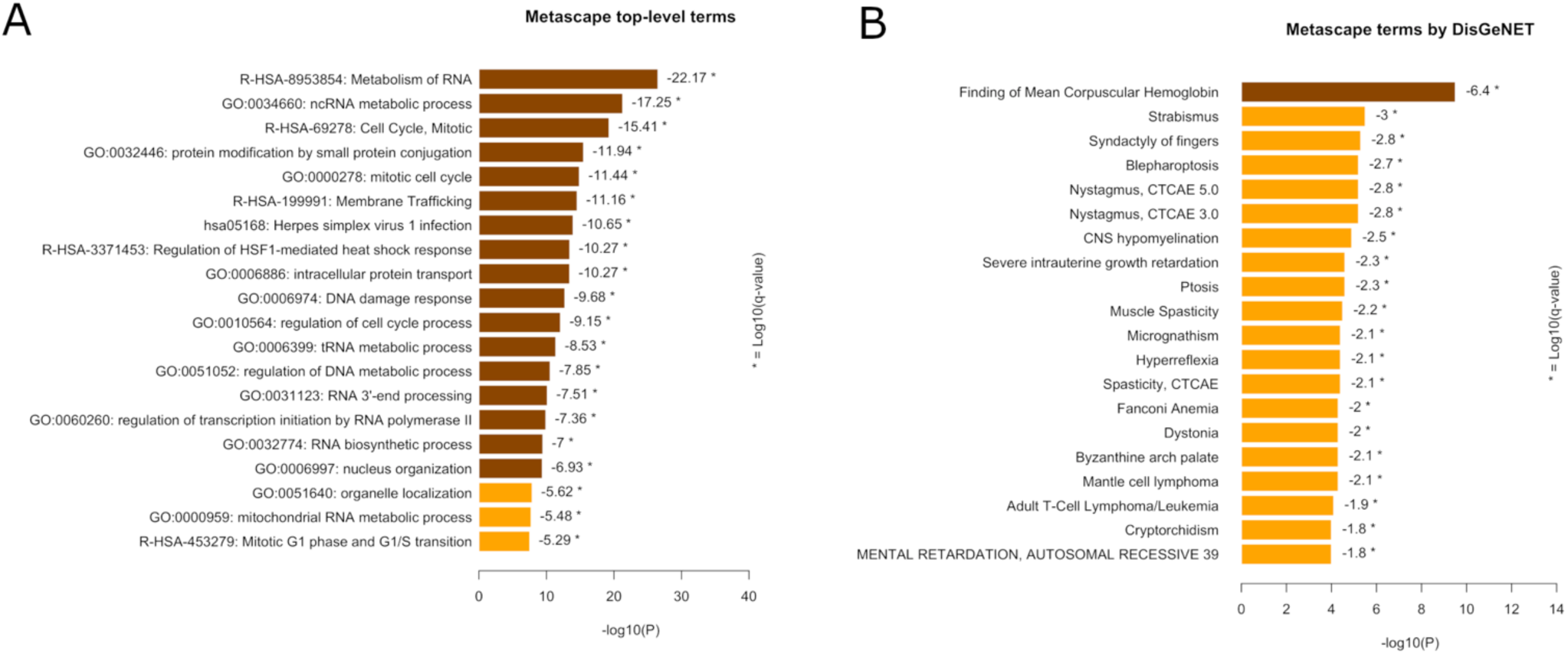
Pathways analysis using Metascape. (A) Bar graph of the 20 top-level enriched terms across input gene list, coloured by decremental p-values and (B) Bar graph of the 20 top-level enriched terms in the DisGeNET, coloured by decremental p-values, using complete DEG list (*P*_adj_ <0.05). Significance of the terms are marked with * at the threshold of the log of q-value from the false discovery rate analysis.

Among the top pathways from the Metascape analysis, enrichments deciphered processes related to immune response and cell function regulation, such as "RNA Metabolism" (109 DEGs among 726 genes) and "Cell Cycle, Mitotic" (82 DEGs among 561 genes). Subsequently to these processes, we identified "ncRNA metabolic process", a direct descendant from the “RNA Metabolism” term and “mitotic cell cycle,” a direct descendant from the “Cell Cycle, Mitotic” term.

The DisGeNET analysis identified several noteworthy pathways. These included "Finding of Mean Corpuscular Hemoglobin" (69 DEGs among 653 genes), "Strabismus" (62 DEGs among 716 genes), and "Nystagmus" (65 DEGs among 772 genes). The latter two, classified as ophthalmologic diseases, shared 32 common genes.

Using Metascape with high confidence DEGs (*P*_adj_ <0.01), we confirmed the top-level pathways previously found with more significant enriched terms (data not shown). By contrast, DisGeNET with the high confidence DEGs uncovered the presence of four RA terms (Fig. 7), of which, three shared the same six DEGs (*AREG*, *DUSP2*, *NR4A2*, *NR4A3*, *CD83* and *ETNK1*). Moreover, “Arthritis, Psoriatic”, the fourth term, added six supplemental DEGs. These findings are of paramount importance, given that 52% of the participants and 85% of the subjects in the CC group fell within the CHIKR subgroup with long-lasting poly-arthralgia/arthritis as cardinal manifestation. This spotlights a potential connection between the molecular profiles of CC individuals and the pathogenesis of RA and other arthritides, far beyond cases diagnosed with RA (n=13), psoriatic arthritis (n=2) and ankylosing spondylarthritis (n=1), respectively.

**Figure 7.**
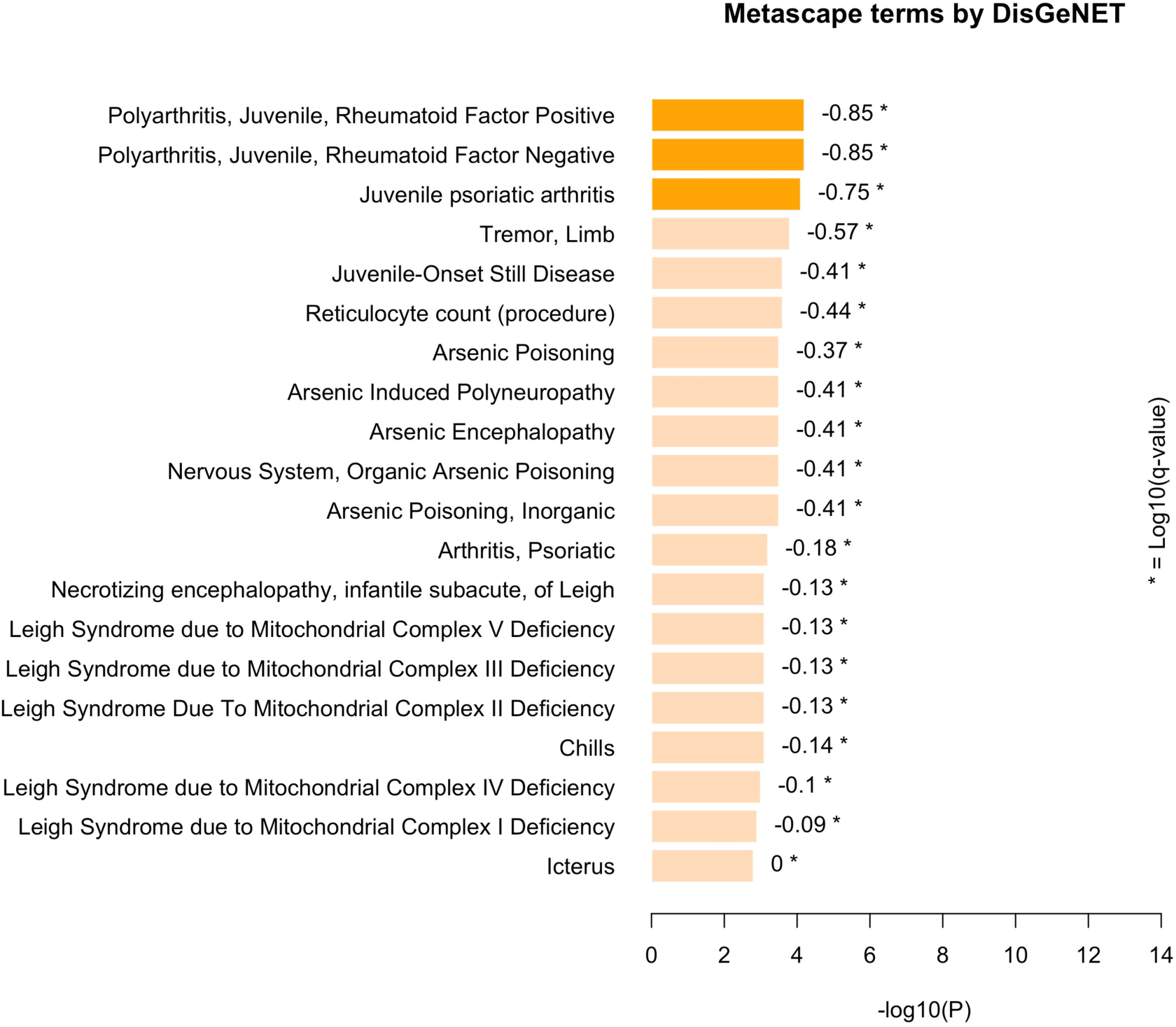
Bar graph of gene set enrichment analysis in the DisGeNET using Metascape. Coloured by p-values, using DEG list with *P*_adj_ <0.01. Significance of the terms are marked with * at the threshold of the log of q-value from the false discovery rate analysis.

Using the STRING tool to map the gene expression data onto the KEGG pathway database, we observed numerous similarities between the findings in STRING and Metascape, particularly the identification of interconnected pathways between RNA modulation and the immune system, such as "RNA transport" and “Viral protein interaction with cytokine and cytokine receptor”. In addition, new pathways were uncovered such as “Neuroactive ligand-receptor interaction”, “Ribosome biogenesis in eukaryotes”, “Protein processing in endoplasmic reticulum” and “Cardiac muscle contraction” (Table 2).

**Table 2.** KEGG pathways using the STRING tool.

| Pathway | Description | enrichment score | number of DEGs | pathway size | FDR |
| --- | --- | --- | --- | --- | --- |
| hsa04080 | Neuroactive ligand-receptor interaction | 0.388267 | 200 | 329 | $1.72^{-06}$ |
| hsa03013 | RNA transport | 0.329103 | 151 | 161 | $2.50^{-04}$ |
| hsa03008 | Ribosome biogenesis in eukaryotes | 0.397400 | 72 | 77 | $4.20^{-04}$ |
| hsa04141 | Protein processing in endoplasmic reticulum | 0.271261 | 155 | 163 | $5.00^{-03}$ |
| hsa04061 | Viral protein interaction with cytokine and cytokine receptor | 0.279189 | 83 | 96 | $8.50^{-03}$ |
| hsa04260 | Cardiac muscle contraction | 0.186342 | 66 | 87 | $8.50^{-03}$ |

DEGs: differentially expressed. FDR: false discovery rate.

#### 3.3.3 Weighted gene co-expression network analysis

The WGCNA identified five distinct clusters of gene expression profiles associated with CC or NCC groups. These five clusters of gene co-expression profiles are named by colours (grey, blue, turquoise, brown, yellow) (Supplementary file 3, S2 fig.). The blue and yellow clusters with 536 genes (*P*_adj_ <3.03^-04^) and 35 genes (*P*_adj_ <6.44^-03^) respectively, were identified as overexpressed in CC compared to NCC individuals. Alternatively, the grey cluster with 62 genes (*P*_adj_ <1.05^-09^), the turquoise cluster with 863 genes (*P*_adj_ <2.72^-03^) and the brown cluster with 53 genes (*P*_adj_ <2.92^-03^) identified as overexpressed in NCC compared to CC individuals.

In the aim to functionally characterise the gene clusters, their gene list was subjected to functional GSEA by Metascape and DisGeNET. The over-expressed clusters in CC individuals and those over-expressed in NCC individuals are described in Supplementary file 3 (S3 fig. and S4 fig., respectively).

All the shared pathways found common through the different Metascape analyses are summarized in Table 3.

**Table 3.**
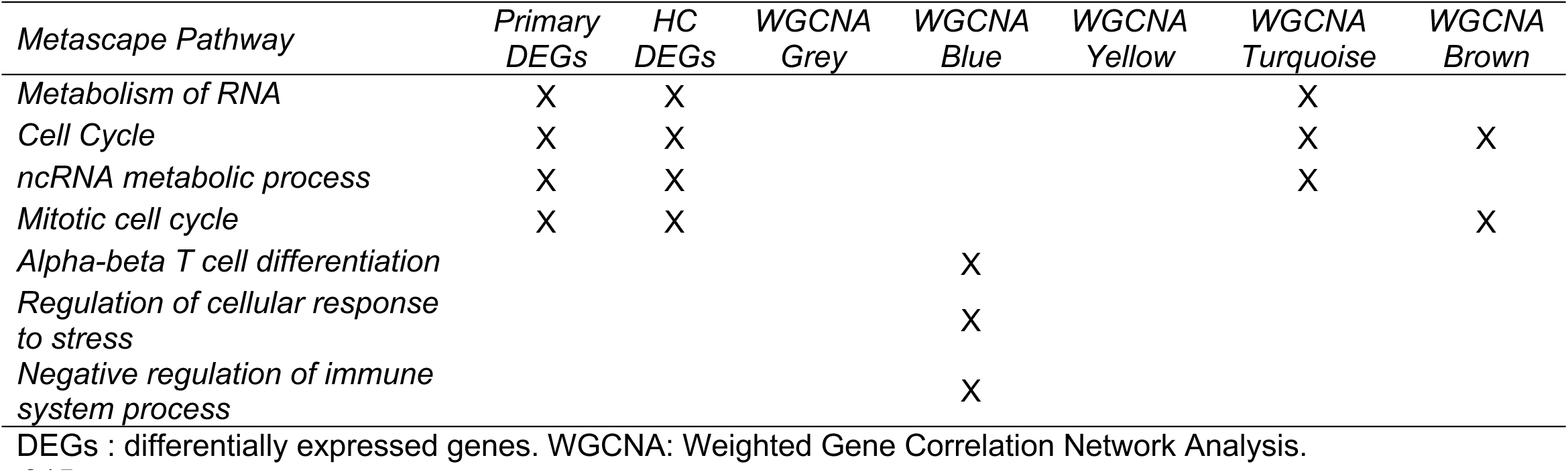
Summary of pathway terms found in Metascape.

| *Metascape Pathway* | *Primary DEGs* | *HC DEGs* | *WGCNA Grey* | *WGCNA Blue* | *WGCNA Yellow* | *WGCNA Turquoise* | *WGCNA Brown* |
| --- | --- | --- | --- | --- | --- | --- | --- |
| *Metabolism of RNA* | X | X |  |  |  | X |  |
| *Cell Cycle* | X | X |  |  |  | X | X |
| *ncRNA metabolic process* | X | X |  |  |  | X |  |
| *Mitotic cell cycle* | X | X |  |  |  |  | X |
| *Alpha-beta T cell differentiation* |  |  |  | X |  |  |  |
| *Regulation of cellular response to stress* |  |  |  | X |  |  |  |
| *Negative regulation of immune system process* |  |  |  | X |  |  |  |

DEGs : differentially expressed genes. WGCNA: Weighted Gene Correlation Network Analysis.

Interestingly, most of the Metascape WGCNA terms were consistent with those found in gene expression mapping and are often observed in viral infections, or immune response to RNA virus infections.

Similarly, the processes found in common by DisGeNET analyses are listed in Table 4. Interestingly, most of the DisGeNET WGCNA terms converged to rheumatoid arthritis and psoriatic arthritis.

**Table 4.**
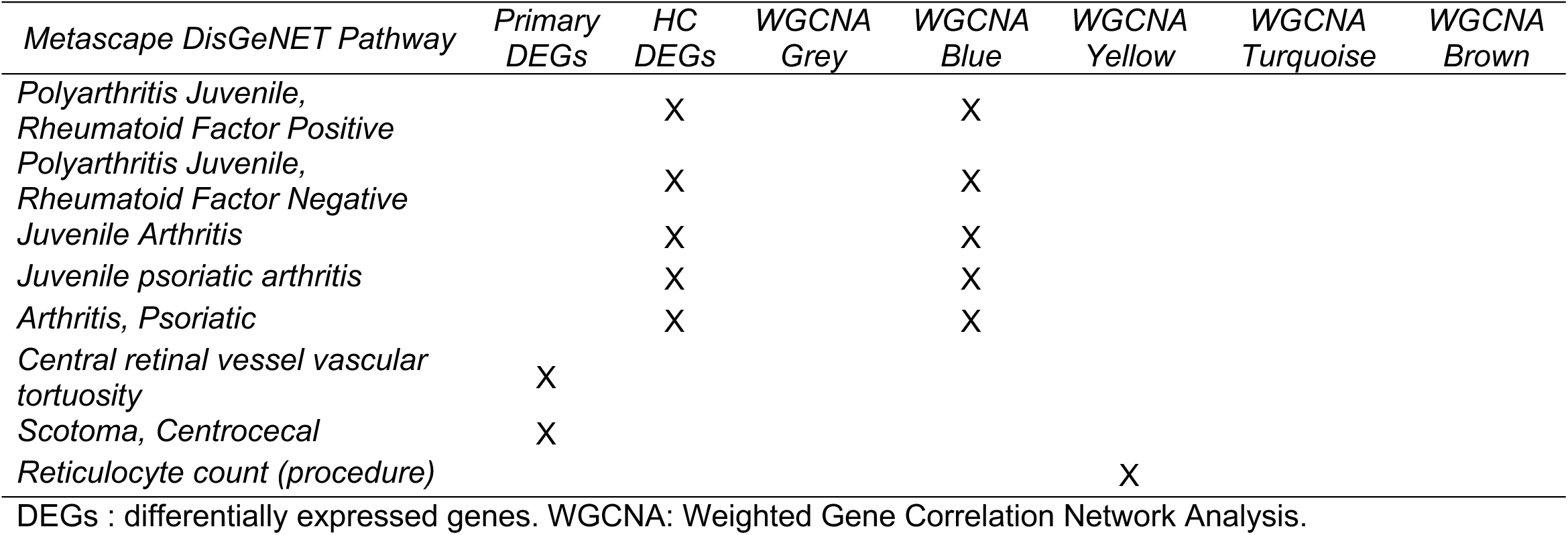
Summary of processes found by DisGeNET discussed in the text.

## 4. Discussion

This study reports the gene expression profiles and related biological processes, associated with the evolution pattern of 219 CHIKV-exposed individuals, 12-to-15 years after the epidemic that hit Reunion Island in 2005-2006, among 133 chronic and 86 nonchronic subjects.

We found first 1549 primary DEGs distinguishing chronic from nonchronic subjects. Among the top 10 ranked DEGs, there were seven genes over-expressed in CC individuals when compared to NCC individuals.

The transcription factor Nuclear Receptor Subfamily 4 Group A Member 2 (*NR4A2,* aka *NURR1*) encodes steroid-thyroid hormone-retinoid receptor superfamily members important in trigger or progression of inflammatory arthritides. *NR4A2* gene is overexpressed in inflamed synovial tissues and cartilages of individuals with RA, psoriatic arthritis, or osteoarthritis and is inactivated by methotrexate or dexamethasone [18]. This has been shown to transactivate the prolactine gene which encodes an immunomodulatory peptide hormone with further roles in autoimmune disease and inflammation [19]. In our study, RA-related terms were identified in the DisGeNET analysis using high confidence DEGs. Although these terms were not significant in the high confidence DEGs analysis, they were significant in the WGCNA blue cluster. *NR4A2* had a larger impact in women, since no differential expression was detected in men. Interestingly, this mirrors the facts that RA is 2 to 3-fold more frequent in women and that *NR4A2* is upregulated by oestrogens [20].

*Tropomodulin 1 (TMOD1)* is an actin-capping protein that regulates tropomyosin in inhibiting depolymerization and elongation of the pointed end of actin filaments [21]. In our study, the strong association between *TMOD1* up-regulation and the enriched pathway named "actin filament capping" in the WGCNA yellow cluster confirms known gene functions. This might be reminiscent of past interactions between viral proteins and the actin cytoskeleton at early stages of infection to subvert the immune system and spread the disease [18], however at late stage, it would more likely indicate osteoarthritis *sequelae* than an active inflammatory mechanism as seen in RA [22].

Another interesting gene is the oxysterol-binding protein 2 (*OSBP2*), a closely related paralog of OSBP, which is a membrane protein lipid exchanger key to cholesterol trafficking and neuronal homeostasis [23]. The *OSBP* gene has been identified as a phosphatidyl-inositol 4 effector of hepatitis C virus replication [23]. A recent review also suggested that it could be useful for dengue virus replication [24]. These findings might underscore a persistent role of *OSBP2* gene in modulating lipid metabolism long after the onset of infection but it does not advocate the possibility of CHIKV latent reservoir reminiscent of acute-stage flavivirus replication processes, as current state-of-art literature yields no result for OSBP genes in CHIKV infection.

The Spectrin Beta Erythrocytic (S*PTB)* gene is a member of the spectrin gene family, which plays a role in cell membrane organization and cytoskeleton stability. In our study, the *SPTB* gene was upregulated in chronic subjects compared to nonchronic peers. Interestingly, it has been associated with RA in the elderly and a senescence associated secretory phenotype in a proteomic study of PBMCs [25]. However, it is still not recognized as a RA term in gene data bases, and it did not correlate to other RA terms in supportive analyses (data not shown).

The tripartite motif-containing 58 (*TRIM58*) has a E3 ubiquitin ligase activity and is involved in several processes, including positive regulations of erythrocyte enucleation and terminal erythropoiesis. In our study, TRIM58 gene was upregulated in chronic subjects and consistently, connections of this gene with the regulation of the erythropoiesis and systemic juvenile idiopathic arthritis were observed in the yellow ("Reticulocyte count (procedure)") and blue clusters (“Juvenile onset Still disease”) of the WGCNA DisGeNET analysis, respectively, where *TRIM58* was among the included genes. These elements confirm previous literature on *TRIM58* [26, 27]. A few members of the TRIM family have also been implicated in the regulation of innate immune pathways, and most of them, including TRIM58, were found to be altered during viral infections at mRNA levels [28], which in turn might indicate a possible signature reminiscent of a positive regulation by TRIM58 of the acute antiviral response. For instance, *TRIM58* was already described as a novel negative mediator of innate immune control through *TLR2* signalling [29].

*Olfactory Receptor Family 2 Subfamily W Member 3 (OR2W3)* gene belongs to the supergene family of G protein-coupled receptors and is detected in the nasal, thyroid, lung, heart, kidney and skin tissues [30]. In a transcriptomic study of infected patients with SARS-CoV-2, olfactory receptors including *OR2W3* gene were identified as DEGs, and the "Olfactory transduction" pathway was downregulated in patients requiring hospitalization or intensive care as compared to outpatients [31]. Indeed, they were linked to the infection severity and COVID-19 pneumonia [31]. In our study, *OR2W3* was more expressed in chronic subjects, which may be indicative of olfaction involvement occurring at a later stage with CHIKV than with SARs-CoV-2. Interestingly, both *TRIM58* and *OR2W3* genes are less than 10 kB apart in the same region of chromosome 1 (cytogenetic band 1q44), which suggests they might share a potential functional or regulatory role in chronic chikungunya.

The Early Growth Response 3 (*EGR3)* gene is a zinc-finger transcription factor of the *EGR* family with a role in immune responses, muscle, lymphocyte and neuronal development, and endothelial cell growth and migration [32]. *EGR3* was found upregulated in human lung epithelial cells infected with SARS-CoV-1 [32], but also in microarray data from severely progressing Covid-19 and type 1 diabetes patients [33], or from patients with CFS/ME (chronic fatigue syndrome/myalgic encephalomyelitis) [34], which suggests its involvement in both early and late immune responses to various infectious pathogens. Consistently, we found *EGR3* upregulated in CC subjects which gives further credence to persistent CFS-like syndrome as CC endotype with a possible immune response disruption even years after the onset of infection.

Among the 10 top DEGs, three were under-expressed in chronic subjects compared to nonchronic subjects, namely *IGHG3*, *IGHV3-49,* and *SEMA6B*.

Immunoglobulin Heavy Constant Gamma 3 (*IGHG3)* gene located on the chromosome 14 encodes heavy chains of IgG3 antibodies (abs). IgG3 neutralizing abs are major humoral effectors involved in the immune response to CHIKV. Their early appearance through the production of interleukin (IL)-6 has been associated with CHIKV clearance and long-term clinical protection [35].

These data align with the correlation observed between low acute-stage IgG3 ab levels and the prevalence of post-acute COVID-19 syndrome or Long Covid [36], despite in this case no link has been yet established with the downregulation of IGHG3 gene. However, the low expression of *IGHG3* thwarts the expected findings for CHIKV-induced RA (*i.e.*, the most severe form of CC arthritis), according to which the level of *IGHG3* expression should correlate with both the diameter of lymphocytic aggregates in the synovium and disease score activity [37].

The role of Immunoglobulin Heavy Variable 3-49 *(IGHV3-49)* in pathology is more elusive. Interestingly, this gene family disclosed a robust positive correlation between the occurrence of SARS-CoV-2 IgG3 neutralizing abs and presence of *IGHV3* genes in the initial human B cell repertoire [38]. Neutralizing IgG3 abs are traditionally sought to inhibit viral infections, and it is interesting that *IGHV3-49* presented in our study a smaller expression among chronic subjects, which suggests these individuals might have not succeeded in getting rid of the virus through the antibody response. Neutralizing IgG abs against CHIKV glycoproteins have indeed been reported to prevent the budding of nascent virions and blocking viral replication in cell lines [39]. Together, these findings fuel the idea of a persistent latent virus reservoir at distance of the acute infection.

Semaphorin 6b (*SEMA6B*) gene is involved in axon guidance and has been associated with epilepsy and intellectual disability. *SEMA6B* gene expression has also been shown to correlate with infiltrating levels of various effector cells and inflammation in colorectal cancer [40], or to represent a dose-dependent marker of smoking in RA patients [41]. In our study population, *SEMA6B* expression levels were significantly higher among smokers than among nonsmokers both in the chronic and nonchronic groups (data not shown), which suggests that untold smoking exposure might have influenced SEMA6B expression and its relationship to disease outcome, albeit without differential misclassification bias.

In summary, this comprehensive analysis of the 10 top DEGs proposes interesting findings. The *SPTB, TRIM58*, *OR2W3* and *EGR3* genes have already exhibited consistent over-expression in chronic when compared to the nonchronic viral infections, and they are already known for their pivotal roles in immune and antiviral responses. Other genes such as *NR4A2* and *OSBP2* are also connected to the chronic group, with an important expression difference between sexes. *IGHG3* and *IGHV3-49* are clearly down-regulated in chronic subjects and remain so even more than 10 years after the infection event, which argues that low IgG3 antibody responses could not be sufficient to neutralize viruses and could thus favour chronicity in viral infections.

The Metascape pathway analysis spotlights RNA metabolism-related and fundamental cellular processes-related terms, suggesting that chronic chikungunya may have a substantial impact on the regulation of RNA processes within cells, which aligns with chronic viral infections, hinting at potential alterations in cellular machinery that could contribute to prolonged manifestations and disease persistence. The related terms “Cell Cycle” and “Metabolism of RNA”, and their direct descendants had already been reported for viral infections, immunity and inflammation regulation [42, 43], especially for hepatitis B [44], *Herpes simplex* [45], SARS-CoV-2 [46, 47], and RA [48], with genes sometimes considered upregulated but indeed identified as downregulated in the majority of severe cases. In our study, these top-level pathways were associated with an under-expression in chronic subjects since they have been identified by WGCNA in the turquoise and brown clusters of overexpressed genes in nonchronic subjects.

The exploration of human diseases potentially linked with the identified DEGs through the DisGeNET analysis unravelled shared processes between various diseases and nonchronic chikungunya, mainly in the neurological (5 terms) and neuro-ophthalmological (5 terms) spheres, which might reflect early reversible manifestations in the course of the disease (a phenomenon of transcriptome resetting also known as “rapid recovery gene downregulation”, whereby mRNA abundance rapidly decreases after a stress condition) [49]. For instance, the “CNS hypomyelination”, “Spasticity” and “Hyperreflexia” might illustrate the propensity of the CNS white matter to be injured under different acute or subacute presentations (*e.g.*, encephalitis, myelitis, encephalomyelitis, etc…) [50]. "Strabismus" and "Blepharoptosis" (or “ptosis”) processes draws attention on third cranial (oculomotor) nerve palsy [51]. By contrast, the pathogenesis of ‘’Nystagmus” is elusive and might reflect a multi-faceted complex ophthalmic involvement [52]. Surprisingly, the literature search on “Finding of Mean Corpuscular Volume” (or “Reticulocyte count” or hereditary spherocytosis) and chikungunya, the strongest pathway, yielded no result.

Importantly, high confidence DEGs and enriched WGCNA within DisGeNET analyses identified multiple RA-related terms, which highlights the strong association between the gene expression patterns found in our study and this autoimmune condition while also suggesting RA may be the ultimate rheumatic condition observed in CHIKR endotype of chronic chikungunya [10]. The DEGs observed in chronic subjects may thus be linked to the development of, progression towards, or recovery from RA. Several studies have discussed similarities between chikungunya arthritis and RA, showing clinical and pathogenic features, or even gene signatures that mimic RA [53–55]. Interestingly, we found no gene signature of the known biomarkers of chronic chikungunya, which might suggest that in our study chronic subjects have recovered from the rheumatic condition, or that was controlled with medications.

In addition, several relevant pathways emerged from the STRING PPI mapping of the gene expression data onto the KEGG pathway database. Among these, the “Neuroactive ligand-receptor interaction” term, a group of neuroreceptor genes involved in environmental information processing and signal molecule interaction, has recently been identified as a target gene for CHIKV miRNAs in the Gene Expression Omnibus (GEO) database [56].

The disruption of "RNA transport" in chronic subjects through hub gene mapping was consistent with the under-expression found in upstream Metascape and WGCNA analyses.

The "Ribosome biogenesis in eukaryotes" hints a possible alteration in the protein synthesis machinery, reflecting the cellular response to potentially persistent CHIKV infection, since specific viral ribosome biogenesis proteins are required for optimal viral replication [57].

The "Protein processing in endoplasmic reticulum" might indicate a potential for the long-lasting disruption of protein folding and maturation. Interestingly, the inhibition of the unfolded protein response by CHIKV non-structural protein2 has been proposed as a virus strategy to subvert type I interferon response and establish chronic chikungunya arthritis [58, 59].

The "Viral protein interaction with cytokine and cytokine receptor" pathway confirms that we are indeed dealing with the host’s immune response following a viral infection. Together with the three abovementioned terms, this pathway advocates a persistent immune dysregulation in chronic chikungunya. Whether the long-term chikungunya manifestations and signalling pathways might result from chronic infection, or the persistence of CHIKV latent reservoir, or the persistence of reactive CHIKV immune debris, or conversely, might denote an increased susceptibility to RNA viral pathogens and a history of new onset viral infections, remains to be deciphered. Interestingly, we found no term related to lymphoid and myeloid cell subsets proliferation and differentiation, which contrasts the effector role of T cells, NK cells, and monocytes in murine models [60, 61].

The “Cardiac muscle contraction” term highlights the CHIKV cardiotropism, since the infection has been associated with cardiovascular manifestations such as arrhythmia, ventricular dysfunction, hypertrophic cardiomyopathy and myocarditis at the acute stage of the infection. CHIKV has been shown replicating in the heart of immunodeficient mice, inducing heart-specific mutations, host responses with likely implications for long-term cardiovascular disease [62, 63].

This study has several strengths. First, the study was largely population-based, since half of the case-mix came from population-based cohorts (Supplemental file 3), which was likely to ensure the capture of mild common phenotypes of the infection. Second, the exposure was defined with certainty on positive CHIKV-specific serology or CHIKV-positive RT-PCR; the case definition of phenotypes (CC, NC, CHIK-R, CFS-like, ICF, etc…) was defined with a high level of confidence from prospective data in the framework of inception cohorts, or from early-collected retrospective data in the framework of follow-up studies of cross-sectional surveys, which likely minimized misclassification through downsize of recall bias at phenotype assessment. Third, with respect to the topic and the difficulties to maintain the cohort 12-to-15 years after the epidemic, the study sample is relatively important and may provide a clear picture of chronic chikungunya.

This study has also several potential limitations. First, the study sample may be prone to referral bias with a selection of more severe cases, the willingness to participate conditioned to information-seeking behaviour and health literacy, a phenomenon likely encouraged by attrition ever growing with time. Second, it was skewed towards the selection of more women and older adults also conditioned to occupational availability, which is inherent to long-term follow-up [64]. Third, we identified two batch-effects with sex and PBMCs isolation technique, which were removed from normalized transformed counts to avoid measurement bias. These sources of potential bias were controlled by predetermined quota selection of identified phenotypes, exclusion of genetic kinship, stratification and multiple adjustment at time of analysis, which makes these unlikely to skew the overall direction of our findings.

## 5. Conclusions

The CHIKGene cohort study delves into the molecular landscape of gene evolution profiles of chikungunya, 12-to-15 years after the unprecedented 2005-2006 Reunion Island’s epidemic. We identified up to 10 top DEGs, which is a quite large gene set that likely indicates a complex disease with rheumatic as well as nervous and cardiac systems involvements. We also uncovered several disrupted pathways, among which some might be suggestive of viral infection, immune response, or auto-immunity, that could represent future molecular targets for diagnostic and/or therapeutic interventions. Last, we unveiled two gene signatures that suggest chronic subjects may have progressed to persistent disease through a defective neutralizing antibody response to the chikungunya virus.

Given the global public health impact of chronic chikungunya, our findings raise new questions about the origin and potential drivers of a long-lasting post-infective immune response, lest we forget the possibility of a latent reservoir and a puzzling chronic infection. A genome-wide association study is under analysis to decipher the genetic basis of these observations.

## Ethical approval

In accordance with the Helsinki declaration on bioethics and the French regulations on biomedical research (*Loi Jardé*), the CHIKGene protocol fulfilled the Reference Methodology MR-001 of the CNIL (*Commission Nationale Informatique et Libertés*), the French data protection authority. The protocol was approved by the CPP (*Comité de Protection des Personnes*) Sud Méditerranée III (IRB: 2018.01.06 bis) as a study on human subjects with minimal constraints (RIPH2), was registered in both the ANSM (*Agence Nationale de Sécurité du Médicamen*t) (ID-RCB 2017-A02675-48) and Clinical Trials (*NCT03690648*) databases. The biobank was declared to the ARS (*Agence Régionale de Santé*) *Océan Indien*, the authority of the French Ministry of Health in regions, the French Ministry of Superior Education and Research and the CNIL.

## Credit authorship contribution statement

**PG:** Conceptualization, Investigation, Funding acquisition, Project administration, Interpretation of data, Writing – review & editing. **RMS:** Methodology, Analysis, Interpretation of data, Writing – original draft. **SLC:** Methodology, Interpretation of data, Writing – review & editing. **LB:** Investigation. **AM:** Investigation. **TL:** Methodology, Interpretation of data. **MR:** Methodology, Interpretation of data. **JLS**: Investigation. **JPM**: Interpretation of data, Writing – review & editing. **CC:** Analysis, Interpretation of data. **CL:** Analysis, Interpretation of data. **NEJ:** Analysis, Interpretation of data. **JS:** Analysis, Interpretation of data. **MJM:** Methodology, Analysis, Interpretation of data, Writing – review & editing. **CF:** Investigation. **CP:** investigation. **NAY:** Investigation. **CC:** Investigation. **CM**: Investigation. **SP**: Investigation. **SM**: Conceptualization. **JN:** Conceptualization, Methodology, Interpretation of data, Writing – review & editing, Funding acquisition. **CM:** Methodology. **HH:** Methodology, Analysis, Interpretation of data, Writing – review & editing. **JFZ:** Conceptualization, Funding acquisition, Project administration, Analysis, Interpretation of Data, Writing – review & editing, Supervision. All authors read and approved the final manuscript.

## Data availability statement

The RNA-Sequencing data generated in this publication will be made available at the NCBI’s Gene Expression when the work is completed. This paper does not report original code. All data produced in the present study are available upon reasonable request to the authors.

## Acknowledgements

This work supported by the Agence Nationale de Recherche (Generic ANR grant 2016). The authors thank Dr Patrick Poubeau and Prof Loic Raffray and colleagues for the loan of consultation premises and hosting the study in their units. They are indebted to Dr Patrick Mavingui for his support through provision of additional funding. They are indebted to Karim Boussaid, Nadège Naty, Martin Rothenburger, Audrey Rivière, Mickael Grondin, Alison Huitelec, Inserm CIC, DRI, DRCI, CRB, nursing and CNAM staffs and all the participants for their dedication and interest in research along the study.

## Supporting information

**Supplementary file 1:** Full methods appendix

**Supplementary file 2:** Strega checklist

**Supplementary file 3:** Additional results

**S1 fig: PCAs from the comparison of chronic vs non chronic (CC:NCC) participants**

Description: PCAs without any batch effect, showing the stratification of the samples by sex and Ficoll/CPT, grouped by (A) CC:NCC, (B) sex and (C) Ficoll/CPT; PCAs after the extraction of the sex and Ficoll/CPT as stratification drivers using the function removeBatchEffect, grouped by (D) CC:NCC, (E) sex and (F) Ficoll/CPT.

**S2 fig:** Heatmap of cluster-trait relationships named by colours

**S3 fig:** Pathways analysis from gene co-expression network analysis using Metascape of clusters blue and yellow (over-expressed in chronic subjects) coloured by p-values

Description: (A) Bar graph of the top-level enriched terms across input gene list of blue cluster; (B) Bar graph of enrichment analysis in the DisGeNET of blue cluster; (C) Bar graph of the top-level enriched terms across input gene list of yellow cluster; and (D) Bar graph of enrichment analysis in the DisGeNET of yellow cluster. Significance of the terms are evaluated and presented with * by a threshold of the log of q-value from the FDR analysis.

**S4 fig.:** Pathways analysis from gene co-expression network analysis using Metascape of clusters grey, turquoise and brown (over-expressed in nonchronic subjects) coloured by p-values

Description: (A) Bar graph of the top-level enriched terms across input gene list of grey cluster; (B) Bar graph of enrichment analysis in the DisGeNET of grey cluster; (C) Bar graph of the top-level enriched terms across input gene list of turquoise cluster; (D) Bar graph of enrichment analysis in the DisGeNET of turquoise cluster; (E) Bar graph of the top-level enriched terms across input gene list of brown cluster; and (F) Bar graph of enrichment analysis in the DisGeNET of brown cluster. Significance of the terms are evaluated and presented with * by a threshold of the log of q-value from the FDR analysis.

**Supplementary file 4:** MutiQC quality assessment report

**Supplementary file 5:** List of DEGs

