## Supplementary figures and images for "Transcriptomic analysis of chronic chikungunya in the Reunionese CHIKGene cohort uncovers a shift in gene expression more than 10 years after infection"

### FigS1.tiff

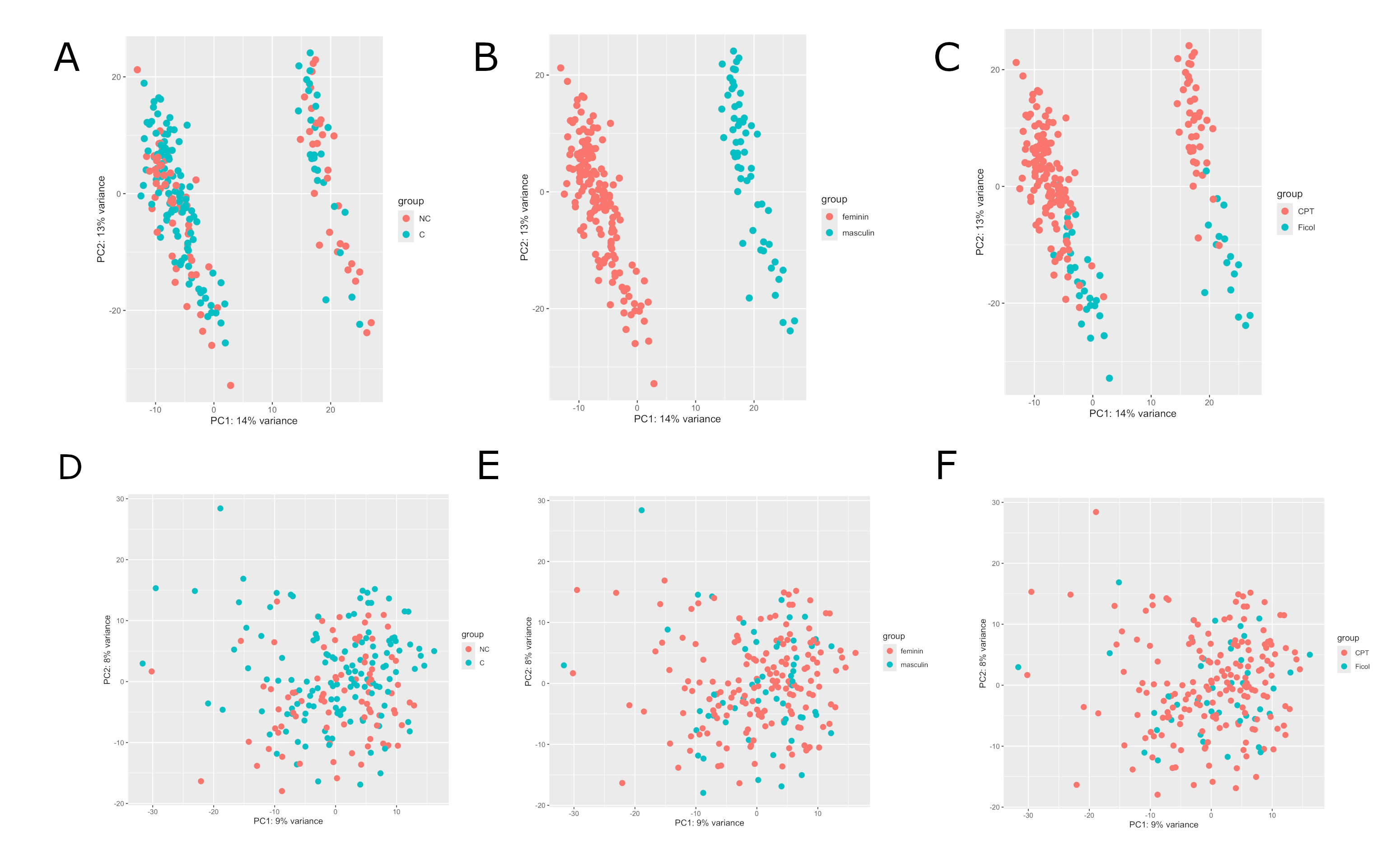

### FigS2.tiff

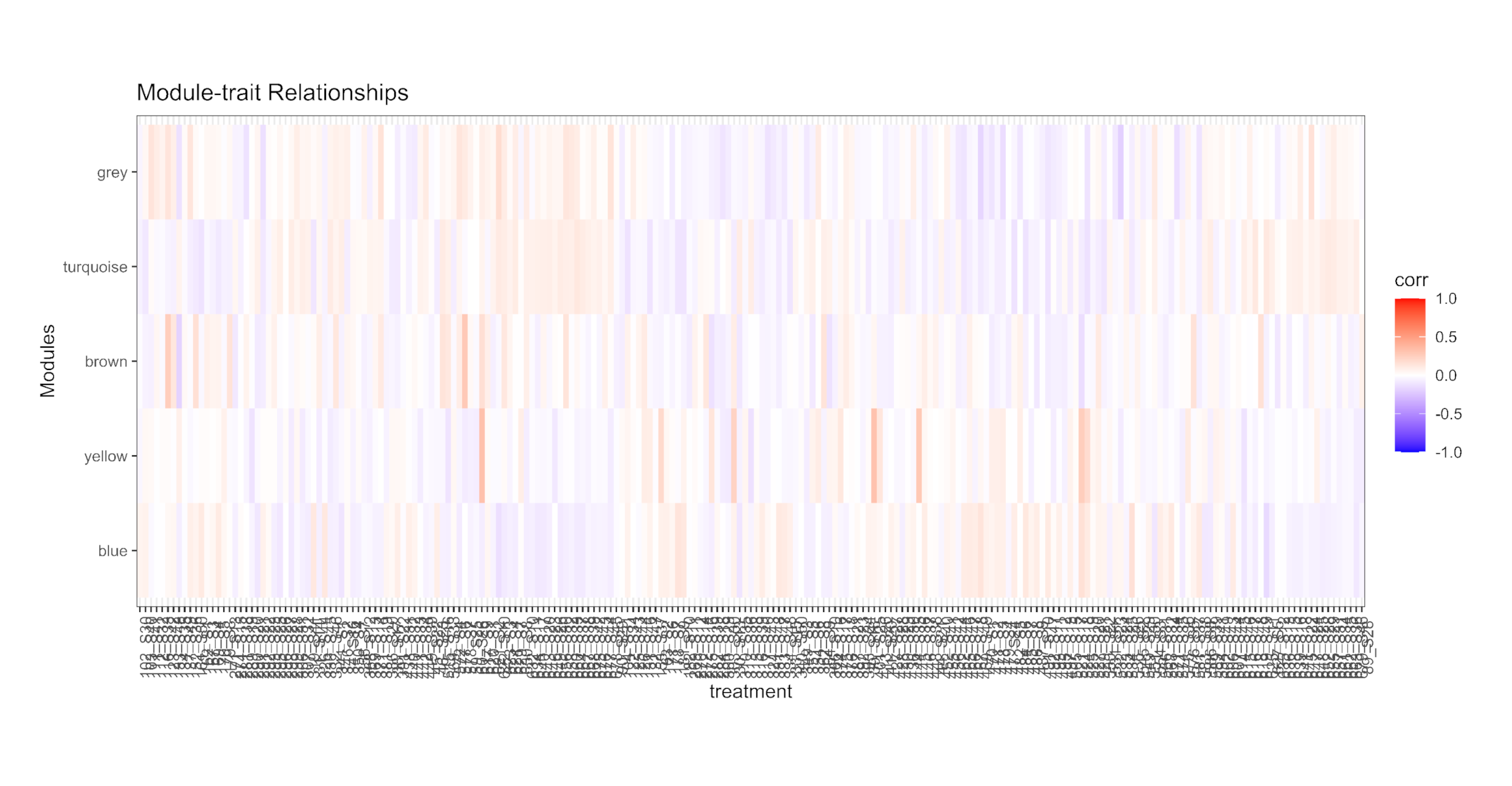

### FigS3.tiff

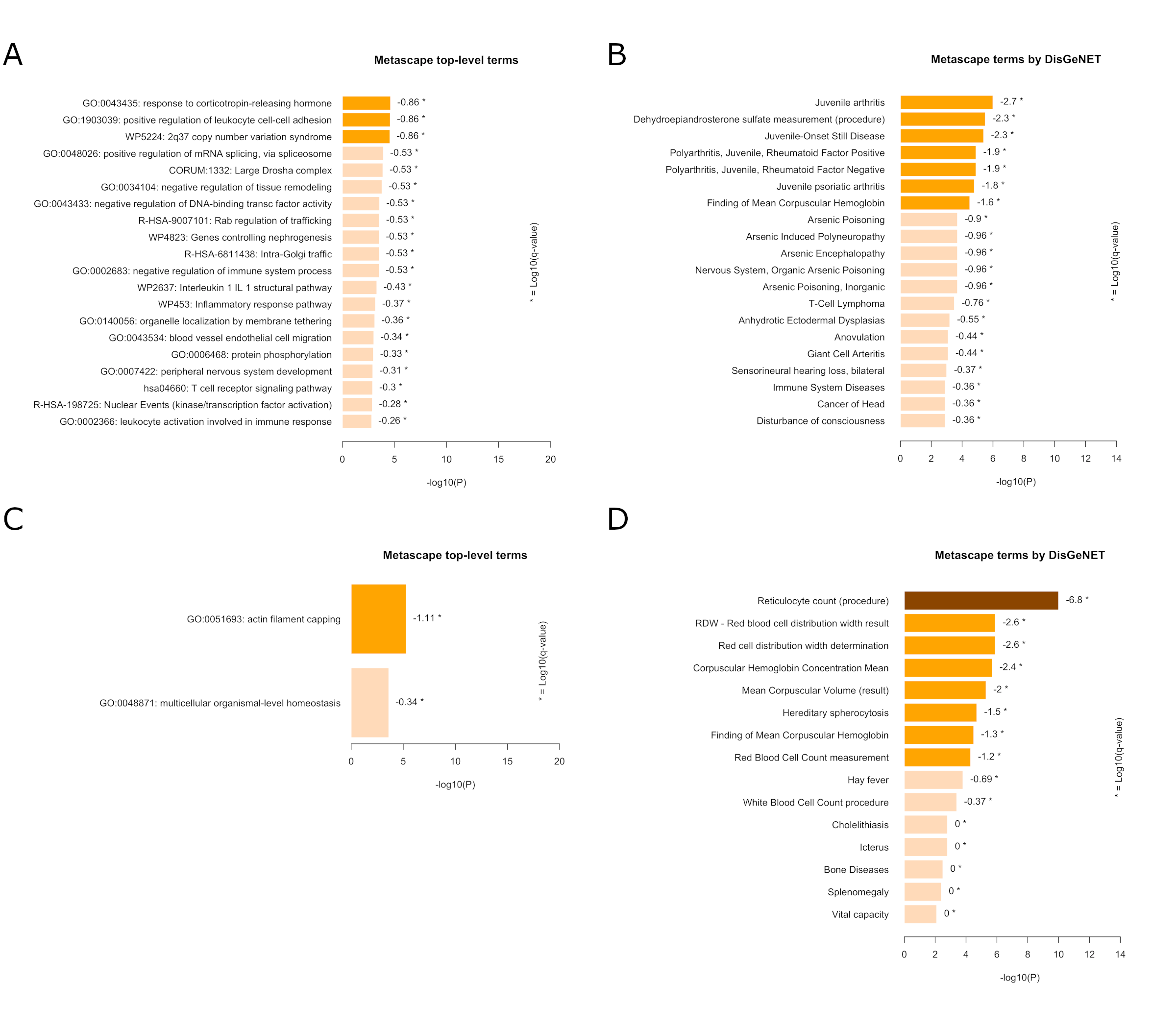

### FigS4.tiff

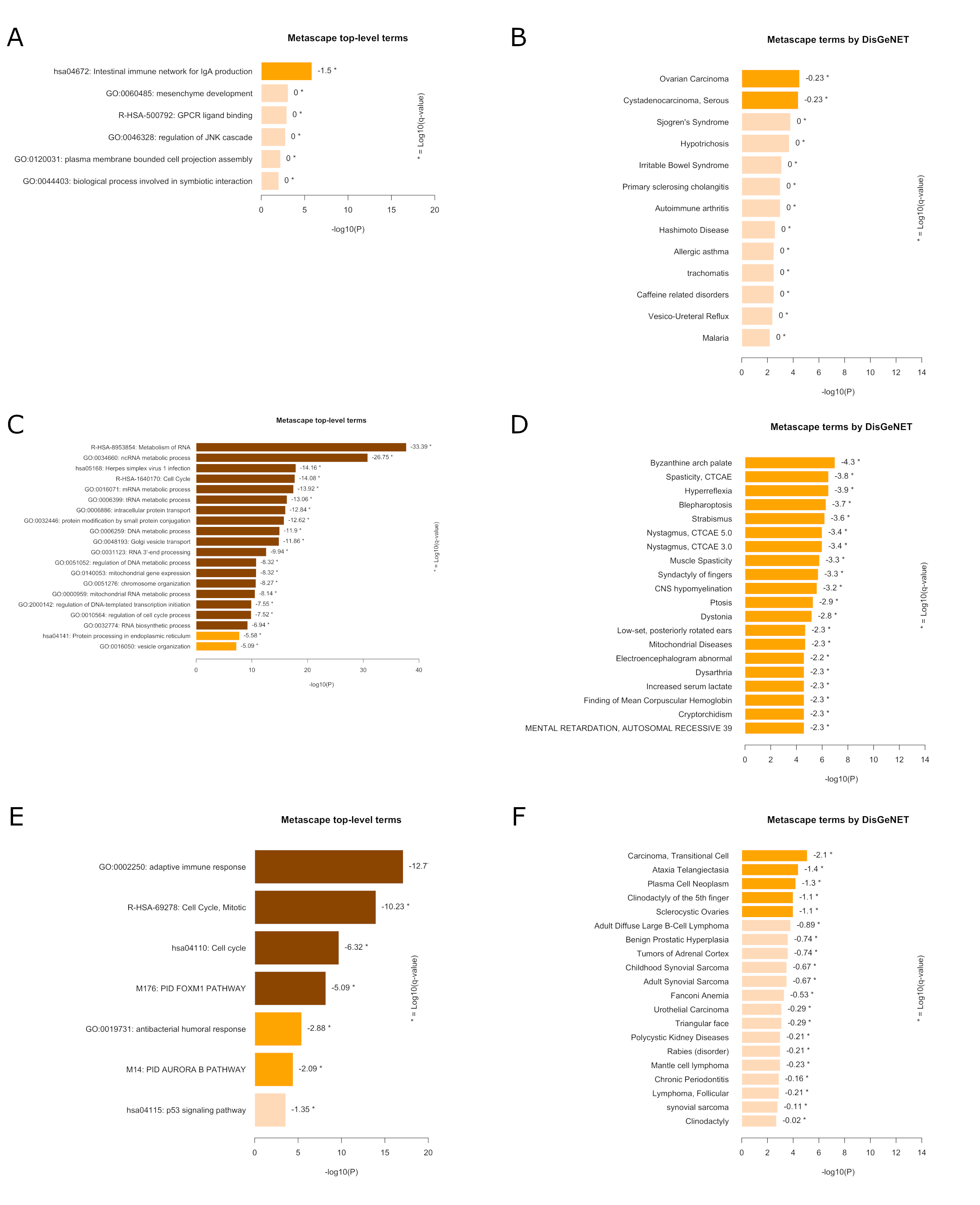
