## Supplemental File 1 for "Transcriptomic analysis of chronic chikungunya in the Reunionese CHIKGene cohort uncovers a shift in gene expression more than 10 years after infection"

Supplementary File 1

Full methods appendix

**Transcriptomic analysis of chronic chikungunya in the Reunionese CHIKGene cohort uncovers a shift in gene expression more than 10 years after infection**

Patrick Gérardin **^a,b,†^**^,✉^, Raissa Medina-Santos **^c,†^**^,✉^, Sigrid Le Clerc **^c^**, Léa Bruneau **^a,d^**, Adrien Maillot **^d^**, Taoufik Labib **^c^**, Myriam Rahmouni **^c^**, Jean-Philippe Meyniel **^e^**, Clémence Cornet ^c,e^, Cécile Lefebvre ^f^, Nora El Jahrani ^f^, Jakub Savara ^g^, Mano Joseph Mathew ^c.g^, Christine Fontaine ^h^, Christine Payet ^h^, Nathalie Ah-You ^h^, Cécile Chabert ^h^, Corinne Mussard ^a^, Sylvaine Porcherat ^a^, Samir Medjane ^i^, Josselin Noirel ^c^, Catherine Marimoutou ^a,d^, Hakim Hocini ^f^, Jean-François Zagury ^c^**^,†^**^,✉^

**^a^** *Clinical Investigation Center, INSERM CIC1410, Centre Hospitalier Universitaire de La Réunion, Saint-Pierre, Réunion, France*

**^b^** *Platform for Clinical and Translational Research, Centre Hospitalier Universitaire de La Réunion, Saint-Pierre, Réunion, France*

**^c^** *Laboratoire Génomique, Bioinformatique et Chimie Moléculaire, EA7528, Conservatoire National des Arts et Métiers, HESAM Université, 2 rue Conté 75003 – Paris, France*

**^d^** *Department of Public Health and Research Support, Centre Hospitalier Universitaire de La Réunion, Saint-Denis, Réunion, France*

**^e^** *AdvanThink, Saint-Aubin, France*

**^f^** *INSERM U955, Equipe 16, Vaccine Research Institute, AP-HP, Groupe Henri Mondor Albert Chenevrier, Créteil, France.*

**^g^** *École d'Ingénieurs Généraliste du Numérique, EFREI, Paris, France*

**^h^** *Biological Resources Center (CRB), Centre Hospitalier Universitaire de La Réunion, Saint-Denis / Saint-Pierre, Réunion, France*

**^i^** *Direction of Clinical Research and Innovation (DRCI), Centre Hospitalier Universitaire de La Réunion, Saint-Pierre, Réunion, France*

^†^ Contributed equally.

^✉^ Correspondence to:

2. **Methods**
   2.1 **Study design, participants, and setting**

The CHIKGene cohort consisted of individuals taken from subsets of five historical cohorts:

1-the regionwide population-based cohort issued from the post-epidemic serosurvey (SEROCHIK/TELECHIK) [1-4];

2- The population-based REDIA (REunion-DIAbète) cohort primarily aimed to assess the prevalence of type-2 diabetes and its related risk factors [5-7];

3- the RHUMATOCHIK cohort of CHIKV-infected outpatients monitored by rheumatologists [8, 9];

4- two catch-up hospital-based inception cohorts (one recruited in the department of infectious and tropical diseases; the other recruited in the emergencies department) for which exposure was well established and reliable follow-up data were available [10-13].

Subjects were selected from preestablished lists of participants to the abovementioned cohorts and offered to participate to the new cohort if they had been tested for CHIKV at the time of the outbreak, were aged between 18 and 75 years, unrelated to second-degree kinship, and had a welfare insurance. Subjects not matching these criteria or pregnant at the time of enrolment were not eligible to the study. Participants were enrolled at the two hospitals of *Centre Hospitalier Universitaire* *Réunion* between August 1, 2018, and September 30, 2020.

2.2 **Ethic statement**

Prior to the study, most of potential participants had given their non-opposition to be recontacted for further follow-up, and each participant provided in due time an *ad hoc* written consent to be enrolled in the CHIKGene cohort study investigating the immune and genetic drivers of the long-term consequences of chikungunya through clinical examination, patient-reported outcomes questionnaires, saliva and whole blood sampling.

2.3 **Exposure and outcome**

Exposure to CHIKV had been defined virologically or serologically with positive RT-PCR and/or positive CHIKV-specific IgM and/or positive CHIKV-specific IgG antibodies as specified at cohort inception or follow-up [1, 8, 10, 12]. In the REDIA follow-up study, a subset of the cohort had been sampled during the epidemic and offered to be tested for CHIKV antibodies using either venipuncture (as indicated to confirm diabetes/prediabetes with oral glucose tolerance test) or the leftover blotting papers from the serosurvey (when the OGTT was not indicated). The Reunionese population had been proved naïve of CHIKV exposure prior to the 2005-2006 outbreak [1].

"Chronic chikungunya" (CC) has been defined as a cluster of three persistent syndromes (>3 months): chikungunya rheumatism (CHIK-R), chronic fatigue (CFS)-like syndrome and idiopathic chronic fatigue (ICF) [4]. These three clinically relevant subgroups of patients were deemed indicative of chronic chikungunya as they reached aetiologic fractions over 60% in the last-in-date evaluation of phenotypes [4]. CHIK-R was defined among subjects who both experienced long-lasting rheumatic musculoskeletal pain in cohorts, reported polyarthralgia/polyarthritis (≥ 4 joints) in the last month and/or exhibited joint pains and/or swollen joints and/or a pathological AIMS2-SF (*Arthritis Impact Measurement Scales 2 Short Form*) score at assessment [4, 14]. CFS-like syndrome was defined in subjects who experienced prolonged fatigue or prolonged post-exertion malaise in the wake of acute chikungunya illness and reported these symptoms plus two of the following among musculoskeletal pain, memory/attention deficits, headaches of new onset, or sleeping disorders/unrefreshing sleep; and/or who exhibited a pathological MFIS-5 (*Modified Fatigue Impact Scale – 5-items*) score at assessment [4, 15]. ICF phenotype was defined in subjects who reported prolonged fatigue alone, either remitting relapsing (recurrent episodes of fatigue ≥ 1 month), or lingering (persistent), over a minimum of six consecutive months [4].

Participants who recovered within 3 months of acute chikungunya illness or who complained of non-specific manifestations (aetiologic fractions ≤ 60% in earlier follow-up evaluation) like musculoskeletal disorders (MSD), mood disorders (ADD), light cerebral disorders or cognitive dysfunction (LCD) were classified as "nonchronic chikungunya" (NCC). Fully asymptomatic subjects were ruled out from this analysis, as done classically for the assessment of chronic chikungunya [16-19].

The clinical data and questionnaires were collected by two investigators (LB, PG) and processed in an electronic case report form (Clinsight v6, Ennov) by two assistant investigators (CM, SP). The samples were collected by nurses (AM) and sent fresh to biological resource operators (CF, CA, NAY, CC) in a timely manner for preanalytical conditioning.

2.4 **Blood sampling, RNA extraction and sequencing**

Heparinized whole blood was collected from the participants. PBMCs were isolated using centrifugation with Ficoll (≤ April 15, 2019) or vacutainer Cell Preparation Tube (CPT; > April 15, 2019), frozen, and stored at -80℃ until RNA extraction. Total RNA was extracted using the R1018 RNeasy Plus Mini Kit (Qiagen), following the manufacturer’s instructions. The quality of the extracted mRNA was assessed using the Bioanalyzer 2100 and the 5067-1513 RNA 6000 Pico Kit (Agilent Technologies). Subsequently, the samples underwent RNA purification using the R1018 ZymoResearch "RNA clean and concentrator-25" as per the manufacturer’s instructions.

Libraries were prepared using the 20020595 TruSeq Stranded mRNA Library Prep Kit (Illumina) according to the manufacturer’s instructions. Sequencing was performed on Illumina HiSeq 2500 sequencer according to the manufacturer’s instructions.

2.5 **Bioinformatic analysis**
    2.5.1 **Quality assessment and Alignment**

The quality of the sequencing reads was assessed using FastQC (v0.11.5) and the results were compiled using the MultiQC tool (v1.20) [20]. Alignment of the reads was performed against the Genome Reference Consortium Human Genome Build 38 (GRCh38) using the Spliced Transcripts Alignment to a Reference Aligner (STAR) (v2.5.2b) [21]. For each participant, the number of reads per gene was calculated based on the total number of reads using the *featureCounts* program (v2.0.1), a tool that counts the total number of mapped reads corresponding to a set of given genomic features.

2.5.2 **Differential gene expression analysis**

For comprehensive differential expression analysis and quality control, we used the DESeq2 package (v1.38.3) [22]. DESeq2 employs a negative binomial distribution model to account for both biological variability and technical noise in RNA-seq data, with the aim to identify genes that exhibit differential expression (DEGs) when comparing disease outcomes [22]. First, we filtered low-count rows from the totality of counts from all samples (<100 counts). Then, we conducted data quality control with the principal component analysis (PCA) using normalized transformed counts (variance-stabilized counts) [22]. The confounders identified by PCA were removed from normalized transformed counts using the *removeBatchEffect* function from the *Limma* package (v3.48.0) [23]. Besides, whenever we observed a correction was needed from visual inspection of the PCA, we used the corresponding variable as a covariate in the differential expression analysis. Subsequently, we proceeded with the differential expression analysis using DESeq2 with the aim to identify genes that exhibited differential expression (DEGs) when comparing chronic and nonchronic subjects. The "Primary DEGs" were characterized first by a *P*_adj_ <0.05. In a second step, we defined "High Confidence DEGs" that were selected from the Primary DEGs by applying a more conservative threshold *P*_adj_ <0.01. In addition, in order to select the Top 10 of the strongly DEGs, the genes with an absolute log2 fold change log2FC >0.7 and an average of normalized count data across the samples (baseMean) higher than 50 were selected for further characterization in specific subgroups: CC (chronic chikungunya), NCC (nonchronic chikungunya), females with CC (fCC), females with NCC (fNCC), males with CC (mCC) and males with NCC (mNCC). Boxplots were generated using the *Boxplot* R function to represent the normalized expression of each of these genes. Stratification by sex was needed given known sex differences in chikungunya recovery time [1], and potential sex-biased gene expression [24].

2.5.3 **Pathway enrichment**

We conducted a gene set enrichment analysis (GSEA) and searched for associations with human diseases (DisGeNET) using the Metascape portal (<https://metascape.org>).to uncover the top-level enriched terms associated with both Primary DEGs and High Confidence DEGs, incorporating default ontologies such as Gene Ontology (GO) processes [25], Kyoto Encyclopedia of Genes and Genomes (KEGG) pathways [26], Reactome gene sets [27], canonical pathways, and CORUM complexes [28]. This analysis was hereafter referred to as “Standard Metascape analysis.”

In addition, while looking for the identification of associations with human diseases also using the Metascape portal, we conducted a comprehensive annotation and another GSEA in DisGeNET using the Primary DEGs and the High Confidence DEGs. To identify the significant terms, a threshold of q-value <0.05 was applied (log10(q-value) <-1.3).

In another approach, the full list of genes (*P*_adj_ <0.01) sorted by the absolute values of log2FC, we used the Search Tool for the Retrieval of Interacting Genes (STRING v12) to provide a protein-protein interaction (PPI) enrichment analysis with KEGG [29], tailored to *Homo Sapiens*.

2.5.4 **Weighted gene co-expression network analysis**

In the aim to elucidate the underlying cellular processes based on the coordinated co-expression of genes encoding the interacting proteins correlated across all individuals (regardless of disease outcome), we performed a gene co-expression network analysis using identified DEGs within the Weighted Correlation Network Analysis (WGCNA, v1.72.5) [30].

We used the normalized and transformed matrix of counts of the DEGs (*P*_adj_ <0.05) generated with DESeq2. In this analysis, we set the minimum cluster size of genes to 30 and merged clusters with >50% similarity. The soft threshold power necessary for achieving a scale-free topology fit was determined as twelve, leading to the identification of co-expression clusters of genes. The LInear Models for MicroArray *(Limma*) data package was used to assess the relationships between gene clusters and phenotypic traits [31]. The gene cluster was represented by the eigengene, which is the first principal component of a cluster's gene expression profile and summarizes the overall expression pattern of the cluster. Additionally, to elucidate the biological relevance of the identified co-expression clusters, we performed an annotation and GSEA using Metascape and DisGeNET tools. To identify the significant terms, a threshold of q-value <0.05 was applied on those results.

The detailed methods of the bulk RNAseq analysis are summarized in Fig. 1.

**2.5.5	Role of funding source and reporting**

The funder of the study had no role in the study design, data collection, data analysis, data interpretation, or writing of the report. The study used the guidance of STREGA checklist, an extension of STROBE guideline for the reporting of genetic association studies [32]. Gene functions were interpreted through Genecards information [33].

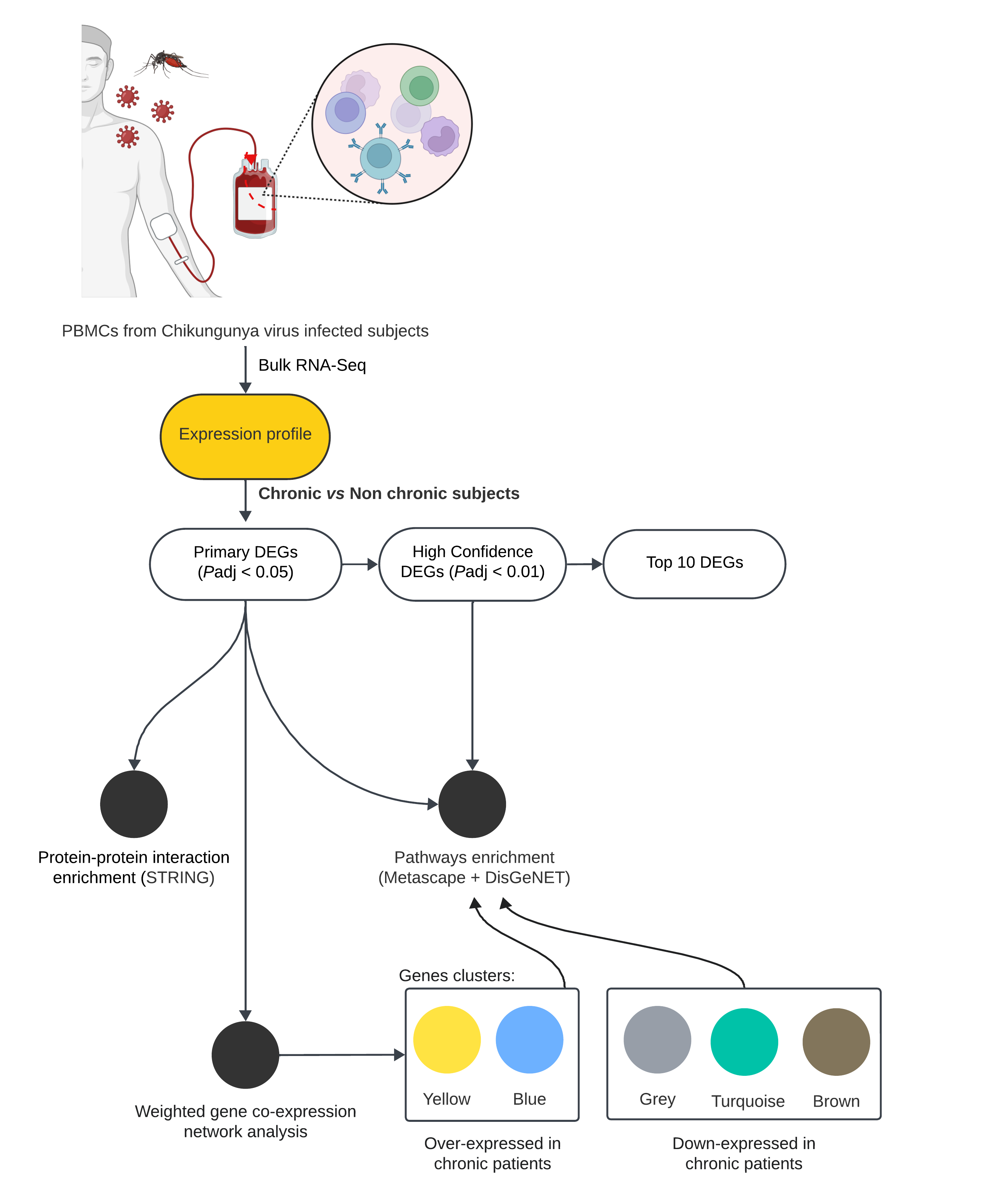

**Figure 1. Methods overview.**

This detailed overview illustrates the different methods used in this bulk RNAseq analysis, from the extraction of PBMCs in CHIKV-exposed subjects. RNA sequencing was performed to identify expression profiles in chronic and nonchronic participants. The resulting list of differentially expressed genes (DEGs), termed Primary DEGs (Padj <0.05), underwent protein-protein interaction enrichment analysis using STRING. Moreover, the Primary DEGs list was employed for pathway enrichment analysis using Metascape. We also subjected the gene clusters resulting from a weighted gene co-expression network analysis to a pathway enrichment in Metascape. High Confidence DEGs (Padj <0.01) were similarly analysed for pathway enrichment using Metascape. Finally, a Top 10 DEGs list (log2FC 0.7 and baseMean >50) was selected for individual gene evaluation and discussion of potential pathomechanisms.
