## Supplemental File 2 for "Transcriptomic analysis of chronic chikungunya in the Reunionese CHIKGene cohort uncovers a shift in gene expression more than 10 years after infection"

### S1 table. STREGA reporting recommendations, extended from STROBE Statement

| **Item** | **Item number** | **STROBE Guideline** | **Extension for Genetic Association Studies (STREGA)** | **Pages, Figures, Tables** |
| --- | --- | --- | --- | --- |
| **Title and Abstract** | 1 | (a) Indicate the study’s design with a commonly used term in the title or the abstract. |  | Pages 1, 2 |
|  |  | (b) Provide in the abstract an informative and balanced summary of what was done and what was found. |  | Pages 2 |
| **Introduction** | | |  |  |
| *Background rationale* | 2 | Explain the scientific background and rationale for the investigation being reported. |  | Pages 3-4 |
| *Objectives* | 3 | State specific objectives, including any pre-specified hypotheses. | ***State if the study is the first report of a genetic association, a replication effort, or both.*** | Page 4 |
| **Methods** | | |  |  |
| *Study design* | 4 | Present key elements of study design early in the paper. |  | Page 4, lines 91-97  Supplement file 1 |
| *Setting* | 5 | Describe the setting, locations and relevant dates, including periods of recruitment, exposure, follow-up, and data collection. |  | Page 4, lines 91-97  Supplement file 1 |
| *Participants* | 6 | (a) **Cohort study –** Give the eligibility criteria, and the sources and methods of selection of participants. Describe methods of follow-up.   **Case-control study –** Give the eligibility criteria, and the sources and methods of case ascertainment and control selection. Give the rationale for the choice of cases and controls.  **Cross-sectional study –** Give the eligibility criteria, and the sources and methods of selection of participants. | ***Give information on the criteria and methods for selection of subsets of participants from a larger study, when relevant***. | Page 4, lines 91-97  Supplement file 1 |
|  |  | (b) **Cohort study –** For matched studies, give matching criteria and number of exposed and unexposed.   **Case-control study –** For matched studies, give matching criteria and the number of controls per case. |  |  |
| *Variables* | 7 | *(a)* Clearly define all outcomes, exposures, predictors, potential confounders, and effect modifiers. Give diagnostic criteria, if applicable. | ***(b)*** ***Clearly define genetic exposures (genetic variants) using a widely-used nomenclature system. Identify variables likely to be associated with population stratification (confounding by ethnic origin).*** | Page 5, lines 102-114  Page 6, lines 139-146  Supplement file 1 |
| *Data sources measurement* | 8***** | *(a)* For each variable of interest, give sources of data and details of methods of assessment (measurement). Describe comparability of assessment methods if there is more than one group. | ***(b)*** ***Describe laboratory methods, including source and storage of DNA, genotyping methods and platforms (including the allele calling algorithm used, and its version), error rates and call rates. State the laboratory/centre where genotyping was done****.* ***Describe comparability of laboratory methods if there is more than one group. Specify whether genotypes were assigned using all of the data from the study simultaneously or in smaller batches.*** | Page 5-6 lines 119-129  Supplement file 1 |
| *Bias* | 9 | *(a)* Describe any efforts to address potential sources of bias. | ***(b) For quantitative outcome variables, specify if any investigation of potential bias resulting from pharmacotherapy was undertaken. If relevant, describe the nature and magnitude of the potential bias, and explain what approach was used to deal with this.*** | Information bias related to exposure:  Supplement file 1  Evaluation bias related to outcome definition:  Supplement file 1  Confounding:  Supplement file 1  (batch effect paragraph) |
| *Study size* | 10 | Explain how the study size was arrived at. |  | Conditioned by funding |
| *Quantitative variables* | 11 | Explain how quantitative variables were handled in the analyses. If applicable, describe which groupings were chosen, and why. | ***If applicable, describe how effects of treatment were dealt with.*** | Page 6, lines 138-146  Supplement file 1 |
| Statistical methods | 12 | (a) Describe all statistical methods, including those used to control for confounding. | ***State software version used and options (or settings) chosen.*** | Pages 6-7, lines 131-166  Supplement file 1 |
|  |  | (b) Describe any methods used to examine subgroups and interactions. |  | Page 6, lines 141-146  Supplement file 1 |
|  |  | (c) Explain how missing data were addressed. |  | No missing data on exposure and outcome variables |
|  |  | (d) **Cohort study –** If applicable, explain how loss to follow-up was addressed.   **Case-control study –** If applicable, explain how matching of cases and controls was addressed.  **Cross-sectional study –** If applicable, describe analytical methods taking account of sampling strategy. |  | NA |
|  |  | (e) Describe any sensitivity analyses. |  | NA |
|  |  |  | ***(f) State whether Hardy-Weinberg equilibrium was considered and, if so, how****.* | NA |
|  |  |  | ***(g) Describe any methods used for inferring genotypes or haplotypes.*** | NA |
|  |  |  | ***(h) Describe any methods used to assess or address population stratification.*** | Supplement file 1. |
|  |  |  | ***(i) Describe any methods used to address multiple comparisons or to control risk of false positive findings.*** | Applying more conservative p-values:  Page 6-7, lines 148-155 |
|  |  |  | ***(j) Describe any methods used to address and correct for relatedness among subjects*** | Eligibility criteria: selection of unrelated subjects  Supplement file 1 |
| **Results** | | |  |  |
| *Participants* | 13***** | (a) Report the numbers of individuals at each stage of the study – e.g., numbers potentially eligible, examined for eligibility, confirmed eligible, included in the study, completing follow-up, and analysed. | ***Report numbers of individuals in whom genotyping was attempted and numbers of individuals in whom genotyping was successful.*** | Page 9, line 191 |
|  |  | (b) Give reasons for non-participation at each stage. |  | Page 9, lines 185-194  Page 9, Fig. 2 |
|  |  | (c) Consider use of a flow diagram. |  | Page 9, Fig. 2 |
| *Descriptive data* | 14***** | (a) Give characteristics of study participants (e.g., demographic, clinical, social) and information on exposures and potential confounders. | ***Consider giving information by genotype****.* | Page 10, Table 1  Supplement file 2 |
|  |  | (b) Indicate the number of participants with missing data for each variable of interest. |  | No missing data for exposure and outcome variables |
|  |  | (c) **Cohort study –** Summarize follow-up time, e.g. average and total amount. |  | NA. No follow-up. No longitudinal analysis |
| *Outcome data* | 15 ***** | **Cohort study-**Report numbers of outcome events or summary measures over time. | ***Report outcomes (phenotypes) for each genotype category over time*** | Page 13, Table 1 |
|  |  | **Case-control study –** Report numbers in each exposure category, or summary measures of exposure. | ***Report numbers in each genotype category*** | NA |
|  |  | **Cross-sectional study –** Report numbers of outcome events or summary measures. | ***Report outcomes (phenotypes) for each genotype category*** | NA |
| *Main results* | 16 | (a) Give unadjusted estimates and, if applicable, confounder-adjusted estimates and their precision (e.g., 95% confidence intervals). Make clear which confounders were adjusted for and why they were included. |  | NA |
|  |  | (b) Report category boundaries when continuous variables were categorized. |  | NA |
|  |  | (c) If relevant, consider translating estimates of relative risk into absolute risk for a meaningful time period. |  | NA |
|  |  |  | ***(d) Report results of any adjustments for multiple comparisons.*** | NA |
| *Other analyses* | 17 | (a) Report other analyses done – e.g., analyses of subgroups and interactions, and sensitivity analyses. |  | NA |
|  |  |  | ***(b) If numerous genetic exposures (genetic variants) were examined, summarize results from all analyses undertaken.*** | NA |
|  |  |  | ***(c) If detailed results are available elsewhere, state how they can be accessed.*** | NA |
| **Discussion** | | |  |  |
| *Key results* | 18 | Summarize key results with reference to study objectives. |  | Page 20, lines 323-328 |
| *Limitations* | 19 | Discuss limitations of the study, taking into account sources of potential bias or imprecision. Discuss both direction and magnitude of any potential bias. |  | Page 27, lines 503-512 |
| *Interpretation* | 20 | Give a cautious overall interpretation of results considering objectives, limitations, multiplicity of analyses, results from similar studies, and other relevant evidence. |  | Pages 20-26, lines 329-492 |
| *Generalizability* | 21 | Discuss the generalizability (external validity) of the study results. |  | Pages 26, lines 493-502 |
| **Other Information** | | |  |  |
| *Funding* | 22 | Give the source of funding and the role of the funders for the present study and, if applicable, for the original study on which the present article is based. |  | Page 7, lines 167-171  Page 27, lines 529-530 |
| *Ethics* | 23 | Report any ethical considerations with specific implications for infectious-disease molecular epidemiology (STROME-ID extension) |  | Page 4, lines 99-101  Supplement file 1  Page 27, ethical approval statement paragraph |

STREGA = STrengthening the REporting of Genetic Association studies; STROBE = STtrengthening the Reporting of Observational Studies in Epidemiology.

* Give information separately for cases and controls in case-control studies and, if applicable, for exposed and unexposed groups in cohort and cross-sectional studies.
