## Supplemental File 3 for "Transcriptomic analysis of chronic chikungunya in the Reunionese CHIKGene cohort uncovers a shift in gene expression more than 10 years after infection"

3. **Results**
   3.1 **Participants**

Women were more represented than men (sex ratio 3:1) which is a well-known sampling bias that occurs in population studies when enrolment is based on the willingness to participate, this being e by sex-differences in occupational availability, health literacy and/or health-information-seeking behaviour. Middle-age (45-64) and elderly (≥ 65) adult participants tended to be overrepresented as compared to young adults (<45), which may illustrate both age-related differences in recreational activity and another health-information-seeking behaviour. These selection incidents on sex and age may have had slight influence in the relationship between these variables and disease outcome, however they were unlikely to bias our findings given female sex and age are known risk factors of chikungunya chronicity.1 Of note, CC was associated with new onset cardiovascular disease and cumulative comorbidities (in a non-linear fashion with a threshold effect from 1 to 2 added comorbidity). Among the 133 CC individuals, we observed a strong overlap between CHIKR and fatigue phenotypes, 51% sharing at least two phenotypes. Of the 113 individuals classified as CHIKR, 13 matched the American College of Rheumatologists criteria for RA. Among the 86 NCC subjects, MSD was the most common phenotype (49%), ahead of ADD (20%) and LCD (15%). Interestingly, ADD was more likely in CC than in NCC subjects but neither in MSD nor in LCD, suggesting mood disorders might complicate CC in the long run.

3.2 **Reads quality, principal component analysis and covariate identification**

The quality assessment of RNA sequences is displayed as a multiQC quality report in Supplementary file 3.

Considering a GC content around 49% and a mean quality value (Phred score) greater than 30 across each base position of the reads, the number of successfully mapped reads to the GRCh38 reference genome ranged from a minimum of 23 million reads to a maximum of 62 million reads, with an average of 43 million reads per individual.

To identify some putative batch effects and check the reliability of sampling thereof, a PCA was conducted (S1 fig.).

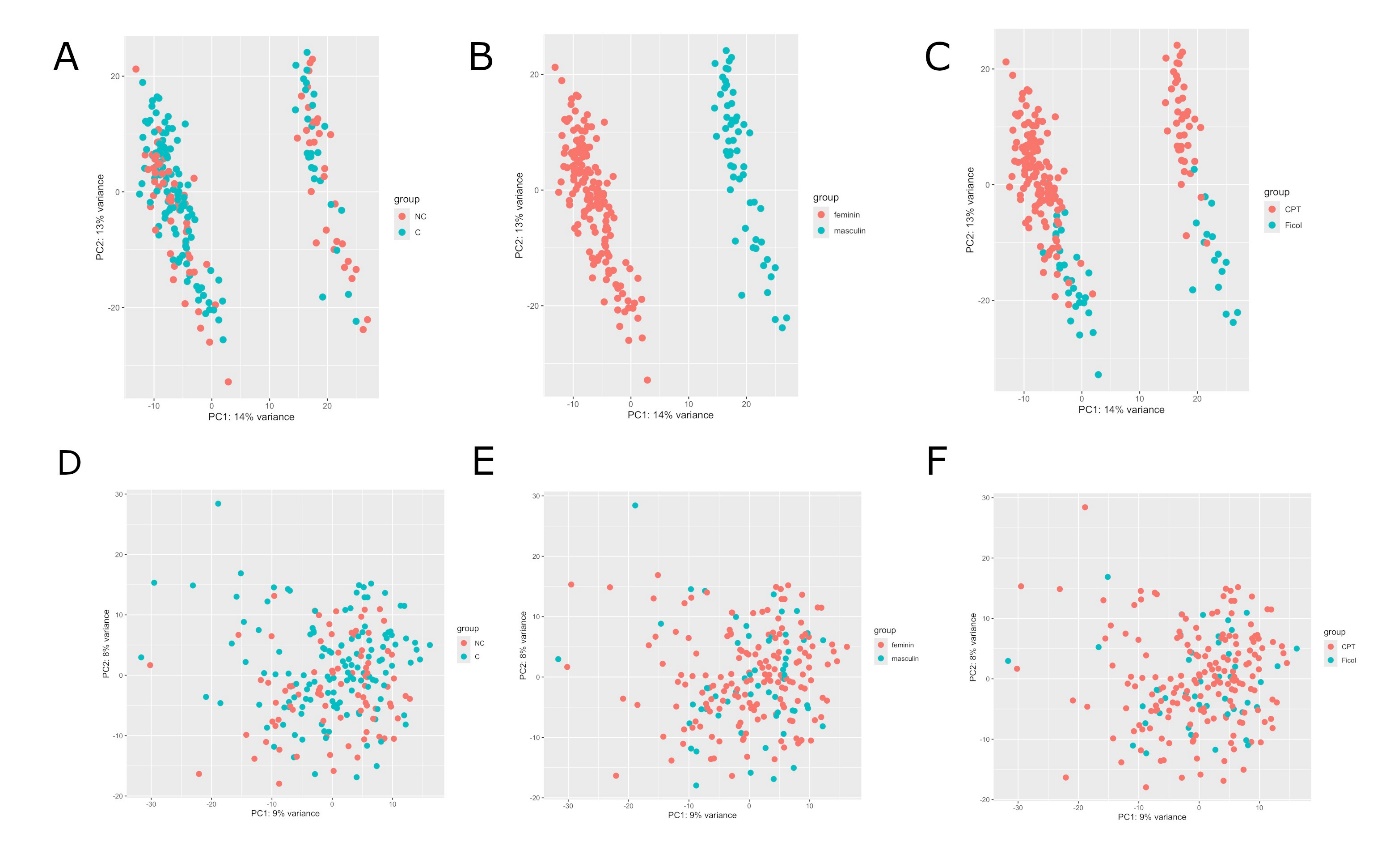

**S1 Fig. 1.** **PCAs from the comparison of chronic vs non chronic (CC:NCC) participants**

PCAs without any batch effect, showing the stratification of the samples by sex and Ficoll/CPT, grouped by (A) CC:NCC, (B) sex and (C) Ficoll/CPT; PCAs after the extraction of the sex and Ficoll/CPT as stratification drivers using the function removeBatchEffect, grouped by (D) CC:NCC, (E) sex and (F) Ficoll/CPT.

Along, the first PCA axis revealed that individuals were stratified (S1 fig.A). Looking for possible reasons for the separation of the individuals and given the abovementioned, we identified sex and PBMCs isolation technique (Ficoll or CPT) as two potential effect modifiers (S1 fig.B and S1 fig.C). After accounting for sex and Ficoll/CPT effects from normalized transform counts, the first PCA no longer showed differentiation (S1 fig.D, S1 fig.E and S1 fig.F). As a result, we used both sex and PBMCs isolation technique as covariates to avoid further bias in subsequent differential expression analysis. Sex-biased stratification was expected in a mixed dataset given inherent biological differences between males and females, including genetic, hormonal, and physiological changes. The Ficoll/CPT effect-modification might have been attributed to solution-specific or operator-dependent variations that arose during the PBMCs isolation process.

3.3 **Bioinformatic analysis**
    3.1.1 **Top genes identification**

All results are found in the article.

3.1.2 **Pathway enrichment**

All results are found in the article.

3.1.3 **Weighted Gene co-expression network analysis**

The WGCNA identified five distinct clusters of gene expression profiles associated with CC or NCC groups. These five clusters of gene co-expression profiles are named by colours (grey, blue, turquoise, brown, yellow) in S2 fig.

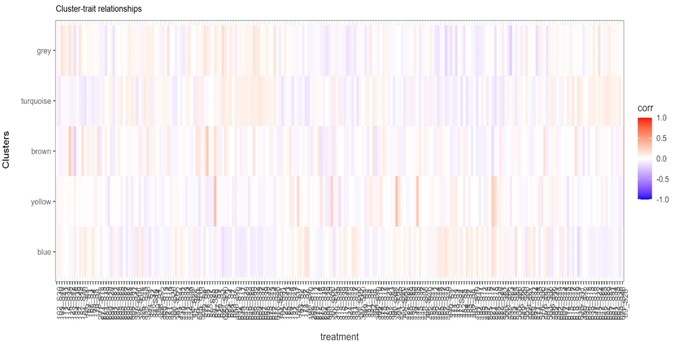

**S2 Fig. Heatmap of cluster-trait relationships named by colours.**

Among the over-expressed enriched terms in CC individuals against the NCC, the Metascape analysis of the blue cluster did not show significant enriched terms (S3.A fig.). By contrast, the DisGeNET analysis of the blue cluster found seven significant terms (S3.B fig.).

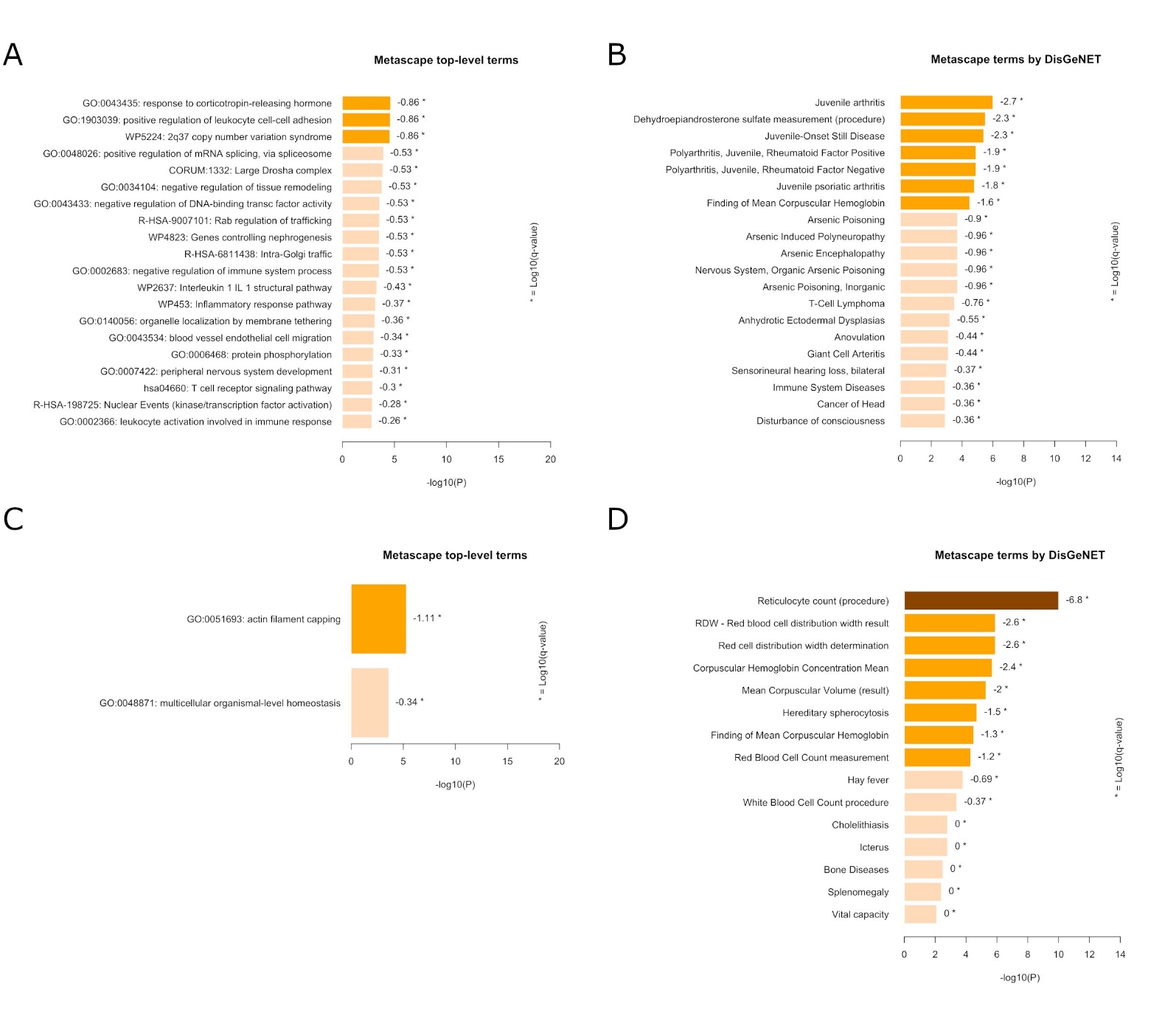

**S3 Fig. Pathways analysis from gene co-expression network analysis using Metascape of clusters blue and yellow (over-expressed in chronic subjects) coloured by p-values.** (A) Bar graph of the top-level enriched terms across input gene list of blue cluster; (B) Bar graph of enrichment analysis in the DisGeNET of blue cluster; (C) Bar graph of the top-level enriched terms across input gene list of yellow cluster; and (D) Bar graph of enrichment analysis in the DisGeNET of yellow cluster. Significance of the terms are evaluated and presented with * by a threshold of the log of q-value from the FDR analysis.

Of these, four were known to be associated with RA, as also identified with the more stringent analysis on high confidence DEGs. In the yellow cluster, also over-expressed, the Metascape analysis displayed only one significantly enriched term, named "actin filament capping" (S3.C fig). This enriched pathway had been yet connected with the top DEG *TMOD1*, an actin-capping protein, responsible for the control actin depolymerization and nucleation. The DisGeNET analysis of the yellow cluster found eight significant terms (S3.D fig.), and the strongest process was "Reticulocyte count (procedure)", with the following present DEGs: *SPTA1*, *SPTB*, *TNS1*, *NPRL3*, *TMCC2*, *TRIM10*, *TRIM58* and *SOX6* (8 DEGs among 234 genes). This enriched pathway underscored a significant association with *TRIM58*, one of the top DEGs, which suggests its involvement as hub gene in a key biological function.

Among the over-expressed clusters in NCC individuals against the CC, the grey cluster displayed only one significantly top-level enriched term in Metascape analysis, named “Intestinal immune network, for IgA production” (S4.A fig.). The DisGeNET analysis of the grey cluster did not show significant enriched terms (S4.B fig.).The turquoise cluster displayed top 20 significant terms in Metascape analysis, with “Metabolism of RNA” (98 DEGs among 726 genes) and “Cell Cycle” (67 genes among 692 genes) amongst the most prominent (S4.C fig.). These pathways included several direct descendants of the RNA metabolism term that had been already detected in the initial Metascape analysis using primary DEGs. That showed a trend toward those terms to be over-expressed in NCC individuals. Similarly, the "Cell cycle" (15 DEGs among 692 genes) pathway and few of its direct descendants were also identified through the Metascape analysis on the brown cluster, as already found in upstream Metascape analysis using primary DEGs. Interestingly, the same “Cell Cycle" pathway was found in the turquoise and brown clusters of genes co-over-expressed among NCC individuals.

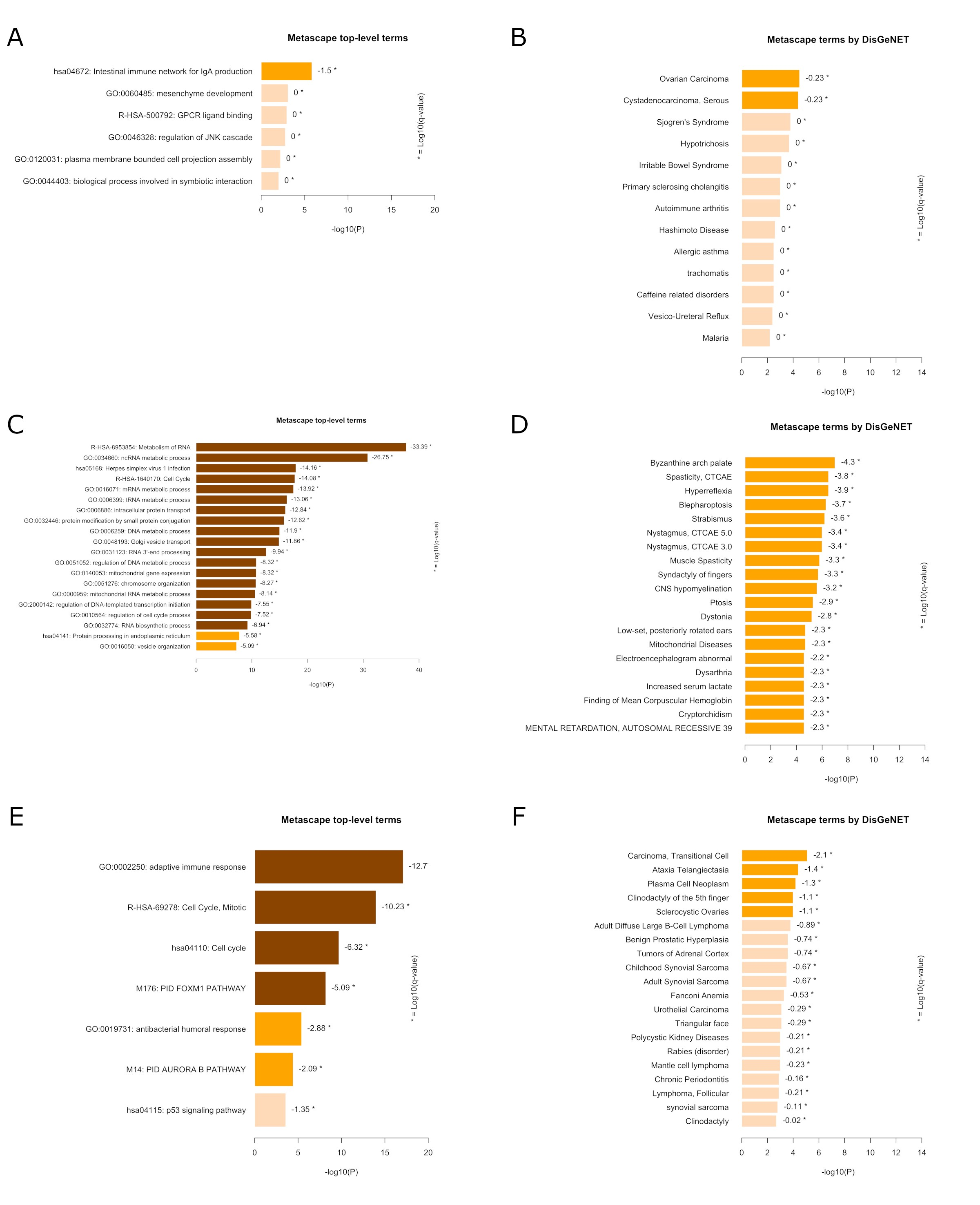

**S4 Fig. Pathways analysis from gene co-expression network analysis using Metascape of grey, turquoise and brown clusters (over-expressed in nonchronic subjects) coloured by p-values.** (A) Bar graph of the top-level enriched terms across input gene list of grey cluster; (B) Bar graph of enrichment analysis in the DisGeNET of grey cluster; (C) Bar graph of the top-level enriched terms across input gene list of turquoise cluster; (D) Bar graph of enrichment analysis in the DisGeNET of turquoise cluster; (E) Bar graph of the top-level enriched terms across input gene list of brown cluster; and (F) Bar graph of enrichment analysis in the DisGeNET of brown cluster. Significance of the terms are evaluated and presented with * by a threshold of the log of q-value from the FDR analysis.

In the turquoise cluster, the DisGeNET analysis presented several significant terms, such as “Strabismus” and “Blepharoptosis” (S4.D fig.), also found with primary DEGs. In addition, the brown cluster identified the “adaptive immune response” (18 DEGs among 653 genes) as overexpressed in NCC individuals (S4.E fig.). This means the adaptive response was in contrast downregulated in CC individuals. The DisGeNET analysis of the brown cluster did not show significant enriched terms (S4.F fig.). In summary, different overexpressed genes were at stake for the NCC group since the clusters were different, but together they supported independently the relevance of the “Cell Cycle” pathway.
