## Supplemental File 4 for "Transcriptomic analysis of chronic chikungunya in the Reunionese CHIKGene cohort uncovers a shift in gene expression more than 10 years after infection"

v1.20

Toolbox

General Stats FastQC

Sequence Counts

Sequence Quality Histograms Per Sequence Quality Scores Per Base Sequence Content Per Sequence GC Content Per Base N Content Sequence Length Distribution Sequence Duplication Levels

##### A modular tool to aggregate results from bioinformatics analyses across many samples into a single report.

Report generated on 2024-06-25, 11:33 CEST based on data in: /media/raissaNFS3/chikgene/fastqc_219

### General Statistics

Configure columns	Table

##### General Statistics

FastQC

% Dups

Export Plot

Overrepresented sequences by sample Top overrepresented sequences Adapter Content

Status Checks

Software Versions

FastQC

% GC

FastQC M Seqs

0%	20%	40%	60%	80%	100%

0%	20%	40%	60%	80%	100%

0 M	10 M	20 M	30 M	40 M	50 M	60 M

Created with MultiQC

FastQC *Version:* 0.11.5

FastQC is a quality control tool for high throughput sequence data, written by Simon Andrews at the Babraham Institute in Cambridge.

Sequence Counts

Sequence counts for each sample. Duplicate read counts are an estimate only.

Percentages

Help

Export Plot

#### Sequence Quality Histograms 219

The mean quality value across each base position in the read.

Help

Export Plot

##### FastQC: Mean Quality Scores

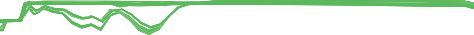
35

30

25

Phred Score

20

15

10

5

0

0	20	40	60	80	100

Position (bp)

Created with MultiQC

#### Per Sequence Quality Scores 219

The number of reads with average quality scores. Shows if a subset of reads has poor quality.

##### FastQC: Per Sequence Quality Scores

Help

Export Plot

45M reads

40M reads

35M reads

30M reads

Count

25M reads

20M reads

15M reads

10M reads

5M reads

0 reads

0	5	10	15	20	25

30	35

Mean Sequence Quality (Phred Score)

Created with MultiQC

Per Base Sequence Content	0	219

The proportion of each base position for which each of the four normal DNA bases has been called.

Help

Click a sample row to see a line plot for that dataset.

Rollover for sample name

Position: -	%T: -	%C: -	%A: -	%G: -

#### Per Sequence GC Content 203 1

The average GC content of reads. Normal random library typically have a roughly normal distribution of GC content.

Percentages Counts

Percentage

Help

Export Plot

#### Per Base N Content 219

The percentage of base calls at each position for which an N was called.

##### FastQC: Per Base N Content

Help

Export Plot

5%

4%

3%

Percentage N-Count

2%

1%

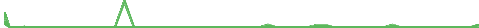
0%

0	20	40	60	80	100

Position in Read (bp)

Created with MultiQC

Sequence Length Distribution	0	219

The distribution of fragment sizes (read lengths) found. See the FastQC help

##### FastQC: Sequence Length Distribution

Export Plot

50M

40M

30M

Read Count

20M

10M

0

40 bp	50 bp	60 bp	70 bp	80 bp	90 bp	100 bp

Sequence Length (bp)

Created with MultiQC

Sequence Duplication Levels	0	219

The relative level of duplication found for every sequence.

% of Library

Overrepresented sequences by sample	34	185

The total amount of overrepresented sequences found in each library.

Help

Export Plot

Help

Export Plot

#### Top overrepresented sequences

Top overrepresented sequences across all samples. The table shows 20 most overrepresented sequences across all samples, ranked by the number of samples they occur in.

Configure columns	Table

##### FastQC: Top overrepresented sequences

Export Plot

FastQC Samples

0	5	10	15	20

FastQC Occurrences

FastQC

% of all reads

0	0.5M	1M	1.5M	2M

0%	20%	40%	60%	80%	100%

Created with MultiQC

#### Adapter Content 219

The cumulative percentage count of the proportion of your library which has seen each of the adapter sequences at each position.

Help

No samples found with any adapter contamination > 0.1%

#### Status Checks

Status for each FastQC section showing whether results seem entirely normal (green), slightly abnormal (orange) or very unusual (red).

###### Min: 0 Max: 1

Help

Export Plot

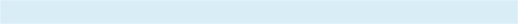

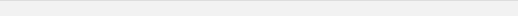

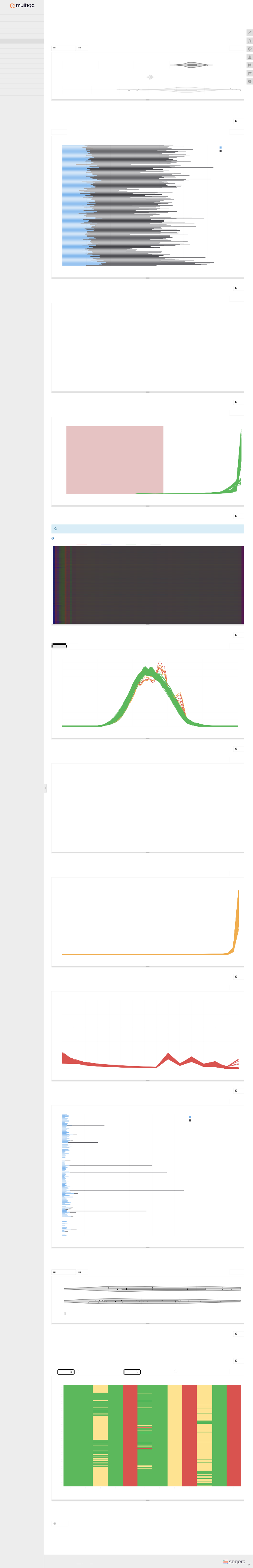

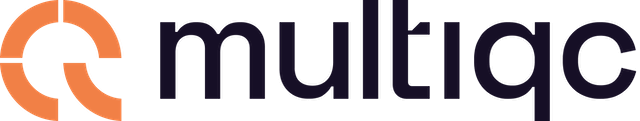

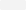

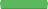

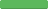

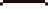

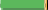

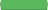

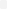

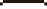

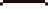

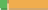

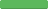

FastQC: Sequence Counts

104_S31

122_S34

131_S46

137_S49

162_S1

170_S5

181_S9

271_S13

281_S38

290_S20

295_S25

300_S29

30_S13

316_S39

324_S43

332_S46

340_S50

34_S16

356_S7

368_S12

375_S17

387_S21

393_S24

41_S20

436_S30

442_S34

448_S38

455_S41

465_S46

46_S23

479_S4

485_S7

492_S47

505_S15

525_S19

531_S1

535_S26

554_S3

571_S34

576_S5

583_S38

59_S25

60_S26

617_S9

621_S2

625_S11

632_S10

638_S15

643_S20

648_S25

656_S30

661_S34

667_S38

677_S44

90_S29

Unique Reads Duplicate Reads

0	10M	20M	30M	40M	50M	60M

Number of reads

Created with MultiQC

FastQC: Per Sequence GC Content

4.5%

4%

3.5%

3%

2.5%

2%

1.5%

1%

0.5%

0%

0% GC

20% GC

40% GC

60% GC

80% GC

100% GC

% GC

Created with MultiQC

FastQC: Sequence Duplication Levels

100%

80%

60%

40%

20%

0%

1	2	3	4	5	6	7	8	9	>10	>50	>100	>500	>1k	>5k	>10k+

Sequence Duplication Level

Created with MultiQC

FastQC: Overrepresented sequences sample summary

104_S31

122_S34

131_S46

137_S49

162_S1

170_S5

181_S9

271_S13

281_S38

290_S20

295_S25

300_S29

30_S13

316_S39

324_S43

332_S46

340_S50

34_S16

356_S7

368_S12

375_S17

387_S21

393_S24

41_S20

436_S30

442_S34

448_S38

455_S41

465_S46

46_S23

479_S4

485_S7

492_S47

505_S15

525_S19

531_S1

535_S26

554_S3

571_S34

576_S5

583_S38

59_S25

60_S26

617_S9

621_S2

625_S11

632_S10

638_S15

643_S20

648_S25

656_S30

661_S34

667_S38

677_S44

90_S29

0%

Top overrepresented sequence

Sum of remaining overrepresented sequences

1%

2%

3%	4%

5%

Percentage of Total Sequences

Created with MultiQC

FastQC: Status Checks

102_S30

116_S33

131_S46

152_S36

167_S3

179_S8

271_S13

282_S39

292_S22

299_S28

30_S13

317_S40

330_S45

33_S15

34_S16

362_S8

371_S14

386_S20

393_S24

427_S27

440_S32

446_S37

455_S41

466_S47

471_S2

484_S6

492_S47

515_S16

529_S21

534_S25

554_S3

572_S4

578_S7

594_S41

60_S26

618_S48

623_S3

630_S8

638_S15

645_S22

652_S28

660_S33

667_S38

678_S45

Basic	Per Base	Per Tile	Per	Per Base	Per Statistics	Sequence	Sequence	Sequence	Sequence	Sequence

Quality	Quality	Quality	Content	GC Content Scores

Per Base N	Sequence	Sequence	Overrepresented	Adapter

Content

Length	Duplication	Sequences	Content Distribution	Levels

Kmer Content

Created with MultiQC

### Software Versions

Software Versions lists versions of software tools extracted from file contents.

Copy table

###### Software Version

FastQC	0.11.5

**MultiQC v1.20** - Written by Phil Ewels, available on GitHub.

This report uses HighCharts, jQuery, jQuery UI, Bootstrap, FileSaver.js and clipboard.js.
