## Supplemental File 5 for "Transcriptomic analysis of chronic chikungunya in the Reunionese CHIKGene cohort uncovers a shift in gene expression more than 10 years after infection"

| Differentially expressed genes between chronic and non-chronic patients | | | | | | | |
| --- | --- | --- | --- | --- | --- | --- | --- |
| gene | baseMean | log2FoldChange | lfcSE | stat | pvalue | padj | selection |
| HSPA12B | 7.45955176796971 | 1.09037975046736 | 0.192493090938808 | 5.66451369838504 | 1,47E-08 | 0.000146144593678815 | High confidence DEG |
| TRIM58 | 1999.92474226596 | 0.862074434914865 | 0.159076746118709 | 5.41923603511195 | 5,99E-08 | 0.000395516896344814 | High confidence DEG |
| OR2W3 | 101.387922765945 | 0.905343466872479 | 0.17037367725884 | 5.31386938075554 | 1,07E-07 | 0.000531885652135975 | High confidence DEG |
| RP11-544A12.8 | 2.28068787343099 | 1.5650926482505 | 0.298165441096121 | 5.24907461608185 | 1,53E-07 | 0.00060607976029319 | High confidence DEG |
| NR4A2 | 326.511970515735 | 1.02739056773801 | 0.199531596500068 | 5.14901191470022 | 2,62E-07 | 0.000766535537678499 | High confidence DEG |
| IGHG3 | 598.938951146387 | -0.967313656534644 | 0.18809074479801 | -5.14280305271499 | 2,71E-07 | 0.000766535537678499 | High confidence DEG |
| MANEA | 526.377974527816 | -0.332695448907839 | 0.067905451878481 | -4.89939231246422 | 9,61E-07 | 0.0021718370070576 | High confidence DEG |
| TRIM10 | 5.98413397582871 | 1.07950079208074 | 0.220557833798099 | 4.89441147245273 | 9,86E-07 | 0.0021718370070576 | High confidence DEG |
| B3GALNT1 | 26.7868953753372 | -0.5185526867807 | 0.107998408021226 | -4.80148454298311 | 1,57E-06 | 0.00284121508268201 | High confidence DEG |
| IGHV3-49 | 97.8748409936067 | -0.891870430804664 | 0.186324436087089 | -4.78665305278476 | 1,70E-06 | 0.00284121508268201 | High confidence DEG |
| TMEM156 | 365.595137582074 | -0.320930796308408 | 0.0673728925355936 | -4.763500337155 | 1,90E-06 | 0.00284121508268201 | High confidence DEG |
| BPGM | 980.097331077349 | 0.550057153167956 | 0.115588187504923 | 4.75876614247048 | 1,95E-06 | 0.00284121508268201 | High confidence DEG |
| TRBJ2-2 | 10.4971091309065 | 0.657483240849282 | 0.138435623872457 | 4.74937897094343 | 2,04E-06 | 0.00284121508268201 | High confidence DEG |
| OSBP2 | 398.893590426356 | 0.838808265681899 | 0.177008420080694 | 4.73880431958834 | 2,15E-06 | 0.00284121508268201 | High confidence DEG |
| CHAC2 | 54.4029133179241 | -0.449201854457251 | 0.0954049216259186 | -4.70837192465359 | 2,50E-06 | 0.00306682154307961 | High confidence DEG |
| MRPS30 | 1090.00265840219 | -0.110395142165415 | 0.0235949880135433 | -4.67875389900811 | 2,89E-06 | 0.00306682154307961 | High confidence DEG |
| TRBJ2-7 | 17.075367998864 | 0.56038957820416 | 0.119984358626521 | 4.67052192985006 | 3,00E-06 | 0.00306682154307961 | High confidence DEG |
| CKS2 | 147.098405370975 | -0.373815800568531 | 0.0801067844454187 | -4.66646867873261 | 3,06E-06 | 0.00306682154307961 | High confidence DEG |
| SLC25A39 | 3523.19923474364 | 0.564032573763567 | 0.12102655073498 | 4.66040360845008 | 3,16E-06 | 0.00306682154307961 | High confidence DEG |
| B3GNT7 | 247.265031135407 | 0.625032635025811 | 0.134287630911684 | 4.65443191440969 | 3,25E-06 | 0.00306682154307961 | High confidence DEG |
| SLC44A2 | 10430.1218027536 | 0.141527055063777 | 0.0306586856128294 | 4.61621404293189 | 3,91E-06 | 0.00352149826603771 | High confidence DEG |
| RTCA | 855.193918170669 | -0.137745751579394 | 0.0300578782620265 | -4.58268379353357 | 4,59E-06 | 0.00381298585294465 | High confidence DEG |
| TMOD1 | 118.327981856546 | 0.803148653907037 | 0.175811104693526 | 4.56824758201187 | 4,92E-06 | 0.00381298585294465 | High confidence DEG |
| RP11-1H8.5 | 3.19616432660758 | 1.43990236619811 | 0.315432656639405 | 4.56484874311592 | 5,00E-06 | 0.00381298585294465 | High confidence DEG |
| GATC | 544.802199220226 | -0.194045016324765 | 0.0425094471155722 | -4.5647504141187 | 5,00E-06 | 0.00381298585294465 | High confidence DEG |
| EGR3 | 165.127476287068 | 0.986423217538975 | 0.217732680676326 | 4.53043252154397 | 5,89E-06 | 0.00416185853437977 | High confidence DEG |
| CELF2-AS1 | 52.0709368126809 | 0.64728601788449 | 0.143064126562465 | 4.52444671796792 | 6,06E-06 | 0.00416185853437977 | High confidence DEG |
| DNAAF2 | 353.575374421582 | -0.200638643036007 | 0.0443566915030027 | -4.52330045901698 | 6,09E-06 | 0.00416185853437977 | High confidence DEG |
| STT3A | 3043.65354547821 | -0.192413033963966 | 0.0428280701370972 | -4.4926851326252 | 7,03E-06 | 0.00423868201872752 | High confidence DEG |
| CDC5L | 2192.84841332222 | -0.156449696209269 | 0.0348570436182404 | -4.48832373516038 | 7,18E-06 | 0.00423868201872752 | High confidence DEG |
| EYA3 | 757.090966866916 | -0.243893673281011 | 0.054358195758073 | -4.48678750057279 | 7,23E-06 | 0.00423868201872752 | High confidence DEG |
| NR4A3 | 27.1949295322092 | 0.714213670700417 | 0.159266631485326 | 4.48438988154415 | 7,31E-06 | 0.00423868201872752 | High confidence DEG |
| CTD-2301A4.6 | 13.3580691031506 | 0.693114704348049 | 0.154589878309231 | 4.48357105865358 | 7,34E-06 | 0.00423868201872752 | High confidence DEG |
| B3GALT2 | 83.4481857686034 | -0.687980544411269 | 0.153777605451646 | -4.473866935245 | 7,68E-06 | 0.00423868201872752 | High confidence DEG |
| FUT8 | 659.988860143605 | -0.240635102335507 | 0.0539444990111183 | -4.46079038172012 | 8,17E-06 | 0.00423868201872752 | High confidence DEG |
| C1R | 44.3827359170116 | 0.294061333257274 | 0.0659861838167754 | 4.45640763335848 | 8,33E-06 | 0.00423868201872752 | High confidence DEG |
| RAMP1 | 38.3118943413561 | 0.493131762318233 | 0.110731209219982 | 4.45341259968148 | 8,45E-06 | 0.00423868201872752 | High confidence DEG |
| NUP35 | 171.1857166158 | -0.259714957115204 | 0.0583195166634191 | -4.45331120650577 | 8,46E-06 | 0.00423868201872752 | High confidence DEG |
| SPTB | 201.301245551734 | 0.723193495497562 | 0.162483916046917 | 4.4508620489473 | 8,55E-06 | 0.00423868201872752 | High confidence DEG |
| CCDC117 | 1367.97675439009 | -0.205008613244524 | 0.0461154297060957 | -4.44555357178911 | 8,77E-06 | 0.00423874981621045 | High confidence DEG |
| INSIG1-DT | 122.215638751766 | 0.290812072648684 | 0.0657043072422839 | 4.42607318841849 | 9,60E-06 | 0.00432443369002112 | High confidence DEG |
| RAB23 | 38.5558077727205 | -0.32321382747741 | 0.0731805783792171 | -4.41666128685861 | 1,00E-05 | 0.00432443369002112 | High confidence DEG |
| BAP1 | 2763.40229891045 | 0.0997348821351128 | 0.0225926067443927 | 4.41449201783084 | 1,01E-05 | 0.00432443369002112 | High confidence DEG |
| TMEM39A | 698.417070432957 | -0.214902607739641 | 0.0487763752706195 | -4.40587490454806 | 1,05E-05 | 0.00432443369002112 | High confidence DEG |
| KLHL9 | 1925.85184812329 | -0.229463926313057 | 0.0521045722263999 | -4.40391152845496 | 1,06E-05 | 0.00432443369002112 | High confidence DEG |
| RACGAP1 | 462.070576594575 | -0.156147269144746 | 0.0354828569407279 | -4.40063970625537 | 1,08E-05 | 0.00432443369002112 | High confidence DEG |
| FAM98A | 501.816068346941 | -0.232309197837174 | 0.0529032293148254 | -4.3912101557111 | 1,13E-05 | 0.00432443369002112 | High confidence DEG |
| LARP4 | 857.429597883705 | -0.245560480858223 | 0.0559480396923483 | -4.38908105107045 | 1,14E-05 | 0.00432443369002112 | High confidence DEG |
| NSUN4 | 654.661141118014 | -0.163604689653294 | 0.0372885772570649 | -4.38752834481763 | 1,15E-05 | 0.00432443369002112 | High confidence DEG |
| TNS1 | 537.649389007716 | 0.591535258052658 | 0.134901581556282 | 4.38493938490901 | 1,16E-05 | 0.00432443369002112 | High confidence DEG |
| TMEM119 | 36.1276767158452 | 0.85463345636051 | 0.194922830034627 | 4.38447079907822 | 1,16E-05 | 0.00432443369002112 | High confidence DEG |
| ZNF497-AS1 | 11.2618821324391 | 0.517814683496747 | 0.118103798965412 | 4.38440327942707 | 1,16E-05 | 0.00432443369002112 | High confidence DEG |
| CDC27 | 1810.43385372937 | -0.134099823359078 | 0.0306050290460902 | -4.38162705734171 | 1,18E-05 | 0.00432443369002112 | High confidence DEG |
| RP11-589P10.5 | 71.5192284019083 | 0.302959598185417 | 0.0695084089419352 | 4.35860355311108 | 1,31E-05 | 0.00468507824656358 | High confidence DEG |
| GFM1 | 1631.60589120781 | -0.127820388909102 | 0.0293807067326676 | -4.35048721162933 | 1,36E-05 | 0.00468507824656358 | High confidence DEG |
| ARMC8 | 1971.27359377606 | -0.109329847976151 | 0.0251677270094815 | -4.34404934283348 | 1,40E-05 | 0.00468507824656358 | High confidence DEG |
| IGLV2-18 | 111.824854877939 | -0.809727085566765 | 0.186419976187049 | -4.34356393627208 | 1,40E-05 | 0.00468507824656358 | High confidence DEG |
| IFI27 | 19.1388318264248 | 1.12929179974785 | 0.260092132458028 | 4.34189142545511 | 1,41E-05 | 0.00468507824656358 | High confidence DEG |
| NUP43 | 653.701098034441 | -0.29313629127622 | 0.0675265086522519 | -4.34105504825984 | 1,42E-05 | 0.00468507824656358 | High confidence DEG |
| RBM18 | 1203.37730087618 | -0.153944160080843 | 0.0355433736680678 | -4.33116342636733 | 1,48E-05 | 0.00475115131981057 | High confidence DEG |
| PIP5K1A | 1779.7283952868 | -0.144738663886905 | 0.0334455513593951 | -4.32759090533717 | 1,51E-05 | 0.00475115131981057 | High confidence DEG |
| CHRM4 | 35.9951595254957 | 0.831400442087958 | 0.192131843977519 | 4.32723917533024 | 1,51E-05 | 0.00475115131981057 | High confidence DEG |
| LINC02863 | 17.5679292944992 | 0.345364143539743 | 0.0799064827212162 | 4.32210418702417 | 1,55E-05 | 0.00478712967784272 | High confidence DEG |
| PMS2P4 | 42.5268426316224 | 0.260735415021874 | 0.0603873156631899 | 4.31771825189458 | 1,58E-05 | 0.00480475509766239 | High confidence DEG |
| FBXO5 | 281.496998893136 | -0.181102946657229 | 0.0419754202980461 | -4.31449989949615 | 1,60E-05 | 0.00480475509766239 | High confidence DEG |
| DBT | 515.685926314362 | -0.314089017271146 | 0.0728633254000988 | -4.31065993140521 | 1,63E-05 | 0.00481599100779909 | High confidence DEG |
| FAM200A | 179.440333111402 | -0.39357377870361 | 0.0913758382993709 | -4.30719746082285 | 1,65E-05 | 0.00482003083109773 | High confidence DEG |
| SOX6 | 11.5450474472064 | 0.873955410905993 | 0.204438595549193 | 4.27490420073688 | 1,91E-05 | 0.0053828242467578 | High confidence DEG |
| PIGHP1 | 14.5249778824914 | 0.636482099405774 | 0.148899639435205 | 4.27457112602843 | 1,92E-05 | 0.0053828242467578 | High confidence DEG |
| GFM2 | 549.338747783849 | -0.214878809499963 | 0.0502865731613427 | -4.27308515954212 | 1,93E-05 | 0.0053828242467578 | High confidence DEG |
| SPTA1 | 15.4967192600872 | 0.757899017816108 | 0.17764562446242 | 4.26635342192984 | 1,99E-05 | 0.00547070632242053 | High confidence DEG |
| NIFK | 715.260525857865 | -0.149537419649413 | 0.0351081055738399 | -4.25934174474051 | 2,05E-05 | 0.00549983058390612 | High confidence DEG |
| GABPB1-IT1 | 794.56930425337 | 0.216190332995109 | 0.0507602608982282 | 4.25904692311492 | 2,05E-05 | 0.00549983058390612 | High confidence DEG |
| CSRNP2 | 586.93501202078 | -0.195006860324485 | 0.0458382570956579 | -4.25423811201053 | 2,10E-05 | 0.00554442913253746 | High confidence DEG |
| ORC5 | 474.101612031703 | -0.147592624542094 | 0.0347285930469038 | -4.24988781845432 | 2,14E-05 | 0.00557882845045852 | High confidence DEG |
| AREG | 59.3128018553527 | 1.0590012610591 | 0.250140363724915 | 4.2336280530227 | 2,30E-05 | 0.0059201968256495 | High confidence DEG |
| CSE1L | 1749.65912248765 | -0.197286833444673 | 0.0466516853166583 | -4.22893261209207 | 2,35E-05 | 0.00595817390339431 | High confidence DEG |
| SLF1 | 308.299804896387 | -0.304287255555775 | 0.0719964333483188 | -4.22642124622525 | 2,37E-05 | 0.00595817390339431 | High confidence DEG |
| METTL18 | 327.674578306247 | -0.251962132930955 | 0.0597123832611924 | -4.21959598947556 | 2,45E-05 | 0.00596462480136506 | High confidence DEG |
| NPLOC4 | 4588.98124302584 | 0.122482077488448 | 0.029047763053883 | 4.2165752061956 | 2,48E-05 | 0.00596462480136506 | High confidence DEG |
| HSPA4 | 3678.62359389875 | -0.128425092000905 | 0.0304769267864294 | -4.2138465239904 | 2,51E-05 | 0.00596462480136506 | High confidence DEG |
| INTS7 | 722.545161153743 | -0.185901768819081 | 0.0441280543213457 | -4.21277964048272 | 2,52E-05 | 0.00596462480136506 | High confidence DEG |
| TIGD2 | 280.174061049021 | -0.173747887116717 | 0.0413014905028857 | -4.20681880971286 | 2,59E-05 | 0.00596462480136506 | High confidence DEG |
| RNF34 | 1228.02619025465 | -0.230424521670401 | 0.0548129083552852 | -4.203836807506 | 2,62E-05 | 0.00596462480136506 | High confidence DEG |
| RP5-1198O20.4 | 5.51530545103918 | 1.02118596035109 | 0.243081974417226 | 4.20099418230957 | 2,66E-05 | 0.00596462480136506 | High confidence DEG |
| CAND1 | 3734.40613041176 | -0.168036427235541 | 0.0400131798660197 | -4.19952695082458 | 2,67E-05 | 0.00596462480136506 | High confidence DEG |
| USO1 | 2736.4435431154 | -0.184461837663173 | 0.0439620454343878 | -4.19593392073799 | 2,72E-05 | 0.00596462480136506 | High confidence DEG |
| METTL6 | 360.058263606668 | -0.128767693716325 | 0.0306962585316913 | -4.19489865787334 | 2,73E-05 | 0.00596462480136506 | High confidence DEG |
| ATF2 | 1610.48354971 | -0.191492599938686 | 0.0456528978694777 | -4.19453329088037 | 2,73E-05 | 0.00596462480136506 | High confidence DEG |
| SLC17A5 | 577.619023690764 | -0.169025147274711 | 0.0402994448992646 | -4.19423016116521 | 2,74E-05 | 0.00596462480136506 | High confidence DEG |
| EED | 594.9318113748 | -0.176394114404315 | 0.0421366775236137 | -4.18623690264763 | 2,84E-05 | 0.00611133697944621 | High confidence DEG |
| RSPRY1 | 1018.14509060597 | -0.189488091337454 | 0.0453193001357375 | -4.18117867597054 | 2,90E-05 | 0.0061817170439774 | High confidence DEG |
| APPBP2 | 1365.53565582478 | -0.138499867428905 | 0.0331830314070352 | -4.17381600041343 | 3,00E-05 | 0.00626557231734799 | High confidence DEG |
| RINT1 | 383.53416286664 | -0.197321233676058 | 0.0472821477298358 | -4.17327137513986 | 3,00E-05 | 0.00626557231734799 | High confidence DEG |
| ITLN1 | 43.5942509266324 | 0.464252932047894 | 0.111489262752854 | 4.16410442211853 | 3,13E-05 | 0.00641794873346625 | High confidence DEG |
| CLIC1P1 | 4.49481305292059 | -0.76503644431313 | 0.183768560478707 | -4.16304313599809 | 3,14E-05 | 0.00641794873346625 | High confidence DEG |
| UBQLN1 | 5561.51854686572 | -0.0996632456882708 | 0.0240438676941583 | -4.1450588131661 | 3,40E-05 | 0.00687218184305181 | High confidence DEG |
| ALKBH1 | 329.416523476364 | -0.26139692540266 | 0.0631248669200291 | -4.14095012245832 | 3,46E-05 | 0.00692578814210773 | High confidence DEG |
| EARS2 | 436.136671241974 | -0.154328603465949 | 0.0373978205929922 | -4.12667372105817 | 3,68E-05 | 0.00722461764613134 | High confidence DEG |
| CIRBP | 7073.10569901455 | 0.260975282047272 | 0.0632414036729715 | 4.12665227035133 | 3,68E-05 | 0.00722461764613134 | High confidence DEG |
| BIN2 | 13629.8256119935 | 0.139095268001652 | 0.0337700565681194 | 4.11889354467249 | 3,81E-05 | 0.00739893792751059 | High confidence DEG |
| BBS12 | 124.979398008321 | -0.339998622747634 | 0.0826021559045378 | -4.11609865413883 | 3,85E-05 | 0.00741647443268851 | High confidence DEG |
| RP11-244H3.5 | 58.2922024659531 | 0.471885602401091 | 0.1148082265455 | 4.11020722643143 | 3,95E-05 | 0.00753510621046172 | High confidence DEG |
| RP11-463O12.5 | 85.2414945497916 | 0.607222700013181 | 0.148013947966595 | 4.10246945206965 | 4,09E-05 | 0.00758623631933009 | High confidence DEG |
| CDPF1P1 | 30.5547684102514 | 0.345920980298637 | 0.0843706480986186 | 4.10001568192649 | 4,13E-05 | 0.00758623631933009 | High confidence DEG |
| LRR1 | 252.352694679413 | -0.179457780223283 | 0.0437793578361951 | -4.09914144686047 | 4,15E-05 | 0.00758623631933009 | High confidence DEG |
| IFNG-AS1 | 88.3023389265742 | -0.531897424257298 | 0.129759098794262 | -4.09911466093521 | 4,15E-05 | 0.00758623631933009 | High confidence DEG |
| INTS13 | 621.597937884031 | -0.243278800797535 | 0.0594251093539184 | -4.09387216014427 | 4,24E-05 | 0.00758623631933009 | High confidence DEG |
| ZNF295-AS1 | 15.37514475519 | 0.63472062763036 | 0.155084298281975 | 4.09274591084847 | 4,26E-05 | 0.00758623631933009 | High confidence DEG |
| PDE12 | 1048.06203723695 | -0.187589835083001 | 0.0458374733945903 | -4.09249945929918 | 4,27E-05 | 0.00758623631933009 | High confidence DEG |
| OR2L1P | 3.90712598515363 | 0.810106714437103 | 0.198020081946964 | 4.09103312387314 | 4,29E-05 | 0.00758623631933009 | High confidence DEG |
| RP11-398C13.8 | 538.539030189803 | -0.175793304431027 | 0.0429894668393006 | -4.089218065629 | 4,33E-05 | 0.00758623631933009 | High confidence DEG |
| RP11-48B3.5 | 41.6346522681121 | 0.43015214631348 | 0.105379264130332 | 4.08194296917346 | 4,47E-05 | 0.00758623631933009 | High confidence DEG |
| CD83 | 818.721988580169 | 0.467278157547508 | 0.114531855453986 | 4.07989686096755 | 4,51E-05 | 0.00758623631933009 | High confidence DEG |
| HARS2 | 914.61351721997 | -0.123521342071578 | 0.0302794708224243 | -4.07937585157865 | 4,52E-05 | 0.00758623631933009 | High confidence DEG |
| IGKV2D-24 | 22.2058034863149 | -0.802922066283981 | 0.196940283826146 | -4.07698237600175 | 4,56E-05 | 0.00758623631933009 | High confidence DEG |
| TAF2 | 1095.6688653833 | -0.150319012304918 | 0.0368785469385166 | -4.07605572300686 | 4,58E-05 | 0.00758623631933009 | High confidence DEG |
| YAE1 | 128.373219952483 | -0.260512789809941 | 0.0639233040719633 | -4.07539618910597 | 4,59E-05 | 0.00758623631933009 | High confidence DEG |
| PPP2R1B | 836.146986828425 | -0.148833971907138 | 0.0365534395031604 | -4.07168173310393 | 4,67E-05 | 0.00758623631933009 | High confidence DEG |
| AURKA | 115.324735788129 | -0.287479004239371 | 0.070617744345715 | -4.07091740047649 | 4,68E-05 | 0.00758623631933009 | High confidence DEG |
| SCYL2P1 | 36.7027019817693 | 0.533640680618615 | 0.131111348949508 | 4.07013340106909 | 4,70E-05 | 0.00758623631933009 | High confidence DEG |
| FAM98B | 748.261267091479 | -0.232305282570531 | 0.0570813844854492 | -4.06972053436701 | 4,71E-05 | 0.00758623631933009 | High confidence DEG |
| DUSP2 | 2100.89498939943 | 0.64320548683867 | 0.158122111177754 | 4.06777699872478 | 4,75E-05 | 0.00758807899976154 | High confidence DEG |
| YIPF4 | 1178.61754871985 | -0.287849208262405 | 0.0709554412662845 | -4.05676017406688 | 4,98E-05 | 0.00779063156887236 | High confidence DEG |
| RPP38 | 284.797130802762 | -0.190989057278545 | 0.0470992637361214 | -4.05503275695733 | 5,01E-05 | 0.00779063156887236 | High confidence DEG |
| PPIL4 | 598.121756070474 | -0.263172974468565 | 0.0649065048465818 | -4.05464714346616 | 5,02E-05 | 0.00779063156887236 | High confidence DEG |
| GABPB1-AS1 | 350.863513657394 | 0.502265700042117 | 0.123887254974233 | 4.05421607046329 | 5,03E-05 | 0.00779063156887236 | High confidence DEG |
| ZBTB11 | 1689.18856095505 | -0.209858382472784 | 0.0518292665026732 | -4.04903246049141 | 5,14E-05 | 0.00790344643960934 | High confidence DEG |
| RAB33B | 601.050886553814 | -0.203483792401544 | 0.0503215753441921 | -4.04366896325771 | 5,26E-05 | 0.00790766556651812 | High confidence DEG |
| SAR1B | 1349.5054434877 | -0.11164665437575 | 0.0276313222054845 | -4.0405831304587 | 5,33E-05 | 0.00790766556651812 | High confidence DEG |
| IGKV2-24 | 220.108923465075 | -0.638740527810944 | 0.158119665117685 | -4.03960207818265 | 5,35E-05 | 0.00790766556651812 | High confidence DEG |
| ERGIC2 | 715.282840719351 | -0.287035430322899 | 0.071065369988283 | -4.03903378495355 | 5,37E-05 | 0.00790766556651812 | High confidence DEG |
| SRSF5 | 15625.8339796295 | 0.294152710656543 | 0.0728812842829807 | 4.03605278845551 | 5,44E-05 | 0.00790766556651812 | High confidence DEG |
| CTC-378H22.2 | 435.302995689918 | 0.260507295748146 | 0.0645517066251096 | 4.0356376208776 | 5,45E-05 | 0.00790766556651812 | High confidence DEG |
| HPS1 | 3221.141483668 | 0.150380225579373 | 0.0372635433272426 | 4.035585780417 | 5,45E-05 | 0.00790766556651812 | High confidence DEG |
| TNFRSF17 | 239.859174974792 | -0.663205841329686 | 0.164375762199877 | -4.03469363398748 | 5,47E-05 | 0.00790766556651812 | High confidence DEG |
| CBX3P2 | 59.4994552452249 | 0.341476829695491 | 0.0846794235749383 | 4.03258330393919 | 5,52E-05 | 0.00790766556651812 | High confidence DEG |
| UFL1 | 1570.51422336573 | -0.1899732662886 | 0.0471234283810009 | -4.03139739224052 | 5,54E-05 | 0.00790766556651812 | High confidence DEG |
| PWWP2A | 1061.3639076421 | -0.112433560049123 | 0.0279174708966322 | -4.02735478673628 | 5,64E-05 | 0.00795514910829724 | High confidence DEG |
| ETNK1 | 1786.56730604879 | -0.274518450859345 | 0.0681757157375786 | -4.02663100620783 | 5,66E-05 | 0.00795514910829724 | High confidence DEG |
| PLCL2 | 5427.530064821 | 0.124146701726354 | 0.0308697955846945 | 4.02162370611573 | 5,78E-05 | 0.00806897387889394 | High confidence DEG |
| UBE3A | 2550.43828656668 | -0.114311282432261 | 0.0284410902933949 | -4.01922996808629 | 5,84E-05 | 0.00809438303579663 | High confidence DEG |
| FAM53B | 4611.5841035439 | 0.195027668265469 | 0.0485782450883937 | 4.01471209819527 | 5,95E-05 | 0.00816158842174392 | High confidence DEG |
| SRP54 | 1758.97009369505 | -0.169333371230752 | 0.0421856306754567 | -4.01400591906451 | 5,97E-05 | 0.00816158842174392 | High confidence DEG |
| LINC00667 | 1224.97798377391 | 0.196474002765635 | 0.0489691739166403 | 4.01219761436226 | 6,02E-05 | 0.0081680016073696 | High confidence DEG |
| MINPP1 | 390.2380152463 | -0.230182230256103 | 0.0573936415279203 | -4.01058765619753 | 6,06E-05 | 0.0081680016073696 | High confidence DEG |
| ZBED8 | 18.4264691663582 | -0.558182871201634 | 0.139485970753923 | -4.00171334926838 | 6,29E-05 | 0.00836929707126637 | High confidence DEG |
| AIF1L | 5.5436373811377 | -0.920812610306423 | 0.230108779478235 | -4.00164049539674 | 6,29E-05 | 0.00836929707126637 | High confidence DEG |
| DNAJA2 | 2612.67796080862 | -0.10719772254734 | 0.0268029316021309 | -3.99947752501885 | 6,35E-05 | 0.00838984399948712 | High confidence DEG |
| TCP1 | 3913.23480614654 | -0.099608353128388 | 0.0249167061749932 | -3.99765331857372 | 6,40E-05 | 0.00839875322268951 | High confidence DEG |
| CTD-2319I12.10 | 18.0870565243813 | -0.591847794594778 | 0.148231048652041 | -3.99273836336467 | 6,53E-05 | 0.00847669137493331 | High confidence DEG |
| SREK1IP1 | 912.382997543188 | -0.159780266806381 | 0.0400479947092372 | -3.9897195344347 | 6,62E-05 | 0.00847669137493331 | High confidence DEG |
| AC015971.2 | 18.0214355329765 | 0.689290763510904 | 0.172799344939947 | 3.98896629932511 | 6,64E-05 | 0.00847669137493331 | High confidence DEG |
| SRPRB | 1086.53672651701 | -0.191526171200729 | 0.048027195464198 | -3.98786915100013 | 6,67E-05 | 0.00847669137493331 | High confidence DEG |
| ALG2 | 835.568636920644 | -0.171631100764203 | 0.0430396722094385 | -3.98774181943154 | 6,67E-05 | 0.00847669137493331 | High confidence DEG |
| TEX10 | 754.563228561263 | -0.138509509132668 | 0.0347800064510074 | -3.98244633242891 | 6,82E-05 | 0.00850718612148861 | High confidence DEG |
| LINC01259 | 33.9964827497464 | 0.595948615611644 | 0.149686541676022 | 3.98131060373819 | 6,85E-05 | 0.00850718612148861 | High confidence DEG |
| STBD1 | 8.82155087054909 | -0.432179460242903 | 0.108588828264191 | -3.97996246162112 | 6,89E-05 | 0.00850718612148861 | High confidence DEG |
| GLS | 3017.87862474123 | -0.212901625120051 | 0.0535089557544769 | -3.97880358751419 | 6,93E-05 | 0.00850718612148861 | High confidence DEG |
| PNRC2 | 4212.03469959519 | -0.13116555573995 | 0.0329717221842666 | -3.97812267757548 | 6,95E-05 | 0.00850718612148861 | High confidence DEG |
| LRRC40 | 443.084906896501 | -0.244273708549896 | 0.0614073553524794 | -3.97792263072983 | 6,95E-05 | 0.00850718612148861 | High confidence DEG |
| DHX15 | 3721.09944516099 | -0.158069990137196 | 0.0397572884367838 | -3.97587452143613 | 7,01E-05 | 0.00852810155631801 | High confidence DEG |
| NDUFV2 | 29.451569681044 | 0.606326933445838 | 0.152671782297033 | 3.97144072285859 | 7,14E-05 | 0.00863258639073613 | High confidence DEG |
| GTPBP8 | 353.30010641989 | -0.13076414688533 | 0.0329374807293468 | -3.97007129840447 | 7,19E-05 | 0.00863258639073613 | High confidence DEG |
| MTERF1 | 393.292044289848 | -0.159455253005113 | 0.0401990447383236 | -3.96664284047271 | 7,29E-05 | 0.00868102702217435 | High confidence DEG |
| ZNF816 | 346.013965031449 | -0.307548888248798 | 0.0775489978071856 | -3.96586541341867 | 7,31E-05 | 0.00868102702217435 | High confidence DEG |
| MTREX | 2430.2112293682 | -0.134384088742861 | 0.0339109664575804 | -3.96285045166624 | 7,41E-05 | 0.00872336268087208 | High confidence DEG |
| PRELID2 | 65.7206474652867 | 0.24901107408791 | 0.0628519966374115 | 3.96186417950151 | 7,44E-05 | 0.00872336268087208 | High confidence DEG |
| RP11-12J10.3 | 8.98947044437064 | 0.461372465869977 | 0.116717158917411 | 3.95291035310794 | 7,72E-05 | 0.00900317083648896 | High confidence DEG |
| NEK2 | 34.6340943077732 | -0.564911046054794 | 0.143168980733899 | -3.94576425115973 | 7,95E-05 | 0.00911150971171205 | High confidence DEG |
| PPP4R3B | 4481.30429707717 | -0.214720858923819 | 0.0544459559229793 | -3.94374302524081 | 8,02E-05 | 0.00911150971171205 | High confidence DEG |
| JADE3 | 93.8726752782106 | -0.253612501813172 | 0.0643269229904468 | -3.94255608729857 | 8,06E-05 | 0.00911150971171205 | High confidence DEG |
| RP5-882C2.2 | 78.2016605985096 | 0.624819803033843 | 0.158515756908026 | 3.94168892242288 | 8,09E-05 | 0.00911150971171205 | High confidence DEG |
| WDR75 | 860.012686391799 | -0.222477225739888 | 0.0564497102166653 | -3.94115797735677 | 8,11E-05 | 0.00911150971171205 | High confidence DEG |
| NOL11 | 1328.30210562716 | -0.229414206703635 | 0.0582309367804973 | -3.93973065500246 | 8,16E-05 | 0.00911150971171205 | High confidence DEG |
| ZNF550 | 391.469662295221 | -0.304256947595333 | 0.077229721451001 | -3.93963543929615 | 8,16E-05 | 0.00911150971171205 | High confidence DEG |
| POLR1B | 1241.51305699848 | -0.179362063347218 | 0.0455437677510143 | -3.93823506934653 | 8,21E-05 | 0.00911150971171205 | High confidence DEG |
| ZNF213-AS1 | 268.810428471791 | 0.26104114556005 | 0.0663188857291678 | 3.93615095745255 | 8,28E-05 | 0.00911150971171205 | High confidence DEG |
| BRAP | 1107.74983556541 | -0.0959362927694228 | 0.0243940955574966 | -3.93276694941618 | 8,40E-05 | 0.00911150971171205 | High confidence DEG |
| THAP6 | 272.003941059141 | -0.287240591002198 | 0.0730621785836207 | -3.93145395566664 | 8,44E-05 | 0.00911150971171205 | High confidence DEG |
| GLDC | 105.053820965754 | -0.759750875594201 | 0.193334560917321 | -3.92972095619834 | 8,50E-05 | 0.00911150971171205 | High confidence DEG |
| RFWD3 | 1271.59012358561 | -0.112268564885094 | 0.0285697088677199 | -3.92963629433212 | 8,51E-05 | 0.00911150971171205 | High confidence DEG |
| SIRT1 | 1671.8649582338 | -0.187724598050061 | 0.047800996597262 | -3.92721096657666 | 8,59E-05 | 0.00911150971171205 | High confidence DEG |
| ATAD1 | 1394.0782952799 | -0.17259343685067 | 0.043964693489763 | -3.92572819575912 | 8,65E-05 | 0.00911150971171205 | High confidence DEG |
| CTD-2553L13.10 | 41.9155196572915 | 0.458110626960005 | 0.116708971874244 | 3.92523916202112 | 8,66E-05 | 0.00911150971171205 | High confidence DEG |
| SLC30A7 | 868.733627985571 | -0.21170485711181 | 0.0539453229923278 | -3.92443395216893 | 8,69E-05 | 0.00911150971171205 | High confidence DEG |
| RRM1 | 1252.36867602913 | -0.166206625924122 | 0.0423569244848342 | -3.92395406289783 | 8,71E-05 | 0.00911150971171205 | High confidence DEG |
| PTER | 661.623120107676 | -0.177078047196414 | 0.0451442578183309 | -3.92249326390546 | 8,76E-05 | 0.00911150971171205 | High confidence DEG |
| C1orf109 | 325.957430224659 | -0.201586214804311 | 0.0514102776827952 | -3.9211267452806 | 8,81E-05 | 0.00911150971171205 | High confidence DEG |
| FPR3 | 321.956266396647 | -0.435056743595968 | 0.110997516946287 | -3.91951780152432 | 8,87E-05 | 0.00911150971171205 | High confidence DEG |
| EIF4E | 639.213295726184 | -0.133894135678508 | 0.0341665641421457 | -3.91886451097213 | 8,90E-05 | 0.00911150971171205 | High confidence DEG |
| MRPL50 | 826.343807745325 | -0.249517619443185 | 0.0636964683121683 | -3.91729127304722 | 8,95E-05 | 0.00911150971171205 | High confidence DEG |
| SNORD3A | 5.38508296616367 | -0.825910998001853 | 0.21089212584647 | -3.91627233443091 | 8,99E-05 | 0.00911150971171205 | High confidence DEG |
| ASTE1 | 428.716086952468 | -0.232846646933061 | 0.0594639750160654 | -3.91575986755262 | 9,01E-05 | 0.00911150971171205 | High confidence DEG |
| CUL2 | 1353.06264231254 | -0.10834699963884 | 0.0276832745299013 | -3.91380721676593 | 9,09E-05 | 0.00911150971171205 | High confidence DEG |
| TDP2 | 1621.33447617084 | -0.110386123139084 | 0.0282071229456765 | -3.91341305356362 | 9,10E-05 | 0.00911150971171205 | High confidence DEG |
| CUL5 | 893.920695902635 | -0.183512443572442 | 0.046893326235967 | -3.9134021470136 | 9,10E-05 | 0.00911150971171205 | High confidence DEG |
| NAA15 | 1517.66329840584 | -0.16061996530651 | 0.0410566429530775 | -3.91215534816322 | 9,15E-05 | 0.00911266787921165 | High confidence DEG |
| POLR2D | 850.318493396967 | -0.100301719307397 | 0.0256468871292837 | -3.91087303506992 | 9,20E-05 | 0.00911538321999269 | High confidence DEG |
| KIFC3 | 224.883864876621 | 0.326434861515674 | 0.0834982009119449 | 3.90948377270936 | 9,25E-05 | 0.00912235072231682 | High confidence DEG |
| TGFBRAP1 | 2228.86442556734 | 0.154158435322034 | 0.039467713255428 | 3.90593785670704 | 9,39E-05 | 0.00912951528009784 | High confidence DEG |
| PPP2R2A | 1400.43161240334 | -0.142066896959884 | 0.0363737154270468 | -3.90575709112866 | 9,39E-05 | 0.00912951528009784 | High confidence DEG |
| WDR12 | 519.370048106256 | -0.122165340625375 | 0.0312849586193681 | -3.90492255756874 | 9,43E-05 | 0.00912951528009784 | High confidence DEG |
| RBBP5 | 1230.11462550751 | -0.226505154737297 | 0.0580181651126131 | -3.90403857649843 | 9,46E-05 | 0.00912951528009784 | High confidence DEG |
| CGAS | 538.147048989426 | -0.293853651093703 | 0.0752978304335674 | -3.90255136704051 | 9,52E-05 | 0.00912951528009784 | High confidence DEG |
| RORA-AS1 | 14.1106228375363 | 0.35501431710784 | 0.0909784076994029 | 3.90218213403805 | 9,53E-05 | 0.00912951528009784 | High confidence DEG |
| UBA6 | 1508.12610757085 | -0.177730148449389 | 0.0455689480284783 | -3.90024690362211 | 9,61E-05 | 0.00915855816618098 | High confidence DEG |
| MZB1 | 609.50361512423 | -0.602705470326407 | 0.15464102979728 | -3.89744863388778 | 9,72E-05 | 0.00922066717865863 | High confidence DEG |
| HOXA-AS2 | 9.67935277229045 | 0.560624945743763 | 0.143946180521248 | 3.89468441408911 | 9,83E-05 | 0.00926235893139214 | High confidence DEG |
| ARIH2 | 2892.28403619133 | 0.126879931576217 | 0.0325830488964367 | 3.89404723847351 | 9,86E-05 | 0.00926235893139214 | High confidence DEG |
| HSPD1 | 4477.32764273815 | -0.125065500524328 | 0.0321538314187079 | -3.8895986887448 | 0,00010041 | 0.00937392298646738 | High confidence DEG |
| ITFG2 | 995.110198177117 | 0.114843203400302 | 0.029539026459123 | 3.88784659369953 | 0,00010114 | 0.00937392298646738 | High confidence DEG |
| WDR48 | 2005.57263233032 | -0.122299524925471 | 0.0314802251643968 | -3.88496347426981 | 0,00010235 | 0.00937392298646738 | High confidence DEG |
| VPS37A | 757.581465853717 | -0.167326149553987 | 0.043078264324399 | -3.88423610324559 | 0,00010265 | 0.00937392298646738 | High confidence DEG |
| ELL2 | 748.803866680305 | -0.271368356792052 | 0.0698949809142819 | -3.88251564336006 | 0,00010338 | 0.00937392298646738 | High confidence DEG |
| RIOK1 | 673.184994454427 | -0.119276777309836 | 0.0307326660021501 | -3.88110739567767 | 0,00010398 | 0.00937392298646738 | High confidence DEG |
| RP11-122G18.13 | 11.6083541335856 | 0.392025971461815 | 0.101013247674533 | 3.88093621863273 | 0,00010406 | 0.00937392298646738 | High confidence DEG |
| TAF5 | 268.953594765582 | -0.185791302826955 | 0.04789005124368 | -3.87953860983754 | 0,00010466 | 0.00937392298646738 | High confidence DEG |
| ADTRP | 217.00567985947 | 0.404893897920679 | 0.104380141761995 | 3.87903188370743 | 0,00010487 | 0.00937392298646738 | High confidence DEG |
| RASSF5 | 14547.8612338791 | 0.08209158184858 | 0.0211646533295364 | 3.87871138593216 | 0,00010501 | 0.00937392298646738 | High confidence DEG |
| WDCP | 400.922242273104 | -0.238485924496281 | 0.0614901850939142 | -3.87843887820536 | 0,00010513 | 0.00937392298646738 | High confidence DEG |
| SPOP | 2363.63715761212 | -0.159878264051871 | 0.0412308896700002 | -3.87763313698757 | 0,00010548 | 0.00937392298646738 | High confidence DEG |
| CRNKL1 | 1309.12856312758 | -0.206984834965132 | 0.0534080904471822 | -3.8755333364676 | 0,00010639 | 0.00937392298646738 | High confidence DEG |
| DDX20 | 818.701990310894 | -0.15352223149349 | 0.0396132228082495 | -3.87552995212294 | 0,00010639 | 0.00937392298646738 | High confidence DEG |
| SYT17 | 56.1140232309283 | 0.34879049077727 | 0.0900394344345897 | 3.8737525726092 | 0,00010717 | 0.00940080607713462 | High confidence DEG |
| CCNB1 | 162.957465126888 | -0.308178310636807 | 0.079674276345093 | -3.86797753018797 | 0,00010974 | 0.00957043644914283 | High confidence DEG |
| ZNF2 | 215.078292961991 | -0.301599871157395 | 0.0779950919974178 | -3.86690833273689 | 0,00011022 | 0.00957043644914283 | High confidence DEG |
| KIAA1586 | 194.649903806572 | -0.195971913983009 | 0.0506888008381239 | -3.86617775016716 | 0,00011055 | 0.00957043644914283 | High confidence DEG |
| GS1-124K5.12 | 602.782158751653 | 0.178288345503582 | 0.046129694060097 | 3.86493665601405 | 0,00011112 | 0.00957740825188339 | High confidence DEG |
| ATAD2 | 627.45101898481 | -0.238014342773173 | 0.0616170869314594 | -3.86279771774884 | 0,0001121 | 0.00957902534581021 | High confidence DEG |
| RIOK2 | 775.917216449903 | -0.127764942208281 | 0.0330758903471119 | -3.86278164752221 | 0,0001121 | 0.00957902534581021 | High confidence DEG |
| DCAF1 | 1387.87726042761 | -0.26046225840985 | 0.0674802864574644 | -3.85982739676172 | 0,00011347 | 0.0096539597921623 | High confidence DEG |
| KLHL24 | 1839.47042431783 | -0.230966681752372 | 0.0598682693375554 | -3.85791479038941 | 0,00011436 | 0.00967325875546887 | High confidence DEG |
| ZNF845 | 714.053110197847 | -0.25134575481044 | 0.0651739091318824 | -3.85653949806556 | 0,000115 | 0.00967325875546887 | High confidence DEG |
| CYP2U1 | 428.481255605507 | -0.180473150075177 | 0.0468006353120445 | -3.85621154225508 | 0,00011516 | 0.00967325875546887 | High confidence DEG |
| LZTS3 | 87.6691987711428 | 0.38282585070959 | 0.0993217499569151 | 3.85440098342666 | 0,00011601 | 0.00970400048227815 | High confidence DEG |
| NOC3L | 471.666672843541 | -0.228487587427369 | 0.0593067748461592 | -3.85263889361818 | 0,00011685 | 0.0097208038958189 | High confidence DEG |
| USP14 | 1645.45079501278 | -0.170445425575252 | 0.0442546347697552 | -3.85147061911216 | 0,00011741 | 0.0097208038958189 | High confidence DEG |
| ZNF765 | 492.557158203648 | -0.197156291997869 | 0.0512073117656251 | -3.85015899487654 | 0,00011804 | 0.0097208038958189 | High confidence DEG |
| LMBRD2 | 644.517527966818 | -0.197070502554506 | 0.0511887369172835 | -3.84988015767911 | 0,00011818 | 0.0097208038958189 | High confidence DEG |
| MED18 | 320.741036234197 | -0.345471365789942 | 0.0898696588065393 | -3.84413794797676 | 0,00012098 | 0.00991011130518652 | High confidence DEG |
| EFNA3 | 26.9464015825028 | -0.278830864299412 | 0.0726236512491643 | -3.83939473578341 | 0,00012334 | 0.0100093011662178 | High confidence DEG |
| MAP3K4 | 1333.89018486617 | -0.185176782607232 | 0.0482332643019096 | -3.83919241808191 | 0,00012344 | 0.0100093011662178 | Primary DEG |
| PRMT3 | 351.767663659526 | -0.158143831722786 | 0.0411975577683097 | -3.83867006418605 | 0,0001237 | 0.0100093011662178 | Primary DEG |
| PIK3R4 | 1471.31290304888 | -0.254193996081707 | 0.0664245200985725 | -3.826809673664 | 0,00012982 | 0.0104611738467077 | Primary DEG |
| TUBB2A | 94.4828924648608 | 0.736052225852919 | 0.192740255374934 | 3.81888165718724 | 0,00013406 | 0.0107593798638898 | Primary DEG |
| INTS14 | 980.931362985768 | -0.174173407594544 | 0.0456474676194683 | -3.81562037672075 | 0,00013584 | 0.010819363061634 | Primary DEG |
| ZNF189 | 645.306964279417 | -0.199922668635549 | 0.0524234959094214 | -3.81360810009657 | 0,00013695 | 0.010819363061634 | Primary DEG |
| KIF18A | 47.1165707107882 | -0.232637854647115 | 0.0610164908707751 | -3.81270458735182 | 0,00013745 | 0.010819363061634 | Primary DEG |
| AASDHPPT | 1115.69752589878 | -0.188966543170442 | 0.0495722160579432 | -3.81194463748737 | 0,00013788 | 0.010819363061634 | Primary DEG |
| SLC5A10 | 17.7056711810606 | 0.447803049299632 | 0.117484397160634 | 3.81159592356217 | 0,00013807 | 0.010819363061634 | Primary DEG |
| MSH6 | 1356.70250507694 | -0.132551069092984 | 0.0347830201832271 | -3.81079815366069 | 0,00013852 | 0.010819363061634 | Primary DEG |
| PPAT | 310.996395125109 | -0.209203087348042 | 0.0549109479892251 | -3.80986114807366 | 0,00013905 | 0.010819363061634 | Primary DEG |
| CABP4 | 44.7572798632432 | 0.408010253040115 | 0.107099539992974 | 3.80963590568999 | 0,00013917 | 0.010819363061634 | Primary DEG |
| ANKRD27 | 1249.87919337002 | -0.166446203718573 | 0.0437234031419456 | -3.80679891677724 | 0,00014078 | 0.0109014278875448 | Primary DEG |
| ZNF225 | 231.168421607415 | -0.207983942671057 | 0.0546656995814524 | -3.80465162365953 | 0,000142 | 0.0109536398899421 | Primary DEG |
| ZMYM4 | 1738.2699339561 | -0.182358543752994 | 0.0480393894647101 | -3.79602126057317 | 0,00014704 | 0.0112952205957327 | Primary DEG |
| ATP7A | 693.765949836575 | -0.232783736482818 | 0.0613799094754289 | -3.79250700224646 | 0,00014913 | 0.0112952205957327 | Primary DEG |
| TOMM70 | 2344.43111559307 | -0.111465225359537 | 0.0294007067901545 | -3.79124305259436 | 0,0001499 | 0.0112952205957327 | Primary DEG |
| ELOCP19 | 18.90583899323 | 0.460178072549325 | 0.121433902207502 | 3.78953541131364 | 0,00015093 | 0.0112952205957327 | Primary DEG |
| NUP160 | 2328.25729513226 | -0.163764280576932 | 0.0432153937850093 | -3.78948949051897 | 0,00015096 | 0.0112952205957327 | Primary DEG |
| KLF11 | 2416.66977685646 | 0.268373216436367 | 0.0708236554440329 | 3.78931608025294 | 0,00015106 | 0.0112952205957327 | Primary DEG |
| PRR7-AS1 | 32.0209716638752 | 0.464848908251439 | 0.122678194482741 | 3.7891730491423 | 0,00015115 | 0.0112952205957327 | Primary DEG |
| ARMC9 | 16.0163083448985 | -0.399059385950503 | 0.105325193118267 | -3.78883127707545 | 0,00015136 | 0.0112952205957327 | Primary DEG |
| TBCD | 1879.37062123192 | 0.142224040799214 | 0.0375486047284764 | 3.78773171007743 | 0,00015203 | 0.0112952205957327 | Primary DEG |
| PIGV | 441.789993384846 | -0.17772205043989 | 0.0469224890429833 | -3.78756656061207 | 0,00015213 | 0.0112952205957327 | Primary DEG |
| FASTKD2 | 959.265070840377 | -0.135702431853484 | 0.0358654907653553 | -3.78364909994673 | 0,00015455 | 0.0114317663847119 | Primary DEG |
| NDUFAF4 | 314.315897677699 | -0.182331085877456 | 0.0482422825163718 | -3.77948712968917 | 0,00015715 | 0.0114796305395818 | Primary DEG |
| PURA | 963.140460427131 | -0.24714854518388 | 0.0653975096334273 | -3.77917365002463 | 0,00015735 | 0.0114796305395818 | Primary DEG |
| HIBCH | 356.938587342562 | -0.147340788934687 | 0.0389949504052946 | -3.77845816966808 | 0,0001578 | 0.0114796305395818 | Primary DEG |
| SSPN | 50.0603741342618 | -0.471233134333647 | 0.124740452563106 | -3.77770903224236 | 0,00015828 | 0.0114796305395818 | Primary DEG |
| DNAJB13 | 33.6886895450876 | 0.722365270876296 | 0.191295963566665 | 3.776165776883 | 0,00015926 | 0.0114796305395818 | Primary DEG |
| BHLHE41 | 77.46359184897 | -0.381562506406319 | 0.101099619485404 | -3.7741240604907 | 0,00016057 | 0.0114796305395818 | Primary DEG |
| SLC38A11 | 23.6817029047092 | 0.574859889270746 | 0.152343076741657 | 3.77345594933462 | 0,000161 | 0.0114796305395818 | Primary DEG |
| LINC00402 | 696.447913223886 | 0.307040237718145 | 0.0813813254179773 | 3.77285865204549 | 0,00016139 | 0.0114796305395818 | Primary DEG |
| FBXO45 | 561.083867749765 | -0.216482525876613 | 0.0573799311958599 | -3.77279166016555 | 0,00016143 | 0.0114796305395818 | Primary DEG |
| ECT2 | 236.889633676313 | -0.180401762715299 | 0.0478218590409362 | -3.77237034137198 | 0,0001617 | 0.0114796305395818 | Primary DEG |
| YARS2 | 342.574332341724 | -0.151896696154841 | 0.0402710960046486 | -3.7718540398629 | 0,00016204 | 0.0114796305395818 | Primary DEG |
| ARPP19 | 4447.08280121933 | -0.128779833529163 | 0.0341437461945726 | -3.77169607562376 | 0,00016214 | 0.0114796305395818 | Primary DEG |
| LINC00653 | 64.3199784064473 | 0.538679811454037 | 0.142857628304143 | 3.77074586669737 | 0,00016276 | 0.011482424613644 | Primary DEG |
| NUP54 | 1032.4040728425 | -0.146951687547868 | 0.0389872033422788 | -3.7692287455897 | 0,00016375 | 0.0114908512735844 | Primary DEG |
| IL4R | 6012.73719933042 | 0.203944110521494 | 0.0541228011513627 | 3.76817360119875 | 0,00016445 | 0.0114908512735844 | Primary DEG |
| MIOS | 1134.44821980247 | -0.0803914104381279 | 0.0213467852506117 | -3.76597269773088 | 0,0001659 | 0.0114908512735844 | Primary DEG |
| AKT2 | 4316.73635058553 | 0.0912165693049961 | 0.0242246354212401 | 3.76544652659737 | 0,00016625 | 0.0114908512735844 | Primary DEG |
| TTI2 | 303.186894057285 | -0.219001293720282 | 0.0581645618103778 | -3.76520147154634 | 0,00016642 | 0.0114908512735844 | Primary DEG |
| RP11-727F15.12 | 18.8500677730992 | 0.307337022524456 | 0.081638409879089 | 3.76461304157735 | 0,00016681 | 0.0114908512735844 | Primary DEG |
| PTRH2 | 452.408034713675 | -0.190523449149667 | 0.0506167962883402 | -3.76403611292079 | 0,00016719 | 0.0114908512735844 | Primary DEG |
| TRIM4 | 2063.04486758334 | -0.152020633450707 | 0.0403928623936152 | -3.76355188620494 | 0,00016752 | 0.0114908512735844 | Primary DEG |
| RP11-573D15.9 | 58.6031257818053 | 0.531323335647608 | 0.141452612100723 | 3.75619317138712 | 0,00017252 | 0.0117930618507767 | Primary DEG |
| CTD-3074O7.5 | 41.139357498159 | 0.260985269072341 | 0.0695201484000678 | 3.75409539649497 | 0,00017397 | 0.0118513869550458 | Primary DEG |
| CLEC20A | 3.19452734049846 | 0.764434272233386 | 0.203738558691947 | 3.75203533951181 | 0,00017541 | 0.0119082992801912 | Primary DEG |
| HSPA13 | 1320.47597428194 | -0.180302157758608 | 0.0480748181594996 | -3.75044908460003 | 0,00017652 | 0.0119429888503015 | Primary DEG |
| CHORDC1 | 821.125007621875 | -0.251647220892104 | 0.0671272283875881 | -3.74880993803453 | 0,00017768 | 0.0119804164050111 | Primary DEG |
| ZC3HC1 | 326.803312594464 | -0.12103001366144 | 0.0322963776245313 | -3.74747951824508 | 0,00017862 | 0.0120032920174688 | Primary DEG |
| ZNF322 | 119.695862656313 | -0.320122071945875 | 0.0854529124575742 | -3.74618094034899 | 0,00017955 | 0.0120069586377971 | Primary DEG |
| RP11-53B2.3 | 3.38749302455752 | 0.809189236146151 | 0.216031099716426 | 3.74570715609157 | 0,00017989 | 0.0120069586377971 | Primary DEG |
| ATP5MGP1 | 3.4395062651192 | 0.614137220226161 | 0.16407734833013 | 3.74297382592076 | 0,00018186 | 0.0120976411078033 | Primary DEG |
| SIKE1 | 1089.1496827779 | -0.126155005647914 | 0.0337297136946509 | -3.74017422116216 | 0,00018389 | 0.0121922737608256 | Primary DEG |
| MSMO1 | 404.721839390688 | -0.223284774304273 | 0.059757460290258 | -3.73651713475973 | 0,00018659 | 0.0122603027737549 | Primary DEG |
| TXNDC15 | 1222.68851377844 | -0.152971773220523 | 0.0409442429622891 | -3.73609968467155 | 0,0001869 | 0.0122603027737549 | Primary DEG |
| SEC22B | 1608.02395516679 | -0.150878552073863 | 0.0403890422055461 | -3.73563085021969 | 0,00018725 | 0.0122603027737549 | Primary DEG |
| POLR3B | 372.341693235779 | -0.160622237754204 | 0.0430000559100331 | -3.73539602111833 | 0,00018742 | 0.0122603027737549 | Primary DEG |
| NPRL3 | 1232.45784819008 | 0.243086387722949 | 0.0650902786516469 | 3.73460358072685 | 0,00018801 | 0.0122603027737549 | Primary DEG |
| OTULIN | 1346.67617066359 | -0.100444501292012 | 0.0269174426984129 | -3.7315766738099 | 0,00019029 | 0.0123679059484605 | Primary DEG |
| RNF220 | 2299.84046939644 | 0.0737487310635504 | 0.0197882886000068 | 3.72688778470338 | 0,00019386 | 0.0125426569308294 | Primary DEG |
| SH3GLB2 | 1509.74562799643 | 0.15190595410658 | 0.0407700241883495 | 3.72592258971456 | 0,0001946 | 0.0125426569308294 | Primary DEG |
| DLGAP1-AS2 | 30.70388779525 | 0.427375756485932 | 0.114714090545149 | 3.7255733315318 | 0,00019487 | 0.0125426569308294 | Primary DEG |
| SLC30A5 | 1042.34692844779 | -0.141959859887587 | 0.0381875244321098 | -3.71744076105182 | 0,00020125 | 0.0129091860241779 | Primary DEG |
| CTBP1-DT | 790.889657961845 | 0.153314507497234 | 0.0412505448074392 | 3.71666624557125 | 0,00020187 | 0.0129091860241779 | Primary DEG |
| CSTF1 | 1029.31705172842 | -0.146853580382003 | 0.0395271292545745 | -3.7152604591189 | 0,00020299 | 0.0129394301067366 | Primary DEG |
| CHIT1 | 27.6855044999807 | 0.844650007889533 | 0.227418962914103 | 3.71407026514566 | 0,00020395 | 0.0129435904812306 | Primary DEG |
| ALG6 | 450.350187390906 | -0.124694250775297 | 0.0335862819128791 | -3.71265420503369 | 0,0002051 | 0.0129435904812306 | Primary DEG |
| ZNF230 | 259.147383333578 | -0.221510707703136 | 0.0596747747203286 | -3.71196554559723 | 0,00020566 | 0.0129435904812306 | Primary DEG |
| CCDC43 | 363.331300604033 | -0.149740999019462 | 0.0403402901046477 | -3.71194650883807 | 0,00020567 | 0.0129435904812306 | Primary DEG |
| E2F4 | 2943.89816969963 | 0.0827535661542656 | 0.0223067273292968 | 3.70980309808065 | 0,00020742 | 0.0129834356973208 | Primary DEG |
| PRIM2 | 318.30023657242 | -0.160346159654288 | 0.0432492594029307 | -3.707489142425 | 0,00020932 | 0.0129834356973208 | Primary DEG |
| ZNRF2 | 395.971367513182 | -0.175220242792619 | 0.0472631834351071 | -3.70733052785574 | 0,00020946 | 0.0129834356973208 | Primary DEG |
| TERF2 | 1271.01180250515 | -0.104644543549186 | 0.0282313959825078 | -3.70667265671253 | 0,00021 | 0.0129834356973208 | Primary DEG |
| TAF1A | 119.97062709603 | -0.262643307916889 | 0.070859781054683 | -3.70652158400271 | 0,00021013 | 0.0129834356973208 | Primary DEG |
| RAE1 | 1208.47025659189 | -0.103042894780046 | 0.0278014146918109 | -3.70639033740964 | 0,00021023 | 0.0129834356973208 | Primary DEG |
| PPM1D | 786.066593130151 | -0.190475886288356 | 0.0514131461986033 | -3.70480899092556 | 0,00021155 | 0.0130168847401363 | Primary DEG |
| VPS50 | 919.531781188997 | -0.167222659976411 | 0.0451445084139308 | -3.70416393602393 | 0,00021209 | 0.0130168847401363 | Primary DEG |
| FBXO8 | 443.771384159946 | -0.200949320129942 | 0.0542658488901901 | -3.70305310318785 | 0,00021302 | 0.0130336846882329 | Primary DEG |
| RP1-102E24.6 | 4.37588517543567 | 0.785752698995209 | 0.212336165678436 | 3.70051279999641 | 0,00021516 | 0.0130831195122292 | Primary DEG |
| SOX30 | 22.2220402001685 | -0.388302403345778 | 0.104939007938238 | -3.70026752658377 | 0,00021537 | 0.0130831195122292 | Primary DEG |
| PLPP6 | 245.942734470055 | -0.266952285798006 | 0.0721540536440229 | -3.69975451573426 | 0,00021581 | 0.0130831195122292 | Primary DEG |
| CXCR4 | 19241.8069483409 | 0.404649846285973 | 0.109425860803158 | 3.69793614887692 | 0,00021736 | 0.0131369987706584 | Primary DEG |
| NIF3L1 | 649.917565285567 | -0.112500243638218 | 0.0304334360854232 | -3.69660012502179 | 0,00021851 | 0.0131661550776059 | Primary DEG |
| CCNH | 747.488053229548 | -0.104673166972443 | 0.0283377843921663 | -3.69376679291057 | 0,00022096 | 0.0132734568943243 | Primary DEG |
| EVL | 8777.26668039351 | 0.189169386031422 | 0.0512312321516011 | 3.69246215807656 | 0,00022209 | 0.0133014485811539 | Primary DEG |
| MFAP1 | 724.20108727564 | -0.294198538346607 | 0.0796929034632074 | -3.69165290209852 | 0,0002228 | 0.01330365887593 | Primary DEG |
| MFAP3 | 969.861993874855 | -0.226481655763783 | 0.0613682556248839 | -3.69053435620139 | 0,00022378 | 0.0133221721216747 | Primary DEG |
| DARS2 | 574.826934999903 | -0.200043921344429 | 0.0542214354390017 | -3.68938814925834 | 0,00022479 | 0.0133422661112612 | Primary DEG |
| SEH1L | 807.311182147142 | -0.169900761067632 | 0.0460709623580883 | -3.6878057755137 | 0,0002262 | 0.013361421132962 | Primary DEG |
| CH17-189H20.1 | 84.8448886531423 | 0.44713099619232 | 0.121264350181278 | 3.68724192661655 | 0,0002267 | 0.013361421132962 | Primary DEG |
| DENND4A | 2332.24692427507 | -0.204491429532895 | 0.0554666259250736 | -3.68674723083264 | 0,00022714 | 0.013361421132962 | Primary DEG |
| CDC23 | 1143.09726192626 | -0.105129826821193 | 0.0285216979513044 | -3.6859596157523 | 0,00022784 | 0.0133631637907525 | Primary DEG |
| TIMM21 | 263.573803916934 | -0.167000435184201 | 0.0453327917369351 | -3.68387714026746 | 0,00022971 | 0.0134331279544064 | Primary DEG |
| RP9P | 46.9864354406238 | 0.247790215193527 | 0.06728244775757 | 3.68283591712294 | 0,00023065 | 0.0134484634758885 | Primary DEG |
| ZNF614 | 380.870944459316 | -0.234604756183662 | 0.0637152322720619 | -3.68208272681024 | 0,00023134 | 0.0134487128015283 | Primary DEG |
| SEC23IP | 2174.02756827496 | -0.169078801004738 | 0.0459551818843051 | -3.67921078912067 | 0,00023396 | 0.0135506859900634 | Primary DEG |
| ZNF701 | 494.923470119974 | -0.188248054055642 | 0.0511729190782784 | -3.67866554119537 | 0,00023446 | 0.0135506859900634 | Primary DEG |
| DUSP28 | 244.566296940965 | 0.157875138314403 | 0.0429324821796699 | 3.67728885680789 | 0,00023573 | 0.0135708328618206 | Primary DEG |
| FAM189B | 190.40970234618 | 0.145818395728835 | 0.0396590169046187 | 3.67680308565221 | 0,00023618 | 0.0135708328618206 | Primary DEG |
| DBR1 | 725.269171856597 | -0.167408665330361 | 0.045556612322422 | -3.6747391168058 | 0,00023809 | 0.0136414656719936 | Primary DEG |
| TXNL4B | 505.720608256147 | -0.148514831417456 | 0.0404442464377598 | -3.67208798527147 | 0,00024058 | 0.0137348878361373 | Primary DEG |
| HUS1 | 688.57598841732 | -0.0833338209623478 | 0.0226973403871118 | -3.67152360325296 | 0,00024111 | 0.0137348878361373 | Primary DEG |
| ARMC1 | 1027.93099184317 | -0.123714896208834 | 0.0337151541794578 | -3.66941511079346 | 0,00024311 | 0.0137718432637308 | Primary DEG |
| MED6 | 504.992369135454 | -0.149915691931953 | 0.0408559524304573 | -3.66937209913613 | 0,00024315 | 0.0137718432637308 | Primary DEG |
| RP11-996F15.5 | 4.52905217984294 | -0.485459124537423 | 0.132395283232721 | -3.66674033004706 | 0,00024566 | 0.0138746556625314 | Primary DEG |
| REM2 | 43.3965228714515 | 0.418924983773629 | 0.114324249770706 | 3.66435803964466 | 0,00024796 | 0.0139646406936744 | Primary DEG |
| MOK | 23.6027201488053 | 0.414934576558999 | 0.113260211756726 | 3.66355112817772 | 0,00024874 | 0.0139680276794659 | Primary DEG |
| INTS2 | 596.858646405384 | -0.208850954333136 | 0.0570187736858719 | -3.66284542497772 | 0,00024943 | 0.0139680276794659 | Primary DEG |
| ZNF217 | 5651.24978577969 | -0.214609780626176 | 0.0586682146748432 | -3.65802473819269 | 0,00025417 | 0.0141684714677606 | Primary DEG |
| ZKSCAN4 | 320.200176095468 | -0.201610558576251 | 0.055141223564023 | -3.65625834077051 | 0,00025592 | 0.0141684714677606 | Primary DEG |
| PELO | 381.694189347245 | -0.236173645938088 | 0.0645983526094557 | -3.65603202555226 | 0,00025615 | 0.0141684714677606 | Primary DEG |
| DACT1 | 239.375679392027 | 0.535258258815013 | 0.146411459862607 | 3.65584947597204 | 0,00025633 | 0.0141684714677606 | Primary DEG |
| KPNA1 | 1569.38093507356 | -0.0874832960230907 | 0.0239335808077414 | -3.65525312429613 | 0,00025693 | 0.0141684714677606 | Primary DEG |
| HSPH1 | 2329.58435342296 | -0.205858123779206 | 0.056324095233614 | -3.65488558538533 | 0,0002573 | 0.0141684714677606 | Primary DEG |
| SLC16A1 | 574.459854462744 | -0.174476533764846 | 0.0477599118549306 | -3.65320049783201 | 0,00025899 | 0.0142039481286456 | Primary DEG |
| RP11-354E11.2 | 82.7626782328856 | 0.544819716848246 | 0.149150356771021 | 3.65282208264956 | 0,00025937 | 0.0142039481286456 | Primary DEG |
| RALA | 1549.52777453571 | -0.134591132048832 | 0.0368552110960054 | -3.65188878441724 | 0,00026032 | 0.0142164088827474 | Primary DEG |
| RP11-37B2.1 | 65.5537455763565 | 0.346155182855433 | 0.0948200185804313 | 3.65065508357611 | 0,00026157 | 0.0142456306051964 | Primary DEG |
| OXSM | 210.026384221336 | -0.170522460971003 | 0.0467318411357376 | -3.64895661773099 | 0,00026331 | 0.0143008467544028 | Primary DEG |
| RP11-172F4.7 | 19.1335165063622 | 0.398250144468385 | 0.109206601917901 | 3.64675887239655 | 0,00026557 | 0.0143842574255374 | Primary DEG |
| BUD13 | 755.651476021672 | -0.223685719031232 | 0.0613734586254644 | -3.64466536579419 | 0,00026774 | 0.0144623339661582 | Primary DEG |
| SEPTIN7P13 | 56.9588770251344 | 0.371073618217222 | 0.101849376984934 | 3.64335678039654 | 0,00026911 | 0.0144965917492436 | Primary DEG |
| STK38L | 1510.41181928286 | -0.207318432132163 | 0.0569330735034908 | -3.64144107061866 | 0,00027112 | 0.0145633866211713 | Primary DEG |
| RP11-981P6.1 | 20.4860192380565 | 0.509362770285529 | 0.139904894551166 | 3.64077877274869 | 0,00027182 | 0.0145633866211713 | Primary DEG |
| PAPOLG | 1266.56920081034 | -0.18174748314242 | 0.0499502654900831 | -3.63856891168083 | 0,00027416 | 0.0146309480296549 | Primary DEG |
| PSTK | 110.329877324149 | 0.188801164108666 | 0.0518941336522492 | 3.6381985943508 | 0,00027455 | 0.0146309480296549 | Primary DEG |
| TOP1 | 2516.65539443728 | -0.192525581862882 | 0.0529304152036586 | -3.63733367898414 | 0,00027548 | 0.014640792565069 | Primary DEG |
| PABIR3 | 193.195693762221 | 0.177541364724312 | 0.0488366149680826 | 3.63541504341251 | 0,00027753 | 0.0147107568362436 | Primary DEG |
| TOB2 | 2142.86452998444 | -0.190519607918887 | 0.0524353092475622 | -3.63342203284029 | 0,00027969 | 0.0147853729169988 | Primary DEG |
| CEP19 | 176.204987499491 | -0.245733623856693 | 0.0676833301563387 | -3.63063731186223 | 0,00028272 | 0.014823395103825 | Primary DEG |
| BTRC | 830.8252007549 | -0.134423720194497 | 0.0370254432759258 | -3.63057693037534 | 0,00028279 | 0.014823395103825 | Primary DEG |
| GALNT3 | 358.682942949464 | -0.27497025884068 | 0.0757815444663118 | -3.62845941946876 | 0,00028512 | 0.014823395103825 | Primary DEG |
| CLINT1 | 4219.65827314604 | -0.103061965235069 | 0.0284135569844639 | -3.62721095748137 | 0,0002865 | 0.014823395103825 | Primary DEG |
| HYLS1 | 197.451638728971 | -0.223030123236169 | 0.061492852131646 | -3.62692761036191 | 0,00028681 | 0.014823395103825 | Primary DEG |
| LINC01144 | 20.3825075774162 | 0.318563367583281 | 0.0878374554784957 | 3.6267372028015 | 0,00028703 | 0.014823395103825 | Primary DEG |
| CCDC85C | 222.565313176575 | 0.266138972284528 | 0.0734015555591397 | 3.62579471589127 | 0,00028807 | 0.014823395103825 | Primary DEG |
| STIM1 | 5019.55627453143 | 0.0891631162354551 | 0.0245928614110965 | 3.6255690114703 | 0,00028833 | 0.014823395103825 | Primary DEG |
| LYRM2 | 1056.93354739272 | -0.103745894530438 | 0.0286166965302472 | -3.62536236217135 | 0,00028856 | 0.014823395103825 | Primary DEG |
| PHC1 | 402.855159560424 | 0.221695477404726 | 0.061154919435454 | 3.62514544130362 | 0,0002888 | 0.014823395103825 | Primary DEG |
| OCRL | 346.458085316735 | -0.153509424816993 | 0.0423457650088382 | -3.62514232025217 | 0,0002888 | 0.014823395103825 | Primary DEG |
| ARCN1 | 6830.23908887556 | -0.133262868909821 | 0.0367722318281102 | -3.62400817912689 | 0,00029007 | 0.014823395103825 | Primary DEG |
| ITCH | 3987.99530158888 | -0.12919143034335 | 0.0356502141590715 | -3.62386126958163 | 0,00029024 | 0.014823395103825 | Primary DEG |
| UBE2G2 | 3285.61134732258 | 0.136385248174398 | 0.0376605400668193 | 3.62143633448741 | 0,00029297 | 0.014823395103825 | Primary DEG |
| TRNT1 | 385.487982902718 | -0.202108996402812 | 0.0558146078732481 | -3.62107706394338 | 0,00029338 | 0.014823395103825 | Primary DEG |
| ADD1 | 13000.961827077 | 0.0893563940794535 | 0.0246769340501745 | 3.62104927207607 | 0,00029341 | 0.014823395103825 | Primary DEG |
| LYSMD3 | 789.895022765764 | -0.232133147338951 | 0.0641157065267311 | -3.62053480986239 | 0,000294 | 0.014823395103825 | Primary DEG |
| BUB1B | 43.0438453799282 | -0.409424520075012 | 0.113099024853654 | -3.62005349387223 | 0,00029454 | 0.014823395103825 | Primary DEG |
| TNPO1 | 3094.23146037183 | -0.145822108080973 | 0.040282452590356 | -3.61999080750817 | 0,00029461 | 0.014823395103825 | Primary DEG |
| C5orf22 | 936.269773763167 | -0.096675788476544 | 0.026715674549696 | -3.61869165222497 | 0,0002961 | 0.0148292007320393 | Primary DEG |
| SEC24A | 1287.00269744885 | -0.128993249771756 | 0.0356597177800009 | -3.61733793204892 | 0,00029765 | 0.0148292007320393 | Primary DEG |
| ZSCAN32 | 582.616555747457 | -0.12163474292428 | 0.0336281148970639 | -3.61705505338629 | 0,00029797 | 0.0148292007320393 | Primary DEG |
| GAPVD1 | 2267.06371125223 | -0.166760827821534 | 0.0461050820929888 | -3.61697279890308 | 0,00029807 | 0.0148292007320393 | Primary DEG |
| FBXO28 | 1291.60540528776 | -0.120699705322334 | 0.0333735738264639 | -3.61662511632553 | 0,00029847 | 0.0148292007320393 | Primary DEG |
| ZNF260 | 468.632892150127 | -0.236402966988865 | 0.0654088681227561 | -3.61423418220898 | 0,00030124 | 0.0148947118034952 | Primary DEG |
| PARG | 827.291633718343 | -0.15189888727747 | 0.0420318585334721 | -3.61389889901026 | 0,00030163 | 0.0148947118034952 | Primary DEG |
| SIK1B | 82.2651254846425 | 0.676144583919088 | 0.187114046832137 | 3.61354262475905 | 0,00030204 | 0.0148947118034952 | Primary DEG |
| BLZF1 | 442.873012840526 | -0.217334154562929 | 0.0601660997568768 | -3.61223605055252 | 0,00030357 | 0.0149328357146448 | Primary DEG |
| MLLT6 | 2787.78125966839 | 0.234046291267945 | 0.0648127987328307 | 3.61111224702274 | 0,00030489 | 0.0149605771933728 | Primary DEG |
| ANKRD13C | 822.094663838834 | -0.115036189708424 | 0.031867913034725 | -3.60978108554127 | 0,00030646 | 0.0150004306618555 | Primary DEG |
| ZNF555 | 245.393103973863 | -0.152648915865698 | 0.0423017946436797 | -3.6085683161082 | 0,00030789 | 0.0150092601008696 | Primary DEG |
| ZNF200 | 426.805296656134 | -0.232101372707361 | 0.0643234024980656 | -3.60835036228597 | 0,00030815 | 0.0150092601008696 | Primary DEG |
| TGDS | 307.297594445178 | -0.163906027541065 | 0.0454708592468977 | -3.60463888863608 | 0,00031259 | 0.0150279870361581 | Primary DEG |
| ARMCX3 | 784.792992613539 | -0.189291340670402 | 0.0525231071197308 | -3.60396311358535 | 0,0003134 | 0.0150279870361581 | Primary DEG |
| RRN3 | 420.94503258637 | -0.186852197075133 | 0.0518467876482715 | -3.60393007070637 | 0,00031344 | 0.0150279870361581 | Primary DEG |
| IGHV1-46 | 33.7631461211094 | -0.677191310789806 | 0.187909048531383 | -3.60382491467252 | 0,00031357 | 0.0150279870361581 | Primary DEG |
| TMEM184C | 1159.10194234505 | -0.204163889682229 | 0.0566537557935302 | -3.60371323705857 | 0,0003137 | 0.0150279870361581 | Primary DEG |
| ATL2 | 937.16896786428 | -0.167749472041575 | 0.0465572576800834 | -3.60307888394673 | 0,00031447 | 0.0150279870361581 | Primary DEG |
| DHX36 | 1189.23424194814 | -0.186599291427122 | 0.0517971666893493 | -3.60249996966515 | 0,00031517 | 0.0150279870361581 | Primary DEG |
| GEMIN6 | 383.985495164434 | -0.229782618592194 | 0.0637864413857717 | -3.60237400927417 | 0,00031532 | 0.0150279870361581 | Primary DEG |
| TRMT1L | 798.250277083088 | -0.29784546771977 | 0.0826809536580707 | -3.60234678655882 | 0,00031536 | 0.0150279870361581 | Primary DEG |
| HAUS6 | 730.976156330318 | -0.12382616832311 | 0.0344106982851052 | -3.59847880148101 | 0,00032008 | 0.0152166521050548 | Primary DEG |
| NRIP2 | 106.26814640713 | 0.314270038452704 | 0.0873674634041512 | 3.59710613319434 | 0,00032178 | 0.0152221174735454 | Primary DEG |
| HACD3 | 964.569514076449 | -0.175301533253505 | 0.0487448588288742 | -3.59630815362346 | 0,00032277 | 0.0152221174735454 | Primary DEG |
| TTC37 | 2265.6435028644 | -0.142964474709786 | 0.0397552279606549 | -3.59611759367284 | 0,000323 | 0.0152221174735454 | Primary DEG |
| EIF2S1 | 2176.33313655959 | -0.136162622802957 | 0.0378710787940931 | -3.59542498230062 | 0,00032386 | 0.0152221174735454 | Primary DEG |
| CDC25A | 25.7306093432404 | -0.421209439164409 | 0.11715610501421 | -3.59528373799488 | 0,00032404 | 0.0152221174735454 | Primary DEG |
| ILF3-DT | 721.84339923361 | 0.218909096594942 | 0.060916748917395 | 3.59357812892789 | 0,00032617 | 0.0152859342760626 | Primary DEG |
| RP11-378J18.8 | 30.8665886005102 | 0.319379250863268 | 0.0889089788145316 | 3.59220469205376 | 0,00032789 | 0.0153305037677081 | Primary DEG |
| RNU6-415P | 27.3138830262413 | 0.609410143277557 | 0.16976205970039 | 3.58978999402514 | 0,00033094 | 0.0154368077216707 | Primary DEG |
| LINC00641 | 398.742968204969 | 0.415326717499469 | 0.115738153620916 | 3.58850305198242 | 0,00033258 | 0.0154502150104315 | Primary DEG |
| EML3 | 2235.15712921764 | 0.113519228362883 | 0.0316355867507604 | 3.58833958912283 | 0,00033279 | 0.0154502150104315 | Primary DEG |
| CCDC77 | 222.825919146576 | -0.212297532431191 | 0.0591892424834649 | -3.58675873391181 | 0,00033481 | 0.015507826109576 | Primary DEG |
| ZNF830 | 641.285422212342 | -0.198123633599212 | 0.0552581499242801 | -3.58541923446043 | 0,00033654 | 0.0155513108736511 | Primary DEG |
| FBXO30 | 620.720527691423 | -0.235827804008925 | 0.0657902524961558 | -3.58454018735838 | 0,00033767 | 0.0155674908049737 | Primary DEG |
| MECOM | 9.32712914026766 | 0.534267103864919 | 0.149212017682231 | 3.58059030474823 | 0,00034282 | 0.0156561559717196 | Primary DEG |
| TMEM9B-AS1 | 51.6105531166008 | 0.410388485959042 | 0.114622556870973 | 3.58034663649149 | 0,00034314 | 0.0156561559717196 | Primary DEG |
| FLJ40194 | 11.460251849521 | 0.4855639006588 | 0.135630395122237 | 3.58005224581988 | 0,00034353 | 0.0156561559717196 | Primary DEG |
| WRN | 455.072770381296 | -0.235629017643663 | 0.0658196697144242 | -3.57991795865888 | 0,0003437 | 0.0156561559717196 | Primary DEG |
| DHFR2 | 324.445298051284 | -0.2836425505147 | 0.0792348160334171 | -3.57977168010429 | 0,0003439 | 0.0156561559717196 | Primary DEG |
| CHUK | 1217.180280004 | -0.114251068971222 | 0.0319187199275878 | -3.57943768517086 | 0,00034433 | 0.0156561559717196 | Primary DEG |
| NUP58 | 1482.99503326697 | -0.185316470527989 | 0.0518175511978302 | -3.57632628798848 | 0,00034846 | 0.0157880075614943 | Primary DEG |
| TTI1 | 884.444630759972 | -0.102957494887858 | 0.0287908557777318 | -3.57604843991787 | 0,00034883 | 0.0157880075614943 | Primary DEG |
| RPP30 | 564.560947528497 | -0.103633373461433 | 0.0289862999146358 | -3.57525361176251 | 0,00034989 | 0.0157997919023277 | Primary DEG |
| ADORA1 | 18.8008087630187 | 0.577786310566777 | 0.161635722931679 | 3.57462014019636 | 0,00035074 | 0.0157997919023277 | Primary DEG |
| CCDC51 | 176.291462006291 | -0.164129200694999 | 0.0459286566812171 | -3.57356849851262 | 0,00035215 | 0.0157997919023277 | Primary DEG |
| RBM48 | 597.876885929406 | -0.203338644029127 | 0.0569129602405457 | -3.57280034582115 | 0,00035318 | 0.0157997919023277 | Primary DEG |
| ELP3 | 1798.95881438318 | -0.136156939893966 | 0.0381195030672126 | -3.57184456612388 | 0,00035448 | 0.0157997919023277 | Primary DEG |
| KIFAP3 | 700.4747792587 | -0.148224671886318 | 0.041500952042977 | -3.57159690536308 | 0,00035481 | 0.0157997919023277 | Primary DEG |
| SMIM15 | 870.681101439985 | -0.178422977486449 | 0.0499599393587718 | -3.57132093786503 | 0,00035519 | 0.0157997919023277 | Primary DEG |
| COPS7B | 1183.45193512694 | 0.0972395353817742 | 0.0272353247323446 | 3.5703460978489 | 0,00035651 | 0.0157997919023277 | Primary DEG |
| BCAR3 | 54.5405019257837 | -0.257228311924568 | 0.0720493999220297 | -3.57016591675899 | 0,00035676 | 0.0157997919023277 | Primary DEG |
| CASTOR3 | 169.957212596219 | 0.198807025277771 | 0.0557023524718505 | 3.56909567469775 | 0,00035822 | 0.0157997919023277 | Primary DEG |
| CASP3 | 1196.97494668278 | -0.199198509577528 | 0.0558136303268965 | -3.56899396098832 | 0,00035836 | 0.0157997919023277 | Primary DEG |
| PCMTD2 | 998.310505346463 | 0.150773537663471 | 0.0422479564346296 | 3.56877705781493 | 0,00035865 | 0.0157997919023277 | Primary DEG |
| RP11-35O15.2 | 3.159392893214 | 0.981649265774954 | 0.275280215192265 | 3.56600006683858 | 0,00036247 | 0.01591856568712 | Primary DEG |
| PREB | 1507.26322032521 | -0.118517007550306 | 0.0332385291521245 | -3.56565138631385 | 0,00036295 | 0.01591856568712 | Primary DEG |
| CCDC127 | 482.935700537599 | -0.186288468743332 | 0.0522716238209318 | -3.56385463328066 | 0,00036545 | 0.0159548705485208 | Primary DEG |
| RP11-81A1.6 | 163.981184011763 | 0.260386320760139 | 0.0730670236350185 | 3.56366398692809 | 0,00036571 | 0.0159548705485208 | Primary DEG |
| PP2D1 | 39.4031623488096 | 0.388279881130008 | 0.108970733981798 | 3.56315743633391 | 0,00036642 | 0.0159548705485208 | Primary DEG |
| STIP1 | 3187.14716488777 | -0.133062095317892 | 0.037348221503688 | -3.56274248038164 | 0,000367 | 0.0159548705485208 | Primary DEG |
| RDX | 850.513129989289 | -0.162596457108449 | 0.0456490339008321 | -3.5618816700847 | 0,00036821 | 0.0159722593183748 | Primary DEG |
| ZNF468 | 744.086828137325 | -0.174231297707137 | 0.0489278966646833 | -3.56098074072534 | 0,00036947 | 0.0159921760200033 | Primary DEG |
| BIK | 40.5338331558017 | -0.292597833492222 | 0.0821969992762859 | -3.5597142969748 | 0,00037126 | 0.0160344836803921 | Primary DEG |
| PPM1N | 99.5412571056953 | 0.298552205882407 | 0.0839133369209037 | 3.55786358685535 | 0,00037388 | 0.0161127482056338 | Primary DEG |
| DCAF11 | 3225.03722052379 | 0.0744104302659084 | 0.020918030997897 | 3.55723874170515 | 0,00037477 | 0.0161160745811686 | Primary DEG |
| DOLPP1 | 353.047762707616 | -0.12946707415821 | 0.0364115391640644 | -3.55566057163507 | 0,00037703 | 0.0161368775053122 | Primary DEG |
| MMUT | 683.652721452162 | -0.185561726815224 | 0.0521988901031606 | -3.55489793841399 | 0,00037813 | 0.0161368775053122 | Primary DEG |
| PEX13 | 584.419096182599 | -0.116651074141702 | 0.0328170890380426 | -3.55458322359051 | 0,00037858 | 0.0161368775053122 | Primary DEG |
| WDR5B | 263.285486879798 | -0.267890237406566 | 0.0753934505311479 | -3.55322956462764 | 0,00038053 | 0.0161368775053122 | Primary DEG |
| NAPEPLD | 25.2937311063367 | -0.359634347145496 | 0.101213425892946 | -3.55322768666959 | 0,00038054 | 0.0161368775053122 | Primary DEG |
| COA6 | 314.908459466856 | -0.207268442580005 | 0.058336943417669 | -3.55295341917464 | 0,00038093 | 0.0161368775053122 | Primary DEG |
| UBTD2 | 443.198953676253 | -0.169892045166812 | 0.0478178680269342 | -3.55289878400931 | 0,00038101 | 0.0161368775053122 | Primary DEG |
| IDE | 1338.7853137125 | -0.114266916666438 | 0.0321701134084618 | -3.55195877663218 | 0,00038238 | 0.0161368775053122 | Primary DEG |
| INO80E | 986.132479630887 | 0.1710861561842 | 0.0481820059243981 | 3.55083091502354 | 0,00038402 | 0.0161368775053122 | Primary DEG |
| NUP153 | 2655.89126056902 | -0.173040120612796 | 0.0487339660248438 | -3.55070877105677 | 0,0003842 | 0.0161368775053122 | Primary DEG |
| NEMP1 | 633.353037620187 | -0.195868979630549 | 0.0551635180241642 | -3.55069775543955 | 0,00038421 | 0.0161368775053122 | Primary DEG |
| MAP4K2 | 2359.20393296866 | 0.126446437886471 | 0.0356216203500286 | 3.54971044674474 | 0,00038566 | 0.0161632691777077 | Primary DEG |
| EDEM3 | 2042.3915800818 | -0.163093807065786 | 0.0459603474512389 | -3.54857646015009 | 0,00038732 | 0.0161987813577049 | Primary DEG |
| VSTM4 | 4.69026634161375 | 1.01236707333647 | 0.285429093518768 | 3.54682510060911 | 0,0003899 | 0.0162725135392418 | Primary DEG |
| ZC4H2 | 258.188891644123 | 0.168704849300771 | 0.04758749650421 | 3.5451507579485 | 0,00039239 | 0.0163418306774765 | Primary DEG |
| CHTOP | 2625.01843582975 | -0.0908549843489282 | 0.0256359637724194 | -3.54404402953148 | 0,00039404 | 0.0163761797901594 | Primary DEG |
| CTC-459F4.3 | 103.093459518078 | -0.224613642957986 | 0.0634225112878381 | -3.54154445160007 | 0,00039779 | 0.0164975421849377 | Primary DEG |
| RP5-1174N9.3 | 9.23846700223942 | 0.4304960137343 | 0.121620941095253 | 3.5396536966207 | 0,00040065 | 0.0165814887405369 | Primary DEG |
| GMNN | 129.993594702931 | -0.18809066793727 | 0.0531529902953357 | -3.53866578140148 | 0,00040216 | 0.016590355082169 | Primary DEG |
| SPATA5L1 | 228.728853655993 | -0.208129319548014 | 0.0588199676964177 | -3.53841268023494 | 0,00040254 | 0.016590355082169 | Primary DEG |
| TRMT5 | 284.296789844475 | -0.226619959236998 | 0.0640897084410409 | -3.53598049904496 | 0,00040627 | 0.016694096269675 | Primary DEG |
| RAB11FIP4 | 2355.74505541007 | 0.119480275073189 | 0.0337928199945225 | 3.53567044989307 | 0,00040674 | 0.016694096269675 | Primary DEG |
| TRIM47 | 94.1326722897533 | -0.252839836261401 | 0.0715470988748703 | -3.53389362025141 | 0,00040949 | 0.0167263909792021 | Primary DEG |
| WDTC1 | 4560.85821266256 | 0.107316033814072 | 0.0303739175305638 | 3.53316406110886 | 0,00041062 | 0.0167263909792021 | Primary DEG |
| CD99P1 | 164.321392130429 | 0.213878867705726 | 0.060542192053594 | 3.5327242118421 | 0,0004113 | 0.0167263909792021 | Primary DEG |
| SEPTIN6 | 10264.1897660604 | 0.134294545902024 | 0.0380152162249645 | 3.53265242810412 | 0,00041141 | 0.0167263909792021 | Primary DEG |
| ZSWIM3 | 165.971266606872 | -0.333553960391301 | 0.0944260009419587 | -3.53243764496951 | 0,00041175 | 0.0167263909792021 | Primary DEG |
| ARMT1 | 857.194133492817 | -0.190625647437856 | 0.0539952370106151 | -3.53041597725332 | 0,00041491 | 0.0168202697840747 | Primary DEG |
| SMAD5 | 751.806355639371 | -0.193509337314534 | 0.0548333869660646 | -3.5290422135385 | 0,00041707 | 0.0168733228849289 | Primary DEG |
| AK6 | 62.6182968979483 | -0.178953778335267 | 0.0507174904825816 | -3.52844308013872 | 0,00041801 | 0.0168771213818558 | Primary DEG |
| HBS1L | 1247.71501701897 | -0.0854449982802553 | 0.0242333317545585 | -3.5259286319217 | 0,000422 | 0.0170035393276071 | Primary DEG |
| G3BP1 | 5008.11029560708 | -0.147321734428814 | 0.0418079341961195 | -3.52377454809733 | 0,00042545 | 0.017092897679365 | Primary DEG |
| RP1-267D11.6 | 73.0963731377929 | 0.239321489896574 | 0.0679353981963238 | 3.52278040977943 | 0,00042705 | 0.017092897679365 | Primary DEG |
| CRPPA | 20.8762449695653 | -0.264988107287924 | 0.075243385197815 | -3.52174621850505 | 0,00042871 | 0.017092897679365 | Primary DEG |
| NFKBIA | 6025.21142462998 | 0.361360979385528 | 0.102612557346533 | 3.52160582223067 | 0,00042894 | 0.017092897679365 | Primary DEG |
| PHF7 | 129.077476555068 | 0.171038167931475 | 0.04858104006144 | 3.52067736127437 | 0,00043045 | 0.017092897679365 | Primary DEG |
| SIT1 | 1128.11131412062 | -0.268460749726996 | 0.0762583292547913 | -3.52041216153616 | 0,00043088 | 0.017092897679365 | Primary DEG |
| AC100830.3 | 4.1077362576408 | 0.719161705849484 | 0.204290388903741 | 3.52029143274255 | 0,00043107 | 0.017092897679365 | Primary DEG |
| IGHG1 | 2557.71341349043 | -0.672760184551361 | 0.191110679255849 | -3.52026473439878 | 0,00043112 | 0.017092897679365 | Primary DEG |
| SUDS3 | 1710.78020335637 | -0.111828042865525 | 0.0317777279084703 | -3.51906980850312 | 0,00043306 | 0.017135799911085 | Primary DEG |
| COPB1 | 4472.59753776046 | -0.127729310709731 | 0.0363161595225154 | -3.51714807923292 | 0,00043621 | 0.017225964395875 | Primary DEG |
| MTO1 | 1058.3155567752 | -0.142457779399237 | 0.0405101302087808 | -3.51659643316472 | 0,00043712 | 0.0172274832637481 | Primary DEG |
| RP11-840I19.3 | 13.3539438161143 | 0.542337812314401 | 0.154303455258086 | 3.51474833410111 | 0,00044017 | 0.0172924636885813 | Primary DEG |
| AF131216.6 | 16.5794613149615 | 0.448051031123511 | 0.12748485942073 | 3.51454308503286 | 0,00044051 | 0.0172924636885813 | Primary DEG |
| ZNF580 | 377.966649847315 | 0.184781983665718 | 0.0525846932217499 | 3.51398805136109 | 0,00044143 | 0.0172943897761858 | Primary DEG |
| UICLM | 104.572487038337 | 0.631328143411778 | 0.179746096447686 | 3.51233298463048 | 0,00044419 | 0.0173681344171036 | Primary DEG |
| NUP133 | 1764.94761333513 | -0.115742948437045 | 0.0329665012832095 | -3.51092605923564 | 0,00044655 | 0.0174015994824742 | Primary DEG |
| CD69 | 2244.80592129879 | 0.427683388436129 | 0.121822001646807 | 3.51072370060125 | 0,00044689 | 0.0174015994824742 | Primary DEG |
| PORCN | 391.346151064502 | 0.135025528739559 | 0.038472857994238 | 3.50963083532244 | 0,00044873 | 0.0174015994824742 | Primary DEG |
| NCK2 | 2640.00150030003 | 0.190187369510292 | 0.0541950348607318 | 3.50931353765208 | 0,00044927 | 0.0174015994824742 | Primary DEG |
| EXOSC3 | 325.141785019578 | -0.204314168247456 | 0.0582222305569621 | -3.50921231105297 | 0,00044944 | 0.0174015994824742 | Primary DEG |
| RASAL3 | 4222.9399169362 | 0.140977392889289 | 0.0401876563066012 | 3.50797746984146 | 0,00045153 | 0.0174174470450648 | Primary DEG |
| RP11-143J12.3 | 11.5554775146008 | 0.715404437185619 | 0.203939015834874 | 3.50793316451459 | 0,0004516 | 0.0174174470450648 | Primary DEG |
| GIN1 | 241.234804429611 | -0.195356587854499 | 0.0557054485995454 | -3.50695655031658 | 0,00045326 | 0.0174448399040074 | Primary DEG |
| HNRNPDL | 10029.7511960635 | 0.212009205598035 | 0.060476856780139 | 3.50562540591 | 0,00045554 | 0.0174448399040074 | Primary DEG |
| PAQR8 | 1631.99500128113 | -0.247617569808089 | 0.0706357666730708 | -3.5055550675079 | 0,00045566 | 0.0174448399040074 | Primary DEG |
| CDKN2D | 2627.9794010914 | 0.235133404748933 | 0.0670853752763435 | 3.50498754430982 | 0,00045663 | 0.0174448399040074 | Primary DEG |
| CXCR3 | 676.217186154576 | -0.259734772264375 | 0.0741053709682739 | -3.50493856073635 | 0,00045671 | 0.0174448399040074 | Primary DEG |
| ASB12 | 4.46764483079401 | 0.576665828034697 | 0.164634512239033 | 3.50270317075094 | 0,00046056 | 0.0175580554870553 | Primary DEG |
| CLDND1 | 1280.08361898728 | -0.170351210039038 | 0.0486542590760487 | -3.5012599775237 | 0,00046306 | 0.0176195361691462 | Primary DEG |
| SHCBP1 | 124.236373221012 | -0.349076200529117 | 0.0997449996818429 | -3.49968621627718 | 0,00046581 | 0.0176899246071896 | Primary DEG |
| PABIR1 | 879.877786818945 | -0.168887085195904 | 0.0482675906112074 | -3.49897484124037 | 0,00046705 | 0.0177032735191401 | Primary DEG |
| EXOC7 | 2434.00250345626 | 0.0906920653149192 | 0.0259390801975311 | 3.49634854529465 | 0,00047167 | 0.0178278504287703 | Primary DEG |
| SMARCA5 | 4694.09527727776 | -0.145080266588884 | 0.0414979069879706 | -3.49608635999255 | 0,00047214 | 0.0178278504287703 | Primary DEG |
| ZFP14 | 308.618680576993 | -0.237294906384787 | 0.067935542083187 | -3.49294197276324 | 0,00047773 | 0.0180048015596324 | Primary DEG |
| TMED5 | 3081.87438569964 | -0.148216914122863 | 0.0424524330135429 | -3.49136441898582 | 0,00048056 | 0.0180770917672556 | Primary DEG |
| ZNF436 | 370.355003302972 | -0.212508420066876 | 0.0608956004171003 | -3.48971713245808 | 0,00048353 | 0.018086736889959 | Primary DEG |
| ZDHHC18 | 2473.36681531173 | 0.107163635524676 | 0.0307131674343518 | 3.48917563627177 | 0,00048451 | 0.018086736889959 | Primary DEG |
| TGS1 | 1134.70566406815 | -0.118044397249655 | 0.0338322800626009 | -3.48910558292949 | 0,00048464 | 0.018086736889959 | Primary DEG |
| SLC33A1 | 168.530771460766 | -0.178370682789652 | 0.0511283851158707 | -3.48868211631203 | 0,00048541 | 0.018086736889959 | Primary DEG |
| MAPK8 | 804.326788151372 | -0.0947315687553217 | 0.0271621529286525 | -3.48763108006038 | 0,00048732 | 0.018086736889959 | Primary DEG |
| IGLL1 | 7.15589774260214 | -0.659909936678525 | 0.189238100375651 | -3.48719383342232 | 0,00048812 | 0.018086736889959 | Primary DEG |
| RP11-104L21.4 | 5.97913914890889 | 0.591874785241044 | 0.169733923676368 | 3.48707419484141 | 0,00048834 | 0.018086736889959 | Primary DEG |
| STRN3 | 398.084302871377 | -0.158627987823658 | 0.0455020493158036 | -3.48617238583502 | 0,00048999 | 0.018086736889959 | Primary DEG |
| DDX5 | 35585.0367559726 | 0.202593395294952 | 0.0581201180002487 | 3.48577054324089 | 0,00049072 | 0.018086736889959 | Primary DEG |
| TCEANC | 320.600143284786 | -0.135400837398209 | 0.0388461164191819 | -3.4855694694708 | 0,00049109 | 0.018086736889959 | Primary DEG |
| RP11-305O6.4 | 19.459589479245 | 0.423814763584787 | 0.121612609499227 | 3.48495740145663 | 0,00049222 | 0.018086736889959 | Primary DEG |
| RP11-157L14.5 | 69.8272658531029 | 0.308742660780763 | 0.088593556368066 | 3.48493359379409 | 0,00049226 | 0.018086736889959 | Primary DEG |
| MAP3K7 | 1995.33258035437 | -0.0630228878689105 | 0.0180855664738493 | -3.48470632424137 | 0,00049268 | 0.018086736889959 | Primary DEG |
| KCTD6 | 305.276149362032 | -0.157840228126799 | 0.0453315242338379 | -3.48190868925115 | 0,00049785 | 0.0182359026501303 | Primary DEG |
| PHAX | 1054.65934514319 | -0.142829681904624 | 0.0410251970300453 | -3.48151117470614 | 0,00049859 | 0.0182359026501303 | Primary DEG |
| TWNK | 399.534712952211 | -0.192031548396287 | 0.0551722411394941 | -3.48058270663259 | 0,00050032 | 0.0182359026501303 | Primary DEG |
| P2RY8 | 11053.9881319852 | 0.159353129111731 | 0.0457841383794155 | 3.48053135326395 | 0,00050042 | 0.0182359026501303 | Primary DEG |
| ATG4C | 634.314635307483 | -0.151493134934746 | 0.0435941860697108 | -3.47507657769995 | 0,00051071 | 0.0185766352962341 | Primary DEG |
| C21orf91 | 1413.34387982042 | -0.173449061535794 | 0.0499262829330271 | -3.47410324474716 | 0,00051256 | 0.0186100075975483 | Primary DEG |
| USF2 | 4694.7052739131 | 0.117507022456444 | 0.0338401392919167 | 3.47241544849411 | 0,0005158 | 0.018693178802132 | Primary DEG |
| COG6 | 803.924320485236 | -0.149249559559542 | 0.0429991054822471 | -3.47099219589956 | 0,00051854 | 0.0187582454652613 | Primary DEG |
| BHLHA15 | 75.7547797203865 | -0.581061738386726 | 0.167437413631749 | -3.47032198947289 | 0,00051984 | 0.0187668552197604 | Primary DEG |
| SGO2 | 159.742315164488 | -0.179947767069735 | 0.0518597813341686 | -3.46989058650686 | 0,00052067 | 0.0187668552197604 | Primary DEG |
| ACSL3 | 1307.0426509651 | -0.166085129242077 | 0.047878236505736 | -3.46890657140598 | 0,00052258 | 0.0188015403765327 | Primary DEG |
| WDR36 | 1096.7063667432 | -0.188133746049353 | 0.0542439814097599 | -3.46828793093537 | 0,00052379 | 0.0188107405578831 | Primary DEG |
| CUL3 | 2553.8877966318 | -0.164890024141688 | 0.0475530095338058 | -3.46749923418552 | 0,00052533 | 0.0188319123629145 | Primary DEG |
| WHRN | 87.3239599973835 | 0.434955647963942 | 0.125489020968491 | 3.46608527668053 | 0,0005281 | 0.0188521061893231 | Primary DEG |
| RP11-165F24.3 | 92.201289956185 | 0.480862197014536 | 0.138746409691451 | 3.46576317242295 | 0,00052873 | 0.0188521061893231 | Primary DEG |
| CACNA1C-AS2 | 28.8502822104624 | 0.315081004805777 | 0.0909165740159249 | 3.46560578438193 | 0,00052904 | 0.0188521061893231 | Primary DEG |
| RP1-68D18.2 | 10.5331439871321 | 0.468595370317007 | 0.135226084469877 | 3.46527352436498 | 0,00052969 | 0.0188521061893231 | Primary DEG |
| ZNF543 | 166.19822592628 | -0.332121523371077 | 0.0958653663025746 | -3.46445787650589 | 0,0005313 | 0.0188754798979574 | Primary DEG |
| KBTBD7 | 334.755434432375 | -0.309105269139 | 0.0892357130577558 | -3.46391885655622 | 0,00053237 | 0.0188795077498873 | Primary DEG |
| RP11-930P14.2 | 31.6304737119824 | 0.219572791881265 | 0.0634519127670128 | 3.46045977664232 | 0,00053925 | 0.0190895841544568 | Primary DEG |
| GIMAP7 | 6675.94608014788 | -0.28407666755442 | 0.0821120510641772 | -3.45962211237899 | 0,00054093 | 0.0191149277602826 | Primary DEG |
| ZFAND2A | 504.353086749007 | 0.210840045268896 | 0.0609722391380774 | 3.45796789242771 | 0,00054427 | 0.0191857347176337 | Primary DEG |
| GRM2 | 22.7726618201144 | 0.587068603196721 | 0.169796654817123 | 3.45748038339752 | 0,00054525 | 0.0191857347176337 | Primary DEG |
| NRBF2 | 1120.15056912018 | -0.211236484808785 | 0.0611006404265565 | -3.45718937369721 | 0,00054584 | 0.0191857347176337 | Primary DEG |
| C17orf80 | 552.325894717448 | -0.143529136616391 | 0.0415224755310274 | -3.45666135703156 | 0,00054691 | 0.0191893396688793 | Primary DEG |
| AP3M1 | 2404.11041954819 | -0.140526024912041 | 0.0406644616253777 | -3.45574536819496 | 0,00054877 | 0.0192206452024549 | Primary DEG |
| CSTF2 | 420.529498394976 | -0.198172021171792 | 0.0573606498667445 | -3.45484267755279 | 0,00055061 | 0.0192367192889345 | Primary DEG |
| CLASP2 | 1534.32747578876 | -0.219053808619661 | 0.063412514310531 | -3.4544255341612 | 0,00055147 | 0.0192367192889345 | Primary DEG |
| PLEKHA8 | 451.019240196245 | -0.157069424079934 | 0.0454734967800692 | -3.45408722007006 | 0,00055216 | 0.0192367192889345 | Primary DEG |
| JOSD1 | 3373.89772265695 | 0.14964950762107 | 0.0433311887021895 | 3.45362110071788 | 0,00055311 | 0.0192367192889345 | Primary DEG |
| PITPNB | 2151.92980075056 | -0.0805228644632692 | 0.0233243679500667 | -3.4523063877081 | 0,00055582 | 0.0192776241790443 | Primary DEG |
| TATDN2P2 | 117.006764187335 | 0.291228996505008 | 0.0843755558009993 | 3.45158018504642 | 0,00055731 | 0.0192776241790443 | Primary DEG |
| TFAM | 1700.99813856152 | -0.113717330442179 | 0.0329488908204595 | -3.45132499487864 | 0,00055784 | 0.0192776241790443 | Primary DEG |
| VPS33A | 940.280189824837 | -0.118190721915877 | 0.0342472740516356 | -3.45109866956645 | 0,00055831 | 0.0192776241790443 | Primary DEG |
| TMEM186 | 176.768272264563 | -0.157666816630422 | 0.0456913678614961 | -3.45069154218267 | 0,00055915 | 0.0192776241790443 | Primary DEG |
| G2E3 | 470.242148195186 | -0.212468436692648 | 0.0615921507311288 | -3.44960249269662 | 0,00056141 | 0.0193134938129115 | Primary DEG |
| RP11-196G11.6 | 13.7652620380223 | 0.38684063823106 | 0.112152026316954 | 3.44925233127582 | 0,00056214 | 0.0193134938129115 | Primary DEG |
| VAMP2 | 4602.19204122857 | 0.195497014905173 | 0.056700625927276 | 3.44788107199939 | 0,000565 | 0.0193514195980656 | Primary DEG |
| DNAJB4 | 140.973197034487 | -0.251957349837981 | 0.0730826588479493 | -3.44756682104555 | 0,00056566 | 0.0193514195980656 | Primary DEG |
| TBC1D25 | 1006.69519663851 | 0.100037998811205 | 0.0290190474854521 | 3.44732193092681 | 0,00056617 | 0.0193514195980656 | Primary DEG |
| C2CD5 | 1977.12875751191 | -0.156584636491122 | 0.0454492654747823 | -3.44526220292832 | 0,00057051 | 0.0194659292271417 | Primary DEG |
| UBQLN2 | 2326.98892213795 | -0.157485564265955 | 0.0457325289567728 | -3.44362246869867 | 0,00057398 | 0.0195507071856479 | Primary DEG |
| SNHG30 | 257.464231965429 | 0.23549839065378 | 0.0684012429960891 | 3.44289636179922 | 0,00057552 | 0.0195696482063813 | Primary DEG |
| RP11-582J16.4 | 253.665247089583 | 0.263480202158817 | 0.0765531302293202 | 3.44179527825373 | 0,00057787 | 0.0195739095446396 | Primary DEG |
| RP11-434P11.2 | 7.2641017698329 | 0.511764437620407 | 0.148695023687982 | 3.44170520927642 | 0,00057806 | 0.0195739095446396 | Primary DEG |
| PSAT1 | 199.437228647952 | -0.274328595305441 | 0.0797235679079545 | -3.44099746792779 | 0,00057957 | 0.0195739095446396 | Primary DEG |
| CNNM3-DT | 22.8464330182928 | 0.417905619349209 | 0.121449317570952 | 3.44098779398298 | 0,0005796 | 0.0195739095446396 | Primary DEG |
| IGLV2-23 | 719.734519890209 | -0.503922400665942 | 0.146503861375151 | -3.43965268857695 | 0,00058246 | 0.01960937735248 | Primary DEG |
| COG4 | 1781.1279605094 | 0.14238890612767 | 0.0414035759123966 | 3.4390485118711 | 0,00058376 | 0.01960937735248 | Primary DEG |
| SLC39A9 | 2219.68989545233 | -0.124418794029614 | 0.0361785187203352 | -3.43902399629426 | 0,00058382 | 0.01960937735248 | Primary DEG |
| PUS3 | 395.667758296681 | -0.149175389988934 | 0.043385554467778 | -3.43836541491535 | 0,00058524 | 0.01960937735248 | Primary DEG |
| RP11-522I20.3 | 28.7187260571398 | -0.27885158047977 | 0.0811164072793656 | -3.43767173414625 | 0,00058674 | 0.01960937735248 | Primary DEG |
| ERCC4 | 443.031021658373 | -0.152351208070319 | 0.0443307800384582 | -3.43669134488837 | 0,00058887 | 0.01960937735248 | Primary DEG |
| TAMALIN | 197.366542366859 | 0.334547988207155 | 0.0973497789411315 | 3.43655621867883 | 0,00058916 | 0.01960937735248 | Primary DEG |
| FBXO38 | 1422.70451538669 | -0.21557795727134 | 0.062731203593584 | -3.43653468962595 | 0,00058921 | 0.01960937735248 | Primary DEG |
| SGCB | 223.141952401603 | -0.163766350456196 | 0.047656673096933 | -3.43637815680288 | 0,00058955 | 0.01960937735248 | Primary DEG |
| ID2 | 4782.05182611165 | 0.233887343735687 | 0.0680782346482025 | 3.43556710811188 | 0,00059132 | 0.0196352234773619 | Primary DEG |
| CNOT11 | 1984.35072055764 | -0.0930114735597813 | 0.027086584727678 | -3.43385755328315 | 0,00059506 | 0.0196894422178741 | Primary DEG |
| ISG20L2 | 1852.19187805816 | -0.09404970355652 | 0.0273960330619066 | -3.43296795357184 | 0,00059701 | 0.0196894422178741 | Primary DEG |
| AICDA | 7.39077682603603 | -0.486177510148864 | 0.141628492711679 | -3.43276625232892 | 0,00059746 | 0.0196894422178741 | Primary DEG |
| RP11-20I23.2 | 38.9832551665472 | 0.483180935452093 | 0.140759938287651 | 3.43265947207709 | 0,00059769 | 0.0196894422178741 | Primary DEG |
| ZBED6CL | 781.511112424504 | 0.211653369859471 | 0.0616605197723018 | 3.43255896384037 | 0,00059791 | 0.0196894422178741 | Primary DEG |
| PRKAA1 | 2914.4669359326 | -0.107022757167443 | 0.0311941585035718 | -3.43085892684647 | 0,00060167 | 0.0197137541738869 | Primary DEG |
| NCBP1 | 1536.50696347379 | -0.171304752245323 | 0.0499334722237351 | -3.43065972816319 | 0,00060212 | 0.0197137541738869 | Primary DEG |
| ZC3H10 | 218.91108190373 | -0.232810088248179 | 0.067871472609158 | -3.43016114574131 | 0,00060322 | 0.0197137541738869 | Primary DEG |
| RP11-331F9.3 | 2.44309954236282 | 0.745909059044942 | 0.217461760151288 | 3.43006999725383 | 0,00060343 | 0.0197137541738869 | Primary DEG |
| ILK | 13.1326224295535 | 0.417877477559594 | 0.121830856309928 | 3.42998063230015 | 0,00060362 | 0.0197137541738869 | Primary DEG |
| GOLPH3L | 605.157584090438 | -0.196804595299685 | 0.0573976521104072 | -3.42879173735403 | 0,00060628 | 0.0197677414742593 | Primary DEG |
| USP50 | 16.7493348889789 | 0.597326698984025 | 0.1742585564116 | 3.42781847436595 | 0,00060845 | 0.0198061669120561 | Primary DEG |
| UBXN4 | 4198.67745864564 | -0.111336855287323 | 0.0324940417837421 | -3.42637755033075 | 0,00061169 | 0.0198789057191995 | Primary DEG |
| SNAPC1 | 143.22173318382 | -0.178697317363466 | 0.052187354553332 | -3.42414975606492 | 0,00061673 | 0.0200097891568345 | Primary DEG |
| TTL | 1494.03427504591 | 0.120612028776263 | 0.0352323898872939 | 3.42332805586258 | 0,00061859 | 0.0200375851633198 | Primary DEG |
| UTP23 | 771.13474388416 | -0.153254549423906 | 0.0447869400233182 | -3.4218580091454 | 0,00062195 | 0.0200767148408944 | Primary DEG |
| RAB39B | 396.800780958055 | -0.190540573590231 | 0.0556871365436844 | -3.42162634706059 | 0,00062248 | 0.0200767148408944 | Primary DEG |
| PYGO2 | 2020.70917877458 | -0.132078148514924 | 0.038607457999451 | -3.42105270222148 | 0,00062379 | 0.0200767148408944 | Primary DEG |
| EIF4H | 8186.33502438617 | 0.101252089873292 | 0.0296005441441462 | 3.42061582990582 | 0,0006248 | 0.0200767148408944 | Primary DEG |
| ALOX12-AS1 | 105.209473747304 | 0.174009717097736 | 0.0508713287841162 | 3.42058525414552 | 0,00062487 | 0.0200767148408944 | Primary DEG |
| DDX11L17 | 22.6133300431772 | 0.677345984598626 | 0.198087213245342 | 3.41943315523196 | 0,00062752 | 0.0201006288600652 | Primary DEG |
| MTRES1 | 250.272735131581 | -0.115962934246978 | 0.033913428378536 | -3.41938104731315 | 0,00062764 | 0.0201006288600652 | Primary DEG |
| SLC30A6 | 1039.43952108023 | -0.128977336589044 | 0.0377387007804666 | -3.41764114613616 | 0,00063166 | 0.0201701354400611 | Primary DEG |
| TMEM170A | 1050.49166937429 | -0.147313709288035 | 0.0431071338977902 | -3.41738584702304 | 0,00063226 | 0.0201701354400611 | Primary DEG |
| RSBN1L | 1515.66986133503 | -0.155703380896382 | 0.0455655986678301 | -3.41712575821615 | 0,00063286 | 0.0201701354400611 | Primary DEG |
| BET1 | 392.956002001264 | -0.199592318002837 | 0.058437180949055 | -3.41550216422725 | 0,00063665 | 0.020226633433558 | Primary DEG |
| NARF-AS2 | 97.5902558022727 | 0.25136347663049 | 0.0735951436730605 | 3.4154899913933 | 0,00063667 | 0.020226633433558 | Primary DEG |
| PSMD11 | 1862.16402309436 | -0.0827811400863704 | 0.0242448203427452 | -3.41438455373587 | 0,00063926 | 0.0202763926981725 | Primary DEG |
| TBX19 | 140.758196640928 | 0.167803283988845 | 0.0491575173691604 | 3.41358337380396 | 0,00064115 | 0.0203036198131896 | Primary DEG |
| RAB22A | 1591.81333736274 | -0.0928848929309553 | 0.0272140557371491 | -3.413122021506 | 0,00064223 | 0.0203055870406311 | Primary DEG |
| RELA-DT | 83.8473186369197 | 0.189259454155552 | 0.0554782919559365 | 3.41141458186691 | 0,00064627 | 0.0203927346505986 | Primary DEG |
| NUP98 | 4920.81535276793 | -0.121702065523466 | 0.0356783869181927 | -3.41108654386527 | 0,00064705 | 0.0203927346505986 | Primary DEG |
| RP11-81A22.4 | 61.5583952169578 | 0.263950689189739 | 0.077404076668147 | 3.41003601556246 | 0,00064954 | 0.0204389534855778 | Primary DEG |
| METTL15 | 212.826980320955 | -0.148887068837289 | 0.0436673266918978 | -3.40957599460547 | 0,00065064 | 0.0204410095691689 | Primary DEG |
| DDIT4 | 1981.34768480631 | 0.426631211559322 | 0.125154913454629 | 3.4088251094831 | 0,00065243 | 0.0204551912811493 | Primary DEG |
| MRPL35 | 741.672002729203 | -0.154759054681029 | 0.0454137488078412 | -3.40775775494464 | 0,00065499 | 0.0204551912811493 | Primary DEG |
| CIRBP-AS1 | 29.0550833191657 | 0.450536716492848 | 0.132213533912313 | 3.40764446090408 | 0,00065526 | 0.0204551912811493 | Primary DEG |
| ALDH9A1 | 2479.27549024839 | -0.130064185296222 | 0.0381722351636724 | -3.40729812489475 | 0,00065609 | 0.0204551912811493 | Primary DEG |
| IL2RB | 11101.7815276954 | 0.209426983693971 | 0.0614654076929509 | 3.4072332968189 | 0,00065625 | 0.0204551912811493 | Primary DEG |
| ZNF227 | 462.360369865186 | -0.11818804984654 | 0.0346999382033976 | -3.40600173849785 | 0,00065922 | 0.0205154310610835 | Primary DEG |
| TAPBPL | 1783.48776047824 | 0.173509800932812 | 0.0509557529830286 | 3.40510719153932 | 0,00066138 | 0.0205504928798845 | Primary DEG |
| CAPRIN1 | 7363.70500200223 | -0.125009453861651 | 0.0367184357558475 | -3.40454192256115 | 0,00066275 | 0.0205608467416788 | Primary DEG |
| HEATR5A | 292.429983002821 | -0.217816882065496 | 0.0639999892929592 | -3.40338935165194 | 0,00066555 | 0.0206111148498589 | Primary DEG |
| RP5-1000K24.2 | 25.0504612112804 | 0.216320235048929 | 0.0635671128511446 | 3.40302123765612 | 0,00066645 | 0.0206111148498589 | Primary DEG |
| AVPR2 | 14.2197511283528 | 0.347804004057625 | 0.102218855561877 | 3.40254253626512 | 0,00066762 | 0.0206142071624277 | Primary DEG |
| TMEM33 | 1619.52157568941 | -0.194556226823905 | 0.057186610884716 | -3.40212899162912 | 0,00066863 | 0.0206142071624277 | Primary DEG |
| THNSL1 | 195.202770488321 | -0.302953363874876 | 0.0890827733206525 | -3.40080750275251 | 0,00067187 | 0.0206347589628903 | Primary DEG |
| RPS6KA5 | 631.268643720404 | -0.15690432449846 | 0.0461394776275542 | -3.40065238200177 | 0,00067225 | 0.0206347589628903 | Primary DEG |
| FAM50B | 196.251890194638 | -0.23736111602602 | 0.0698000977727516 | -3.40058429142603 | 0,00067242 | 0.0206347589628903 | Primary DEG |
| CYP4F22 | 269.771429352696 | 0.416680034736721 | 0.122625471371562 | 3.39798926011185 | 0,00067883 | 0.0207765292454328 | Primary DEG |
| VPS13B-DT | 28.6826657552758 | 0.289908271827739 | 0.0853206817571956 | 3.3978663303788 | 0,00067914 | 0.0207765292454328 | Primary DEG |
| LDHAP2 | 10.6394573821733 | 0.672591216251826 | 0.198016768860971 | 3.39663766922718 | 0,00068219 | 0.0208126492220164 | Primary DEG |
| NODAL | 9.06285954274948 | 0.511213231640194 | 0.150509653072851 | 3.39654780409832 | 0,00068242 | 0.0208126492220164 | Primary DEG |
| KIFBP | 211.231424959064 | -0.248675643397832 | 0.0732520827276812 | -3.39479280503597 | 0,00068681 | 0.0209112748932332 | Primary DEG |
| CES4A | 55.5779978290742 | 0.276438394100893 | 0.0814413963494303 | 3.39432286886161 | 0,00068799 | 0.0209112748932332 | Primary DEG |
| ASF1A | 1021.03127573893 | -0.183320564615181 | 0.0540153364388743 | -3.39386138643481 | 0,00068915 | 0.0209112748932332 | Primary DEG |
| TBC1D23 | 1158.04200428011 | -0.165456959152127 | 0.0487559624017043 | -3.39357385234064 | 0,00068987 | 0.0209112748932332 | Primary DEG |
| TNFAIP3 | 6507.50176755813 | 0.387621570178691 | 0.114316311969418 | 3.39078092619351 | 0,00069694 | 0.0210932767102454 | Primary DEG |
| GIMAP1 | 3206.74349286148 | -0.165102900042849 | 0.0487039925792445 | -3.38992536955191 | 0,00069912 | 0.0211269606324394 | Primary DEG |
| THAP2 | 159.818941783369 | -0.178872856109865 | 0.0527760436268185 | -3.38928126887801 | 0,00070076 | 0.0211401046824213 | Primary DEG |
| MT-CO3 | 155185.167787307 | 0.220339772063027 | 0.0650208168488135 | 3.38875736635794 | 0,0007021 | 0.0211401046824213 | Primary DEG |
| ABCD3 | 1079.79364197363 | -0.179700079071912 | 0.0530322796807722 | -3.38850375947662 | 0,00070275 | 0.0211401046824213 | Primary DEG |
| RARS1 | 1082.45417817243 | -0.222985592705263 | 0.0658272517515328 | -3.38743585326834 | 0,00070549 | 0.0211904180405423 | Primary DEG |
| HP1BP3 | 8861.84834261356 | 0.135134234320911 | 0.0399065760090839 | 3.38626481736121 | 0,00070851 | 0.0212194101060815 | Primary DEG |
| PARTICL | 54.8851576936872 | 0.235697306392868 | 0.0696046237399829 | 3.38623059400976 | 0,0007086 | 0.0212194101060815 | Primary DEG |
| MED14 | 1670.11929132733 | -0.154303641868327 | 0.0455817978077101 | -3.38520306985844 | 0,00071126 | 0.0212668806328124 | Primary DEG |
| KRAS | 2063.98315892874 | -0.117891885578734 | 0.0348300021937871 | -3.38477973451758 | 0,00071235 | 0.0212676268830505 | Primary DEG |
| IGLV1-47 | 433.798994362956 | -0.510381914614474 | 0.150856816852157 | -3.38322076034959 | 0,00071641 | 0.021356565073404 | Primary DEG |
| SRP68 | 2912.20273035137 | -0.0876198903561015 | 0.0259154571776157 | -3.38098956756135 | 0,00072225 | 0.0214540283099746 | Primary DEG |
| IGHV3OR16-12 | 2.67022958125123 | -0.912476536471935 | 0.269885071682402 | -3.38098187789256 | 0,00072227 | 0.0214540283099746 | Primary DEG |
| BUB1 | 157.865999084483 | -0.374582720721909 | 0.110800153970226 | -3.38070577792309 | 0,000723 | 0.0214540283099746 | Primary DEG |
| KLHL25 | 103.430591708316 | -0.212021681912663 | 0.0627256379826011 | -3.38014388903423 | 0,00072448 | 0.0214540283099746 | Primary DEG |
| AHCTF1 | 1722.63375788078 | -0.17339386476322 | 0.0513012975431212 | -3.37991187488921 | 0,00072509 | 0.0214540283099746 | Primary DEG |
| COPB2 | 5261.57600992639 | -0.155770972572613 | 0.0461169764299409 | -3.37773602328968 | 0,00073085 | 0.0215896961466498 | Primary DEG |
| FAM114A2 | 808.440019987638 | -0.131811213928626 | 0.0390278931407939 | -3.37735919930687 | 0,00073185 | 0.0215896961466498 | Primary DEG |
| EPB41L4A-AS1 | 664.919736030928 | 0.182646477690867 | 0.0540954792235838 | 3.37637230157375 | 0,00073449 | 0.0216351000622636 | Primary DEG |
| OSBP | 2273.75227808346 | -0.098393331670939 | 0.0291458324891808 | -3.37589711007444 | 0,00073575 | 0.0216403456307993 | Primary DEG |
| FOSL2 | 6107.93339063744 | 0.264973055141362 | 0.0785075373077368 | 3.37512886313973 | 0,00073781 | 0.0216487092883194 | Primary DEG |
| SLF2 | 1450.37187464026 | -0.188074293016921 | 0.0557261216440607 | -3.37497545977105 | 0,00073822 | 0.0216487092883194 | Primary DEG |
| WDR83 | 68.1047180385606 | 0.151156264087391 | 0.0448160779315199 | 3.37281330861574 | 0,00074404 | 0.0217871858381171 | Primary DEG |
| ABCC5 | 951.993006352527 | 0.197580545680841 | 0.0585957131665239 | 3.37192833747639 | 0,00074644 | 0.0218250742821337 | Primary DEG |
| FASTKD3 | 229.840409397547 | -0.130987936482767 | 0.0388576372599322 | -3.37097018036745 | 0,00074904 | 0.0218688690126702 | Primary DEG |
| RAB33A | 137.817773483052 | -0.22004812417933 | 0.0653140680079869 | -3.36907699811964 | 0,0007542 | 0.0219842840149185 | Primary DEG |
| ZNF691 | 191.6127623084 | -0.195050133342103 | 0.0579005559976337 | -3.36870915971989 | 0,00075521 | 0.0219842840149185 | Primary DEG |
| KLF10 | 5657.95672261609 | 0.315036825849905 | 0.0935826768835733 | 3.36640109410254 | 0,00076156 | 0.0221365732976337 | Primary DEG |
| GPRC5C | 20.3749006817438 | 0.450037926983132 | 0.133747016028945 | 3.36484461743608 | 0,00076587 | 0.0222292276556409 | Primary DEG |
| CCNJL | 123.359918129131 | 0.267699615374716 | 0.0795729988064439 | 3.36420166878313 | 0,00076765 | 0.0222396111185001 | Primary DEG |
| TMEM60 | 425.211525326977 | -0.232641388553805 | 0.0691662688773399 | -3.36350929911187 | 0,00076958 | 0.0222396111185001 | Primary DEG |
| RUNX3 | 11484.017687077 | 0.193662102011547 | 0.0575825313781992 | 3.36320924725563 | 0,00077042 | 0.0222396111185001 | Primary DEG |
| GGPS1 | 458.033816119809 | -0.161883658283388 | 0.0481351908557978 | -3.36310411167486 | 0,00077071 | 0.0222396111185001 | Primary DEG |
| UBXN8 | 184.564532001317 | -0.148673403894149 | 0.0442401844693149 | -3.36059638262288 | 0,00077774 | 0.0223957318793736 | Primary DEG |
| INPP5F | 411.217979661971 | -0.157529897348004 | 0.0468787417354409 | -3.36036957299365 | 0,00077838 | 0.0223957318793736 | Primary DEG |
| MMAA | 300.532787553743 | -0.136372475477689 | 0.0406047634672035 | -3.35853392146558 | 0,00078357 | 0.0224419679313392 | Primary DEG |
| MOCS3 | 286.033947049197 | -0.232605939302258 | 0.0692596575622626 | -3.35846216238005 | 0,00078378 | 0.0224419679313392 | Primary DEG |
| RNU6-5P | 13.6363338183372 | 0.457256415563452 | 0.136151221512446 | 3.35844519413037 | 0,00078382 | 0.0224419679313392 | Primary DEG |
| RP11-275I4.1 | 2.58100438739216 | 0.762782924887233 | 0.227140402115724 | 3.35820011667766 | 0,00078452 | 0.0224419679313392 | Primary DEG |
| KLHL7 | 774.621768773282 | -0.0955091296343407 | 0.0284761606830545 | -3.35400304477057 | 0,00079652 | 0.0227259835896993 | Primary DEG |
| TCTA | 1437.19566301375 | 0.156677152778774 | 0.0467145650720654 | 3.35392510958996 | 0,00079674 | 0.0227259835896993 | Primary DEG |
| DCUN1D4 | 491.056259187341 | -0.14191012779742 | 0.0423253080236949 | -3.35284335598868 | 0,00079986 | 0.0227314568067801 | Primary DEG |
| NR4A1 | 575.260433730684 | 0.394654990556439 | 0.117714908830538 | 3.3526338717603 | 0,00080047 | 0.0227314568067801 | Primary DEG |
| SLC25A17 | 398.055462086378 | -0.099245838907318 | 0.0296048255651768 | -3.35235344281365 | 0,00080128 | 0.0227314568067801 | Primary DEG |
| TP73-AS1 | 525.630401986985 | 0.163112725869944 | 0.0486607582297822 | 3.3520383118509 | 0,00080219 | 0.0227314568067801 | Primary DEG |
| PKNOX1 | 727.58556616774 | -0.118931024102593 | 0.0354819459733633 | -3.35187433608844 | 0,00080266 | 0.0227314568067801 | Primary DEG |
| NAT1 | 207.50692109594 | -0.194320270484887 | 0.0579841141679108 | -3.35126738199662 | 0,00080443 | 0.0227333734872131 | Primary DEG |
| TDG | 1053.95021325209 | -0.0958609008345431 | 0.0286061341375004 | -3.35106101278036 | 0,00080503 | 0.0227333734872131 | Primary DEG |
| LDAH | 556.853396784247 | -0.0946899514196033 | 0.0282647691582337 | -3.35010524549143 | 0,00080781 | 0.0227774001620899 | Primary DEG |
| CD80 | 22.9627335825416 | -0.313792582824008 | 0.093718565039606 | -3.34824357043236 | 0,00081326 | 0.0227774001620899 | Primary DEG |
| TMEM99 | 113.695341967545 | -0.222685428593002 | 0.0665094442254646 | -3.34817755863523 | 0,00081345 | 0.0227774001620899 | Primary DEG |
| SMTNL1 | 13.6469707051336 | 0.437011017719618 | 0.13055041642802 | 3.3474502010537 | 0,00081559 | 0.0227774001620899 | Primary DEG |
| LARS2 | 720.685164149128 | -0.0944405487046458 | 0.0282140059249063 | -3.34729314780774 | 0,00081605 | 0.0227774001620899 | Primary DEG |
| ASCC3 | 2168.8227004071 | -0.129806189744618 | 0.0387889722528221 | -3.3464714893335 | 0,00081847 | 0.0227774001620899 | Primary DEG |
| CD274 | 174.845004765177 | -0.235640048634965 | 0.070416902694515 | -3.34635633801201 | 0,00081881 | 0.0227774001620899 | Primary DEG |
| CRTC2 | 2638.85057548114 | 0.119645764397506 | 0.0357567083868696 | 3.34610678094301 | 0,00081955 | 0.0227774001620899 | Primary DEG |
| PAICS | 1188.89777760495 | -0.131471985086662 | 0.0392987648859294 | -3.34544827218563 | 0,0008215 | 0.0227774001620899 | Primary DEG |
| C10orf88 | 240.384960758586 | -0.134258135193938 | 0.0401319022003208 | -3.34542166787362 | 0,00082158 | 0.0227774001620899 | Primary DEG |
| SGIP1 | 2.33398823900742 | 1.23626428288016 | 0.369590586066787 | 3.34495609327225 | 0,00082296 | 0.0227774001620899 | Primary DEG |
| HEATR1 | 1268.5018818693 | -0.191444766291275 | 0.057234552873578 | -3.34491590620348 | 0,00082308 | 0.0227774001620899 | Primary DEG |
| UGDH | 602.910008617568 | -0.174209332251122 | 0.0520830436286806 | -3.34483778431087 | 0,00082331 | 0.0227774001620899 | Primary DEG |
| RAB8B | 4358.82893124833 | -0.167372024214871 | 0.0500399800343318 | -3.3447660071015 | 0,00082352 | 0.0227774001620899 | Primary DEG |
| CYTH1 | 11197.4346877911 | 0.101585159103132 | 0.0303722949665624 | 3.34466523570146 | 0,00082382 | 0.0227774001620899 | Primary DEG |
| CEP44 | 339.252334557074 | -0.138661362533242 | 0.0414785888883728 | -3.34296238732729 | 0,00082889 | 0.0228857129902488 | Primary DEG |
| SLC37A1 | 905.222513192029 | 0.106196176654327 | 0.0317788583325994 | 3.34172409665795 | 0,0008326 | 0.0229560751909449 | Primary DEG |
| RP11-10L12.8 | 4.66490199682669 | 0.554550909698343 | 0.166007893517655 | 3.34050928511641 | 0,00083625 | 0.023024718167999 | Primary DEG |
| EIF2AK3-DT | 67.1773107713292 | 0.23796956258679 | 0.0712500233059961 | 3.33992259293427 | 0,00083802 | 0.0230414116388541 | Primary DEG |
| SEC24D | 2103.57424837801 | -0.119473818662999 | 0.0357773562701542 | -3.3393696773136 | 0,00083969 | 0.0230553503572706 | Primary DEG |
| TBC1D16 | 153.135750587871 | 0.197956849861182 | 0.0593098376726845 | 3.33767310161349 | 0,00084483 | 0.0231644892662119 | Primary DEG |
| ADGRL1-AS1 | 97.2581059596354 | 0.173028880450075 | 0.0518531164427766 | 3.33690416931881 | 0,00084717 | 0.02318063986786 | Primary DEG |
| WDR13 | 1096.96955776612 | 0.138737030837721 | 0.0415810092824808 | 3.33654793935401 | 0,00084826 | 0.02318063986786 | Primary DEG |
| KMT5B | 1983.27489582287 | -0.112892854672957 | 0.03383745146286 | -3.33632852925902 | 0,00084893 | 0.02318063986786 | Primary DEG |
| SLC38A2 | 5198.80679458799 | -0.136661040083581 | 0.040967047493471 | -3.3358772097345 | 0,00085031 | 0.0231863662339835 | Primary DEG |
| CENPA | 17.8202427099803 | -0.429484864914173 | 0.12876436018504 | -3.33543275714635 | 0,00085167 | 0.0231915606887798 | Primary DEG |
| METTL2A | 285.417197317367 | -0.141481622242753 | 0.0424309092713631 | -3.33439996154501 | 0,00085484 | 0.0232459224067095 | Primary DEG |
| ZNF776 | 749.00770023235 | -0.181953008721813 | 0.0546081557157492 | -3.33197498316789 | 0,00086232 | 0.0234173057052543 | Primary DEG |
| CLCN3 | 2193.82615697183 | -0.144525203519471 | 0.0433842493802106 | -3.33128279465854 | 0,00086447 | 0.0234435026078138 | Primary DEG |
| SLC35B3 | 517.064583186095 | -0.118193415261858 | 0.0355034769175866 | -3.32906592602798 | 0,00087138 | 0.0235543409250847 | Primary DEG |
| SNPH | 456.276563660682 | 0.245819873591883 | 0.0738426940653711 | 3.32896675430429 | 0,00087169 | 0.0235543409250847 | Primary DEG |
| VPS26B | 3860.81997539031 | 0.105387123655003 | 0.0316608157115266 | 3.32862945210332 | 0,00087274 | 0.0235543409250847 | Primary DEG |
| PRPF40A | 3792.56757290001 | -0.122701294759203 | 0.0368670851427507 | -3.3282071062602 | 0,00087407 | 0.0235543409250847 | Primary DEG |
| RUNDC1 | 838.850890554646 | -0.122516929763225 | 0.0368131943089465 | -3.32807114576988 | 0,0008745 | 0.0235543409250847 | Primary DEG |
| GNAS | 31716.7687077575 | 0.112099387299908 | 0.0336893192567757 | 3.3274459019341 | 0,00087646 | 0.0235734619499391 | Primary DEG |
| YIPF6 | 672.790829612323 | -0.144370053870449 | 0.0433971700812018 | -3.32671585728548 | 0,00087876 | 0.0235734619499391 | Primary DEG |
| LINC01215 | 1039.53882355728 | 0.249127433699084 | 0.0748876330787154 | 3.32668323803508 | 0,00087886 | 0.0235734619499391 | Primary DEG |
| UTP11 | 658.90956542954 | -0.144419913770895 | 0.0434171314868594 | -3.32633476291738 | 0,00087996 | 0.0235734619499391 | Primary DEG |
| SUSD6 | 4528.65298211133 | 0.114186511176818 | 0.0343345933679782 | 3.32569866062001 | 0,00088197 | 0.023595435416309 | Primary DEG |
| PDP2 | 166.650334273585 | -0.254067494492337 | 0.0764223068477224 | -3.32452009069272 | 0,00088571 | 0.0236634577246249 | Primary DEG |
| ZNF28 | 564.284405002316 | -0.241824959550641 | 0.0727693580411994 | -3.3231701647516 | 0,00089001 | 0.0237462717788252 | Primary DEG |
| QPRT | 101.744347577549 | 0.207854890660271 | 0.0625695014389717 | 3.32198412773044 | 0,0008938 | 0.0238098164109284 | Primary DEG |
| MAPK3 | 2396.110589662 | 0.123083073496474 | 0.0370606635686583 | 3.32112438484678 | 0,00089656 | 0.0238098164109284 | Primary DEG |
| GTF2H1 | 1459.94294900606 | -0.1012106959869 | 0.030479788504427 | -3.32058393292986 | 0,00089829 | 0.0238098164109284 | Primary DEG |
| USP16 | 1485.66544325796 | -0.117877712358732 | 0.0355017152934016 | -3.32033850715493 | 0,00089908 | 0.0238098164109284 | Primary DEG |
| IGF1 | 2.65838136717397 | -0.890152162020357 | 0.268127929306791 | -3.31987855320304 | 0,00090057 | 0.0238098164109284 | Primary DEG |
| RP11-446N19.1 | 18.1558507727665 | 0.312372273200134 | 0.0940939114174844 | 3.31979262520157 | 0,00090084 | 0.0238098164109284 | Primary DEG |
| RP4-671O14.7 | 17.0269000504649 | 0.427275667190179 | 0.128709281737473 | 3.31969584028671 | 0,00090116 | 0.0238098164109284 | Primary DEG |
| WDR47 | 571.807682246985 | -0.126072529002258 | 0.0379801102973204 | -3.31943556812558 | 0,000902 | 0.0238098164109284 | Primary DEG |
| MLYCD | 320.034980401691 | -0.178986506754593 | 0.0539322511463941 | -3.31872864473525 | 0,00090428 | 0.0238135767700984 | Primary DEG |
| LIN9 | 159.716931270035 | -0.131356845727233 | 0.0395888981573326 | -3.31802226991012 | 0,00090657 | 0.0238135767700984 | Primary DEG |
| MED23 | 1578.31713450529 | -0.166384035805503 | 0.0501543363824159 | -3.31744068024071 | 0,00090846 | 0.0238135767700984 | Primary DEG |
| CCDC47 | 2084.77472304744 | -0.1076872983014 | 0.0324615131431153 | -3.31738381469255 | 0,00090865 | 0.0238135767700984 | Primary DEG |
| RP11-434H6.6 | 61.3146698008866 | 0.308198516000191 | 0.0929070626799243 | 3.31727757944486 | 0,00090899 | 0.0238135767700984 | Primary DEG |
| RP11-434H14.1 | 2.88120936491557 | 0.840196899420782 | 0.253287340618586 | 3.31716894089073 | 0,00090935 | 0.0238135767700984 | Primary DEG |
| CILP | 6.67025288783135 | 0.559557389362556 | 0.168715795994664 | 3.31656787714324 | 0,0009113 | 0.0238333772106371 | Primary DEG |
| RP11-660L16.2 | 209.049128604787 | 0.345528538848491 | 0.10421116037239 | 3.31565772431448 | 0,00091428 | 0.0238796226726571 | Primary DEG |
| IRAK1 | 3797.46927938141 | 0.125836707144878 | 0.0379579678824351 | 3.31515921860266 | 0,00091591 | 0.0238907737138765 | Primary DEG |
| MED10 | 1044.53314390351 | 0.116326658421652 | 0.0351001201954725 | 3.31413846373827 | 0,00091926 | 0.0239164754088395 | Primary DEG |
| CTB-50L17.9 | 4.17374011740988 | 0.648788508486258 | 0.195764711383487 | 3.31412389853724 | 0,00091931 | 0.0239164754088395 | Primary DEG |
| FOXO4 | 1375.75104424875 | 0.134770997303159 | 0.0406776452697219 | 3.31314648155101 | 0,00092253 | 0.0239687618991085 | Primary DEG |
| FAM219A | 492.314035501504 | 0.0947394078722086 | 0.028601609091804 | 3.31238034783703 | 0,00092506 | 0.0239793190353996 | Primary DEG |
| TIPRL | 1406.15191435943 | -0.118470393207741 | 0.0357678238092013 | -3.3122057925499 | 0,00092563 | 0.0239793190353996 | Primary DEG |
| BIRC2 | 2567.69959242619 | -0.121120008771268 | 0.0365708718883966 | -3.31192565331476 | 0,00092656 | 0.0239793190353996 | Primary DEG |
| PGM3 | 284.790406771468 | -0.144451547849442 | 0.0436214572844472 | -3.31147918574799 | 0,00092804 | 0.0239863017090668 | Primary DEG |
| TRAPPC8 | 2724.19714583671 | -0.106826165357272 | 0.0322870991720703 | -3.3086331103316 | 0,00093753 | 0.0241915871822726 | Primary DEG |
| ZNF45 | 661.642714227863 | -0.104465408217447 | 0.0315761449990258 | -3.30836485013194 | 0,00093843 | 0.0241915871822726 | Primary DEG |
| BLACAT1 | 2.9671873897572 | 0.542679257815669 | 0.164158599940938 | 3.30582289329294 | 0,00094698 | 0.0243138880184623 | Primary DEG |
| KRT8P33 | 24.7933268037064 | 0.312328879686355 | 0.0945011269551833 | 3.30502809595567 | 0,00094967 | 0.0243138880184623 | Primary DEG |
| RP11-83M8.1 | 8.40148349711693 | 0.32762716596272 | 0.0991354658501217 | 3.30484315732216 | 0,0009503 | 0.0243138880184623 | Primary DEG |
| PER1 | 1267.10888847915 | 0.378658801920979 | 0.114592268138302 | 3.30440096939154 | 0,0009518 | 0.0243138880184623 | Primary DEG |
| PGAM1P8 | 52.3361614849976 | 0.407088663638531 | 0.123213385408101 | 3.30393213602721 | 0,00095339 | 0.0243138880184623 | Primary DEG |
| RP11-394B2.5 | 8.35536325830139 | 0.419461869178098 | 0.126960271425582 | 3.30388289555577 | 0,00095356 | 0.0243138880184623 | Primary DEG |
| HSP90AA2P | 15.3893996549325 | -0.278298116907961 | 0.084234358185264 | -3.30385513588025 | 0,00095365 | 0.0243138880184623 | Primary DEG |
| ARR3 | 5.12154252808254 | 0.521950730075382 | 0.157995034291163 | 3.30358946037188 | 0,00095456 | 0.0243138880184623 | Primary DEG |
| TOR1AIP1 | 4183.83012252269 | -0.131122951325244 | 0.0396958857992441 | -3.30318743832493 | 0,00095593 | 0.0243138880184623 | Primary DEG |
| STAG3L2 | 98.9298136046925 | 0.281903627448268 | 0.0853448152160705 | 3.30311368926821 | 0,00095618 | 0.0243138880184623 | Primary DEG |
| N6AMT1 | 142.350045200716 | -0.216812135194313 | 0.0656415348024135 | -3.3029717517565 | 0,00095666 | 0.0243138880184623 | Primary DEG |
| ZNF212 | 405.746456611776 | -0.193613143945223 | 0.0586281999522089 | -3.30238936387349 | 0,00095865 | 0.0243332358899924 | Primary DEG |
| KPNB1 | 6603.11632715533 | -0.105619634738256 | 0.0320095426692377 | -3.2996296082593 | 0,00096813 | 0.0245423430820691 | Primary DEG |
| GMPPB | 378.545055956477 | -0.135029027194467 | 0.0409316237716818 | -3.2988925127345 | 0,00097067 | 0.0245708869371152 | Primary DEG |
| TTC9C | 499.705927107193 | -0.177345369293643 | 0.0537640529049848 | -3.29858631764719 | 0,00097173 | 0.0245708869371152 | Primary DEG |
| TRMT2B | 1295.55735330779 | 0.105849525165668 | 0.0321035622383835 | 3.29712710320703 | 0,00097679 | 0.0246456996450993 | Primary DEG |
| C2orf81 | 17.7266379635289 | 0.247945633945371 | 0.0752030147955863 | 3.29701720894205 | 0,00097718 | 0.0246456996450993 | Primary DEG |
| PRR34 | 60.7391716399413 | 0.174317837145014 | 0.0529076486660954 | 3.29475683648594 | 0,00098507 | 0.0247806468952613 | Primary DEG |
| BAK1 | 776.018867901501 | -0.167775583865434 | 0.0509253947178444 | -3.29453673938132 | 0,00098584 | 0.0247806468952613 | Primary DEG |
| ZNF184 | 377.295235921713 | -0.16333527214912 | 0.0495838430932478 | -3.2941228827695 | 0,00098729 | 0.0247806468952613 | Primary DEG |
| ZFAND2A-DT | 79.9252324273602 | 0.282516833215872 | 0.0857656256048404 | 3.29405669489954 | 0,00098753 | 0.0247806468952613 | Primary DEG |
| TAF6 | 835.401304668747 | 0.0839770383003381 | 0.0255050089837576 | 3.29257042621774 | 0,00099276 | 0.0248805060789683 | Primary DEG |
| TMEM41B | 270.958233116516 | -0.295447456116118 | 0.0897595108824215 | -3.29154485370507 | 0,00099639 | 0.0249188717321393 | Primary DEG |
| LRRC8C | 3236.1051566385 | -0.130998660814444 | 0.039799960414916 | -3.29142691220741 | 0,00099681 | 0.0249188717321393 | Primary DEG |
| S100PBP | 963.490567974516 | -0.126649580145984 | 0.0384847445655302 | -3.29090348853246 | 0,00099866 | 0.0249338405272945 | Primary DEG |
| BBS10 | 309.910596710386 | -0.276248600253536 | 0.083966354620001 | -3.28999158655546 | 0,0010019 | 0.0249642595416868 | Primary DEG |
| LIN7C | 556.200465277872 | -0.144479094127939 | 0.0439165853153031 | -3.28985263974051 | 0,0010024 | 0.0249642595416868 | Primary DEG |
| TSNARE1 | 372.263301144896 | 0.183327125553592 | 0.0557315830059896 | 3.28946560756922 | 0,00100378 | 0.0249672517328633 | Primary DEG |
| CDC37L1 | 428.053606658908 | -0.155967004114491 | 0.0474373301245558 | -3.28785375789425 | 0,00100954 | 0.0250791650045104 | Primary DEG |
| RPRD1B | 905.588703977821 | -0.182630225487253 | 0.0555598203345997 | -3.2870917218125 | 0,00101228 | 0.0251156575258066 | Primary DEG |
| ATXN7L3 | 3847.43001391673 | 0.0588651965131837 | 0.0179100470439931 | 3.28671367353704 | 0,00101364 | 0.0251179597516388 | Primary DEG |
| STX1B | 21.4090088400287 | 0.367312113486756 | 0.111771540780274 | 3.28627583481951 | 0,00101522 | 0.0251256312083822 | Primary DEG |
| CEP76 | 172.788470232011 | -0.167843509931595 | 0.0510929647819254 | -3.28506107735151 | 0,0010196 | 0.0252027471910975 | Primary DEG |
| PNPLA8 | 1927.06084734114 | -0.14807779401336 | 0.0450848133776068 | -3.28442734747814 | 0,0010219 | 0.0252280375888701 | Primary DEG |
| UBE2F | 789.709672873912 | 0.114798631577006 | 0.0349641184049591 | 3.28332693098086 | 0,0010259 | 0.0252642200619957 | Primary DEG |
| ANGEL2 | 959.145182524063 | -0.167567797731923 | 0.0510360484856588 | -3.2833223320377 | 0,00102591 | 0.0252642200619957 | Primary DEG |
| FOXN3 | 7258.86861812059 | 0.130434972492891 | 0.0397327833442168 | 3.28280481543148 | 0,0010278 | 0.0252792406711173 | Primary DEG |
| NEDD1 | 937.633803448058 | -0.128756611675619 | 0.0392329608239643 | -3.2818479403923 | 0,00103129 | 0.0253337461378188 | Primary DEG |
| SEC14L2 | 58.8776746129165 | 0.370377269808916 | 0.112936222516246 | 3.27952592672945 | 0,00103982 | 0.0255115388014865 | Primary DEG |
| SLA | 5083.96622232249 | -0.169650871931269 | 0.0517373400514459 | -3.27907990172232 | 0,00104146 | 0.0255203109205006 | Primary DEG |
| VMA21 | 1113.30858727391 | -0.118984324229272 | 0.0363026397340894 | -3.27756673070641 | 0,00104706 | 0.0256030987531664 | Primary DEG |
| ARRDC5 | 62.3017783742898 | 0.309681828167448 | 0.0944913265209126 | 3.27735718789925 | 0,00104784 | 0.0256030987531664 | Primary DEG |
| SOAT1 | 2935.73218718827 | -0.157788392520346 | 0.0481540631256407 | -3.27674099086205 | 0,00105013 | 0.0256030987531664 | Primary DEG |
| KRBA1 | 255.697717766132 | 0.156150657866343 | 0.0476548153584731 | 3.27670261004546 | 0,00105027 | 0.0256030987531664 | Primary DEG |
| CYTH2 | 922.063899378867 | 0.0851696802625507 | 0.0259974273492716 | 3.27608109519102 | 0,00105258 | 0.0256030987531664 | Primary DEG |
| CA14 | 23.9444986920044 | 0.354823447520024 | 0.108328590255264 | 3.27543676774455 | 0,00105499 | 0.0256030987531664 | Primary DEG |
| TM2D2 | 829.654174804459 | -0.12252085890267 | 0.0374080345089711 | -3.27525518276794 | 0,00105567 | 0.0256030987531664 | Primary DEG |
| TARDBP | 3123.74387960461 | -0.118557745660808 | 0.0361991497833015 | -3.27515276934759 | 0,00105605 | 0.0256030987531664 | Primary DEG |
| CSTF3 | 445.992448575456 | -0.129050267300849 | 0.0394045743225238 | -3.27500726805425 | 0,00105659 | 0.0256030987531664 | Primary DEG |
| RRP8 | 578.363189097948 | -0.137882759907865 | 0.0421055069501357 | -3.27469658710333 | 0,00105776 | 0.0256030987531664 | Primary DEG |
| RABL2B | 494.455432035354 | 0.120422940520609 | 0.0367859957307723 | 3.27360828838112 | 0,00106184 | 0.0256705559511412 | Primary DEG |
| AASDH | 608.373956054114 | -0.116842823437084 | 0.035726792253112 | -3.27045379863082 | 0,00107375 | 0.0259269621611363 | Primary DEG |
| UBR1 | 1602.42726101563 | -0.136485518834714 | 0.0417436754710304 | -3.26960952275115 | 0,00107696 | 0.0259728251904099 | Primary DEG |
| TSPAN17 | 1307.9427234103 | 0.11202954270355 | 0.0342677828517452 | 3.26923814091593 | 0,00107838 | 0.0259753415324322 | Primary DEG |
| ZNF235 | 160.213185833489 | -0.184582978946476 | 0.0564687810964702 | -3.26876152384338 | 0,00108019 | 0.0259875561890344 | Primary DEG |
| ACTL6A | 951.558439581145 | -0.111251290082607 | 0.0340447431581539 | -3.26779642794755 | 0,00108388 | 0.0260447222519317 | Primary DEG |
| TP53RK | 852.872221689059 | -0.184618458538644 | 0.056515736376186 | -3.26667350328354 | 0,00108819 | 0.0261165846728368 | Primary DEG |
| MSANTD4 | 556.741170100448 | -0.171761530734397 | 0.0525916619289941 | -3.26594605369762 | 0,00109099 | 0.0261521063501725 | Primary DEG |
| LBR | 6254.76737794993 | -0.125845270791515 | 0.0385489611327215 | -3.26455673755354 | 0,00109636 | 0.0262489638113428 | Primary DEG |
| SMC4 | 964.357984603789 | -0.166693984009398 | 0.0510742639135111 | -3.26375695382859 | 0,00109945 | 0.0262914154645793 | Primary DEG |
| GPR171 | 669.318738532663 | -0.24996425523853 | 0.0766184205526062 | -3.26245638366957 | 0,00110451 | 0.0263262533627246 | Primary DEG |
| WDR76 | 410.74430696005 | -0.150066978733748 | 0.0459991012576584 | -3.2623893648088 | 0,00110477 | 0.0263262533627246 | Primary DEG |
| ZNF222 | 91.1985132815575 | -0.238210584590971 | 0.0730179172208152 | -3.26235797538011 | 0,0011049 | 0.0263262533627246 | Primary DEG |
| MT-RNR1 | 92718.6943985797 | 0.319767054290253 | 0.0981572458210926 | 3.25770198231804 | 0,00112318 | 0.0267209969583347 | Primary DEG |
| NOX1 | 4.8944743323512 | -0.478279691557282 | 0.146847197606769 | -3.25698889289009 | 0,00112601 | 0.0267209969583347 | Primary DEG |
| LAS1L | 1322.34338454872 | 0.0854813820598318 | 0.0262466372469325 | 3.25685082075884 | 0,00112656 | 0.0267209969583347 | Primary DEG |
| FYN | 12563.4170338464 | 0.100138845477357 | 0.0307507405230409 | 3.25646939794264 | 0,00112807 | 0.0267209969583347 | Primary DEG |
| RP11-568N6.1 | 94.6008310754116 | -0.203213121223454 | 0.0624102809762698 | -3.25608406250761 | 0,0011296 | 0.0267209969583347 | Primary DEG |
| ZNF8-ERVK3-1 | 16.362939061733 | 0.293254498755598 | 0.0900706501554649 | 3.25582748930346 | 0,00113062 | 0.0267209969583347 | Primary DEG |
| RP11-188C12.2 | 15.9820355188106 | 0.518482638704178 | 0.15926251588182 | 3.25552209089121 | 0,00113184 | 0.0267209969583347 | Primary DEG |
| RBPJ | 2892.0812481924 | -0.212699677108848 | 0.0653370809337885 | -3.25542056775378 | 0,00113225 | 0.0267209969583347 | Primary DEG |
| C5orf51 | 1081.57954931044 | -0.154711211605132 | 0.047541128574514 | -3.25426038977271 | 0,00113688 | 0.0267966145801119 | Primary DEG |
| ZW10 | 698.658725540752 | -0.122726426339804 | 0.0377162248786015 | -3.25394248058569 | 0,00113815 | 0.0267966145801119 | Primary DEG |
| NUP155 | 975.020442619678 | -0.138576047823749 | 0.0426121125584673 | -3.25203421054544 | 0,00114582 | 0.0269451734289209 | Primary DEG |
| LINC02390 | 16.4188258154782 | 0.510774523128692 | 0.157122652978461 | 3.25080129087885 | 0,0011508 | 0.0270302260005642 | Primary DEG |
| PRMT9 | 343.518685946988 | -0.155114310999482 | 0.0477209021989945 | -3.25044799766486 | 0,00115223 | 0.0270318044093222 | Primary DEG |
| MRPL30 | 836.195955805989 | -0.0875350235055996 | 0.0269370247876896 | -3.24961736478053 | 0,0011556 | 0.0270788299645531 | Primary DEG |
| ARMC2 | 126.600065849867 | 0.150748602400389 | 0.0464140111505847 | 3.24791153928289 | 0,00116255 | 0.0271645341167386 | Primary DEG |
| SLC26A2 | 668.720195567312 | -0.189439435500622 | 0.0583287499605968 | -3.24778836557608 | 0,00116306 | 0.0271645341167386 | Primary DEG |
| PPME1 | 536.352442333835 | -0.136522069390626 | 0.0420384392434719 | -3.24755323574071 | 0,00116402 | 0.0271645341167386 | Primary DEG |
| BBS7 | 291.878266998157 | -0.140263046099161 | 0.04319272822979 | -3.24737639523362 | 0,00116474 | 0.0271645341167386 | Primary DEG |
| ARHGAP25 | 5847.49690979551 | -0.131784639754702 | 0.0405910007293832 | -3.24664672924177 | 0,00116773 | 0.0272022639157672 | Primary DEG |
| IGLV10-54 | 201.933526722443 | -0.76537503076914 | 0.235826069975764 | -3.24550644823873 | 0,00117242 | 0.0272686539044682 | Primary DEG |
| CENPL | 169.940008489337 | -0.14250688337321 | 0.0439119849591871 | -3.24528448225831 | 0,00117333 | 0.0272686539044682 | Primary DEG |
| MORF4L2 | 2133.73009695597 | -0.0977704949178943 | 0.0301365205881089 | -3.24425292004254 | 0,00117759 | 0.0273355625842511 | Primary DEG |
| SORBS3 | 955.939796078302 | 0.154838617512442 | 0.0477382328372572 | 3.24349286326322 | 0,00118074 | 0.0273763117658379 | Primary DEG |
| PDAP1 | 2407.20376515431 | 0.0875516742063866 | 0.0269957736531157 | 3.24316225685504 | 0,00118211 | 0.0273763117658379 | Primary DEG |
| LDHD | 82.6810696029104 | 0.318688990804373 | 0.0982842577429905 | 3.24252324963101 | 0,00118476 | 0.0274057619382972 | Primary DEG |
| INIP | 1422.31882414626 | -0.147944335895786 | 0.0456407759381779 | -3.24149475671014 | 0,00118905 | 0.0274727880596271 | Primary DEG |
| JCHAIN | 8332.02310072045 | -0.520801038533524 | 0.160723526559071 | -3.24035347956426 | 0,00119382 | 0.0275501960791998 | Primary DEG |
| SLC4A7 | 1175.56807331734 | -0.203321687476616 | 0.062753048084502 | -3.24002887003715 | 0,00119518 | 0.0275501960791998 | Primary DEG |
| CKAP2 | 977.84293978404 | -0.133222986729084 | 0.0411395167302707 | -3.23832162644384 | 0,00120235 | 0.0276569207993979 | Primary DEG |
| CWC22 | 1255.4202869284 | -0.136086868560569 | 0.0420261243514605 | -3.23814938114415 | 0,00120308 | 0.0276569207993979 | Primary DEG |
| FPGT | 314.511756462091 | -0.250498301098665 | 0.0773653715183201 | -3.23786076616134 | 0,0012043 | 0.0276569207993979 | Primary DEG |
| CDH23 | 779.431772456187 | 0.212342824861532 | 0.0655870248378482 | 3.23757367233092 | 0,00120551 | 0.0276569207993979 | Primary DEG |
| RNF145 | 2446.74179929843 | -0.157731137452385 | 0.0487234674482089 | -3.23727242155019 | 0,00120678 | 0.0276569207993979 | Primary DEG |
| NDEL1 | 1696.52109711104 | -0.0984903975643671 | 0.0304306643503861 | -3.23655101414562 | 0,00120984 | 0.0276645253373618 | Primary DEG |
| IFNAR1 | 3487.51690978755 | -0.136752912635245 | 0.0422528775713806 | -3.23653489408429 | 0,0012099 | 0.0276645253373618 | Primary DEG |
| RP11-617F9.2 | 32.0326011792236 | 0.322038217145718 | 0.099533817581248 | 3.23546534204662 | 0,00121445 | 0.0276924432575177 | Primary DEG |
| LSM14B | 1577.64381485289 | 0.064289934471878 | 0.01987150868486 | 3.2352820055812 | 0,00121523 | 0.0276924432575177 | Primary DEG |
| KRT18P63 | 7.70870085527938 | 0.393609794395961 | 0.121671865439958 | 3.23501076417866 | 0,00121638 | 0.0276924432575177 | Primary DEG |
| C9orf40 | 84.0081883435428 | -0.254202876591411 | 0.0785805703563463 | -3.23493295401974 | 0,00121671 | 0.0276924432575177 | Primary DEG |
| RP11-429O22.1 | 19.8820521387327 | 0.412669977271335 | 0.127638513291809 | 3.23311488537855 | 0,00122448 | 0.0278373311749487 | Primary DEG |
| LINC00852 | 62.0742247270989 | 0.355492042228915 | 0.10999873770218 | 3.23178292455868 | 0,00123021 | 0.0279345179259148 | Primary DEG |
| POLR3H | 458.052501230619 | 0.111122814802338 | 0.0343877716503197 | 3.23146308903983 | 0,00123158 | 0.0279345179259148 | Primary DEG |
| LINC01948 | 62.9422424046263 | 0.223561871460446 | 0.0691898319430477 | 3.23113765682314 | 0,00123299 | 0.0279345179259148 | Primary DEG |
| NFS1 | 329.254496013813 | -0.134962537306386 | 0.0417803313211109 | -3.23028882344433 | 0,00123665 | 0.0279856092665629 | Primary DEG |
| FNTB | 153.456994913907 | -0.20695220671668 | 0.0640778096177787 | -3.22970163854134 | 0,0012392 | 0.0280111682202112 | Primary DEG |
| ST8SIA4 | 2864.36655043862 | -0.149582431897678 | 0.0463403588211077 | -3.22790836547308 | 0,00124699 | 0.0281063556949768 | Primary DEG |
| PPP6R2 | 1982.3952556501 | 0.113848140090187 | 0.0352701129084376 | 3.2278927029732 | 0,00124706 | 0.0281063556949768 | Primary DEG |
| DAZAP1 | 2999.31176984341 | 0.101660437182107 | 0.031497370116194 | 3.22758493191911 | 0,0012484 | 0.0281063556949768 | Primary DEG |
| ZNF180 | 528.815065818436 | -0.0973948771092772 | 0.030177225547853 | -3.22742980314194 | 0,00124908 | 0.0281063556949768 | Primary DEG |
| ZNF770 | 1176.22201765255 | -0.208577759136646 | 0.0646616665345123 | -3.22567867973709 | 0,00125674 | 0.0282468090534352 | Primary DEG |
| DUS4L-BCAP29 | 190.085683742105 | 0.149054827186168 | 0.0462370317593927 | 3.2237109847755 | 0,00126541 | 0.0284093955440166 | Primary DEG |
| ZNF77 | 131.701616008692 | 0.16459380846431 | 0.0510635382603822 | 3.22331381787561 | 0,00126717 | 0.0284166375363481 | Primary DEG |
| FNTA | 2071.75544499877 | -0.102792171020474 | 0.0319011975898306 | -3.22220414236869 | 0,00127209 | 0.0284946962100556 | Primary DEG |
| MCM6 | 1247.59213139405 | -0.152657803167367 | 0.047397726875984 | -3.22078321533848 | 0,00127841 | 0.02860402035139 | Primary DEG |
| DHRS12 | 127.571879112757 | 0.141895541497289 | 0.0440767017951499 | 3.21928673694234 | 0,0012851 | 0.0287213132726058 | Primary DEG |
| CHAMP1 | 1010.33423330175 | -0.164092138468731 | 0.0510039004065541 | -3.2172468607449 | 0,00129427 | 0.0288642640581309 | Primary DEG |
| USP8 | 1899.20922606057 | -0.139095177467275 | 0.0432400322818789 | -3.216814838632 | 0,00129622 | 0.0288642640581309 | Primary DEG |
| SEC62 | 3659.16209177174 | -0.127797037415776 | 0.0397326600969375 | -3.21642288998482 | 0,00129799 | 0.0288642640581309 | Primary DEG |
| TCTE1 | 4.76482035254449 | 0.595244123001737 | 0.185067451228059 | 3.21636311005449 | 0,00129827 | 0.0288642640581309 | Primary DEG |
| AC007238.1 | 6.41552385936634 | -0.357542770865313 | 0.111168856026244 | -3.21621345802917 | 0,00129894 | 0.0288642640581309 | Primary DEG |
| MAK16 | 389.624165984442 | -0.134510734824404 | 0.041826404862249 | -3.2159286763326 | 0,00130023 | 0.0288642640581309 | Primary DEG |
| CTC-523E23.11 | 304.788523745968 | 0.236856487331275 | 0.0736795198600157 | 3.21468554329997 | 0,00130588 | 0.0289571185571843 | Primary DEG |
| MIA2 | 1679.99963949871 | -0.0851110562533433 | 0.0264823226682315 | -3.21388185317458 | 0,00130954 | 0.0290058443478668 | Primary DEG |
| EHHADH | 88.3607263624987 | -0.260788675152916 | 0.0811806789764579 | -3.21244757300618 | 0,00131609 | 0.0290782868144506 | Primary DEG |
| PACS2 | 1331.65014186074 | 0.143635595005797 | 0.0447133066625638 | 3.21236798901424 | 0,00131646 | 0.0290782868144506 | Primary DEG |
| TMEM68 | 348.660131023657 | -0.120113397364414 | 0.0373944207354805 | -3.21206733523347 | 0,00131784 | 0.0290782868144506 | Primary DEG |
| FBXW4 | 1127.75663080911 | 0.137227315972145 | 0.0427262064283643 | 3.21178329281875 | 0,00131914 | 0.0290782868144506 | Primary DEG |
| TMEM168 | 672.802297752175 | -0.130459565110515 | 0.0406218019252068 | -3.21156519227577 | 0,00132014 | 0.0290782868144506 | Primary DEG |
| CTD-2561B21.11 | 10.6596540245456 | 0.359862585797756 | 0.112085018634662 | 3.21062163508861 | 0,00132448 | 0.0291415442804304 | Primary DEG |
| MYORG | 41.9900237971303 | -0.234629642443832 | 0.0731129008075922 | -3.20914147643103 | 0,00133132 | 0.0292595132937011 | Primary DEG |
| FICD | 116.213229512728 | -0.19884335364685 | 0.0619895938091784 | -3.207689249569 | 0,00133806 | 0.029345588585137 | Primary DEG |
| RNF166 | 2854.20019289254 | 0.148558775452119 | 0.0463149081297593 | 3.20758005253742 | 0,00133857 | 0.029345588585137 | Primary DEG |
| IGKV2-30 | 246.867026506347 | -0.521206950490479 | 0.162504334478092 | -3.20734183592343 | 0,00133968 | 0.029345588585137 | Primary DEG |
| KPNA2 | 1022.51011031869 | -0.124235879846371 | 0.0387453211561779 | -3.20647438552879 | 0,00134372 | 0.0293752860592476 | Primary DEG |
| TUNAR | 4.9648778130483 | 0.772213949780425 | 0.240834003482982 | 3.20641578270732 | 0,001344 | 0.0293752860592476 | Primary DEG |
| MTAP | 443.168001995654 | -0.132672947152665 | 0.0413987818423782 | -3.20475485626132 | 0,00135178 | 0.0295127743082917 | Primary DEG |
| CCR9 | 116.888045671674 | -0.371778460811886 | 0.116100928293282 | -3.20220058768814 | 0,00136382 | 0.0297429776966573 | Primary DEG |
| TMEM81 | 62.6070254314344 | -0.187702863234798 | 0.0586616230597593 | -3.19975570814982 | 0,00137544 | 0.0298864917892445 | Primary DEG |
| RRP15 | 414.19204842811 | -0.120113075466118 | 0.0375393408889383 | -3.19965861471775 | 0,00137591 | 0.0298864917892445 | Primary DEG |
| HDAC2 | 2246.53265177004 | -0.0890325704503858 | 0.0278275825244172 | -3.19943604056387 | 0,00137697 | 0.0298864917892445 | Primary DEG |
| PDCD4-AS1 | 220.534827244618 | 0.153982020674474 | 0.0481317886794731 | 3.19917511688369 | 0,00137821 | 0.0298864917892445 | Primary DEG |
| NECTIN1 | 611.674391181003 | 0.198348365712923 | 0.0620138014210927 | 3.19845520138456 | 0,00138166 | 0.0298864917892445 | Primary DEG |
| NGRN | 218.795501290334 | -0.150812720624541 | 0.0471561547530723 | -3.19815560480396 | 0,0013831 | 0.0298864917892445 | Primary DEG |
| IGKV1-5 | 1852.98150294833 | -0.446478093151384 | 0.139606422490458 | -3.19812000899815 | 0,00138327 | 0.0298864917892445 | Primary DEG |
| UPF3A | 1002.10621623788 | 0.180869172536362 | 0.0565579374327811 | 3.19794498785116 | 0,00138411 | 0.0298864917892445 | Primary DEG |
| UTP25 | 903.429192853525 | -0.142637616018196 | 0.0446060055894301 | -3.19772223792205 | 0,00138518 | 0.0298864917892445 | Primary DEG |
| SKOR1 | 10.7388426548142 | 0.405132991194325 | 0.126709967251968 | 3.19732535632893 | 0,00138708 | 0.0298864917892445 | Primary DEG |
| ARHGAP9 | 4589.62999165822 | 0.126653810345722 | 0.039624346357648 | 3.19636339745644 | 0,00139172 | 0.0298864917892445 | Primary DEG |
| EIF3J | 1694.6418404347 | -0.0895911604462759 | 0.0280328853617612 | -3.19593075383116 | 0,00139381 | 0.0298864917892445 | Primary DEG |
| MIS12 | 733.573916594693 | -0.141274417954429 | 0.0442104589672689 | -3.19549765495584 | 0,0013959 | 0.0298864917892445 | Primary DEG |
| LAMB2 | 198.203508025141 | 0.320417315710281 | 0.100289250441843 | 3.19493180274679 | 0,00139864 | 0.0298864917892445 | Primary DEG |
| PCDHGB3 | 6.28480589724423 | 0.449186265101005 | 0.140595650351018 | 3.19488023974814 | 0,00139889 | 0.0298864917892445 | Primary DEG |
| PSMD5 | 1273.41640507538 | -0.138570021299104 | 0.0433743319605735 | -3.19474710123632 | 0,00139953 | 0.0298864917892445 | Primary DEG |
| KBTBD6 | 187.207666107985 | -0.323948710835097 | 0.101401756975505 | -3.1947051066714 | 0,00139974 | 0.0298864917892445 | Primary DEG |
| ZNF576 | 149.878785477752 | -0.173748783773926 | 0.0543865772674573 | -3.19469973113918 | 0,00139976 | 0.0298864917892445 | Primary DEG |
| AQR | 3004.41382002614 | -0.0989663083205989 | 0.0309789412715795 | -3.19463171620367 | 0,00140009 | 0.0298864917892445 | Primary DEG |
| C17orf75 | 256.181845207085 | -0.159857295413552 | 0.0500408309805653 | -3.19453718655545 | 0,00140055 | 0.0298864917892445 | Primary DEG |
| RP11-640F22.1 | 2.29810378406365 | 0.86927594325114 | 0.272191397180734 | 3.19362019613701 | 0,00140501 | 0.0299493390367655 | Primary DEG |
| SPTY2D1 | 1516.38843151052 | -0.138547774739927 | 0.0433920614957555 | -3.1929290742151 | 0,00140838 | 0.0299789971045915 | Primary DEG |
| MYH10 | 86.35155019036 | 0.287420962941137 | 0.0900240217111523 | 3.19271409428192 | 0,00140942 | 0.0299789971045915 | Primary DEG |
| PGBD2 | 317.960943783382 | -0.170562469325603 | 0.0534440611089596 | -3.19142044572299 | 0,00141575 | 0.0300338342403835 | Primary DEG |
| CCDC92 | 209.572124660322 | 0.141104986349593 | 0.0442157700358595 | 3.19128189411051 | 0,00141643 | 0.0300338342403835 | Primary DEG |
| PANK4 | 502.432805100715 | -0.134866227256728 | 0.0422615838946523 | -3.19122509920396 | 0,00141671 | 0.0300338342403835 | Primary DEG |
| SACM1L | 2358.98810252929 | -0.166638381155904 | 0.0522243744713345 | -3.19081622025689 | 0,00141872 | 0.0300338342403835 | Primary DEG |
| TSC22D1 | 2509.80126988994 | 0.160880138128906 | 0.0504239396582744 | 3.19055074274637 | 0,00142002 | 0.0300338342403835 | Primary DEG |
| LINC00092 | 45.6584150100161 | 0.244981916233819 | 0.0767888376618235 | 3.19033239326677 | 0,00142109 | 0.0300338342403835 | Primary DEG |
| RP11-678G14.2 | 2.80781977648433 | 0.741491284577819 | 0.232454249553603 | 3.18983750996918 | 0,00142353 | 0.0300532701304436 | Primary DEG |
| ATP5MC2P3 | 2.89591007741542 | 0.730172434417585 | 0.228970803420116 | 3.18893249056677 | 0,00142799 | 0.0300672240168476 | Primary DEG |
| GLI3 | 17.0728308307718 | 0.476323011887103 | 0.149369458494296 | 3.18889160266517 | 0,00142819 | 0.0300672240168476 | Primary DEG |
| BNC2 | 65.1178820070258 | 0.398687027711205 | 0.125036536513745 | 3.18856422952325 | 0,00142981 | 0.0300672240168476 | Primary DEG |
| DISP1 | 236.244910363796 | 0.126016600492678 | 0.0395225357675217 | 3.18847457647781 | 0,00143026 | 0.0300672240168476 | Primary DEG |
| TFAP2E-AS1 | 107.548069313769 | 0.257986517499807 | 0.0809230064264099 | 3.18804909620376 | 0,00143236 | 0.0300796048990857 | Primary DEG |
| RNF2 | 1007.22525386389 | -0.117443781945567 | 0.0368481602670308 | -3.18723597309818 | 0,0014364 | 0.0301323821053269 | Primary DEG |
| GTSCR1 | 34.8551898775635 | 0.405576521940253 | 0.127266905057097 | 3.18681845652091 | 0,00143847 | 0.0301406733277986 | Primary DEG |
| SSU72 | 4404.046518006 | 0.105637105477567 | 0.0331509860903514 | 3.18654489461212 | 0,00143983 | 0.0301406733277986 | Primary DEG |
| ZBTB33 | 1909.64860268447 | -0.134499063560311 | 0.042228419734585 | -3.18503662712616 | 0,00144736 | 0.0302435464675371 | Primary DEG |
| PERP | 199.631833062777 | -0.251052663329774 | 0.0788247126914175 | -3.1849486634045 | 0,0014478 | 0.0302435464675371 | Primary DEG |
| ZNF395 | 1661.86761950756 | 0.130190407551289 | 0.0408875787059778 | 3.18410655929243 | 0,00145202 | 0.0302903589575416 | Primary DEG |
| PAAF1 | 342.351184644995 | -0.150706527381622 | 0.0473340624479032 | -3.18389167520732 | 0,00145309 | 0.0302903589575416 | Primary DEG |
| SEMA3F-AS1 | 44.3655755605302 | 0.508759910047928 | 0.159923658318195 | 3.1812673334152 | 0,00146632 | 0.0305227734318937 | Primary DEG |
| GIT1 | 2643.23773270854 | 0.131060669106832 | 0.0412038911836809 | 3.18078378866167 | 0,00146877 | 0.0305227734318937 | Primary DEG |
| SERBP1 | 7816.06074766695 | -0.0734153017636832 | 0.0230810132036097 | -3.18076598787274 | 0,00146886 | 0.0305227734318937 | Primary DEG |
| FANCF | 593.254810279484 | -0.163363860920743 | 0.0513726771603789 | -3.17997562032327 | 0,00147287 | 0.0305740971253466 | Primary DEG |
| C11orf54 | 299.227845619727 | -0.148900329666627 | 0.0468323789222377 | -3.17943126301286 | 0,00147564 | 0.0305995390471028 | Primary DEG |
| RP11-314N13.3 | 10.2700657578532 | 0.379239226786132 | 0.119298703008718 | 3.17890486000019 | 0,00147833 | 0.0306231303172502 | Primary DEG |
| ZNF182 | 395.711028433638 | -0.149357578090353 | 0.0470004920401263 | -3.17778754236956 | 0,00148404 | 0.0307092908556278 | Primary DEG |
| MED20 | 372.046929428203 | -0.187640260645543 | 0.0590671536860499 | -3.17672765548983 | 0,00148947 | 0.0307585230683617 | Primary DEG |
| TRAV34 | 21.0883583967663 | -0.393367973697777 | 0.123830648257333 | -3.17666086089057 | 0,00148981 | 0.0307585230683617 | Primary DEG |
| TNFSF14 | 430.877972138911 | -0.323579870396698 | 0.101869480098531 | -3.17641623461435 | 0,00149107 | 0.0307585230683617 | Primary DEG |
| CIAO1 | 3101.5501833295 | -0.0918854962333358 | 0.028940357207433 | -3.1749952350186 | 0,00149839 | 0.0308774176526128 | Primary DEG |
| DEAF1 | 700.30450039846 | 0.101113241651273 | 0.0318566377547781 | 3.17400858275155 | 0,00150349 | 0.0309390113368025 | Primary DEG |
| ID1 | 34.3091274434448 | 0.851290524838787 | 0.268228396682915 | 3.17375242653796 | 0,00150482 | 0.0309390113368025 | Primary DEG |
| NSMF | 1197.75119403431 | 0.160011608435223 | 0.0504233407806137 | 3.17336388184621 | 0,00150684 | 0.0309390113368025 | Primary DEG |
| AC003104.1 | 4.97337273972928 | 0.599605644647888 | 0.188958553379015 | 3.17321250573501 | 0,00150762 | 0.0309390113368025 | Primary DEG |
| CTD-2616J11.3 | 4.96084203260154 | 0.51599656568486 | 0.162636172456946 | 3.17270480416314 | 0,00151026 | 0.0309611102838881 | Primary DEG |
| HOMEZ | 505.988599673242 | -0.201312559741965 | 0.0634724478723924 | -3.17165268537762 | 0,00151574 | 0.0309851516629227 | Primary DEG |
| TRAPPC10 | 3156.10392871547 | 0.117485205682057 | 0.0370433113673503 | 3.17156326865551 | 0,00151621 | 0.0309851516629227 | Primary DEG |
| BANF1P2 | 3.06529170626788 | 0.671697436657713 | 0.211800138398341 | 3.17137392702938 | 0,0015172 | 0.0309851516629227 | Primary DEG |
| UBE2S | 196.867449853339 | -0.181165543783194 | 0.0571269362926176 | -3.17128058216218 | 0,00151769 | 0.0309851516629227 | Primary DEG |
| MESD | 1915.98991426754 | -0.0805095465126157 | 0.0253911169603471 | -3.17077608828104 | 0,00152032 | 0.0310070778858438 | Primary DEG |
| LRIF1 | 520.295547406976 | -0.206083174153031 | 0.0650017903972082 | -3.17042304363792 | 0,00152217 | 0.031012874844869 | Primary DEG |
| PEX12 | 163.596875453047 | -0.274335708132607 | 0.0865523973150187 | -3.16959109906715 | 0,00152654 | 0.0310698654592563 | Primary DEG |
| TYW3 | 675.8591854486 | -0.116033825648437 | 0.0366143132681426 | -3.16908376237103 | 0,0015292 | 0.0310922295064833 | Primary DEG |
| FUS | 6236.364645533 | 0.108171444713901 | 0.0341555070379429 | 3.16702792887059 | 0,00154006 | 0.0312463917103929 | Primary DEG |
| CTA-363E19.2 | 25.4040834901641 | 0.355180064124933 | 0.112152104317837 | 3.16694961976245 | 0,00154047 | 0.0312463917103929 | Primary DEG |
| TNFSF9 | 38.2130769990965 | 0.472135624624007 | 0.149091413295922 | 3.16675262636953 | 0,00154151 | 0.0312463917103929 | Primary DEG |
| CASP2 | 2020.95440943881 | -0.163860019424499 | 0.051783863242433 | -3.16430658441544 | 0,00155453 | 0.0314336690926012 | Primary DEG |
| ZCCHC4 | 280.935477229562 | -0.154962674657502 | 0.0489806563881285 | -3.16375251139061 | 0,00155749 | 0.0314336690926012 | Primary DEG |
| ST3GAL3 | 32.8554671991588 | 0.267423450745996 | 0.0845280412589953 | 3.16372468547575 | 0,00155764 | 0.0314336690926012 | Primary DEG |
| FURIN | 5792.32249336184 | 0.139702510269825 | 0.0441629103319366 | 3.16334474380868 | 0,00155968 | 0.0314336690926012 | Primary DEG |
| PARS2 | 113.663016442181 | -0.320482017015432 | 0.101321326380014 | -3.1630262696467 | 0,00156138 | 0.0314336690926012 | Primary DEG |
| RGP1 | 1418.30616327933 | 0.0976807903615696 | 0.0308838247856649 | 3.16284628084373 | 0,00156235 | 0.0314336690926012 | Primary DEG |
| ODR4 | 482.331691075381 | -0.13473424323773 | 0.0426063269644501 | -3.16230599624674 | 0,00156525 | 0.0314336690926012 | Primary DEG |
| RMI1 | 417.306638109292 | -0.258232768689977 | 0.0816642666891232 | -3.16212682926559 | 0,00156621 | 0.0314336690926012 | Primary DEG |
| CHMP4B | 5168.76435284784 | 0.0992704368801816 | 0.0313942455386765 | 3.16205837015208 | 0,00156658 | 0.0314336690926012 | Primary DEG |
| METTL14 | 841.683698046035 | -0.148506631422437 | 0.0469660717837193 | -3.16199813572479 | 0,00156691 | 0.0314336690926012 | Primary DEG |
| SRSF1 | 1254.030053657 | -0.151341516567386 | 0.0478662464651079 | -3.16175860327185 | 0,0015682 | 0.0314336690926012 | Primary DEG |
| VIRMA | 3142.42444545331 | -0.0734932469908076 | 0.0232467477090228 | -3.16144210410484 | 0,0015699 | 0.0314360628568868 | Primary DEG |
| HGD | 123.23822316429 | 0.336524615891921 | 0.106482005641392 | 3.16038953121584 | 0,00157558 | 0.0315180274458811 | Primary DEG |
| SPATS2 | 256.215150504067 | -0.164767202253692 | 0.0521688402705293 | -3.15834512324344 | 0,00158668 | 0.031707927122991 | Primary DEG |
| DENND4C | 2691.35071127117 | -0.145078572201892 | 0.0459622543562932 | -3.15647207113174 | 0,0015969 | 0.0318801421807976 | Primary DEG |
| LRRC58 | 1420.4406042101 | -0.140980434284715 | 0.0446796903311445 | -3.15535835722753 | 0,00160301 | 0.0319699055762242 | Primary DEG |
| FXR2 | 837.18424795031 | -0.127283253765697 | 0.0403462214998693 | -3.1547750702283 | 0,00160622 | 0.0319874528081158 | Primary DEG |
| CRIPT | 350.628195032069 | -0.141060155346053 | 0.0447155350318779 | -3.15461181992994 | 0,00160712 | 0.0319874528081158 | Primary DEG |
| MXD4 | 3012.54209333227 | 0.122067979280566 | 0.038705112479765 | 3.15379471754238 | 0,00161162 | 0.0320449705994853 | Primary DEG |
| SRP19 | 385.347661874583 | -0.101635114026031 | 0.0322340648248561 | -3.15303436219618 | 0,00161583 | 0.0320963640669496 | Primary DEG |
| TMEM263 | 723.888799033688 | -0.176929861132121 | 0.0561334819744607 | -3.15194879969532 | 0,00162185 | 0.0321594513467533 | Primary DEG |
| TERF1 | 908.011531456385 | -0.0858667920397476 | 0.0272430694143405 | -3.15187656478048 | 0,00162225 | 0.0321594513467533 | Primary DEG |
| POLR2B | 5224.90914655532 | -0.111593914525158 | 0.035411088589425 | -3.15138333698344 | 0,00162499 | 0.0321780183839372 | Primary DEG |
| AP5M1 | 1529.69148143029 | -0.143739953430507 | 0.0456154468930427 | -3.15112452515357 | 0,00162643 | 0.0321780183839372 | Primary DEG |
| CAAP1 | 449.323572659089 | -0.107548919795381 | 0.0341338290358366 | -3.15080150200751 | 0,00162823 | 0.0321815081738509 | Primary DEG |
| ZNF174 | 285.245734000083 | -0.133028108584148 | 0.0422306225639588 | -3.15003901215699 | 0,00163249 | 0.0322334808109554 | Primary DEG |
| RBM33 | 3364.57487974903 | 0.208825919263807 | 0.0663343777325976 | 3.14807987052522 | 0,00164347 | 0.0324057476726619 | Primary DEG |
| ENTPD7 | 394.999301614588 | -0.135633006808187 | 0.043093447787765 | -3.14741599410143 | 0,0016472 | 0.0324057476726619 | Primary DEG |
| PBX2 | 1163.67041005656 | 0.0991855252216067 | 0.031518844203033 | 3.14686428800147 | 0,00165032 | 0.0324057476726619 | Primary DEG |
| ARRDC3-AS1 | 24.4237232148808 | 0.241803918623455 | 0.0768397265774409 | 3.14686073719639 | 0,00165034 | 0.0324057476726619 | Primary DEG |
| PCDHGA10 | 14.2805203546316 | 0.46665322359542 | 0.148295523654928 | 3.14677889186518 | 0,0016508 | 0.0324057476726619 | Primary DEG |
| ACOX1 | 1570.1946180644 | -0.135719152670366 | 0.0431349533443683 | -3.14638459410982 | 0,00165303 | 0.0324057476726619 | Primary DEG |
| RPRD1A | 952.896933392679 | -0.0866194089043825 | 0.0275300983416813 | -3.14635305073496 | 0,0016532 | 0.0324057476726619 | Primary DEG |
| WASH8P | 158.43065733736 | 0.348463175495752 | 0.110758211186735 | 3.14616109958885 | 0,00165429 | 0.0324057476726619 | Primary DEG |
| TRAK1 | 3005.50944918664 | 0.151071954774985 | 0.0480297176736226 | 3.14538502602842 | 0,00165868 | 0.0324597699979158 | Primary DEG |
| SRPRA | 7433.9924337432 | -0.0709451121794803 | 0.0225644149683365 | -3.14411484982146 | 0,0016659 | 0.0325688464870535 | Primary DEG |
| RPF2 | 454.456774682184 | -0.136079132050113 | 0.043288385482334 | -3.1435483336667 | 0,00166913 | 0.032599805957148 | Primary DEG |
| NT5DC2 | 202.722444313724 | -0.294745934534918 | 0.0937953042942108 | -3.14243806502699 | 0,00167547 | 0.0326098802605614 | Primary DEG |
| C2orf42 | 521.633894951276 | -0.193133527733653 | 0.0614614482722714 | -3.14235237149115 | 0,00167596 | 0.0326098802605614 | Primary DEG |
| YKT6 | 2881.66261007909 | 0.0921503388987886 | 0.0293256720301458 | 3.14230953698388 | 0,00167621 | 0.0326098802605614 | Primary DEG |
| TUBGCP3 | 1285.95256823915 | -0.0940367066504707 | 0.0299260126481246 | -3.14230658645277 | 0,00167622 | 0.0326098802605614 | Primary DEG |
| CSNK1E | 596.752661290724 | 0.166906770368349 | 0.0531261319970714 | 3.14170755698061 | 0,00167966 | 0.0326446298764112 | Primary DEG |
| GCLM | 512.588520936137 | -0.185288262495544 | 0.0589830391021184 | -3.14138208739552 | 0,00168153 | 0.032648924710576 | Primary DEG |
| RADIL | 3.02624921839439 | 0.587008828778395 | 0.18694792080448 | 3.13995911937593 | 0,00168971 | 0.0327431755693336 | Primary DEG |
| RP11-129M16.4 | 32.0672554276088 | 0.310438043391582 | 0.0988688882185867 | 3.13989616941219 | 0,00169008 | 0.0327431755693336 | Primary DEG |
| CCDC121 | 58.6679034184042 | -0.358181923874226 | 0.114086325195039 | -3.13956929773913 | 0,00169196 | 0.0327431755693336 | Primary DEG |
| KNDC1 | 179.788887954059 | 0.435019562147162 | 0.138579425396521 | 3.13913527136102 | 0,00169447 | 0.0327431755693336 | Primary DEG |
| CCDC28A | 953.734523316785 | -0.140281507796025 | 0.0446883538638062 | -3.13910662772568 | 0,00169464 | 0.0327431755693336 | Primary DEG |
| POP1 | 221.126784130822 | -0.195024125223913 | 0.0621438351691566 | -3.13826986527391 | 0,00169948 | 0.0327526263765968 | Primary DEG |
| NUP107 | 980.618173737767 | -0.16605867989973 | 0.0529142268983322 | -3.13826147774568 | 0,00169953 | 0.0327526263765968 | Primary DEG |
| VPS4B | 3065.4706750461 | -0.147411203317794 | 0.0469839605489605 | -3.13747929283615 | 0,00170407 | 0.0327526263765968 | Primary DEG |
| DENND1C | 3490.67953196354 | 0.136734897087774 | 0.0435817208111244 | 3.13743685524395 | 0,00170432 | 0.0327526263765968 | Primary DEG |
| ZNF823 | 111.656305661031 | -0.163051568829612 | 0.0519713282339315 | -3.13733695809525 | 0,0017049 | 0.0327526263765968 | Primary DEG |
| USP1 | 2374.53368592446 | -0.088984443706553 | 0.0283632659836453 | -3.13731302163378 | 0,00170504 | 0.0327526263765968 | Primary DEG |
| SBNO1-AS1 | 15.3960515796248 | 0.335253976519361 | 0.10689069995148 | 3.13641857216334 | 0,00171025 | 0.0328208910418325 | Primary DEG |
| ORC2 | 679.988416800687 | -0.139982964381647 | 0.0446479589301623 | -3.13526010451243 | 0,00171702 | 0.0329094124511927 | Primary DEG |
| AP4E1 | 1036.5640922184 | -0.0922424588521261 | 0.0294228566254197 | -3.13506129015474 | 0,00171818 | 0.0329094124511927 | Primary DEG |
| TRAV25 | 119.122472465077 | -0.327581971479787 | 0.104504031215999 | -3.13463478554917 | 0,00172068 | 0.0329254799043147 | Primary DEG |
| LINC02773 | 17.1641725317511 | 0.335406027296949 | 0.107034162549074 | 3.13363527409457 | 0,00172655 | 0.0330059687959509 | Primary DEG |
| ORAI2 | 2893.01657694932 | 0.137033162135629 | 0.0437382312378866 | 3.1330293488624 | 0,00173012 | 0.0330423188262952 | Primary DEG |
| FAM117A | 3008.11020918016 | 0.118194458424835 | 0.0377437523155219 | 3.13149729885834 | 0,00173917 | 0.0331832363655865 | Primary DEG |
| TIGAR | 192.312549033832 | -0.208510106950394 | 0.066607147638001 | -3.13044642121011 | 0,00174541 | 0.033270168476895 | Primary DEG |
| HCCS | 729.21398094519 | -0.173009672558465 | 0.0552728786166451 | -3.13010063684947 | 0,00174746 | 0.0332773597949237 | Primary DEG |
| HSF2 | 458.038996715913 | -0.157550887668557 | 0.0503515008835099 | -3.12902068268146 | 0,0017539 | 0.0333678561901867 | Primary DEG |
| CTC-378H22.1 | 119.170826484178 | 0.139649808748015 | 0.0446473702466522 | 3.12783951163363 | 0,00176096 | 0.033452866043914 | Primary DEG |
| PHF23 | 1309.97137370364 | -0.154960769389228 | 0.0495444912721153 | -3.12770936607525 | 0,00176174 | 0.033452866043914 | Primary DEG |
| WDR3 | 1209.95948545859 | -0.132964348724714 | 0.0425274194775733 | -3.12655576938621 | 0,00176867 | 0.033552253773419 | Primary DEG |
| CCNA2 | 114.799111850392 | -0.33704023843051 | 0.107824555886457 | -3.1258207897042 | 0,0017731 | 0.0335883715050539 | Primary DEG |
| NARS2 | 307.191902655806 | -0.165424770324171 | 0.0529244590623402 | -3.125677111396 | 0,00177396 | 0.0335883715050539 | Primary DEG |
| TSR1 | 784.039927797512 | -0.0882202957829028 | 0.0282287540526774 | -3.12519268892548 | 0,00177689 | 0.0336077512514546 | Primary DEG |
| ANKRD44-IT1 | 5.23842959055125 | 0.581314468623265 | 0.186024295332617 | 3.12493842583227 | 0,00177842 | 0.0336077512514546 | Primary DEG |
| PITPNM2 | 991.293354735862 | 0.197878397786736 | 0.0633278592899643 | 3.12466582646817 | 0,00178007 | 0.0336077512514546 | Primary DEG |
| MMAB | 289.026515282298 | 0.122932687681872 | 0.0393499702006337 | 3.12408591556932 | 0,00178358 | 0.0336420201310695 | Primary DEG |
| GSKIP | 750.144395125921 | -0.140463306828133 | 0.044966663303757 | -3.12372091919031 | 0,0017858 | 0.0336517580803572 | Primary DEG |
| KPNA4 | 3542.2672309834 | -0.0690807045183347 | 0.0221208160086488 | -3.12288228839866 | 0,00179089 | 0.0337157401048233 | Primary DEG |
| RP11-667F14.1 | 64.7290731187492 | 0.181329595987377 | 0.0580703965702657 | 3.12258236032479 | 0,00179272 | 0.0337180921763903 | Primary DEG |
| ZNF773 | 192.757829571436 | -0.181149040622875 | 0.0580280806955316 | -3.12174792706568 | 0,00179781 | 0.0337674205873368 | Primary DEG |
| ACKR3 | 117.665788911816 | -0.225230170532274 | 0.0721522973044433 | -3.12159389162518 | 0,00179875 | 0.0337674205873368 | Primary DEG |
| ACSS1 | 2617.51158874081 | 0.0932086619061041 | 0.0298624257360722 | 3.12126893943224 | 0,00180074 | 0.0337727250987121 | Primary DEG |
| AMMECR1L | 975.68555333098 | -0.0871190594779161 | 0.0279176583172868 | -3.12057187919559 | 0,001805 | 0.0338207634114682 | Primary DEG |
| DNAJC19P5 | 9.53812408670737 | 0.396989056095957 | 0.1272467554722 | 3.11983637321651 | 0,00180952 | 0.0338682711896995 | Primary DEG |
| CTD-2600H12.2 | 15.372627193647 | 0.484764808694064 | 0.155393152925655 | 3.11960211609832 | 0,00181096 | 0.0338682711896995 | Primary DEG |
| ZNF419 | 256.769220448194 | -0.185590267287578 | 0.0594998676292478 | -3.11917109537146 | 0,00181361 | 0.0338858890909798 | Primary DEG |
| GPR18 | 490.90910648968 | -0.329113628379214 | 0.105559925306234 | -3.11778951552343 | 0,00182213 | 0.0339943363697998 | Primary DEG |
| IQCN | 152.887814371329 | 0.342407680230498 | 0.109827911641303 | 3.11767450653891 | 0,00182284 | 0.0339943363697998 | Primary DEG |
| CTD-2260A17.2 | 91.1186580155164 | 0.145181009298142 | 0.0465715540370906 | 3.11737523687779 | 0,00182469 | 0.0339968858281905 | Primary DEG |
| LIN52 | 393.827314257492 | -0.153221212983279 | 0.0491643904960705 | -3.1165079326169 | 0,00183007 | 0.0340650323620502 | Primary DEG |
| TRIM32 | 500.266208365486 | -0.166336344588429 | 0.0533839435777706 | -3.11584970012767 | 0,00183416 | 0.0340865957702079 | Primary DEG |
| CPOX | 410.437002648443 | -0.125384495328706 | 0.0402423073146755 | -3.11573822912933 | 0,00183485 | 0.0340865957702079 | Primary DEG |
| LMAN1 | 2626.24670700519 | -0.134171651889709 | 0.043068343269862 | -3.1153195526701 | 0,00183746 | 0.0340865957702079 | Primary DEG |
| GOPC | 1041.09451821099 | -0.117898308975307 | 0.0378459525322518 | -3.11521579156545 | 0,0018381 | 0.0340865957702079 | Primary DEG |
| RP11-578C11.2 | 7.66107297689823 | 0.470651706292981 | 0.15111223725798 | 3.11458366862428 | 0,00184205 | 0.0341084883681005 | Primary DEG |
| COA7 | 325.119977048362 | -0.133516909931001 | 0.0428697962540907 | -3.11447502898409 | 0,00184273 | 0.0341084883681005 | Primary DEG |
| CD40LG | 1152.05868254968 | -0.197305783777113 | 0.0633681000250419 | -3.1136452520928 | 0,00184792 | 0.0341278247578075 | Primary DEG |
| ZCCHC10 | 559.595258662335 | -0.12337550867053 | 0.0396243624872608 | -3.11362759994423 | 0,00184803 | 0.0341278247578075 | Primary DEG |
| ARRDC2 | 2540.97577441733 | 0.134296072089584 | 0.0431352757296182 | 3.1133699696597 | 0,00184964 | 0.0341278247578075 | Primary DEG |
| KCTD17 | 486.298660603152 | 0.104101808613218 | 0.0334401958822913 | 3.11307412730636 | 0,0018515 | 0.0341278247578075 | Primary DEG |
| OTUD6B-AS1 | 582.199854550712 | -0.144297136263592 | 0.046354070564134 | -3.11293343836864 | 0,00185238 | 0.0341278247578075 | Primary DEG |
| ST7-AS1 | 5.42230714223945 | 0.500353822567461 | 0.160752908836091 | 3.11256465708897 | 0,00185469 | 0.0341284787095521 | Primary DEG |
| PACC1 | 514.7356868942 | -0.100821921698272 | 0.032396928095387 | -3.11208276912612 | 0,00185772 | 0.0341284787095521 | Primary DEG |
| ATXN2L | 5353.06619657233 | 0.158960646392789 | 0.0510826330140237 | 3.11183345520089 | 0,00185929 | 0.0341284787095521 | Primary DEG |
| RNF6 | 2093.35352065254 | -0.104406366989061 | 0.0335530777262131 | -3.11167779722021 | 0,00186027 | 0.0341284787095521 | Primary DEG |
| TRIT1 | 494.664978856647 | -0.140722472184582 | 0.0452257087331433 | -3.1115592464216 | 0,00186102 | 0.0341284787095521 | Primary DEG |
| AARD | 3.34771869978899 | 0.658087946014825 | 0.211601823958775 | 3.11002964767938 | 0,00187069 | 0.0342610657501296 | Primary DEG |
| IMPA1 | 684.983266552425 | -0.128842645760264 | 0.0414379931675017 | -3.10928777944078 | 0,00187539 | 0.0342610657501296 | Primary DEG |
| SLFN5 | 9522.36136041963 | -0.176634513142399 | 0.0568104456138069 | -3.10919077000639 | 0,00187601 | 0.0342610657501296 | Primary DEG |
| BMP6 | 406.758911325879 | 0.30335974362054 | 0.0975724719634719 | 3.10907100656534 | 0,00187677 | 0.0342610657501296 | Primary DEG |
| SBDSP1 | 502.384188580297 | 0.206128452178349 | 0.0663025882812834 | 3.10890505969188 | 0,00187782 | 0.0342610657501296 | Primary DEG |
| LRRC32 | 32.7189500604995 | 0.352998653022916 | 0.113549026561482 | 3.10877744805485 | 0,00187863 | 0.0342610657501296 | Primary DEG |
| RBM15B | 3458.76652503789 | 0.0945466561447087 | 0.030415679768998 | 3.1084840734376 | 0,0018805 | 0.0342610657501296 | Primary DEG |
| SCOC-AS1 | 22.3480497324185 | 0.23867636369453 | 0.0767887704587701 | 3.10821962988301 | 0,00188218 | 0.0342610657501296 | Primary DEG |
| TIMM17A | 834.607022894189 | -0.121183476175143 | 0.0389912651048445 | -3.10796471592521 | 0,00188381 | 0.0342610657501296 | Primary DEG |
| IRF1-AS1 | 379.771612635714 | 0.197538571897224 | 0.0635648096468662 | 3.10767188629444 | 0,00188567 | 0.0342635934554262 | Primary DEG |
| YWHAG | 4226.40493266694 | -0.185519675598324 | 0.0597046986543575 | -3.10728769727714 | 0,00188813 | 0.0342767397420099 | Primary DEG |
| MRM1 | 32.5223953610623 | -0.284559548773095 | 0.091601331233176 | -3.1065001451642 | 0,00189316 | 0.0343075686065391 | Primary DEG |
| HCFC2 | 528.252055249263 | -0.175004775203662 | 0.0563399925852713 | -3.10622645075413 | 0,00189492 | 0.0343075686065391 | Primary DEG |
| RP11-631N16.2 | 42.000491607899 | 0.310884632217731 | 0.100084845870774 | 3.10621083055005 | 0,00189502 | 0.0343075686065391 | Primary DEG |
| CCDC69 | 12028.0327021495 | 0.111683187090825 | 0.0359638217701549 | 3.10543155854219 | 0,00190002 | 0.0343448897278607 | Primary DEG |
| GIGYF2 | 2253.07324512666 | -0.175398984413405 | 0.0564910788193688 | -3.10489705771501 | 0,00190345 | 0.0343448897278607 | Primary DEG |
| ZNF613 | 201.59114004437 | -0.197932089367482 | 0.0637523852996335 | -3.10470092118451 | 0,00190472 | 0.0343448897278607 | Primary DEG |
| LINC02100 | 2.28968339115889 | 0.581437573299985 | 0.187280604146853 | 3.10463315701428 | 0,00190515 | 0.0343448897278607 | Primary DEG |
| RP11-322M19.4 | 9.54980215236788 | -0.317767398781452 | 0.102355648848481 | -3.10454188270399 | 0,00190574 | 0.0343448897278607 | Primary DEG |
| UTP15 | 386.618300758256 | -0.127330357161065 | 0.0410356596433596 | -3.10291971099505 | 0,00191622 | 0.0344829539375695 | Primary DEG |
| EIF1 | 26109.9417751826 | 0.139024174475474 | 0.044805787928545 | 3.10281731228978 | 0,00191688 | 0.0344829539375695 | Primary DEG |
| PREPL | 1142.97311157454 | -0.175020999056819 | 0.0564315272188211 | -3.10147549929227 | 0,00192559 | 0.0345784909708853 | Primary DEG |
| GDAP2 | 776.269474524887 | -0.154174187640702 | 0.0497135136364914 | -3.10125308719947 | 0,00192704 | 0.0345784909708853 | Primary DEG |
| CTD-3065J16.9 | 9.71855321937514 | 0.283852754780925 | 0.0915353126494941 | 3.10101912108883 | 0,00192856 | 0.0345784909708853 | Primary DEG |
| AP1AR | 680.133488970955 | -0.101608323104781 | 0.0327682782309417 | -3.10081360969515 | 0,0019299 | 0.0345784909708853 | Primary DEG |
| ZBTB32 | 79.8404680728184 | -0.277476569942254 | 0.0894895752762612 | -3.1006580273251 | 0,00193091 | 0.0345784909708853 | Primary DEG |
| TMEM169 | 88.4737903497888 | -0.19523236664588 | 0.0629742706235002 | -3.10019258203246 | 0,00193395 | 0.0346016208212746 | Primary DEG |
| MT-RNR2 | 381943.391211482 | 0.26636123361587 | 0.0859265613958515 | 3.09987074181615 | 0,00193605 | 0.0346080049049944 | Primary DEG |
| TPBGL | 11.9993114692176 | 0.445695477363611 | 0.143791708967711 | 3.09959093304672 | 0,00193788 | 0.0346095038911143 | Primary DEG |
| ZC3H7A | 3093.82718187697 | 0.0886657394195934 | 0.0286161609591987 | 3.09844984259118 | 0,00194536 | 0.0347117876764692 | Primary DEG |
| GSTCD | 164.031238933503 | -0.166346573346193 | 0.0536963365567872 | -3.09791289337348 | 0,00194889 | 0.0347434679733973 | Primary DEG |
| CDKN2B | 118.973405720285 | -0.325942175480743 | 0.105227094796384 | -3.09751187288259 | 0,00195153 | 0.0347592517727323 | Primary DEG |
| CFAP20 | 493.660183664765 | -0.11195490080142 | 0.0361623237816867 | -3.09589896593193 | 0,00196217 | 0.0349175036993328 | Primary DEG |
| CNNM2 | 389.0364305321 | 0.129654975253885 | 0.0418869007232507 | 3.0953585253424 | 0,00196575 | 0.0349498231943661 | Primary DEG |
| LSM11 | 194.734338220698 | -0.153958554248833 | 0.0497550973029406 | -3.09432726684134 | 0,0019726 | 0.0350321099703263 | Primary DEG |
| PAK1IP1 | 425.018083426196 | -0.129288238329829 | 0.0417850134780572 | -3.09412939157535 | 0,00197391 | 0.0350321099703263 | Primary DEG |
| MED7 | 349.140406323877 | -0.123026870028049 | 0.0397752857476762 | -3.09304805020129 | 0,00198112 | 0.0350780296683115 | Primary DEG |
| NAA30 | 814.741647963302 | -0.125985051979999 | 0.0407333144678669 | -3.09292414884098 | 0,00198195 | 0.0350780296683115 | Primary DEG |
| CBR4 | 450.58594905742 | -0.178393000276164 | 0.0576807397642903 | -3.09276547085143 | 0,00198301 | 0.0350780296683115 | Primary DEG |
| OLFM4 | 150.801600389406 | 0.86313366845856 | 0.279089227284233 | 3.09267998932657 | 0,00198358 | 0.0350780296683115 | Primary DEG |
| ASB3 | 335.805893152385 | -0.100773842241593 | 0.0325929198586049 | -3.09189365907601 | 0,00198884 | 0.0351397330841188 | Primary DEG |
| LIX1-AS1 | 13.528627980678 | 0.505540446831173 | 0.163567307658966 | 3.09071815185229 | 0,00199673 | 0.0352477195477616 | Primary DEG |
| UTP3 | 1585.80779424031 | -0.105616820551948 | 0.0341815271759983 | -3.08988009833891 | 0,00200237 | 0.0353158761329391 | Primary DEG |
| SPOCD1 | 177.292612261698 | 0.38235132011123 | 0.123769979977005 | 3.08920887102242 | 0,0020069 | 0.0353558754357486 | Primary DEG |
| WHAMMP2 | 73.4292742378779 | 0.283053634535704 | 0.0916351924562994 | 3.08891842695362 | 0,00200887 | 0.0353558754357486 | Primary DEG |
| PRPF38A | 2459.01360795491 | -0.0860993189210321 | 0.0278751155894407 | -3.08875199619431 | 0,00200999 | 0.0353558754357486 | Primary DEG |
| POGLUT1 | 927.798683082676 | -0.108359880886974 | 0.0350854990592847 | -3.08845203267241 | 0,00201202 | 0.0353602114038294 | Primary DEG |
| RP5-906C1.1 | 78.6084275409657 | 0.227826443316087 | 0.0737807938216627 | 3.0878827878536 | 0,00201588 | 0.0353966324523229 | Primary DEG |
| KDSR | 1429.44704796115 | -0.101899386924448 | 0.0330083447812318 | -3.08707957335645 | 0,00202134 | 0.0354314035317583 | Primary DEG |
| HILPDA | 51.0146494041126 | -0.184909109709759 | 0.05990156631891 | -3.08688271564254 | 0,00202267 | 0.0354314035317583 | Primary DEG |
| KCNC3 | 267.89842891838 | 0.266780917936709 | 0.0864263082021324 | 3.08680219584025 | 0,00202322 | 0.0354314035317583 | Primary DEG |
| SLC9A3R2 | 68.1373946282751 | 0.176090075612506 | 0.057059743486767 | 3.0860649707152 | 0,00202825 | 0.0354880217111188 | Primary DEG |
| LAX1 | 958.071270219729 | -0.186729964443239 | 0.0605370022493316 | -3.08455915398918 | 0,00203854 | 0.0356235565369897 | Primary DEG |
| ARHGEF9 | 826.233202790219 | 0.107689902906558 | 0.0349159831634085 | 3.0842580717995 | 0,00204061 | 0.0356235565369897 | Primary DEG |
| HMMR | 53.3207100248002 | -0.342682135408616 | 0.111110910296039 | -3.08414479276238 | 0,00204138 | 0.0356235565369897 | Primary DEG |
| SIGLEC10-AS1 | 4.65029003855485 | 0.479880987007236 | 0.155624664527697 | 3.08357925437861 | 0,00204527 | 0.0356599455553513 | Primary DEG |
| RP11-677M14.3 | 15.3726960228065 | 0.304445085355736 | 0.0987457504591059 | 3.08312088307858 | 0,00204842 | 0.0356792496644643 | Primary DEG |
| METTL21EP | 16.5601274734892 | 0.418913630403422 | 0.135885903665473 | 3.08283360601343 | 0,0020504 | 0.0356792496644643 | Primary DEG |
| CEBPG | 1456.37657855367 | -0.110700434323068 | 0.0359149415613789 | -3.08229470828682 | 0,00205411 | 0.0356792496644643 | Primary DEG |
| ALKBH3-AS1 | 8.40537249601668 | -0.306818445016035 | 0.0995536424795902 | -3.08194092525481 | 0,00205656 | 0.0356792496644643 | Primary DEG |
| ELAVL1 | 2206.37466388819 | -0.0871928721152892 | 0.0282934378731935 | -3.08173480034746 | 0,00205798 | 0.0356792496644643 | Primary DEG |
| POLK | 869.241731058797 | -0.164073567388389 | 0.0532437952281547 | -3.08155282104362 | 0,00205924 | 0.0356792496644643 | Primary DEG |
| LTN1 | 1543.77785684944 | -0.168297629451994 | 0.0546186641412344 | -3.08132086527794 | 0,00206084 | 0.0356792496644643 | Primary DEG |
| ZNF473 | 319.353805904384 | -0.120150983694845 | 0.0389961159632663 | -3.08110130270475 | 0,00206237 | 0.0356792496644643 | Primary DEG |
| MED24 | 2028.80879530755 | 0.071425157043524 | 0.0231819218476288 | 3.08107142768363 | 0,00206257 | 0.0356792496644643 | Primary DEG |
| LYRM9 | 114.250604706005 | 0.187669693915306 | 0.0609317750593429 | 3.07999715636924 | 0,00207003 | 0.0357769744915161 | Primary DEG |
| RETREG2 | 5546.8072060441 | 0.102681071754174 | 0.0333444847021465 | 3.07940196621375 | 0,00207417 | 0.0358173091359599 | Primary DEG |
| PCDHGB6 | 21.3692034240209 | 0.468951883925403 | 0.152364922716699 | 3.07782050857829 | 0,00208521 | 0.0359692095462952 | Primary DEG |
| RP11-47I22.1 | 100.311262719773 | 0.392987600026853 | 0.127691949839223 | 3.07762236007566 | 0,00208659 | 0.0359692095462952 | Primary DEG |
| RP11-80H8.4 | 7.42055346860631 | 0.474847910345515 | 0.15433004701382 | 3.07683383458694 | 0,00209212 | 0.0360057470472007 | Primary DEG |
| BRIP1 | 96.3763208200501 | -0.174248851175082 | 0.0566331086146599 | -3.07680181147563 | 0,00209234 | 0.0360057470472007 | Primary DEG |
| FMNL3 | 1960.02365475244 | 0.137225620006734 | 0.0446122279824099 | 3.07596428631273 | 0,00209823 | 0.0360757213112633 | Primary DEG |
| DMAP1 | 961.804910807815 | 0.135628194055466 | 0.0441022760941106 | 3.07531052968892 | 0,00210284 | 0.0360956403716559 | Primary DEG |
| DNAJA1 | 5042.01470168898 | -0.162024799375001 | 0.0526861449717928 | -3.07528287487624 | 0,00210303 | 0.0360956403716559 | Primary DEG |
| ACTR1B | 2281.88854329022 | 0.0900631438512362 | 0.0292918455594782 | 3.07468314580451 | 0,00210726 | 0.0361212679309129 | Primary DEG |
| IGKV2D-40 | 6.66712620731038 | -0.554414807506212 | 0.180323583672082 | -3.07455517584662 | 0,00210817 | 0.0361212679309129 | Primary DEG |
| H3-3B | 22423.6820415504 | 0.15674310123894 | 0.0509968365950858 | 3.07358478886598 | 0,00211504 | 0.0361963634962214 | Primary DEG |
| MED21 | 449.199939595128 | -0.183287977933802 | 0.0596373023034104 | -3.07337808476491 | 0,0021165 | 0.0361963634962214 | Primary DEG |
| TMCC2 | 313.016689758831 | 0.427361400908432 | 0.139071089971166 | 3.07297081655891 | 0,00211939 | 0.0361963634962214 | Primary DEG |
| CCR4 | 1128.48919225576 | -0.287191641825428 | 0.0934606997357078 | -3.07285995758176 | 0,00212018 | 0.0361963634962214 | Primary DEG |
| GPATCH2 | 298.542813824673 | -0.167004266273681 | 0.0543518870429232 | -3.07264890622459 | 0,00212168 | 0.0361963634962214 | Primary DEG |
| ETFDH | 556.338361579549 | -0.11736104307796 | 0.038202331962936 | -3.0720910752732 | 0,00212565 | 0.0362124151475924 | Primary DEG |
| IGLV3-21 | 836.182151769949 | -0.50655672394516 | 0.16491444719735 | -3.07163339873415 | 0,00212891 | 0.0362124151475924 | Primary DEG |
| VTA1 | 1699.73343180416 | -0.162273455176923 | 0.0528337449125824 | -3.07139793791671 | 0,00213059 | 0.0362124151475924 | Primary DEG |
| UBR2 | 2949.13193026249 | -0.173056515136363 | 0.0563464765051497 | -3.07129257888108 | 0,00213134 | 0.0362124151475924 | Primary DEG |
| RP11-345P4.9 | 122.155267258747 | 0.216867212349876 | 0.0706202053216651 | 3.07089467330316 | 0,00213418 | 0.0362124151475924 | Primary DEG |
| PLA2G4A | 237.118457170813 | -0.35029147540824 | 0.114070393354348 | -3.07083604349549 | 0,0021346 | 0.0362124151475924 | Primary DEG |
| IGKV4-1 | 2057.10785332804 | -0.518964290699238 | 0.169015052999737 | -3.07052112512159 | 0,00213686 | 0.0362124151475924 | Primary DEG |
| LCMT2 | 239.663462881136 | -0.246385789757595 | 0.0802498354242686 | -3.07023420615122 | 0,00213891 | 0.0362124151475924 | Primary DEG |
| PTCD1 | 100.890409120617 | 0.210769276804031 | 0.0686497215799543 | 3.07021313347285 | 0,00213906 | 0.0362124151475924 | Primary DEG |
| SEC23A | 2126.663176797 | -0.0927497894229528 | 0.0302204757865869 | -3.06910420861471 | 0,00214702 | 0.0363160939451916 | Primary DEG |
| MAPKAPK2 | 6039.7763446341 | 0.1054175176344 | 0.0343561645160421 | 3.06837271038147 | 0,00215228 | 0.0363740821645429 | Primary DEG |
| FNDC3A | 1837.90358452806 | -0.137934653114888 | 0.0449578214180355 | -3.06809024023466 | 0,00215432 | 0.0363774709129735 | Primary DEG |
| IGKV1D-16 | 41.3269377664225 | -0.522345538832683 | 0.170304587063284 | -3.06712548287724 | 0,00216128 | 0.0364640305699368 | Primary DEG |
| SH3PXD2A | 573.707508569105 | 0.214339663356515 | 0.0699070326595072 | 3.06606724963552 | 0,00216895 | 0.036543421187647 | Primary DEG |
| ZNF12 | 992.91409014762 | -0.149098401508513 | 0.0486301409325945 | -3.06596688081114 | 0,00216967 | 0.036543421187647 | Primary DEG |
| GTF2A1 | 1533.90434892248 | -0.12262097418093 | 0.0400129948204933 | -3.0645287794886 | 0,00218013 | 0.0366295741282701 | Primary DEG |
| ATAD2B | 1037.04202040659 | -0.172558928499446 | 0.056309383665152 | -3.06447908443787 | 0,00218049 | 0.0366295741282701 | Primary DEG |
| RN7SL834P | 13.5527817743865 | 0.573096774339238 | 0.187012818173598 | 3.0644785739085 | 0,0021805 | 0.0366295741282701 | Primary DEG |
| NFYA | 1817.48757413877 | -0.0643555792721886 | 0.0210032831126932 | -3.06407236082514 | 0,00218346 | 0.0366295741282701 | Primary DEG |
| YIPF5 | 1420.02717303968 | -0.126338370416112 | 0.0412332199753053 | -3.06399477149194 | 0,00218403 | 0.0366295741282701 | Primary DEG |
| TIMM10B | 1007.07293162952 | -0.133657565246922 | 0.0436264661179869 | -3.06368076858318 | 0,00218632 | 0.0366370431933614 | Primary DEG |
| GLMN | 155.323978220345 | -0.149331022897589 | 0.048758941632296 | -3.06263872632293 | 0,00219395 | 0.0367337977029593 | Primary DEG |
| TRBJ2-2P | 2.82489688208169 | 0.612359356642231 | 0.199991200570666 | 3.06193149946044 | 0,00219914 | 0.0367646987463485 | Primary DEG |
| DDX19B | 602.97279145229 | 0.071951188680136 | 0.0234990085765643 | 3.06188188517423 | 0,0021995 | 0.0367646987463485 | Primary DEG |
| ANKS6 | 317.07379243793 | 0.183362660883077 | 0.0599088967623622 | 3.0606916633837 | 0,00220826 | 0.0368800509245806 | Primary DEG |
| EIF2AK4 | 2263.14643988364 | 0.0928020716653362 | 0.0303526249211078 | 3.05746445016028 | 0,00223218 | 0.0372481140359887 | Primary DEG |
| DPM1 | 934.135027657569 | -0.0956008220166534 | 0.0312798808427226 | -3.05630390656988 | 0,00224084 | 0.0373458357338529 | Primary DEG |
| SMNDC1 | 1210.33542981643 | -0.0801738787318494 | 0.0262355407137535 | -3.05592629504371 | 0,00224366 | 0.0373458357338529 | Primary DEG |
| LYNX1 | 206.739413774009 | 0.320166316854253 | 0.104769106111155 | 3.0559229599093 | 0,00224369 | 0.0373458357338529 | Primary DEG |
| GEMIN2 | 199.169103339064 | -0.123306699018184 | 0.0403746081712762 | -3.05406552789553 | 0,00225763 | 0.0375463171407209 | Primary DEG |
| MTF2 | 1277.06943323279 | -0.0726169498064518 | 0.0237824817847142 | -3.05337981392359 | 0,00226279 | 0.0376006845612441 | Primary DEG |
| CHASERR | 1456.94653351741 | 0.208658319754632 | 0.0683553615134121 | 3.05255235485355 | 0,00226904 | 0.0376539608208153 | Primary DEG |
| RP11-797A18.5 | 6.25040615804904 | 0.648449945344043 | 0.212435737461027 | 3.05245225259242 | 0,0022698 | 0.0376539608208153 | Primary DEG |
| ZNF250 | 177.345994389336 | -0.166692430996983 | 0.0546189560966857 | -3.05191535887114 | 0,00227386 | 0.0376898345595098 | Primary DEG |
| LAMTOR3 | 1164.9249369872 | -0.152879855238711 | 0.0501008950958724 | -3.0514395989565 | 0,00227747 | 0.0377180773388925 | Primary DEG |
| RP11-1099M24.6 | 16.7402340419864 | 0.330083057771315 | 0.108185184499257 | 3.05109298744674 | 0,0022801 | 0.0377301270409609 | Primary DEG |
| ADO | 625.975320498375 | -0.176823981557727 | 0.0579711588422097 | -3.05020608677186 | 0,00228684 | 0.0378058016801982 | Primary DEG |
| CCDC174 | 742.59247012758 | -0.114180258858655 | 0.0374362682357142 | -3.04999040341651 | 0,00228849 | 0.0378058016801982 | Primary DEG |
| RP11-521B24.4 | 28.697952454278 | 0.181092717222162 | 0.0593858847896438 | 3.0494235770609 | 0,00229281 | 0.0378360313902579 | Primary DEG |
| ABHD3 | 874.736663728035 | -0.121100508295658 | 0.0397178856788209 | -3.04901699136098 | 0,00229592 | 0.0378360313902579 | Primary DEG |
| SEC22C | 1263.22763037055 | -0.0909696109419297 | 0.0298358811903451 | -3.04900030810444 | 0,00229604 | 0.0378360313902579 | Primary DEG |
| TOGARAM1 | 442.248705231193 | -0.131670481283412 | 0.0431885263177746 | -3.0487375353954 | 0,00229805 | 0.0378376865239677 | Primary DEG |
| ADPRS | 702.549585964137 | -0.140539457890132 | 0.0461078091470399 | -3.04806193332556 | 0,00230323 | 0.0378913884482028 | Primary DEG |
| EXOC5 | 2061.72692180333 | -0.124726729281058 | 0.0409239646003187 | -3.0477675000258 | 0,00230548 | 0.0378970818677207 | Primary DEG |
| VMAC | 242.379104373499 | 0.150346939127932 | 0.0493477909399746 | 3.04668023155909 | 0,00231384 | 0.0380029063362236 | Primary DEG |
| NFKBIZ | 1743.8130704638 | 0.3647949167617 | 0.119752308811183 | 3.04624537416546 | 0,00231719 | 0.0380264114083902 | Primary DEG |
| CROCC2 | 30.7708671172977 | 0.565220462697149 | 0.185564353510386 | 3.04595388071402 | 0,00231943 | 0.0380268997846547 | Primary DEG |
| EVC2 | 34.761285650385 | -0.257289872813308 | 0.0844773486375995 | -3.04566699787252 | 0,00232165 | 0.0380268997846547 | Primary DEG |
| ARL14EP | 1279.32349212496 | -0.113701845009524 | 0.0373344295466566 | -3.04549570972905 | 0,00232297 | 0.0380268997846547 | Primary DEG |
| BCDIN3D | 165.452142471963 | -0.174841735437082 | 0.0574253187800045 | -3.04468027607992 | 0,00232928 | 0.0380986832939534 | Primary DEG |
| GNA12 | 2505.12665873094 | 0.119459070802715 | 0.0392398519524038 | 3.04433031367227 | 0,00233199 | 0.0381021739811521 | Primary DEG |
| CLIC5 | 151.533266347956 | -0.287776112957405 | 0.094533933944113 | -3.04415674827977 | 0,00233334 | 0.0381021739811521 | Primary DEG |
| HNRNPCP2 | 17.4645879734067 | -0.251083381559125 | 0.0824918283913839 | -3.04373640947629 | 0,0023366 | 0.0381240437137975 | Primary DEG |
| TTC32 | 216.206435529085 | 0.200331214868087 | 0.0658369055560891 | 3.04284068602551 | 0,00234356 | 0.0382062554809481 | Primary DEG |
| CHM | 992.565632787601 | -0.111540337467331 | 0.0366749625616705 | -3.04132109964042 | 0,00235543 | 0.0383570957762809 | Primary DEG |
| LINC02288 | 4.8911086984823 | 0.459991409329037 | 0.151255250052876 | 3.04115995423785 | 0,00235669 | 0.0383570957762809 | Primary DEG |
| KNSTRN | 266.051487777573 | -0.154785194447515 | 0.0509160504529028 | -3.04000787709744 | 0,00236572 | 0.0384725394409051 | Primary DEG |
| PFKP | 1456.49812406828 | 0.144251065152065 | 0.0474599573236137 | 3.03942677757725 | 0,00237029 | 0.038515239335752 | Primary DEG |
| RP11-301N24.6 | 14.694859972637 | 0.436565091399786 | 0.143657567564334 | 3.03892860502653 | 0,00237421 | 0.03854283209183 | Primary DEG |
| RP11-613D13.4 | 3.73263370367304 | -0.457786334698909 | 0.150651162386624 | -3.038717574074 | 0,00237588 | 0.03854283209183 | Primary DEG |
| PLK1 | 90.0857482193015 | -0.329851027401218 | 0.108586312367504 | -3.0376851392177 | 0,00238403 | 0.0386434934461976 | Primary DEG |
| FBXO41 | 813.991324723971 | 0.196366281302616 | 0.064658404043255 | 3.03698002151819 | 0,00238961 | 0.0387023584780306 | Primary DEG |
| RP11-517H2.8 | 12.9742621422819 | 0.334830725167792 | 0.110265299817788 | 3.03659198062396 | 0,00239269 | 0.0387205757887352 | Primary DEG |
| CENPBD1 | 390.591238356855 | -0.196611279769287 | 0.0647738899319033 | -3.03534773001875 | 0,00240259 | 0.0388489775896918 | Primary DEG |
| GPR15 | 918.346586998339 | -0.512055844612814 | 0.168754727363255 | -3.03432000165887 | 0,00241079 | 0.0389106561068627 | Primary DEG |
| MRPS14 | 699.163307969599 | -0.116606663932611 | 0.0384307283821746 | -3.03420384784319 | 0,00241171 | 0.0389106561068627 | Primary DEG |
| AKAP8L | 1267.66755401473 | 0.158098011235861 | 0.0521155554042498 | 3.03360503422686 | 0,00241651 | 0.0389106561068627 | Primary DEG |
| TBC1D13 | 1388.5909971154 | 0.0692522542101108 | 0.0228285722786244 | 3.03357798135083 | 0,00241672 | 0.0389106561068627 | Primary DEG |
| NCOA5 | 1857.48964977132 | -0.12169411311384 | 0.0401181009169444 | -3.03339665468663 | 0,00241818 | 0.0389106561068627 | Primary DEG |
| DFFBP1 | 3.99690891069906 | 0.565553101828511 | 0.186449689961191 | 3.03327456294634 | 0,00241915 | 0.0389106561068627 | Primary DEG |
| GART | 1647.41475385146 | -0.147522662020451 | 0.0486367571719713 | -3.0331516860558 | 0,00242014 | 0.0389106561068627 | Primary DEG |
| MTA2 | 5126.90229818344 | -0.0700146460916649 | 0.0230873682715798 | -3.03259536851811 | 0,00242461 | 0.0389508618906566 | Primary DEG |
| RP11-345F18.2 | 40.4097293719132 | 0.23705109979719 | 0.0781858118552974 | 3.03189407607499 | 0,00243025 | 0.0389987616278184 | Primary DEG |
| RFK | 696.26681114759 | -0.138691247674823 | 0.0457518999844774 | -3.03137678920171 | 0,00243441 | 0.0389987616278184 | Primary DEG |
| MLC1 | 955.309937545497 | 0.197174626250649 | 0.0650460497387862 | 3.03130823535739 | 0,00243497 | 0.0389987616278184 | Primary DEG |
| TARS1 | 2307.02424223556 | -0.106831007307065 | 0.0352459863552547 | -3.03101199184168 | 0,00243736 | 0.0389987616278184 | Primary DEG |
| MRPL17 | 500.516169949323 | -0.149148394961421 | 0.0492081638528552 | -3.03096850773405 | 0,00243771 | 0.0389987616278184 | Primary DEG |
| SCYL2 | 2548.3645677247 | -0.0961954846462413 | 0.0317420154336356 | -3.03054117175897 | 0,00244116 | 0.0389987616278184 | Primary DEG |
| PHF6 | 971.883967046197 | -0.110002432736651 | 0.0362982437415221 | -3.03051666962106 | 0,00244136 | 0.0389987616278184 | Primary DEG |
| ADAMTS14 | 4.53455257904898 | 0.604575930270088 | 0.199587338392219 | 3.02912967896795 | 0,00245259 | 0.0391245950750891 | Primary DEG |
| IGHA1 | 4999.11567307456 | -0.500521894501294 | 0.165240154509056 | -3.02905728930351 | 0,00245318 | 0.0391245950750891 | Primary DEG |
| DIP2A-IT1 | 2.99979252205667 | 0.73617727194608 | 0.24307802426528 | 3.02856366457325 | 0,00245719 | 0.0391346759261726 | Primary DEG |
| EXD2 | 390.915691281156 | -0.131101101963212 | 0.0432892104870389 | -3.02849371675338 | 0,00245776 | 0.0391346759261726 | Primary DEG |
| FZD7 | 19.099813344767 | -0.352438756413678 | 0.116392371902803 | -3.02802280469026 | 0,0024616 | 0.0391642546929407 | Primary DEG |
| CAPS2 | 47.7039821559457 | 0.185744667950487 | 0.0613861110181689 | 3.0258419187935 | 0,00247942 | 0.0393957858306847 | Primary DEG |
| PRR29 | 159.618457129565 | 0.153269768066172 | 0.0506550294124011 | 3.02575617552892 | 0,00248012 | 0.0393957858306847 | Primary DEG |
| DDX23 | 3952.14847101098 | -0.0796734072167086 | 0.0263382377124693 | -3.0250090414739 | 0,00248626 | 0.0394394461879562 | Primary DEG |
| HNRNPUL1 | 11686.6997792176 | 0.0926155866890088 | 0.0306173610637364 | 3.0249369465974 | 0,00248685 | 0.0394394461879562 | Primary DEG |
| SMARCD1 | 2647.02170925652 | 0.0740116805944167 | 0.0244746628145532 | 3.02401226751151 | 0,00249446 | 0.0395285684836372 | Primary DEG |
| LINC01145 | 237.800353016716 | 0.183096737426494 | 0.0605543495010006 | 3.02367606844606 | 0,00249724 | 0.0395409107254557 | Primary DEG |
| MFSD14B | 3529.45162966825 | -0.143636990998747 | 0.0475133922266013 | -3.023084319337 | 0,00250213 | 0.0395866952010648 | Primary DEG |
| CNIH3 | 19.8938174661958 | 0.235562139350492 | 0.0779307800105499 | 3.02270988842409 | 0,00250522 | 0.0396040973321935 | Primary DEG |
| ZCWPW1 | 183.199739227109 | 0.122742071369421 | 0.0406173970240452 | 3.02190884602375 | 0,00251186 | 0.0396774084005058 | Primary DEG |
| HES6 | 47.7938485863088 | 0.246791869757819 | 0.0816959201425129 | 3.02085917298327 | 0,00252059 | 0.03971466820739 | Primary DEG |
| PET117 | 22.1453290812749 | 0.221555917481508 | 0.0733467236218398 | 3.02066549862272 | 0,0025222 | 0.03971466820739 | Primary DEG |
| BAG4 | 639.419322531132 | -0.16936536411914 | 0.0560712108229648 | -3.02054051684173 | 0,00252324 | 0.03971466820739 | Primary DEG |
| AP003068.23 | 395.091823532827 | 0.354183732275917 | 0.117281034855661 | 3.01995742714766 | 0,0025281 | 0.03971466820739 | Primary DEG |
| ZNF547 | 35.3560630279438 | -0.349151598780736 | 0.115620278000901 | -3.01981282883628 | 0,00252931 | 0.03971466820739 | Primary DEG |
| RP11-473M20.16 | 10.1363489177332 | 0.31185811667174 | 0.103273033009069 | 3.01974395043044 | 0,00252989 | 0.03971466820739 | Primary DEG |
| PAFAH2 | 952.110678610029 | -0.156609664600796 | 0.0518619986033914 | -3.01973832127933 | 0,00252993 | 0.03971466820739 | Primary DEG |
| ALDH1B1 | 299.952413094588 | -0.14524893835304 | 0.0481030985969803 | -3.01953393002751 | 0,00253164 | 0.03971466820739 | Primary DEG |
| PPP2R3B | 98.4433070849846 | 0.190667815655545 | 0.063149671091864 | 3.0193002173864 | 0,00253359 | 0.03971466820739 | Primary DEG |
| RP1-152L7.5 | 16.018070949792 | -0.421676798353469 | 0.139664095347213 | -3.0192212057448 | 0,00253425 | 0.03971466820739 | Primary DEG |
| C3 | 337.9945404818 | 0.308906937572342 | 0.102353070928871 | 3.0180524606537 | 0,00254405 | 0.0398366662510628 | Primary DEG |
| MAP3K8 | 939.152033741775 | 0.202116658366857 | 0.0670181147704242 | 3.01585114799519 | 0,00256259 | 0.0400402932359446 | Primary DEG |
| FADD | 72.9866853130902 | -0.266808665397845 | 0.0884720888263967 | -3.01573828466272 | 0,00256354 | 0.0400402932359446 | Primary DEG |
| PARP3 | 370.291444510446 | 0.146250789564684 | 0.0485017961669244 | 3.01536852493763 | 0,00256667 | 0.0400402932359446 | Primary DEG |
| SUCLA2 | 794.338762424468 | -0.132048951969499 | 0.0437926366982886 | -3.0153231667519 | 0,00256706 | 0.0400402932359446 | Primary DEG |
| KB-318B8.7 | 6.89995555411052 | 0.540538830747893 | 0.17926465233977 | 3.01531185145959 | 0,00256715 | 0.0400402932359446 | Primary DEG |
| AC103563.9 | 15.9499732023319 | 0.422362852244326 | 0.140103300727073 | 3.01465311703902 | 0,00257273 | 0.0400958032124023 | Primary DEG |
| GIMAP8 | 5269.45206244462 | -0.253225478126587 | 0.084041994571242 | -3.01308267870688 | 0,00258609 | 0.0402722226424808 | Primary DEG |
| OBI1 | 377.447062126172 | -0.141941204937822 | 0.047124866915767 | -3.01202346505367 | 0,00259513 | 0.0403407985928273 | Primary DEG |
| MAT2A | 3107.32268430628 | -0.134848187895813 | 0.0447703612603222 | -3.01199686801104 | 0,00259535 | 0.0403407985928273 | Primary DEG |
| POLR1G | 35.6259237126151 | -0.203087005868406 | 0.0674343511565322 | -3.0116254162065 | 0,00259853 | 0.0403407985928273 | Primary DEG |
| DNAH17 | 43.1789071979397 | 0.324084622045371 | 0.107611606326954 | 3.01161401736455 | 0,00259863 | 0.0403407985928273 | Primary DEG |
| HSPA8 | 55291.1498115546 | -0.132664034024239 | 0.0440631146233935 | -3.0107729596084 | 0,00260584 | 0.0404210454906803 | Primary DEG |
| TNPO3 | 2643.6403780607 | -0.0762579738199713 | 0.0253365604960525 | -3.00979976472547 | 0,0026142 | 0.0405190733999881 | Primary DEG |
| SNORD14A | 13.8298279442338 | 0.455158970875205 | 0.15129246440474 | 3.00847086248497 | 0,00262566 | 0.0406524246080724 | Primary DEG |
| ZBTB5 | 574.200645986929 | -0.143662884981841 | 0.0477555295858387 | -3.00829843638551 | 0,00262715 | 0.0406524246080724 | Primary DEG |
| RAB3GAP2 | 1997.85029950026 | -0.11527885018844 | 0.038322942926784 | -3.00808970774193 | 0,00262896 | 0.0406524246080724 | Primary DEG |
| SPIN4 | 178.139478527911 | -0.151754914733795 | 0.0504622848274506 | -3.00729376905349 | 0,00263585 | 0.0407272576408539 | Primary DEG |
| GNAQ | 2810.26391085113 | -0.128530643623672 | 0.0427473309960012 | -3.00675248322978 | 0,00264055 | 0.0407369596142906 | Primary DEG |
| CROT | 288.155999242585 | -0.150945069732017 | 0.050202104251777 | -3.00674786409324 | 0,00264059 | 0.0407369596142906 | Primary DEG |
| ZNF561 | 1382.01065561273 | -0.129091878554439 | 0.0429391526919954 | -3.0063909150798 | 0,00264369 | 0.0407531016168665 | Primary DEG |
| POLR1F | 591.271074223838 | -0.149208009420061 | 0.0496374119831393 | -3.00595868033458 | 0,00264745 | 0.0407793651563758 | Primary DEG |
| ZNF350-AS1 | 16.594954605212 | 0.44524642023067 | 0.148142956786541 | 3.00551865501256 | 0,00265128 | 0.0407881707352053 | Primary DEG |
| PLEKHM2 | 2709.245050079 | 0.112866705806198 | 0.0375552939475051 | 3.00534742089792 | 0,00265278 | 0.0407881707352053 | Primary DEG |
| DGCR11 | 66.1839515676497 | 0.224249347783197 | 0.0746208138831797 | 3.0051849626602 | 0,00265419 | 0.0407881707352053 | Primary DEG |
| CDC6 | 61.8023153149236 | -0.333563547153205 | 0.111015518139982 | -3.00465694113689 | 0,00265881 | 0.0408273898040719 | Primary DEG |
| ZNF525 | 333.979295788268 | -0.181622475297579 | 0.0604699963847473 | -3.0035139103033 | 0,00266881 | 0.0409493486821428 | Primary DEG |
| ACTG2 | 2.73145040308566 | 0.645166575053497 | 0.214872645083938 | 3.00255332548947 | 0,00267725 | 0.0410470353911549 | Primary DEG |
| JRKL | 407.651895826718 | -0.153385136107631 | 0.0510909580463775 | -3.00219729621036 | 0,00268038 | 0.0410633166010208 | Primary DEG |
| RCAN3AS | 26.9187924985224 | 0.302926946259528 | 0.100912369517659 | 3.00188121344747 | 0,00268317 | 0.0410742339518604 | Primary DEG |
| LRRN1 | 37.2963785236084 | -0.439071194473772 | 0.146313585128486 | -3.00089150360302 | 0,00269191 | 0.0411761704280002 | Primary DEG |
| CARD17 | 31.2088206185454 | -0.332803936511847 | 0.110915750264511 | -3.00051107005252 | 0,00269527 | 0.0411871868448105 | Primary DEG |
| CPSF2 | 3290.41605043368 | -0.136613503963585 | 0.0455358881547926 | -3.00012823949293 | 0,00269866 | 0.0411871868448105 | Primary DEG |
| CISD1 | 249.275182102359 | -0.146452132224502 | 0.048815654558535 | -3.00010587892232 | 0,00269886 | 0.0411871868448105 | Primary DEG |
| ZNF430 | 850.43333562766 | -0.140655189581781 | 0.0469015649118137 | -2.9989444882329 | 0,00270917 | 0.0413127070736729 | Primary DEG |
| LIPT1 | 193.190704324365 | -0.138378323190102 | 0.0461540697573831 | -2.99818247702774 | 0,00271595 | 0.0413843146351127 | Primary DEG |
| CCT6B | 41.3125634017131 | 0.180473143220402 | 0.0602116228999374 | 2.99731404882278 | 0,0027237 | 0.0414670377323316 | Primary DEG |
| SUPV3L1 | 546.094092727794 | -0.120946894174692 | 0.0403545649332869 | -2.99710564033186 | 0,00272556 | 0.0414670377323316 | Primary DEG |
| SCRN2 | 459.605393162014 | 0.181191099381274 | 0.0604661101225947 | 2.9965727746321 | 0,00273033 | 0.0414999011159718 | Primary DEG |
| CAMTA2-AS1 | 3.36858901040592 | 0.565830377383553 | 0.188836948726726 | 2.9963965272622 | 0,00273191 | 0.0414999011159718 | Primary DEG |
| ERMN | 46.1659732083712 | 0.371101459652636 | 0.123870969433618 | 2.99587111774004 | 0,00273662 | 0.0415396394086195 | Primary DEG |
| SF3B2 | 9820.13665850942 | 0.0727390478757379 | 0.0242863247870162 | 2.99506197473836 | 0,00274389 | 0.0415999820296597 | Primary DEG |
| FAM126A | 1191.96116112827 | -0.131533008001025 | 0.0439180934454819 | -2.99496170443519 | 0,00274479 | 0.0415999820296597 | Primary DEG |
| TTC17 | 3245.43859361586 | 0.0886302733261663 | 0.0295967385224413 | 2.99459595046136 | 0,00274809 | 0.04161807353994 | Primary DEG |
| WASHC2A | 1882.3728560092 | 0.129062991322193 | 0.0431025484368816 | 2.99432390897235 | 0,00275054 | 0.0416205673724941 | Primary DEG |
| FAM209B | 9.78244692804444 | 0.420536621538241 | 0.140454543870903 | 2.99411190231604 | 0,00275245 | 0.0416205673724941 | Primary DEG |
| FAM217B | 1178.76485493741 | -0.127786301666134 | 0.0426934160164769 | -2.99311494814134 | 0,00276146 | 0.0416693536571637 | Primary DEG |
| LAMB1 | 26.6756431757844 | 0.301724714700689 | 0.100807856760944 | 2.99306744925844 | 0,00276189 | 0.0416693536571637 | Primary DEG |
| CDK18 | 123.874402197299 | 0.190967177706797 | 0.0638033891063248 | 2.99305695797069 | 0,00276198 | 0.0416693536571637 | Primary DEG |
| ZNF383 | 368.155572886672 | -0.141065908481738 | 0.0471449653397721 | -2.99217334163003 | 0,00276999 | 0.0417583789339796 | Primary DEG |
| KCNK7 | 12.9252164188335 | 0.35593893989438 | 0.118972388573032 | 2.99177770710963 | 0,00277358 | 0.0417807586314399 | Primary DEG |
| SLAMF1 | 1018.48484423315 | -0.162324566411547 | 0.0542633226862862 | -2.9914232740593 | 0,0027768 | 0.04179229341915 | Primary DEG |
| ZNF624 | 351.869033940487 | -0.116143855283686 | 0.0388314598267989 | -2.99097319033912 | 0,0027809 | 0.04179229341915 | Primary DEG |
| OSMR | 10.6379175167622 | 0.509229909192286 | 0.170265334970093 | 2.99080202838548 | 0,00278246 | 0.04179229341915 | Primary DEG |
| CALCOCO1 | 3195.04561130854 | 0.0747418893544924 | 0.0249908789796812 | 2.99076672794347 | 0,00278278 | 0.04179229341915 | Primary DEG |
| FAM8A1 | 1399.68448094845 | -0.113890799755286 | 0.0380983589217765 | -2.98938859779042 | 0,00279536 | 0.0419494992325331 | Primary DEG |
| DTWD2 | 197.852243428675 | -0.163291101198295 | 0.0546350492513078 | -2.98876094075062 | 0,00280111 | 0.042003967051036 | Primary DEG |
| ALG8 | 794.587788156366 | -0.0914859585519047 | 0.030622768513416 | -2.98751429061106 | 0,00281256 | 0.0421437779659329 | Primary DEG |
| GADD45B | 1663.23576182539 | 0.202647310071154 | 0.0678535025614955 | 2.98654162896742 | 0,00282152 | 0.042246144883881 | Primary DEG |
| TBCCD1 | 388.50589455127 | -0.0854061294732646 | 0.0286084967011576 | -2.9853413957892 | 0,00283262 | 0.0423163357352523 | Primary DEG |
| SH2D1B | 1782.58815361985 | 0.256774895043547 | 0.0860122097972755 | 2.98533075302619 | 0,00283272 | 0.0423163357352523 | Primary DEG |
| AC026271.5 | 21.9126772879351 | -0.202108812005102 | 0.0677038627834882 | -2.98518878681163 | 0,00283403 | 0.0423163357352523 | Primary DEG |
| TMCO4 | 660.465771926454 | 0.0874969910303953 | 0.0293135748298501 | 2.9848625266031 | 0,00283706 | 0.0423163357352523 | Primary DEG |
| C7orf26 | 1559.77621292202 | 0.0839573718236019 | 0.0281283001111048 | 2.98480076975772 | 0,00283763 | 0.0423163357352523 | Primary DEG |
| RP4-635A23.6 | 51.2046435916651 | 0.316357765283684 | 0.105994891054713 | 2.98465107266712 | 0,00283902 | 0.0423163357352523 | Primary DEG |
| ABRAXAS2 | 1053.32654204019 | -0.0796786996570089 | 0.0267055553855784 | -2.98360017257073 | 0,00284879 | 0.042430027591715 | Primary DEG |
| DEPDC1 | 14.4854452433993 | -0.387123988725626 | 0.12982545015101 | -2.98188058100573 | 0,00286484 | 0.0426314874507244 | Primary DEG |
| NCAM1-AS1 | 5.29837649482624 | 0.552960198917352 | 0.185459458138148 | 2.98156914976779 | 0,00286775 | 0.0426314874507244 | Primary DEG |
| MEG3 | 22.9541060232454 | 0.826218643302222 | 0.277118710864342 | 2.98146105228773 | 0,00286877 | 0.0426314874507244 | Primary DEG |
| GPN3 | 468.641821035766 | -0.143079309219152 | 0.0480155469289183 | -2.97985378425377 | 0,00288386 | 0.042762875850407 | Primary DEG |
| ASNSD1 | 648.515429336704 | -0.136640715116462 | 0.0458560714306514 | -2.97977368870566 | 0,00288461 | 0.042762875850407 | Primary DEG |
| GRK5 | 1536.88414997938 | 0.109363655007857 | 0.0367028938038115 | 2.97970115360494 | 0,0028853 | 0.042762875850407 | Primary DEG |
| VPS41 | 2379.83771859588 | -0.0970238305153507 | 0.0325656369427166 | -2.97933157843702 | 0,00288878 | 0.042762875850407 | Primary DEG |
| DENND2D | 4761.72846454876 | -0.165115932043233 | 0.0554223546910959 | -2.97922982456321 | 0,00288974 | 0.042762875850407 | Primary DEG |
| HERC4 | 2055.58568048329 | -0.101942299480835 | 0.0342203707084608 | -2.97899459796413 | 0,00289196 | 0.042762875850407 | Primary DEG |
| CAPN15 | 1647.12981612073 | 0.203426908241728 | 0.0682889183286943 | 2.97891536753557 | 0,00289271 | 0.042762875850407 | Primary DEG |
| MTFR1L | 975.957107157781 | 0.109559917064894 | 0.0367875196878259 | 2.97818167668288 | 0,00289964 | 0.0428163637947151 | Primary DEG |
| ZNF232 | 189.825146723446 | -0.168132020329799 | 0.0564565995586379 | -2.97807557742069 | 0,00290065 | 0.0428163637947151 | Primary DEG |
| ZNF433 | 28.9830856359508 | 0.207938026844431 | 0.0698301757657391 | 2.97776748467605 | 0,00290356 | 0.0428275340898257 | Primary DEG |
| VDAC1P8 | 45.9671254152712 | 0.262713251769789 | 0.0882336508764569 | 2.97747230404911 | 0,00290636 | 0.0428369228481348 | Primary DEG |
| YTHDC1 | 1413.77465140176 | -0.103912208104583 | 0.0349022539665769 | -2.97723488586414 | 0,00290861 | 0.0428382616850193 | Primary DEG |
| NUF2 | 48.1845567894517 | -0.25796756178401 | 0.0867078550882369 | -2.97513485394712 | 0,0029286 | 0.0431006099401528 | Primary DEG |
| HELQ | 617.793087821727 | -0.126397672450225 | 0.0424950963151699 | -2.97440607059181 | 0,00293556 | 0.0431710710441449 | Primary DEG |
| KHDC4 | 922.160499677455 | -0.196710784097459 | 0.0661412680874537 | -2.97410058478714 | 0,00293849 | 0.0431820408349791 | Primary DEG |
| DCLRE1B | 639.458974688444 | -0.147390885396459 | 0.0495693050970531 | -2.97343053544686 | 0,00294491 | 0.0432443755357936 | Primary DEG |
| APAF1 | 3291.3321755428 | -0.200270204385426 | 0.0673726930009828 | -2.97257235038095 | 0,00295316 | 0.0432475173110499 | Primary DEG |
| RP11-983P16.4 | 24.1143957804376 | 0.41523928883706 | 0.139693680890777 | 2.97249872856974 | 0,00295386 | 0.0432475173110499 | Primary DEG |
| ZNF407-AS1 | 448.424266247664 | -0.113595756571857 | 0.0382167519314696 | -2.97240740855104 | 0,00295474 | 0.0432475173110499 | Primary DEG |
| C14orf119 | 1495.54565842843 | -0.179439183173668 | 0.0603697358449723 | -2.97233672902698 | 0,00295542 | 0.0432475173110499 | Primary DEG |
| Y_RNA | 127.00700702933 | 0.230021795088157 | 0.0773891755983381 | 2.97227349057195 | 0,00295603 | 0.0432475173110499 | Primary DEG |
| DTNB-AS1 | 3.61741138863584 | -0.609259060064363 | 0.204999896042176 | -2.97199692208144 | 0,0029587 | 0.0432504389139178 | Primary DEG |
| RP11-542H15.1 | 2.833452308237 | 0.678159727500554 | 0.228207105093359 | 2.97168542242854 | 0,0029617 | 0.0432504389139178 | Primary DEG |
| RP11-325K4.2 | 11.3724970574753 | 0.409784366809241 | 0.137919211108079 | 2.97119134830403 | 0,00296647 | 0.0432504389139178 | Primary DEG |
| RP11-19B4.2 | 6.890470471366 | 0.435736276715482 | 0.146657885535591 | 2.97110704360821 | 0,00296728 | 0.0432504389139178 | Primary DEG |
| SLC25A44 | 1332.04025857165 | -0.126168192550716 | 0.0424664478479997 | -2.97100885391474 | 0,00296823 | 0.0432504389139178 | Primary DEG |
| CLEC2B | 1906.96184260316 | 0.148491763085118 | 0.0499850449653384 | 2.9707238072525 | 0,00297099 | 0.0432504389139178 | Primary DEG |
| TNFAIP1 | 588.485389823247 | -0.0832428391551416 | 0.0280215659286059 | -2.97067049597548 | 0,0029715 | 0.0432504389139178 | Primary DEG |
| NEK1 | 475.556426729021 | -0.111424434029889 | 0.037518151381521 | -2.96988070912175 | 0,00297915 | 0.0433299714279832 | Primary DEG |
| RP11-109N23.6 | 5.13891562410031 | 0.490296428936148 | 0.165133326603097 | 2.96909436164023 | 0,00298679 | 0.043375525824548 | Primary DEG |
| RPL17 | 67.4870791368831 | 0.235739225071615 | 0.0794018084356051 | 2.96894025106241 | 0,00298829 | 0.043375525824548 | Primary DEG |
| PSMC6 | 1686.29490112071 | -0.141694573575755 | 0.0477265711526094 | -2.968882326004 | 0,00298885 | 0.043375525824548 | Primary DEG |
| RBM38-AS1 | 35.0931899388641 | 0.250451512277962 | 0.0843713333942141 | 2.96844321646263 | 0,00299312 | 0.043388821073679 | Primary DEG |
| ALG1L6P | 11.6092939179317 | 0.354818285346687 | 0.119534306306125 | 2.96833851562249 | 0,00299414 | 0.043388821073679 | Primary DEG |
| TMEM41A | 481.551683676985 | -0.0882341238935064 | 0.0297294505137148 | -2.96790295040274 | 0,00299839 | 0.0433965425178866 | Primary DEG |
| WDR53 | 328.589303401508 | -0.113085587222523 | 0.0381037340067489 | -2.9678347849713 | 0,00299906 | 0.0433965425178866 | Primary DEG |
| NEPRO | 1854.67957958861 | -0.0782460909790417 | 0.0263731580607474 | -2.96688363216917 | 0,00300835 | 0.0434199538936178 | Primary DEG |
| KMT2E-AS1 | 79.0448273555712 | 0.231193194068953 | 0.0779296629300025 | 2.96669054345345 | 0,00301024 | 0.0434199538936178 | Primary DEG |
| PI4KA | 5512.85864898943 | 0.104903299116763 | 0.0353612262599747 | 2.96661937981215 | 0,00301093 | 0.0434199538936178 | Primary DEG |
| GTSE1 | 38.022596082872 | -0.357369803474319 | 0.12046482294093 | -2.96659053447955 | 0,00301122 | 0.0434199538936178 | Primary DEG |
| TTC5 | 600.083811008047 | -0.105950089997524 | 0.0357149305896727 | -2.96654895440733 | 0,00301162 | 0.0434199538936178 | Primary DEG |
| ESR2 | 26.8663355399222 | 0.336630246728068 | 0.113492740068885 | 2.96609498126266 | 0,00301607 | 0.0434524975389147 | Primary DEG |
| MICAL1 | 2872.37238580799 | 0.145645067027325 | 0.0491138953145187 | 2.96545541938047 | 0,00302235 | 0.0435113255983764 | Primary DEG |
| CRKL | 5308.47880856146 | -0.100094270445954 | 0.0337589323389154 | -2.96497144640356 | 0,00302711 | 0.0435482100932853 | Primary DEG |
| PHC1P1 | 23.7964872965253 | 0.308999991122 | 0.104238919764941 | 2.96434375777104 | 0,00303329 | 0.0436055019361839 | Primary DEG |
| FUT7 | 325.302679604104 | -0.194807414805624 | 0.0657252267538713 | -2.96396717709535 | 0,00303701 | 0.043627262711322 | Primary DEG |
| NEURL1 | 368.30616950619 | 0.220596046518393 | 0.0744321288043345 | 2.96372077571891 | 0,00303944 | 0.0436305920166778 | Primary DEG |
| RP11-463O9.1 | 2.77342967229847 | 0.566279412313456 | 0.191113440789843 | 2.96305382799403 | 0,00304603 | 0.0436936017664634 | Primary DEG |
| RP11-447D11.3 | 67.5741660755279 | -0.212910578366138 | 0.071865872651186 | -2.96261035331107 | 0,00305042 | 0.0437249558956401 | Primary DEG |
| ZNF273 | 240.718459736169 | -0.150713932628147 | 0.0508986809688796 | -2.9610577280047 | 0,00306585 | 0.0438478096460384 | Primary DEG |
| TRIM17 | 11.5498497294276 | 0.368807279984306 | 0.124553571880585 | 2.96103334826799 | 0,00306609 | 0.0438478096460384 | Primary DEG |
| MED8 | 984.86367292217 | -0.117841754377331 | 0.0397998130385606 | -2.96086200865211 | 0,00306779 | 0.0438478096460384 | Primary DEG |
| MTATP6P1 | 16051.6038584844 | 0.205236393697714 | 0.0693165485349787 | 2.96085708298282 | 0,00306784 | 0.0438478096460384 | Primary DEG |
| FAM135A | 419.573081939849 | -0.140289283278106 | 0.0473878371505399 | -2.96044917248366 | 0,00307191 | 0.0438742833899138 | Primary DEG |
| MARCKSL1 | 2036.56372093545 | 0.111815853510524 | 0.0377952841813343 | 2.95846045168105 | 0,0030918 | 0.0441265844040702 | Primary DEG |
| CAB39 | 6847.69602326478 | -0.0668882814318614 | 0.0226113175831582 | -2.9581770803875 | 0,00309464 | 0.044135396589769 | Primary DEG |
| GYPB | 9.83913136045168 | 0.916682138842667 | 0.30990871237465 | 2.95791019174216 | 0,00309732 | 0.0441418697776884 | Primary DEG |
| WDR44 | 1159.64416191698 | -0.100956901103166 | 0.0341398288088499 | -2.95715897312865 | 0,00310488 | 0.0442177727875081 | Primary DEG |
| SMIM13 | 335.666501065199 | -0.0898674152128719 | 0.0303969180847573 | -2.95646469692387 | 0,00311188 | 0.0442856281596467 | Primary DEG |
| TASL | 917.751980962608 | -0.225637346601169 | 0.0763328661419548 | -2.95596586379392 | 0,00311692 | 0.0443254950035987 | Primary DEG |
| RP11-256E16.3 | 9.50959001778252 | 0.373352902916061 | 0.126360459762424 | 2.95466559411082 | 0,00313008 | 0.0444808104100949 | Primary DEG |
| SNRPA | 1355.75080075434 | 0.113880165529731 | 0.0385493610317573 | 2.95413886201423 | 0,00313543 | 0.0445248868568602 | Primary DEG |
| AKAP10 | 1674.61336957042 | -0.0894892931784859 | 0.0303004047356574 | -2.95340256868501 | 0,00314292 | 0.0445752303484599 | Primary DEG |
| RPL32P11 | 3.44303458687989 | 0.576408269305689 | 0.195174584416439 | 2.95329574303497 | 0,00314401 | 0.0445752303484599 | Primary DEG |
| CASS4 | 545.988875158368 | -0.135057221310335 | 0.0457336229705859 | -2.95312753588752 | 0,00314572 | 0.0445752303484599 | Primary DEG |
| CCL25 | 2.95290840483496 | -0.715926566746459 | 0.242455661133426 | -2.95281439665984 | 0,00314891 | 0.0445885982132926 | Primary DEG |
| FAM220A | 523.744264476414 | -0.11148686691753 | 0.0377612532098478 | -2.95241437825096 | 0,003153 | 0.0446145414192716 | Primary DEG |
| IDI2-AS1 | 21.7091171715898 | 0.533928762589697 | 0.180882833284607 | 2.95179345045749 | 0,00315934 | 0.0446563739388668 | Primary DEG |
| BYSL | 211.450935641036 | -0.139898903292338 | 0.0473978751810453 | -2.95158596789345 | 0,00316147 | 0.0446563739388668 | Primary DEG |
| AP1G1 | 2985.49214918294 | -0.156651726250762 | 0.0530759315545468 | -2.95146447104313 | 0,00316271 | 0.0446563739388668 | Primary DEG |
| BAG2 | 185.848372642922 | -0.175471854420831 | 0.0594804987329088 | -2.95007369068591 | 0,00317698 | 0.0448053699765623 | Primary DEG |
| POLR3A | 955.663959031236 | -0.103089785187948 | 0.0349457390689508 | -2.94999584883706 | 0,00317778 | 0.0448053699765623 | Primary DEG |
| NVL | 645.682197711502 | -0.0815594362767789 | 0.0276597595355584 | -2.94866758230233 | 0,00319147 | 0.0449214931401597 | Primary DEG |
| C3orf35 | 11.2536716523349 | 0.420676647114112 | 0.142671051109035 | 2.94857747135131 | 0,0031924 | 0.0449214931401597 | Primary DEG |
| CWF19L2 | 877.504091659108 | -0.146614910390785 | 0.0497246244411983 | -2.94853731000349 | 0,00319282 | 0.0449214931401597 | Primary DEG |
| RABL2A | 181.685435996268 | 0.182377578999464 | 0.0618667880755761 | 2.94790766859712 | 0,00319933 | 0.0449559572042428 | Primary DEG |
| GEMIN5 | 906.474319879233 | -0.173999223958374 | 0.0590293250322384 | -2.94767429346932 | 0,00320174 | 0.0449559572042428 | Primary DEG |
| RP4-647C14.3 | 5.41291539320666 | 0.517526171803046 | 0.175572895470277 | 2.94764274643211 | 0,00320207 | 0.0449559572042428 | Primary DEG |
| GPR137 | 378.838110016891 | 0.101352877041631 | 0.034391879859068 | 2.94700020635563 | 0,00320873 | 0.0450175947647189 | Primary DEG |
| YTHDF2 | 3005.43357334342 | -0.0666842034763755 | 0.0226299955239672 | -2.94671748413521 | 0,00321167 | 0.0450269049056864 | Primary DEG |
| CTD-2013N17.6 | 20.6130738874118 | 0.39984304287983 | 0.13570748752004 | 2.94635948381834 | 0,00321539 | 0.0450471992415417 | Primary DEG |
| CARNMT1 | 560.112228345567 | -0.115382357754083 | 0.0391639746309357 | -2.94613503459229 | 0,00321772 | 0.0450480651295648 | Primary DEG |
| UQCC1 | 565.789426589919 | -0.125831039474462 | 0.0427145839422973 | -2.94585661057698 | 0,00322062 | 0.045049531515989 | Primary DEG |
| YPEL3 | 2827.47787387348 | 0.120419894474596 | 0.0408800521366881 | 2.9456883780861 | 0,00322237 | 0.045049531515989 | Primary DEG |
| DCLRE1C | 1277.90200801744 | -0.09873674872838 | 0.0335228681606483 | -2.94535504107864 | 0,00322584 | 0.0450663176496029 | Primary DEG |
| UMAD1 | 432.864072322727 | -0.114053962646038 | 0.0387437832552767 | -2.94380034842118 | 0,00324209 | 0.0452404606320729 | Primary DEG |
| STK39 | 1217.60031086158 | -0.135507274585349 | 0.0460325740788225 | -2.9437257702191 | 0,00324287 | 0.0452404606320729 | Primary DEG |
| PCYOX1 | 1606.6956401299 | -0.106096792158111 | 0.0360496080781531 | -2.94307754825241 | 0,00324967 | 0.0452936961804351 | Primary DEG |
| SCAT1 | 14.8443510970687 | 0.367994469273392 | 0.125043723607305 | 2.94292635133823 | 0,00325126 | 0.0452936961804351 | Primary DEG |
| LINC00899 | 49.5565599044075 | 0.218657104106608 | 0.0743150809556346 | 2.9422978659896 | 0,00325786 | 0.0453538587684543 | Primary DEG |
| ERF | 674.814573831814 | -0.113922679493712 | 0.0387550541444551 | -2.93955671095382 | 0,00328682 | 0.0456774783440531 | Primary DEG |
| SKA1 | 7.88572646849781 | -0.396044352830093 | 0.134732144590255 | -2.93949416477066 | 0,00328749 | 0.0456774783440531 | Primary DEG |
| RP3-475N16.9 | 34.5489136693906 | 0.186720252729743 | 0.0635223150426721 | 2.93944344761854 | 0,00328802 | 0.0456774783440531 | Primary DEG |
| NKRF | 392.859361999876 | -0.0922099448529639 | 0.0313787740707506 | -2.93860890310933 | 0,00329689 | 0.0457685641055508 | Primary DEG |
| CWF19L1 | 1083.4314764134 | -0.0937639169303948 | 0.0319195033533848 | -2.93751177430058 | 0,00330858 | 0.0458986789445234 | Primary DEG |
| THAP5 | 726.803382800874 | -0.141985318061225 | 0.0483487466804508 | -2.93669076883507 | 0,00331735 | 0.0459881745001668 | Primary DEG |
| MCM10 | 39.1831971955464 | -0.368891699411887 | 0.125638548496303 | -2.93613468021515 | 0,0033233 | 0.0460338487381099 | Primary DEG |
| RP11-755F10.1 | 99.9388033481275 | 0.197536493997274 | 0.0672819828266708 | 2.93594935372461 | 0,00332529 | 0.0460338487381099 | Primary DEG |
| MOB3C | 803.245037511843 | -0.136749831496777 | 0.0465905802466713 | -2.93513905112927 | 0,00333398 | 0.0461031660015335 | Primary DEG |
| ZSCAN26 | 436.073285906728 | -0.139662705831755 | 0.0475862815341441 | -2.9349363162899 | 0,00333616 | 0.0461031660015335 | Primary DEG |
| NFYB | 774.493963683745 | -0.105145395572865 | 0.0358271644949743 | -2.93479534467805 | 0,00333768 | 0.0461031660015335 | Primary DEG |
| CRBN | 2251.27591012775 | -0.0729587183902819 | 0.0248632029663157 | -2.93440545408109 | 0,00334187 | 0.0461031660015335 | Primary DEG |
| RP11-501J20.5 | 127.626549650895 | 0.310834776636587 | 0.105927845605494 | 2.93440100532418 | 0,00334192 | 0.0461031660015335 | Primary DEG |
| CBX8 | 125.691216005447 | -0.163079275359801 | 0.0555909484274157 | -2.93355806966905 | 0,00335101 | 0.0461883422282351 | Primary DEG |
| DIS3 | 1815.17363295349 | -0.076037990410869 | 0.025921485042748 | -2.93339638085828 | 0,00335276 | 0.0461883422282351 | Primary DEG |
| PSMD10P1 | 6.05131421459747 | 0.607932964401407 | 0.207284571956178 | 2.93284231751666 | 0,00335874 | 0.0462032263612146 | Primary DEG |
| KRT72 | 59.0265934428878 | 0.676870768582435 | 0.230798051403262 | 2.9327403956274 | 0,00335985 | 0.0462032263612146 | Primary DEG |
| RNASEL | 999.57731934302 | -0.178782856675573 | 0.0609673582429802 | -2.93243568079577 | 0,00336315 | 0.0462032263612146 | Primary DEG |
| STAM2 | 1068.38581958338 | -0.126334519694853 | 0.0430818022875076 | -2.93243348669017 | 0,00336317 | 0.0462032263612146 | Primary DEG |
| HUS1B | 12.7737895772008 | 0.375796993069275 | 0.128172177116997 | 2.93197011646485 | 0,00336819 | 0.0462032263612146 | Primary DEG |
| ZNF501 | 89.4646676430495 | -0.258099918128447 | 0.0880329548185789 | -2.93185567450675 | 0,00336943 | 0.0462032263612146 | Primary DEG |
| ESCO2 | 33.3232642524509 | -0.318216252427952 | 0.108539937490551 | -2.93178953097936 | 0,00337015 | 0.0462032263612146 | Primary DEG |
| FCER2 | 1054.09164826904 | 0.291830534003662 | 0.0995624813475903 | 2.93112957866959 | 0,00337732 | 0.0462626575645713 | Primary DEG |
| ZFYVE16 | 972.631566250089 | -0.163512807657844 | 0.0557881228986609 | -2.93096091357769 | 0,00337915 | 0.0462626575645713 | Primary DEG |
| MHENCR | 193.469053578711 | 0.267158192372578 | 0.0911791558416703 | 2.93003581691949 | 0,00338923 | 0.046368587203526 | Primary DEG |
| RBM3 | 12407.7507848139 | 0.14608619344561 | 0.0498643522284742 | 2.92967193830686 | 0,0033932 | 0.04639080454597 | Primary DEG |
| NCBP2AS2 | 583.735019128441 | 0.168339185080382 | 0.0574642705745286 | 2.92945831204894 | 0,00339553 | 0.04639080454597 | Primary DEG |
| STK38 | 8768.42900154318 | -0.116853928659928 | 0.0399049629249321 | -2.92830565661092 | 0,00340815 | 0.0464541190692897 | Primary DEG |
| NEK7 | 3022.31946194789 | -0.128991396280177 | 0.0440535303513494 | -2.92806036772546 | 0,00341084 | 0.0464541190692897 | Primary DEG |
| GOSR2 | 1237.51261060555 | -0.0681893268047016 | 0.0232883929997532 | -2.92803916549434 | 0,00341107 | 0.0464541190692897 | Primary DEG |
| ZNF17 | 214.869930924888 | -0.178365792442789 | 0.060918189568859 | -2.92795622629548 | 0,00341198 | 0.0464541190692897 | Primary DEG |
| SMIM24 | 26.5559341137843 | 0.422393732880413 | 0.144269351541067 | 2.92781334613662 | 0,00341355 | 0.0464541190692897 | Primary DEG |
| RP11-439K3.3 | 4.85802122418762 | 0.466596920468805 | 0.159370396174201 | 2.92775152518783 | 0,00341423 | 0.0464541190692897 | Primary DEG |
| RABGAP1 | 1839.86777415589 | -0.0874431277312599 | 0.0298741824719497 | -2.9270467171232 | 0,00342198 | 0.0465068693315359 | Primary DEG |
| PHKG2 | 802.073685868768 | 0.0926532728551373 | 0.0316549909719394 | 2.92697201958674 | 0,0034228 | 0.0465068693315359 | Primary DEG |
| ERI1 | 559.341897767107 | -0.121133910840626 | 0.0413890238146253 | -2.92671582164308 | 0,00342562 | 0.0465133137165256 | Primary DEG |
| IQUB | 6.36969401264005 | 0.333249817420823 | 0.113896316166744 | 2.92590514457857 | 0,00343456 | 0.0466027695060934 | Primary DEG |
| HDHD2 | 38.8535636885456 | 0.160560332418255 | 0.0549077801058298 | 2.92418182102407 | 0,00345363 | 0.0468043202086521 | Primary DEG |
| GALNT11 | 1185.85902060555 | 0.0914970161557828 | 0.0312902684473012 | 2.92413650301151 | 0,00345413 | 0.0468043202086521 | Primary DEG |
| MANBAL | 366.443670834719 | -0.114339917293181 | 0.0391165186955687 | -2.92305964605521 | 0,0034661 | 0.0469124520831002 | Primary DEG |
| TRBV7-3 | 23.6890389599508 | -0.278773073795271 | 0.0953728474833015 | -2.92298155241805 | 0,00346697 | 0.0469124520831002 | Primary DEG |
| MKLN1 | 1590.94470999452 | -0.119312976865788 | 0.0408217394401748 | -2.92278032494534 | 0,00346921 | 0.0469124520831002 | Primary DEG |
| SERINC3 | 6402.95248735116 | -0.0881842142076165 | 0.0301955089136602 | -2.92044139609525 | 0,00349536 | 0.0471930409490332 | Primary DEG |
| RP11-36B6.2 | 13.4970927205026 | 0.365163553919742 | 0.125042824957571 | 2.9203079348507 | 0,00349686 | 0.0471930409490332 | Primary DEG |
| MAN2B2 | 3593.91389490375 | 0.11067191498731 | 0.0378976294496131 | 2.92028595441449 | 0,0034971 | 0.0471930409490332 | Primary DEG |
| ZNF3 | 784.516680554063 | -0.0824612232557491 | 0.0282410124337634 | -2.91991030594792 | 0,00350132 | 0.0472178204266244 | Primary DEG |
| RP11-43F13.1 | 144.20006920179 | 0.234816703646042 | 0.0804518672269528 | 2.91872285553832 | 0,00351469 | 0.0473163296328208 | Primary DEG |
| MAP2K5-DT | 22.0867537873403 | 0.25820968387727 | 0.0884686017673435 | 2.91865903517176 | 0,00351541 | 0.0473163296328208 | Primary DEG |
| DLGAP5 | 79.411271057209 | -0.376766899180219 | 0.129090538657328 | -2.91862519979368 | 0,00351579 | 0.0473163296328208 | Primary DEG |
| ZC3H12A | 536.64394259062 | 0.250693959483559 | 0.0859046012370501 | 2.91828325693265 | 0,00351965 | 0.0473172926027091 | Primary DEG |
| NAA25 | 647.016952463412 | -0.173774001911546 | 0.0595484378209245 | -2.91819581286286 | 0,00352063 | 0.0473172926027091 | Primary DEG |
| THOP1 | 528.545545467378 | 0.112051829442025 | 0.0384028827922966 | 2.91779734474784 | 0,00352513 | 0.0473456984303163 | Primary DEG |
| GRWD1 | 854.671157674628 | -0.129245553582716 | 0.0443055951741768 | -2.91713841275844 | 0,00353259 | 0.0473539725328253 | Primary DEG |
| IPO7 | 4454.65357185469 | -0.094652902735382 | 0.0324479733627667 | -2.91706670481904 | 0,0035334 | 0.0473539725328253 | Primary DEG |
| WDHD1 | 161.744074561048 | -0.156606927020858 | 0.0536882397337759 | -2.91696892648046 | 0,00353451 | 0.0473539725328253 | Primary DEG |
| KIAA0232 | 2405.13446113763 | 0.111046191329227 | 0.0380719368033834 | 2.91674657642735 | 0,00353703 | 0.0473539725328253 | Primary DEG |
| RP11-549B18.1 | 17.0986568320826 | 0.273493591313244 | 0.0937685421678609 | 2.91668810232376 | 0,00353769 | 0.0473539725328253 | Primary DEG |
| XPR1 | 1387.07424666615 | -0.0936207720508905 | 0.032112641098164 | -2.91538686477716 | 0,00355248 | 0.0474713656748038 | Primary DEG |
| XXbac-B444P24.10 | 5.66213704069187 | 0.395591615680464 | 0.135693445977877 | 2.9153332559994 | 0,00355309 | 0.0474713656748038 | Primary DEG |
| ACCS | 184.908386051921 | 0.356679820549379 | 0.122348214718832 | 2.91528422681985 | 0,00355365 | 0.0474713656748038 | Primary DEG |
| GOLPH3-DT | 60.120410713477 | 0.192691976928645 | 0.0661583409216403 | 2.91258780441418 | 0,00358447 | 0.0478509190155642 | Primary DEG |
| LPXN | 4838.72525636791 | -0.0911350790128197 | 0.0312949785142522 | -2.91213106189913 | 0,00358972 | 0.0478833445930633 | Primary DEG |
| ZNF22 | 647.483766183427 | -0.105840527707245 | 0.0363468860757001 | -2.91195585467238 | 0,00359173 | 0.0478833445930633 | Primary DEG |
| IGLV8-61 | 240.920288359562 | -0.461763372607132 | 0.158587920312593 | -2.91171844423554 | 0,00359447 | 0.0478875464733512 | Primary DEG |
| PDS5B | 1999.99690981245 | -0.0900360655232651 | 0.0309381091864823 | -2.91019935900299 | 0,00361198 | 0.0480886145684475 | Primary DEG |
| NUP205 | 2502.7480704991 | -0.0847305118176293 | 0.0291200208888388 | -2.90969955485523 | 0,00361776 | 0.0481024642940634 | Primary DEG |
| SNHG3 | 726.365663595988 | 0.279142570406421 | 0.0959355073911156 | 2.90968983223694 | 0,00361788 | 0.0481024642940634 | Primary DEG |
| LINC02019 | 36.6186651401493 | 0.231012433062527 | 0.0794008611951036 | 2.90944493026195 | 0,00362071 | 0.0481079010936044 | Primary DEG |
| DUOX1 | 25.5203480916712 | 0.302803531263706 | 0.104092984976141 | 2.90897154436596 | 0,0036262 | 0.048127073830777 | Primary DEG |
| BCR | 3269.57625308657 | 0.15725484567107 | 0.054059871319162 | 2.90890159065788 | 0,00362701 | 0.048127073830777 | Primary DEG |
| RP11-151A6.4 | 4.45538784807994 | 0.597885036473309 | 0.205555626003918 | 2.90862891031702 | 0,00363018 | 0.0481326270039039 | Primary DEG |
| CYP2F2P | 8.63795607505153 | 0.320436431100447 | 0.110174400505921 | 2.90844724027543 | 0,00363229 | 0.0481326270039039 | Primary DEG |
| ATP6V1C1 | 1584.61384997181 | -0.10926507665045 | 0.0375750382006619 | -2.90791658193244 | 0,00363845 | 0.0481384987116469 | Primary DEG |
| SEPTIN9 | 29950.8454929133 | 0.080374749174057 | 0.0276406482322559 | 2.90784602801977 | 0,00363928 | 0.0481384987116469 | Primary DEG |
| CDKN2AIP | 1388.6504805557 | -0.112907921397107 | 0.0388295604888587 | -2.9077826268341 | 0,00364001 | 0.0481384987116469 | Primary DEG |
| ASB14 | 22.3357024011005 | 0.266416363083667 | 0.09164310289733 | 2.90710762360523 | 0,00364788 | 0.0482103382817651 | Primary DEG |
| FBXL3 | 2046.89813376765 | -0.146598525411725 | 0.0504428133558704 | -2.90623214009661 | 0,0036581 | 0.0482580281351636 | Primary DEG |
| ZNF175 | 391.405850770837 | -0.161453026951975 | 0.0555563424653229 | -2.9061133218543 | 0,00365949 | 0.0482580281351636 | Primary DEG |
| ZNF330 | 1018.00782438375 | -0.101830685608057 | 0.0350417950911739 | -2.90597799978876 | 0,00366107 | 0.0482580281351636 | Primary DEG |
| PEX11B | 842.390331300802 | -0.132822695750727 | 0.0457069131296006 | -2.90596512991618 | 0,00366122 | 0.0482580281351636 | Primary DEG |
| SCFD1 | 1600.77193263968 | -0.0959062865766311 | 0.0330097673733003 | -2.9053911677731 | 0,00366794 | 0.0483144972840455 | Primary DEG |
| CRK | 80.6894105252031 | -0.154844543821156 | 0.0533036723838076 | -2.90495076410897 | 0,00367311 | 0.0483504036887849 | Primary DEG |
| TFIP11 | 1362.52229735306 | -0.110100100082265 | 0.0379082324597487 | -2.90438495646482 | 0,00367975 | 0.0483563869528722 | Primary DEG |
| MT-ND5 | 121426.862179455 | 0.213114421278427 | 0.0733771145450398 | 2.90437178675941 | 0,00367991 | 0.0483563869528722 | Primary DEG |
| RP5-956O18.2 | 2.77416879676932 | 0.643543931886907 | 0.221583978955663 | 2.9042890867831 | 0,00368088 | 0.0483563869528722 | Primary DEG |
| PPID | 850.288027565973 | -0.0973690160439783 | 0.0335329978491442 | -2.90367763961978 | 0,00368808 | 0.0483881640541343 | Primary DEG |
| AARS1 | 2268.21036885767 | -0.0969053699084872 | 0.0333734248083597 | -2.90366872638775 | 0,00368818 | 0.0483881640541343 | Primary DEG |
| PPP2CA | 4379.67184105481 | -0.0894807109094923 | 0.0308302430510966 | -2.90236800148449 | 0,00370353 | 0.0485282526808 | Primary DEG |
| NBPF1 | 377.501051888822 | 0.207095167523816 | 0.0713543270206006 | 2.90234911001298 | 0,00370376 | 0.0485282526808 | Primary DEG |
| CTD-2252P21.1 | 6.92700514988941 | 0.508846882899584 | 0.175397401623303 | 2.90110844396898 | 0,00371845 | 0.0486072582874504 | Primary DEG |
| ZNF394 | 1374.22179129432 | 0.163996925435303 | 0.0565301180828987 | 2.90105400443016 | 0,0037191 | 0.0486072582874504 | Primary DEG |
| E2F8 | 18.4634899618317 | -0.399909963471689 | 0.137860728369101 | -2.90082584215711 | 0,00372181 | 0.0486072582874504 | Primary DEG |
| SMG8 | 853.416289676204 | -0.15541060212158 | 0.0535747136294224 | -2.90082002484528 | 0,00372188 | 0.0486072582874504 | Primary DEG |
| RP11-372K14.2 | 45.7764788698095 | 0.314426273970344 | 0.10839273548853 | 2.90080578327701 | 0,00372205 | 0.0486072582874504 | Primary DEG |
| RP11-527J8.1 | 12.91580688968 | 0.383963706164406 | 0.132415909643969 | 2.89967955660904 | 0,00373544 | 0.0487383297139678 | Primary DEG |
| RGS1 | 386.686490816027 | 0.442306186965952 | 0.152543095023739 | 2.89954905462695 | 0,003737 | 0.0487383297139678 | Primary DEG |
| RN7SKP23 | 8.9003764922586 | 0.581117841521464 | 0.20049068248283 | 2.89847804558814 | 0,00374979 | 0.0488729453100638 | Primary DEG |
| RP11-248J18.3 | 57.3379522212137 | 0.288112541223111 | 0.0994326897808528 | 2.89756358656398 | 0,00376074 | 0.0489834491101677 | Primary DEG |
| TIGD4 | 4.43919590942508 | -0.40851454331522 | 0.140998358634618 | -2.89729999179522 | 0,0037639 | 0.0489924388354434 | Primary DEG |
| SHARPIN | 982.48645109744 | 0.103753609568432 | 0.0358148457718278 | 2.89694419541645 | 0,00376817 | 0.0490043716024681 | Primary DEG |
| GTPBP6 | 1358.63549371081 | 0.100472403088791 | 0.0346837858061161 | 2.89681188929133 | 0,00376976 | 0.0490043716024681 | Primary DEG |
| XIAP | 1786.57822581242 | -0.124504223730861 | 0.0429998766735536 | -2.89545536783913 | 0,00378609 | 0.0491355702717489 | Primary DEG |
| FZD2 | 341.858299305027 | -0.238858303823639 | 0.0824958214919441 | -2.89539881540502 | 0,00378677 | 0.0491355702717489 | Primary DEG |
| AC000120.7 | 3.2546385401036 | 0.653372324018536 | 0.225662160442766 | 2.89535614981515 | 0,00378729 | 0.0491355702717489 | Primary DEG |
| POC5 | 431.188152389171 | -0.123251027670165 | 0.0425807323351532 | -2.89452578457448 | 0,00379732 | 0.0492334966525738 | Primary DEG |
| WAC-AS1 | 930.50061914991 | 0.146698831438595 | 0.0506907974086739 | 2.89399336640723 | 0,00380376 | 0.0492848218759788 | Primary DEG |
| PAIP1 | 894.405724370017 | -0.0652883960375008 | 0.0225659168457364 | -2.89323037410006 | 0,00381302 | 0.0493425976083377 | Primary DEG |
| HLTF | 389.587114972479 | -0.201106583620841 | 0.0695097213701537 | -2.89321521733495 | 0,0038132 | 0.0493425976083377 | Primary DEG |
| ILRUN | 7018.17521782359 | 0.0984191063086366 | 0.0340238576981002 | 2.89264983359403 | 0,00382007 | 0.0493992518876408 | Primary DEG |
| LEO1 | 701.47111670575 | -0.136464066397199 | 0.0471798849944906 | -2.89242049685231 | 0,00382286 | 0.049403103599905 | Primary DEG |
| TSPYL4 | 865.611183634781 | 0.0961973450671842 | 0.0332609814140284 | 2.89219803437944 | 0,00382557 | 0.0494058931207347 | Primary DEG |
| NCAPG | 66.5141727645526 | -0.339707325346621 | 0.11746734190667 | -2.8919299597034 | 0,00382883 | 0.0494158750513985 | Primary DEG |
| NAA35 | 770.855150157735 | -0.088222933773698 | 0.0305090826856346 | -2.89169407952204 | 0,00383171 | 0.0494208126213438 | Primary DEG |
| BRCA1 | 201.542234973544 | -0.146081890962655 | 0.0505253211955011 | -2.89126100549485 | 0,00383699 | 0.049456795079368 | Primary DEG |
| DERL3 | 60.646644089202 | -0.40179693181853 | 0.139035069668423 | -2.8898962885893 | 0,00385369 | 0.0496354674614301 | Primary DEG |
| MON1B | 2194.09501304581 | -0.0992038344092635 | 0.0343299236195933 | -2.88971905409787 | 0,00385586 | 0.0496354674614301 | Primary DEG |
| BTF3L4P2 | 2.37263493653521 | -0.450065183945098 | 0.155775822037666 | -2.88918509983067 | 0,00386242 | 0.0496875686973544 | Primary DEG |
| PDE4A | 1105.47008146633 | 0.171515139853571 | 0.0593793164997695 | 2.88846605120886 | 0,00387126 | 0.0497277143703641 | Primary DEG |
| DEGS1 | 2797.79053449648 | -0.111285434503465 | 0.0385277142358762 | -2.88845151368566 | 0,00387144 | 0.0497277143703641 | Primary DEG |
| YY1 | 6076.18534554416 | 0.0627661638501651 | 0.0217313630539986 | 2.88827551655193 | 0,0038736 | 0.0497277143703641 | Primary DEG |
| AGAP7P | 11.5261937203942 | -0.385796360401878 | 0.133580638639874 | -2.8881158551874 | 0,00387557 | 0.0497277143703641 | Primary DEG |
| FKBP7 | 38.8421186766026 | -0.159563591727779 | 0.0552650097625103 | -2.88724443211844 | 0,00388632 | 0.049833405086842 | Primary DEG |
| ARL8B | 3085.10493270882 | -0.10741313243709 | 0.0372070558313587 | -2.8869022296185 | 0,00389055 | 0.0498553872767661 | Primary DEG |
| CDIP1 | 233.393490982368 | 0.118951663907177 | 0.0412110812813566 | 2.88639997322734 | 0,00389677 | 0.0499027696215392 | Primary DEG |
| CTD-2649C14.2 | 8.10985011039435 | 0.520818395303981 | 0.180460031090989 | 2.88605954545903 | 0,00390098 | 0.0499245292210274 | Primary DEG |
| RP11-158H5.7 | 72.967734713567 | 0.180579275226055 | 0.0625764837530433 | 2.8857370116657 | 0,00390498 | 0.0499434743517951 | Primary DEG |
